# Functional Overlap of Inborn Errors of Immunity and Metabolism Genes Define T Cell Immunometabolic Vulnerabilities

**DOI:** 10.1101/2023.01.24.525419

**Authors:** Andrew R. Patterson, Gabriel A. Needle, Ayaka Sugiura, Channing Chi, KayLee K. Steiner, Emilie L. Fisher, Gabriella L. Robertson, Caroline Bodnya, Janet G. Markle, Vivian Gama, Jeffrey C. Rathmell

**Affiliations:** Department of Pathology, Microbiology, and Immunology, Vanderbilt University Medical Center, Nashville, TN USA; Department of Cell and Developmental Biology, Vanderbilt University, Nashville, TN USA; Vanderbilt Center for Immunobiology, Vanderbilt University Medical Center, Nashville, TN USA

**Keywords:** Inborn errors of immunity, inborn errors of metabolism, Gfpt1, Bcl11b, T cells, immunometabolism, CRISPR screening

## Abstract

Inborn Errors of Metabolism (IEM) and Immunity (IEI) are Mendelian diseases in which complex phenotypes and patient rarity can limit clinical annotations. Few genes are assigned to both IEM and IEI, but immunometabolic demands suggest functional overlap is underestimated. We applied CRISPR screens to test IEM genes for immunologic roles and IEI genes for metabolic effects and found considerable crossover. Analysis of IEM showed N-linked glycosylation and the *de novo* hexosamine synthesis enzyme, *Gfpt1*, are critical for T cell expansion and function. Interestingly, *Gfpt1*-deficient T_H_1 cells were more affected than T_H_17 cells, which had increased *Nagk* for salvage UDP-GlcNAc synthesis. Screening IEI genes showed the transcription factor *Bcl11b* promotes CD4^+^ T cell mitochondrial activity and *Mcl1* expression necessary to prevent metabolic stress. These data illustrate a high degree of functional overlap of IEM and IEI genes and point to potential immunometabolic mechanisms for a previously unappreciated set of these disorders.

**HIGHLIGHTS:**

- Inborn errors of immunity and metabolism have greater overlap than previously known
- *Gfpt1* deficiency causes an IEM but also selectively regulates T cell subset fate
- Loss of *Bcl11b* causes a T cell deficiency IEI but also harms mitochondrial function
- Many IEM may have immune defects and IEI may be driven by metabolic mechanisms

## INTRODUCTION

Activated T cells increase nutrient uptake and a diverse array of metabolic pathways to support cell growth, proliferation, and effector functions. Pharmacologically targeting immune cell metabolism may prevent immunopathology in inflammatory diseases or enhance tumor immunotherapy(Johnson et al., 2018; Macintyre et al., 2014; Sinclair et al., 2019; Sugiura et al., 2022). Resting naïve T cells primarily utilize oxidative phosphorylation but undergo metabolic reprogramming after activation to primarily rely on high rates of glucose and glutamine uptake for their biosynthetic and energetic needs. Differentiation of T cells into functional subsets further requires appropriate metabolic reprogramming, with each subset utilizing different metabolic pathways to balance its energetic needs with nutrient availability and synthesis of macromolecules for growth and posttranslational modifications(Gerriets et al., 2015; Johnson et al., 2018). Understanding how metabolic pathways regulate T cell function and, conversely, how immunologic signals influence T cell metabolism, will provide a clearer understanding of T cell function and enable development of targeted therapies for immune-mediated disorders.

Forward genetic screens offer the potential to identify novel immunometabolic regulators(Hanna and Doench, 2020). Whole genome-based forward genetic screens are a powerful and unbiased approach to achieve this goal. The number of cells required for whole genome CRISPR screens, however, limits the range of phenotypes that can be tested. In contrast, targeted CRISPR libraries provide greater flexibility, including *in vivo* assays, but are biased by a focus on selected biological processes or pathways. The Inborn Errors of Metabolism (IEM) and Inborn Errors of Immunity (IEI) are human monogenic disorders in which causal gene mutations are identified through unbiased genome-wide approaches. Thus, IEM and IEI provide two gene sets ascertained through unbiased human genetics and known to have severe, deleterious impacts on health when mutated (hereafter referred to as “IEM genes” and “IEI genes”). IEM and IEI genes represent diverse biology of enzymes, kinases, adaptors, and transcription factors and offer an unbiased yet focused opportunity to identify novel immunometabolic regulators. Importantly, the monogenic nature of IEM and IEI disorders is ideally suited to CRISPR screening, where single genes are typically disrupted.

The IEM are comprised of >1500 disorders caused by genetic defects in ∼1400 genes that affect a range of metabolic processes(Ferreira et al., 2019; Lee et al., 2018). Despite the critical role for metabolic reprogramming in immune cell activation and differentiation, only ∼5% of IEM disorders have clinically recognized immunologic phenotypes. While this low frequency of described immune defects in IEM patients may reflect the distinct biology of metabolic and immune processes, it was also possible that the rarity of these patients and the potential for presentation with complicated or severe non-immunologic symptoms can obscure immune phenotypes. In the latter case, the limited clinical overlap of IEM and IEI may reflect a significant underrepresentation of the immunological impacts of IEM. Consistent with this possibility, a variety of IEM associated with mitochondrial dysfunction, but that lacked clinically noted immunological deficiencies, were recently shown to have immune defects when directly assessed(Kapnick et al., 2018; Kruk et al., 2019; Tarasenko et al., 2017).

The IEI are a rapidly growing set of 485 genetic disorders caused by inherited or *de novo* mutations in 450 unique genes that impair immune system development, function, or regulation(Tangye et al., 2022). The remaining IEI are caused by either chromosomal deletions, mutations in the same gene with different pathogenic effects, or are genetically undefined. Notably, 72 (∼16%) of the 450 unique identified IEI genes are also noted as IEM genes, and many others are in pathways that have been described to influence mitochondria and metabolism when activated(Buck et al., 2015; Saravia et al., 2020; Shyer et al., 2020). Understanding how genes canonically considered immunologic may regulate metabolism will provide new mechanistic insight. Recent findings that members of the complement cascade regulate effector T cell differentiation via intracellular regulation of metabolic programs, for example, demonstrate how immunologic factors shape metabolism, which in turn can shape immunity(Arbore et al., 2018; Kolev et al., 2015; West and Kemper, 2019).

We leveraged IEI and IEM to create pooled CRISPR libraries targeting disease relevant genes and test for overlap in immune and metabolic regulatory mechanisms. These are presented in a CRISPR screening data resource to share these and related findings at the Functional ImmunoGenomics reSource (FIGS; https://figs.app.vumc.org/). Screens of IEM genes identified a surprising large number of genes that led to CD4^+^ T cell functional defects when deleted. The N-linked glycosylation pathway was particularly impactful to CD4^+^ T cell function *in vitro* and *in vivo* and *de novo* synthesis of UDP-GlcNAc via Gfat was required for CD4^+^ T cell expansion and T_H_1 polarization. Conversely, metabolic screens of IEI genes identified the transcription factor B-cell leukemia/lymphoma 11B (Bcl11b) as a key regulator of mitochondrial function and nutrient uptake. These data highlight an underappreciated overlap of IEM and IEI gene functions at the interface of immunity and metabolism. They also suggest a set of IEM be considered as candidate IEI and point to metabolic mechanisms for a greater portion of IEI than previously acknowledged.

## RESULTS

### CRISPR Screening of Mendelian disorders of immunity and metabolism identifies metabolic and immunologic regulators in CD4^+^ T cells

We hypothesized that screening IEM genes for effects on T cell function and IEI genes for metabolic alterations would provide an unbiased but disease-relevant approach to identify new immunometabolic regulators of CD4^+^ T cells. Two custom pooled CRISPR libraries were generated comprising of non-targeting control (NTC) sgRNAs and sgRNAs targeting genes mutated in IEM or IEI (**Supplemental Table 1, 2**) to screen for effects on CD4^+^ T cell function or metabolism, respectively (**Fig. 1A, B**). The IEM library included genes compiled in 2019 in the online resource IEMBase(Ferreira et al., 2019) and encompassed 1038 genes spanning a diverse range of metabolic and biological processes. The IEI library was designed from pathogenic gene variants identified by the International Union of Immunological Scientists in their 2017 biennial report on the classification of Inborn Errors of Immunity(Picard et al., 2018). This included sgRNAs for 318 genes causing primary immune deficiencies. Because our CRISPR approach relies on gene disruption, genes associated with gain-of-function variants, multigenic disorders, or those without a mouse homolog were excluded. Notably, sgRNAs for 43 genes were present in both libraries. An online searchable portal, the Functional ImmunoGenomics resource (FIGS; https://figs.app.vumc.org/), has been created to make available all library information and CRISPR screening results.

**Figure 1.**
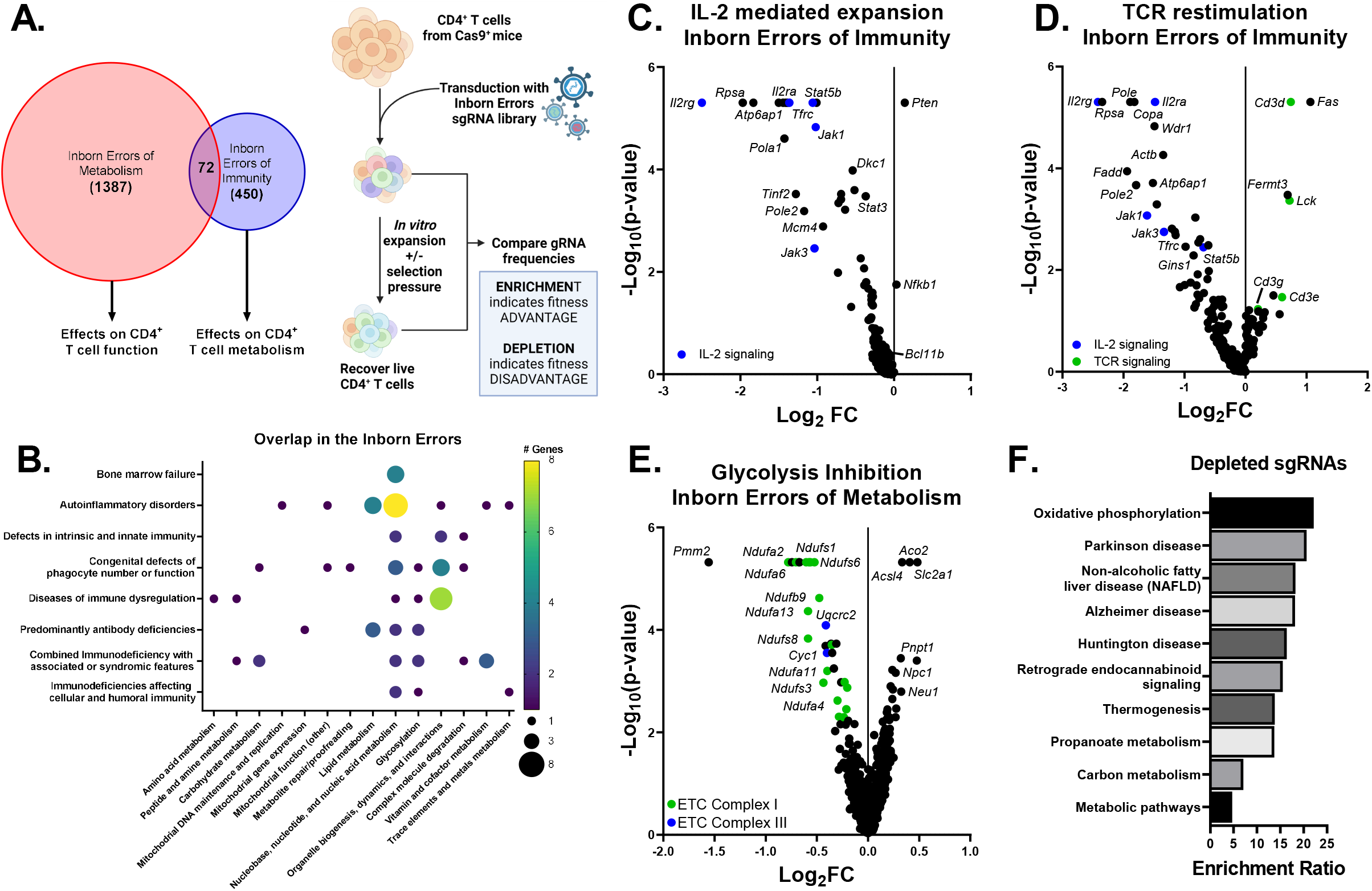
Screening the Inborn Errors identifies novel immunometabolic regulators. **(A)** Workflow for pooled CRISPR screens targeting the Inborn Errors of Metabolism and Inborn Errors of Immunity for changes in CD4^+^ T cell function and metabolism, respectively. **(B)** Overlap between in the Inborn Errors covers a range of immunological phenotypes and metabolic pathways. **(C and D)** Change in sgRNA abundance from IEI-targeted *in vitro* CRISPR screens after **(C)** IL-2 mediated expansion alone or **(D)** with restimulation with αCD3/αCD28. sgRNAs marked in blue are associated with IL-2 signaling; sgRNAs in green are associated with TCR signaling. Representative of 3 independent experiments. **(E)** Comparison of sgRNA abundance samples treated 2 days with 2mM 2-DG relative to vehicle treated samples from a screen of IEM genes. Genes associated with the electron transport chain are in marked in green (C1) or blue (C3). **(F)** Over enrichment analysis significantly and trending (-log10(p-value)>0.1) depleted genes in the 2-DG treated sample relative to vehicle treated cells generated using WebGestalt.

To validate the CRISPR libraries and this approach, the IEI library was first screened for effects on T cell proliferation following activation and culture in the T cell growth factor IL-2. Briefly, CD4^+^ T cells isolated from Cas9 transgenic mice were activated and retrovirally transduced with the IEI library. The following day, DNA was collected to represent guide abundance in the initially transduced, or unselected, population. Transduced T cells were then expanded *in vitro* with IL-2 for an additional 4 days at which time DNA was again collected to reflect guide abundance in the selected T cell population. The frequencies of sgRNA were normalized to NTC and changes for each guide were determined by comparing unselected to selected CD4^+^ T cell samples. As expected, sgRNAs targeting genes associated with IL-2 signaling (*Il2rg, Il2ra, Stat5b, Jak1, Jak3*) were depleted, indicating that loss of those genes decreased T cell proliferation or fitness in IL-2 cultures (**Fig. 1C**). Notably, patients with loss-of-function or autosomal dominant negative mutations in *IL2RG, JAK3*, and *STAT5B* present with T cell lymphopenia(Tangye et al., 2022). This CRISPR library was further validated by restimulating the IL-2 expanded cells 4 days post transduction with αCD3 and αCD28 for 24 hours prior to isolation to promote activation-induced cell death (AICD). As observed without restimulation, sgRNAs for *Il2rg, Il2ra, Jak3*, and *Jak1* were all highly depleted. However, sgRNAs targeting TCR signaling pathway genes (*Cd3d, Lck, Cd3g*, and *Cd3e*) and *Fas* were all enriched, consistent with T cell escape from AICD (**Fig. 1D**). Notably, these sgRNAs were enriched only in the context of TCR restimulation, supporting the specificity of these screens (**Fig. S1A**).

The IEM Library was next validated by screening to identify metabolic genes that modify the response of activated T cells to the glycolytic inhibitor 2-deoxyglucose (2-DG). Cas9 transgenic CD4^+^ T cells were transduced, DNA was collected the following day as an unselected sample and T cells were expanded in IL-2 for 48 hours, and then selected by culture with 2-DG or vehicle for an additional 48 hours in IL-2 prior to determination of sgRNA abundance in each group. sgRNAs for 101 genes were differentially abundant in 2-DG-treated samples but not vehicle-treated samples (**Fig. 1E, F, S1B**). Among sgRNAs depleted in 2-DG treated samples relative to vehicle-treated samples, genes associated with electron transport and oxidative phosphorylation were particularly prevalent, indicating that these pathways were conditionally essential or generated synthetic lethality upon glycolysis inhibition. Notably, sgRNAs for *Slc2a1* (Glut1) were enriched in 2-DG-treated samples but depleted in vehicle-treated samples, consistent with adaptation of Glut1-deficient T cells to low glucose uptake and resistance to 2-DG treatment. Together, these findings validated our screening approach to use IEM and IEI genes to identify immunometabolic regulators.

### N-linked glycosylation is critical for CD4^+^ T cell expansion and survival *in vitro* and *in vivo*

The IEM library was next examined in a pair of focused screens for effects on T cell expansion and survival after expansion in IL-2 with or without additional TCR restimulation (**Fig. 2A-D**). Gene set analyses of depleted sgRNA showed widespread pathways were critical, with N-Glycan biosynthesis most important. This approach identified 313 IEM genes not previously associated with IEI as potential modifiers of normal T cell expansion and survival (**Fig. 2E, Supplemental Table 3**). When genes identified that regulate T cell expansion during glycolysis inhibition are also included, 343 IEM genes are shown to be necessary for T cell expansion. Of these, sgRNAs for *Mat2a* (methionine adenosyltransferase 2A) were consistently the most depleted in each condition. Among other strongly depleted genes were those encoding parts of the V-ATPase complex, including *Atp6ap2, Atp6ap1, Atp6v1a*, and *Atp6v1e1* (**Fig. S2A**). Notably, mutations in X-linked *ATP6AP1* also cause an IEI and was consistently highly depleted in screens using that library (**Fig. 1C, D**). The V-ATPase complex is a proton pump that regulates a range of cellular functions, including organelle trafficking, pH, and membrane fusion. Clinically, variants of *ATP6AP1* and X-linked *ATP6AP2* have been shown to cause reduced T cell numbers, although autosomal recessive mutations in *ATP6V1E1* and *ATP6V1A* have no published immunological consequences(Jansen et al., 2016; Rujano et al., 2017; Van Damme et al., 2017). Consistent with a role for the V-ATPase in activated T cells, pharmacologic inhibition of this complex reduced T cell expansion and survival (**Fig. S2B, C**).

**Figure 2.**
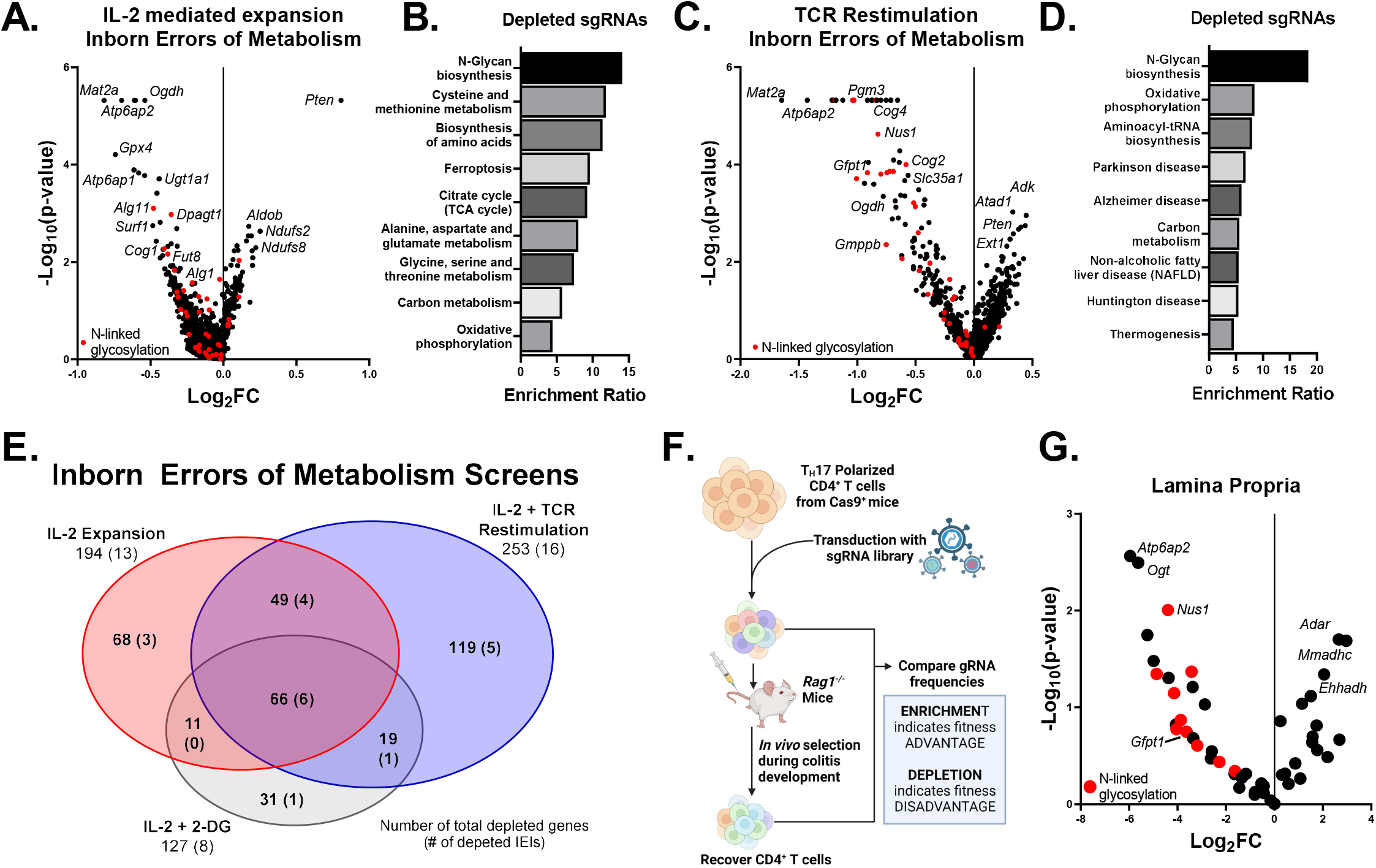
N-linked glycosylation is critical for CD4^+^ T cell expansion *in vitro* and *in vivo*. **(A)** Change in sgRNA abundance from IEM-targeted *in vitro* pooled CRISPR screens after IL-2 mediated expansion. Genes associated with N-linked glycosylation are marked in red. Representative of 3 independent screens. **(B)** Over enrichment analysis of significantly and trending (-log10(p-value)>0.1) depleted genes in Fig. 2A generated using WebGestalt. **(C)** Change in sgRNA abundance from IEM-targeted *in vitro* pooled CRISPR screens after IL-2 mediated expansion and restimulation with αCD3/αCD28. Genes associated with N-linked glycosylation are marked in red. Representative of 2 independent screens. **(D)** Over enrichment analysis of significantly and trending (-log10(p-value)>0.1) depleted genes in Fig. 2C generated using WebGestalt. **(E)** Comparison sgRNAs depleted in screens of IEM genes after expansion and treatment with 2-DG, expansion with IL-2 alone, or expansion and restimulation with αCD3/αCD28 across all biological replicates. Numbers shown describe number of total genes depleted and (number of IEI genes depleted). **(F)** Schematic of *in vivo* screen using a model of colitis. **(G)** Screen for CD4^+^ T cells isolated from the lamina propria after colitis development. Screen represents pooled populations from 5 mice. Genes associated with N-linked glycosylation are marked in red.

Analysis of sgRNAs depleted in the screens of the IEM Library identified the N-Glycan biosynthesis pathway as critical for T cell expansion and survival. T cells rendered deficient in *Pgm3, Gfpt1, Cog4, Nus1*, and *Alg11* were all depleted (**Fig. 2A-D**). Disruption of this pathway was particularly notable when cells were restimulated with anti-CD3 and anti-CD28 after IL-2 expansion (**Fig. 2C, D**). To confirm the *in vivo* impact of this pathway, we generated a targeted IEM library of 48 genes focused on key hits from the *in vitro* screen and the N-Glycan biosynthesis pathway (**Supplemental Table 4**). This targeted IEM library was then tested in an *in vivo* colitis screen. T_H_17-polarized Cas9 transgenic CD4^+^ T cells were transduced with the focused IEM library and a sample of DNA was collected to reflect the unselected population prior to transfer into *Rag1*^*-/-*^ mice (**Fig. 2F**). After colitis was apparent, sgRNA abundance was determined in CD4^+^ T cells from the lamina propria, mesenteric lymph node, and spleen and compared to input to establish the relative depletion or enrichment of each knockout (**Fig. 2G, Fig. S3A, B**). Several sgRNA, including sgRNAs targeting adenosine deaminase, *Adar*, were consistently enriched to indicate that loss of these genes increased T cell fitness. Notably, sgRNAs for *Atp6ap2, Gfpt1*, and N-linked glycosylation genes were consistently depleted, confirming this importance of the N-Glycan biosynthesis pathway in CD4^+^ T cell expansion and persistence *in vitro* and *in vivo*. In addition, O-linked N-Acetylglucosamine (GlcNac) Transferase, *Ogt*, which mediates O-GlcNacylation as a hexosamine-dependent post-translational modification that has been shown to influence T cells (Chang et al., 2020; Golks et al., 2007), was also highly depleted.

### CD4^+^ T cell subsets differentially require *Gfpt1* for expansion and polarization

To further interrogate the importance of N-linked glycosylation in CD4^+^ T cells, we focused on *Gfpt1* (Gfat), a gene highly depleted in both *in vitro* T cell expansion and *in vivo* colitis screens. Patients with pathogenic autosomal recessive mutations in *GFPT1* present with a neuromuscular disorder characterized by early onset proximal muscle weakness(Bauche et al., 2017; Senderek et al., 2011; Zoltowska et al., 2013). However, while autoimmune conditions such as myasthenia gravis and Lamber-Eaton myasthenic syndrome can present similarly, no clinical immunological phenotype was noted in the 13 unrelated families with pathogenic *GFPT1* variants. In contrast, patients with defects in PGM3 function, an enzyme downstream of GFAT in the hexosamine biosynthesis pathway (HBP), are described to develop Combined Immunodeficiency associated with hyper IgE syndrome.

Gfat is the entry and rate-limiting enzyme of the HBP for the *de novo* synthesis of UDP-GlcNAc, which is used as a post-translational modification as the primarily building block for N-linked glycosylation of proteins (**Fig. S4A**). Notably, UDP-GlcNAc can also be obtained through a hexosamine salvage pathway, in which hyaluronic acid (HA) is taken up via CD44, broken down, and shunted into the HBP by Nagk as GlcNAc-6P. Interestingly, Gfat was more highly expressed in T_H_1-polarized cells, while Nagk was more highly expressed in T_H_17 and iTreg cells (**Fig. 3A, Fig. S4B, C**), suggesting potential differential usage of these pathways in CD4^+^ T cell subsets. To test the requirement of IEM genes in CD4^+^ T cell subsets, the targeted IEM library was used to screen T_H_1, T_H_17, and Treg T cells. When these IEM genes were screened in T_H_1, T_H_17, and iTreg-polarizing conditions, *Gfpt1* and N-linked glycosylation genes were depleted in all conditions (**Fig. S5A**). However, *Gfpt1* and genes in the hexosamine biosynthesis pathway were more depleted in T_H_1-polarizing conditions than in T_H_17 or Treg, with *Gfpt1* deficient T cells enriched in T_H_17 relative to levels observed in T_H_1 (**Fig. S5B**) and depletion of *Gfpt1* and *Pgm3* sgRNAs only reaching significance in T_H_1 conditions (**Fig. S5C**).

**Figure 3.**
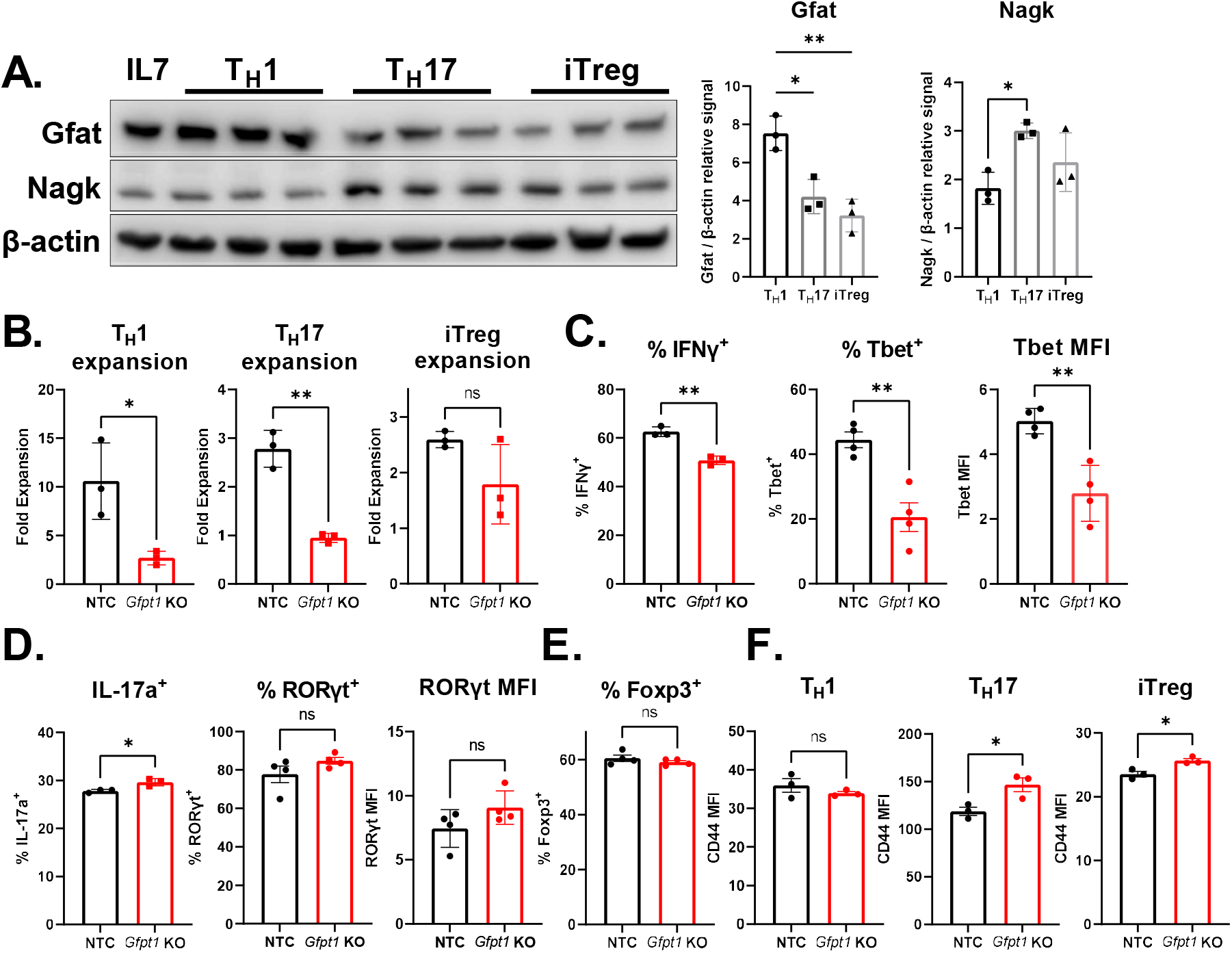
*Gfpt1* is required for T cell expansion and T_H_1 polarization. **(A)** Expression of Gfat and Nagk in polarized CD4^+^ T cells 3 days post activation (*n*=3; mean ± SD). One-way ANOVA with Sidak’s multiple comparisons test. **(B)** Expansion of T_H_1, T_H_17, and iTreg polarized WT and *Gfpt1* KO cells over 4 days post gene deletion. **(C)** Expression of IFNγ and Tbet in T_H_1-polarized cells, **(D)** IL-17a and RORγt in T_H_17-polarized cells, and **(E)** Foxp3 in iTreg-polarized cells (*n*=3-4; mean ± SD). **(F)** Expression of CD44 in polarized WT and *Gfpt1* KO cells (*n*=3; mean ± SD). Student’s unpaired two-tailed t-test, * p<0.05, ** p<0.005.

To directly test for differential roles in CD4 T cell subsets, *Gfpt1* was disrupted as a single gene and compared to NTC. While CRISPR knockout of *Gfpt1* blocked the expansion of both T_H_1 and T_H_17 cells (**Fig. 3B**), only T_H_1 cell polarization was affected, consistent with greater dependence of T_H_1 on *Gfpt1. Gftp1*-deficiency reduced IFNγ and Tbet expression, yet T_H_17 and iTreg polarized cells expressed IL-17a, RORγt, and Foxp3 comparable to WT cells (**Fig. 3C-E**). Interestingly, despite reduced expansion, Gfat-deficient T_H_17 and iTreg cells upregulated CD44, which may allow increased HA uptake to support salvage UDP-GlcNAc synthesis (**Fig. 3F**).

### Gfat is critical for CD4^+^ T cell persistence and pathogenicity *in vivo*

We next sought to determine if Gfat was essential for CD4^+^ T cells *in vivo* using a colitis model. To determine the importance of *Gfpt1* in a cell intrinsic fashion independent of disease development or inflammatory signals differing between mice, T_H_17-polarized Cas9-transgenic T cells were transduced with a *Gfpt1* sgRNA (BFP) or NTC (GFP), mixed at a 1:1 ratio, and adoptively transferred into *Rag1*^*-/-*^ KO mice (**Fig. 4A**). After onset of colitis, the NTC and *Gfpt1* KO CD4^+^ T cells in the spleen, mesenteric lymph node, and lamina propria were characterized. *Gfpt1* KO cells exhibited reduced recruitment to each site, consistent with *de novo* synthesis of UDP-GlcNAc as important for CD4^+^ T cells *in vivo* (**Fig. 4B**). Similar to *in vitro* studies, *Gfpt1*-deficient cells expressed higher levels of CD44, which may compensate in part to support HA uptake and salvage UDP-GlcNAc synthesis (**Fig. 4C**). As previously observed in this model, adoptively transferred T_H_17-polarized cells adopt a T_H_1 or T_H_1* phenotype become IFNγ^+^ or IFNγ^+^IL-17a^+^, respectively(Harbour et al., 2015). In contrast, *Gfpt1* KO cells isolated from each site exhibited both reduced IL-17a and IFNγ production (**Fig. 4D-G**). Notably, the reduction of IFNγ-producing cells was more marked than that observed for IL-17a-producing cells (**Fig. 4H**). Furthermore, *Gfpt1* KO cells retained a higher expression of Il23r, consistent with a T_H_17 phenotype (**Fig. 4I**). These data show that the ability of *Gfpt1-*deficient cells to adopt a T_H_1 or T_H_1* phenotype was impaired and suggest T_H_17 cells are less dependent on Gfat, possibly due to increased capacity for salvage UDP-GlcNAc synthesis.

**Figure 4.**
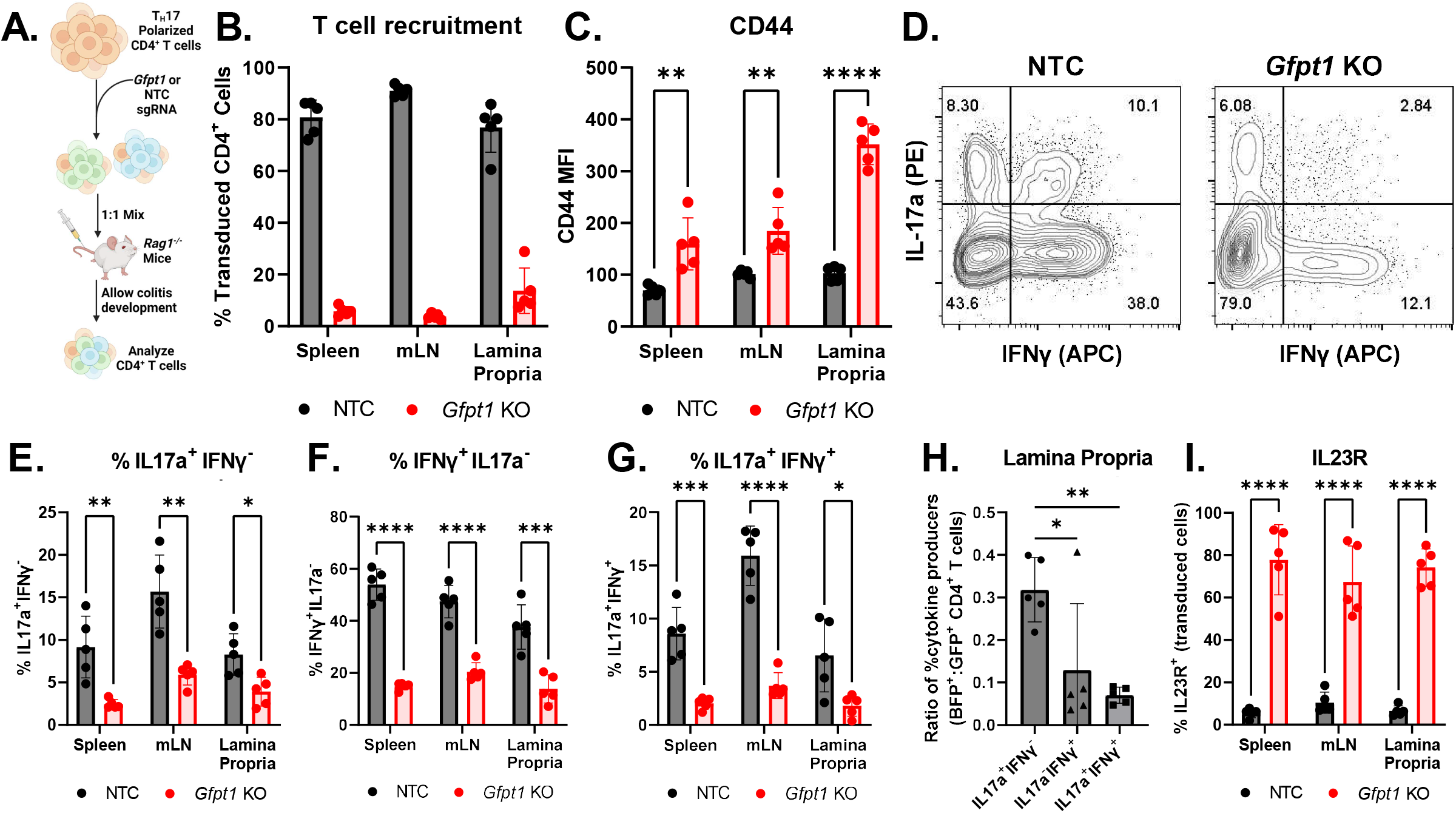
Impaired expansion of *Gfpt1*-deficient CD4^+^ T cells during colitis. **(A)** Schematic of colitis induction via co-transfer of WT and *Gfpt1* KO T_H_17 cells into *Rag1*^*-/-*^ mice. **(B)** Percentage of CD4^+^ T cells that are either WT or *Gfpt1* KOs in the spleen, mLN, or lamina propria of mice (*n*=5; mean ± SD). Expression of **(C)** CD44 and (**D-G)** IL-17a and IFNγ by transduced cells. Student’s unpaired two-tailed t-test. (**H)** Ratio of cytokine production by *Gfpt1* KO and WT cells (*Gfpt1* KO:WT). One-way ANOVA with Sidak’s multiple comparisons test. (**I)** Expression of IL23R by transduced cells. Representative of two independent experiments. * p<0.05, ** p<0.005, *** p<0.0005, **** p<0.00005.

### Immunoregulator *Bcl11b* regulates metabolic function of CD4^+^ T cells

We next screened genes associated with IEI to identify immunologic regulators of CD4^+^ T cell metabolism. CD4^+^ T cells retrovirally transduced with a pooled IEI CRISPR library were expanded in IL-2 and stained for different metabolic readouts measuring mitochondrial features or nutrient uptake (**Fig. 5A**). The top and bottom quintile for each readout was isolated by cell sorting and sgRNA abundances in the top and bottom quintiles were compared. An “enriched” sgRNA would suggest increased signal, while a “depleted” sgRNA would indicate a reduced signal for that readout. To validate this approach, we first tested our IEM library to validate if sorting for mitochondrial membrane potential led to expected gene depletions. Notably, the most “depleted” sgRNAs targeted genes involved in the electron transport chain, including *Ndufa13, Ndufaf4, Atp5e, Ndufb8*, and *Cyc1* (**Fig 5B**). Targeting these genes is consistent with a reduced mitochondrial membrane potential, confirming this method to identify IEI regulators of mitochondria.

**Figure 5.**
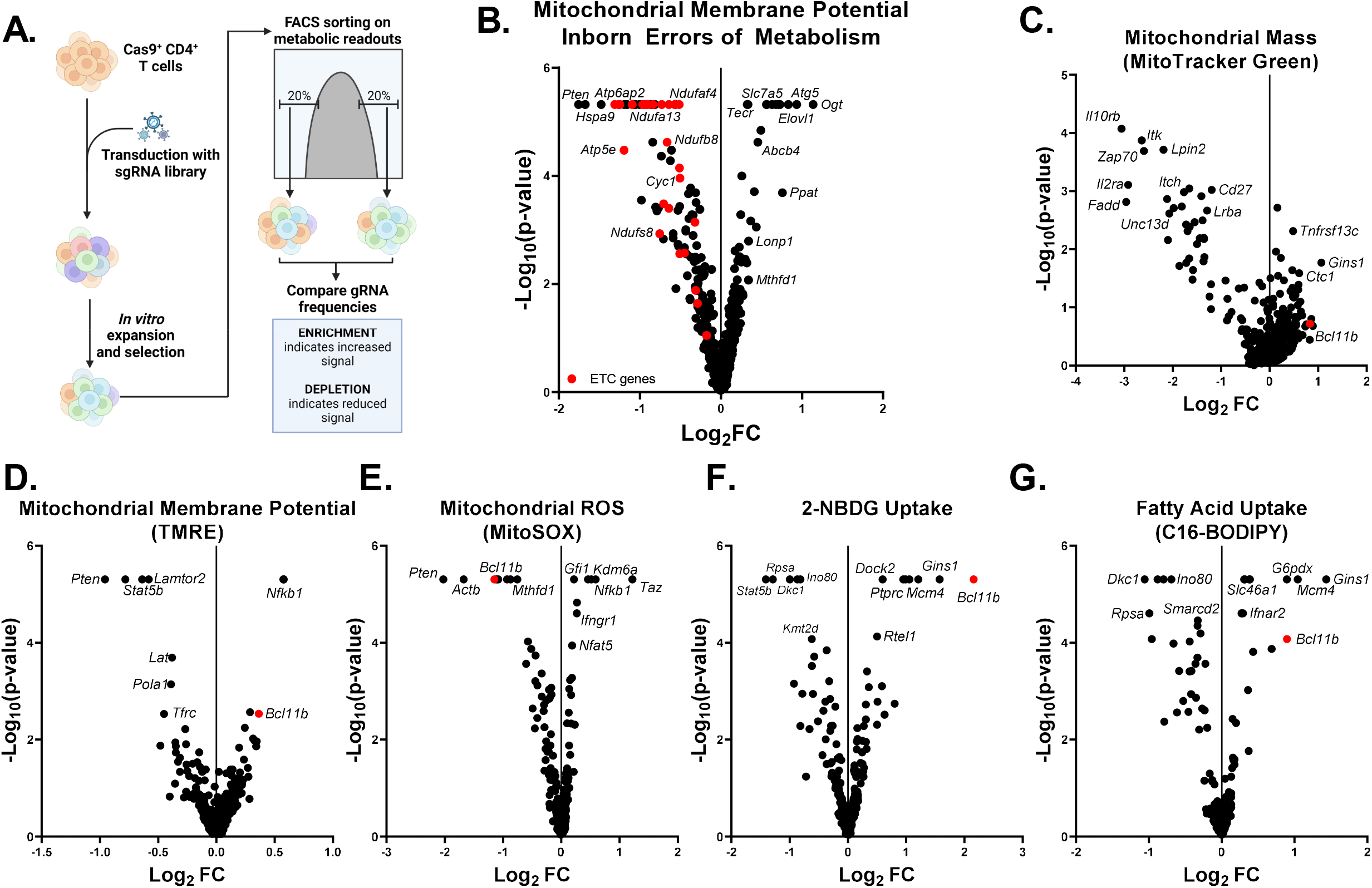
*In vitro* CD4^+^ T cell screens identify *Bcl11b* as a key metabolic regulator in CD4^+^ T cells. **(A)** Schematic of CD4^+^ T cell screen for metabolic readouts. Activated Cas9 Tg CD4^+^ T cells were transduced with a pooled CRISPR library and expanded in IL-2. Cells were stained with for metabolic readouts associated with mitochondria function or nutrient uptake and the sgRNA abundance in the top and bottom quintile determined. Enrichment relative to the bottom quintile indicates that a loss of that gene promotes increased signal, while depletion is associated with reduced signal. **(B)** Screen of the IEM library for effects on mitochondrial membrane potential. Genes associated with electron transport chain marked in red. (**C-G**) Depletion-Enrichment plots of CD4^+^ T cells transduced with the IEI library after 7 days expansion in IL-2 and selection for the top and bottom quintiles for **(C)** mitochondrial mass (MTG), **(D)** mitochondrial membrane potential (TMRE), **(E)** mitochondrial ROS (MitoSOX), **(F)** 2-NDBG uptake, and **(G)** C16-BODIPY uptake.

Next, we applied this approach to screening the IEI genes for effects on mitochondria by staining for mitochondrial mass (MitotrackerGreen), mitochondrial membrane potential (TMRE), or mitochondrial reactive oxygen species (MitoSOX) (**Fig. 5C-E**). Of note, sgRNAs for *Pten* were depleted in mitochondrial potential and reactive oxygen species (mROS) screens, despite consistently providing an advantage for survival and expansion (**Fig. 1C, E, 2A**). *Taz* sgRNAs were enriched in the mitochondrial ROS screen, consistent with Tafazzin’s role to regulate mitochondrial function via cardiolipin synthesis(Liu et al., 2021; Paradies et al., 2014) (**Fig. 5E**). We further screened the IEI for changes in nutrient uptake using 2-NBDG and C16-BODIPY (fatty acid uptake). Notably, several sgRNAs were differentially abundant across multiple screens, even after accounting for significant effects on expansion or survival (**Fig. 5F, G**). Among differentially abundant sgRNAs, *Bcl11b* was one of the most significant hits in each screen. Furthermore, *Bcl11b* deficiency did not affect expansion or survival in initial T cell activation screens (**Fig 1C**), suggesting the finding was specific to the metabolic selection. These data demonstrate that Bcl11b is a novel regulator of metabolic activity in activated CD4^+^ T cells.

### Bcl11b-deficient thymocytes exhibit altered mitochondria

Patients with autosomal dominant-negative *BCL11B* mutations patients develop combined immunodeficiency with syndromic features characterized by low peripheral T cell numbers, congenital abnormalities, and neurocognitive deficits(Alfei et al., 2022; Che et al., 2022; Lessel et al., 2018; Punwani et al., 2016; Tangye et al., 2022; Yang et al., 2020). Molecularly, the mutant protein was shown to form nonfunctional heterodimers with WT BCL11B, functionally resulting in BCL11B-deficiency(Lessel et al., 2018; Punwani et al., 2016). Several different Bcl11b-deficient mouse models have demonstrated a clear role of Bcl11b in guiding T cell specification during thymic development and promoting regulatory T cell function in the periphery(Albu et al., 2007; Avram and Califano, 2014; Drashansky et al., 2019; Hasan et al., 2019; Lessel et al., 2018; Punwani et al., 2016). However, Bcl11b has not been previously shown to regulate metabolic function of T cells either in patients or mouse models.

Based on our CRISPR screens pointing to Bcl11b as a mitochondrial regulator, we asked if mitochondria were dysregulated in thymic development of *Bcl11b*^*-/-*^ T cells. A conditional Bcl11b allele was crossed to the constitutive CD4-cre mouse (*Bcl11b*^*fl/fl*^*Cd4*^*cre*^, cKO) and, consistent with reports of compromised T cell development in *BCL11B* patients, CD4 and CD8 SP thymocytes were decreased but not completely absent in cKO mice (**Fig. S6A**)(Albu et al., 2007; Lessel et al., 2018; Punwani et al., 2016). Strikingly, the remaining CD4 SP cKO cells exhibit significantly altered mitochondrial mass, membrane potential, and mROS consistent with Bcl11b as a metabolic regulator (**Fig. S6B-D**). These data demonstrate that loss of Bcl11b leads to mitochondrial dysfunction *in vivo*, which may contribute to the impaired thymic development of T cells in these mice and patients.

### Bcl11b-deficiency rewires metabolic programing of CD4^+^ T cells

Because altered thymic development and lymphopenia-induced T cell proliferation resulting in a memory-like phenotype in *Bcl11b*^*fl/fl*^*Cd4*^*cre*^ mice(Albu et al., 2007) that may complicate evaluation of the role of Bcl11b in peripheral effector T cell metabolism, we generated tamoxifen-inducible conditional knockout mice (*Bcl11b*^*fl/fl*^*Cd4*^*cre-ert2*^, KO). Treatment with tamoxifen induced robust reduction in Bcl11b protein levels observed in resting and stimulated CD4^+^ T cells (**Fig. S7A-B**). To determine how loss of Bcl11b may rewire the metabolic function of CD4^+^ T cells, we first performed RNAseq on resting T cells or T cells activated *in vitro* for 3 days. While few differences were observed in resting T cells, gene expression in activated Bcl11b-deficient T cells was markedly different (**Fig. S7C**). Among the most differentially expressed genes with *Bcl11b* deficiency were *Tox, Irf7*, and *Ctla4* (**Fig. S7D**). Interestingly, gene set enrichment analysis showed that *Bcl11b* KO T cells predominantly upregulated genes associated with hypoxia and glycolysis (**Fig. S7E-G**). However, Bcl11b-deficient CD4^+^ T cells did not exhibit increased glycolytic activity, and instead had a reduced glycolytic reserve compared to WT littermates, suggesting these increased gene signatures may be compensatory (**Fig. S8A-D**).

We next asked if Bcl11b-deficiency affected the mitochondrial function of CD4^+^ T cells. Notably, *ex vivo* resting Bcl11b KO T cells exhibited only a slight increase in mitochondrial mass, showed no changes in mitochondrial potential or level of mROS (**Fig. S9A-C**), and were not proliferating at increased levels *in vivo* (**Fig. S9D**). Significant changes were observed, however, following activation. Bcl11b KO CD4^+^ T cells activated for 3 days *in vitro* exhibited a reduction in mitochondrial mass, potential, and mROS (**Fig. 6A-C**). This corresponded to reduced basal and maximal mitochondrial respiration (**Fig. 6D-F**) and with negligible spare respiratory capacity (**Fig. 6G**), suggesting a potentially profound mitochondrial defect in activated Bcl11b-deficient T cells. Notably, Bcl11b KO T cells stimulated for 3 days exhibited enhanced proliferation and viability compared to WT controls, indicating reduced mitochondrial function was not a result of poor viability (**Fig. S9E**). This correlated with increased expression of several pro-survival factors and reduced expression of pro-apoptotic factors (**Fig. S9F**). Interestingly, both *Tox* and *Tcf7* expression were notably reduced in Bcl11b KO cells. While Bcl11b-deficient T cells initially survived and expanded well, WT T cell expansion surpassed that of Bcl11b KO cells after culture in IL-2 for an additional 2 days, suggesting the metabolic deficiencies ultimately impaired sustained expansion (**Fig. S9G**). Consistent with this metabolic defect, activated Bcl11b KO CD4^+^ T cells exhibited increased AMPK phosphorylation, consistent with metabolic stress (**Fig. 7A**), that corresponded with reduced mTOR activity and cMyc levels (**Fig. 7A-B**).

**Figure 6.**
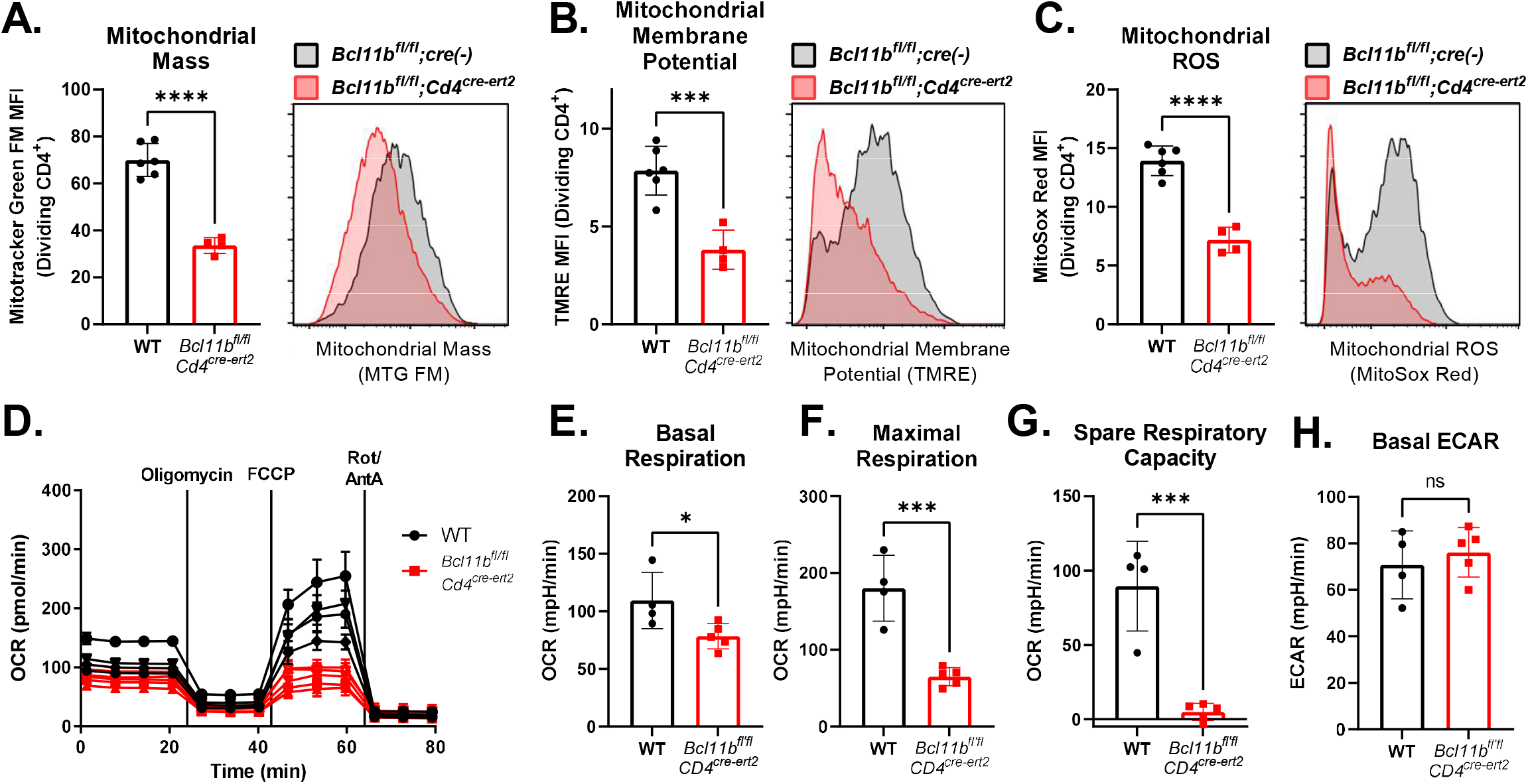
Bcl11b-deficient CD4^+^ T cells exhibit impaired mitochondrial function. **(A)** Mitochondrial mass, **(B)** membrane potential, and **(C)** reactive oxygen species in dividing CD4^+^ T cells stimulated 3 days with αCD3/αCD28 (*n*=4-6; mean ± SD). **(D)** Seahorse XP Cell Mito stress test performed on WT or Bcl11b-deficient CD4^+^ T cells stimulated 3 days with αCD3/αCD28 (*n*=4-5). **(E)** Basal OCR, **(F)** maximal OCR, **(G)** SRC, and **(H)** basal ECAR measured in **(D)** (mean ± SD). Student’s unpaired two-tailed t-test, * p<0.05, *** p<0.0005, **** p<0.00005.

**Figure 7.**
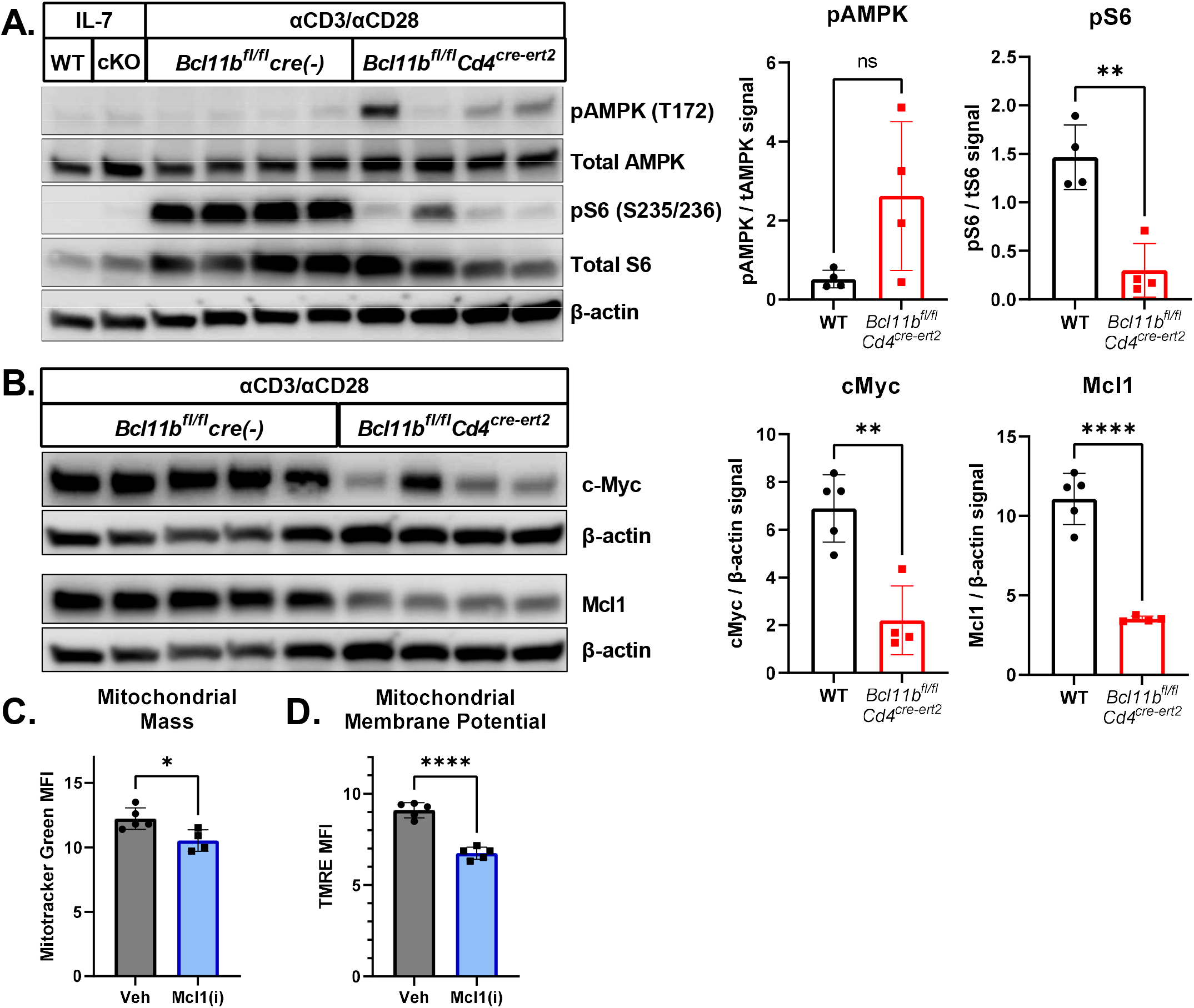
Bcl11b-deficient CD4^+^ T cells are metabolically stressed with reduced expression of c-Myc and Mcl1. **(A-B)** Western blots of CD4^+^ T cells from Bcl11b KO mice or litter mate controls stimulated for 3 days with αCD3/αCD28 (*n*=3-4; mean signal relative to β-actin ± SD). **(C-D)** WT CD4^+^ T cells stimulated 3 with αCD3/αCD28 followed by 2 days expansion in IL-2 ± 500 nM S63845 (Mcl1 inhibitor) were evaluated for mitochondrial **(C)** mass and **(D)** membrane potential (*n*=5; mean ± SD). Student’s unpaired two-tailed t-test, * p<0.05, *** p<0.0005, **** p<0.00005.

No obvious metabolic transcriptional targets of Bcl11b were apparent in either published gene expression, chromosome immunoprecipitation and sequencing databases of Bcl11b targets, or our RNAseq study of Bcl11b-deficient T cells (**Fig. S7C**)(Drashansky et al., 2019; Hasan et al., 2019; Kastner et al., 2010; Longabaugh et al., 2017). However, the Bcl-2 family member *Mcl1* has been shown to be a direct transcriptional target of Bcl11b(Hasan et al., 2019), and can localize to the inner mitochondrial membrane to promote regulate mitochondrial fusion and respiration(Morciano et al., 2016; Perciavalle et al., 2012). Consistent with Bcl11b as a regulator of Mcl1 transcription, Bcl11b-deficient CD4^+^ T cells had significantly lower levels of Mcl1 mRNA and protein (**Fig. 7B, Fig. S9F**). Indeed, Mcl1 was the only anti-apoptotic Bcl2 family member with decreased expression upon loss of Bcl11b. Because Mcl1 can regulate mitochondrial fusion, we asked if loss of Bcl11b affected mitochondrial morphology in activated T cells. Mitochondrial morphology of activated cells was imaged by structured illumination microscopy and quantified as previously described(Rasmussen et al., 2020). Trends in the reduction of mitochondrial surface area and major axis length were observed in Bcl11b KO cells (**Fig. S10A-D**). These changes in mitochondrial morphology observed in Bcl11b-deficient T cells were similar to those seen in cardiomyocytes after MCL1 inhibition(Rasmussen et al., 2020). Notably, pharmacologically inhibiting Mcl1 in activated WT CD4^+^ T cells partially mimicked the loss of Bcl11b, with Mcl1 inhibition leading to reduced mitochondrial mass and potential (**Fig. 7C-D**). These data suggest that *Bcl11b* deficiency leads to loss of Mcl1 which impairs mitochondrial dynamics and membrane potential that may contribute to T cell dysfunction in this IEI.

## DISCUSSION

Productive immune responses require cells to rapidly transition from quiescent and metabolically homeostatic to highly active states with significant metabolic demands for growth and function(Makowski et al., 2020; O’Neill et al., 2016). This shift is governed by immunological signals, but metabolic changes and environment also directly regulate the function of the cell. Understanding of the complexity of this immunometabolic interface has grown significantly over the last two decades and underscores the importance of metabolism in regulating immune responses in health and in disease. This is highlighted by genes underlying Inborn Errors of Immunity. Of the 450 identified genes that are currently known to cause monogenic IEI in patients when mutated, 72 are also annotated as IEM genes in metabolism, and many of the remaining IEI may influence metabolic function through signaling or transcriptional regulation(Tangye et al., 2022). Interestingly, of 1387 genes mutated in IEM, only ∼5% have been clinically shown to result in an immune defect. Given the growing list of IEI and our current understanding of immunometabolism, we hypothesized that the overlap of IEI and IEM genes is significantly underappreciated. As with most Mendelian disorders, patients with IEM are rare and have significant morbidities, and may not receive an adequate immunological analysis. Conversely, IEI may not be screened for metabolic defects, although these may contribute to immune mechanisms or uncharacterized effects on other tissues of some IEI. Here we sought to leverage two gene sets derived from the IEI and IEM to identify novel immunometabolic regulators in activated CD4^+^ T cells through a series of pooled CRISPR screens that are presented in an online resource.

CRISPR screening has opened new opportunities to identify novel metabolic regulators of immunologic function. These studies have taken a variety of approaches, including whole genome CRISPR screens or targeted screens of all enzymes or genes associated with specific metabolic pathways(Covarrubias et al., 2020; Henriksson et al., 2019; Huang et al., 2022; Long et al., 2021; Schmidt et al., 2022; Sugiura et al., 2022). While each approach has provided new insight, each has drawbacks. CRISPR screens of large gene sets, such as in screens of the whole genome or all enzymes, can robustly identify the gene knockouts with the most significant hits. However, the scale of these screens limits experimental flexibility and largely restricts work to *in vitro* selections. Also, genes that result in subtle phenotypes may be overlooked in screens driven a handful of large effects. In contrast, targeted screens are inherently biased to examine a single area of biology, yet can identify modest effects of biological significance and can readily be performed *in vivo* in a variety of disease settings(Sugiura et al., 2022). To avoid each of these barriers, we screened gene sets derived from the IEI and IEM. These gene sets span a wide range of metabolic and immunologic pathways and were not identified by arbitrary selections but rather by unbiased natural mutation and clinical observations. In this way, screening IEM and IEI provides a diverse and unbiased yet targeted set of genes that reflect the breadth of whole genome screens while providing the flexibility targeted screens.

Strikingly, our screens identified 343 unique IEM genes not previously associated with IEI as critical for CD4^+^ T cell function (**Supplemental Table 3, FIGS:** https://figs.app.vumc.org/). Some of these hits have recently been shown to play important roles with T cells in individual pre-clinical studies, such as Mat2a that generates S-adenosylmethionine(Roy et al., 2020). A few select others encode subunits of complexes, such as the v-ATPase, that have been shown to be necessary for T cell function in patients(Jansen et al., 2016; Peterson et al., 2019; Rujano et al., 2017; Van Damme et al., 2017). However, given that only 72 IEM genes are recognized as also causing IEI when mutated, these data provide a significant expansion of genes to be consider as candidate IEI and that may contribute to previously undescribed potential immune dysfunction in IEM patients.

Our screens using IEM genes to identify pathways critical for CD4^+^ T cell expansion and survival *in vitro* and expansion, recruitment, and persistence *in vivo* in a colitis model identified a wide range of genes associated with N-linked glycosylation as a critical metabolic process for CD4^+^ T cell function. N-linked glycosylation utilizes UDP-N-Acetylglucosamine (UDP-GlcNAc) generated through the HBP as a substrate for posttranslational modifications to regulate protein stability, trafficking, and function. Mutations in over 40 genes associated with N-linked glycosylation have been recognized as IEM(Ferreira et al., 2019). These cases result in developmental disorders of glycosylation characterized primarily by growth failure, developmental delay, and facial dysmorphisms. While CD4^+^ T cells have only rarely been evaluated in these patients, a growing number exhibit a range of immune disorders(Monticelli et al., 2016). Top hits in these screens included *Gfpt1* and *Pgm3* (UDP-GlcNAc synthesis via the HBP), *Nus1* and *Dpagt1* (dolichol synthesis), *Alg11* and *Alg13* (acetylglucosamine transferases), and *Slc35a1* and *Slc35a2* (sugar transporters). Of these, mutations in only 7 glycosylation-related genes have been recognized to date as causing IEI, most notably *PGM3* and *MAGT1* (**Fig. 1B**). Patients with mutations in *PGM3*, which is involved in the production UDP-GlcNAc used for N-linked glycosylation, exhibit several immunologic defects, including T cell lymphopenia and hyper-IgE syndrome, although the mechanisms remain unclear(Sassi et al., 2014; Stray-Pedersen et al., 2014; Zhang et al., 2014). Previous work has shown that N-linked glycosylation is important for surface translocation of several cytokine receptors, including IL2Rα (CD25) and IL23R, suggesting that modulation of N-linked glycosylation could have significant effects on CD4^+^ T cell polarization(Araujo et al., 2017; Chien et al., 2015). Patients with bi-allelic mutations in *IL23R* that disrupt its N-linked glycosylation exhibit normal T_H_17 responses but exhibit impaired ability to develop pathogenic IL-17A^+^IFNγ^+^ T_H_1^*^ cells(Harbour et al., 2015; Martinez-Barricarte et al., 2018). *IL23R* variants have also been reported to protect against inflammatory bowel disease (IBD) development, indicating a potential role for N-linked glycosylation in pathogenic CD4^+^ T cell function(Sivanesan et al., 2016).

Here we show that Gfat (*Gfpt1*), the enzyme regulating the first step for *de novo* synthesis of UDP-GlcNAc, is critical for CD4^+^ T cell expansion *in vitro* and *in vivo* but differentially required for polarization. Patients with autosomal recessive *GFPT1* mutations present with congenital myasthenia syndrome, but no immunological phenotype has been reported. Notably, while *Gfpt1* was broadly required for optimal CD4^+^ T cell expansion, it appeared dispensable for T_H_17 and iTreg polarization. In contrast, T_H_1 cells were highly reliant on *Gfpt1* for both expansion and polarization, with reductions in both IFNγ and Tbet expression upon Gfat deficiency. The differential requirement for Gfat in different CD4^+^ T cell subsets may be attributable to several factors. Previous work has suggested that T_H_1 cells are more heavily glycosylated than T_H_17 cells(Araujo et al., 2017). Additionally, Gfat is more highly expressed in T_H_1 cells. In contrast, T_H_17 cells express higher levels of Nagk, an enzyme that salvages GlcNAc to support HBP by generating GlcNAc-6P as a substrate for Pgm3. These data suggest that T_H_17 cells both require less UDP-GlcNAc and may be more adept at salvaging hyaluronic acid or free GlcNAc to shunt into the HBP. This may also explain why *PGM3* mutations are more likely to present with a more notable immunological phenotype. While loss of Gfat function will impair *de novo* UDP-GlcNAc synthesis, reduced Pgm3 function would block both *de novo* synthesis and the GlcNAc salvage pathway.

Our metabolic screens of IEI genes identified multiple potentially novel immunometabolic regulators of CD4^+^ T cells. Among established IEI genes, *Bcl11b* was one of the most consistently differentially abundant in our mitochondrial and nutrient uptake screens. Bcl11b is primarily recognized for its role as a regulator of T cell specification during thymic development, where it promotes the transcription of machinery necessary for positive selection while suppressing the mature transcriptional program(Albu et al., 2007; Albu et al., 2011; Kastner et al., 2010; Longabaugh et al., 2017). Loss of Bcl11b in developing thymocytes results in impaired thymic development and reduced total T cell populations in the periphery that exhibit a memory-like phenotype. Patients with *BCL11B* mutations develop combined immunodeficiency with low T cell numbers, due to this block in productive development(Albu et al., 2007; Avram and Califano, 2014; Drashansky et al., 2019; Hasan et al., 2019; Lessel et al., 2018; Punwani et al., 2016). Further interrogation of Bcl11b in conditional KO mouse models demonstrated that loss of Bcl11b interrupts T cell development specifically at the CD4^+^CD8^+^ double positive (DP) stage in the thymus and is critical for regulatory T cell function in the periphery. However, Bcl11b has not been previously described to affect to T cell metabolism.

Despite enhanced early proliferation Bcl11b deficient CD4^+^ T cells exhibit impaired mitochondrial function and ultimately decreased proliferation and function. *In vivo*, Bcl11b-deficient CD4 SP thymocytes also exhibited an altered mitochondrial phenotype and mitochondrial imaging showed altered mitochondrial morphology suggestive of mitochondrial fission. Further analysis showed that Bcl11b-deficient CD4^+^ T cells failed to upregulate Mcl1, a key regulator of mitochondrial morphology and function that can promote mitochondrial fusion(Morciano et al., 2016; Perciavalle et al., 2012). Interestingly, Mcl1 has two isoforms with different functions; while the full form functions as an anti-apoptotic protein like other Bcl2 family members, the truncated form localizes to the inner mitochondrial membrane, where it regulates mitochondrial fusion and function(Perciavalle et al., 2012; Prew et al., 2022). Loss of Bcl11b resulted in reduced Mcl1 levels, altered mitochondrial morphology, and reduced mitochondrial function, phenotypes that were recapitulated by pharmacological targeting of Mcl1. These data show that Bcl11b regulates mitochondrial function during T cell development and activation, which may be essential for thymopoiesis as well as contribute to the defects observed in Bcl11b-deficient Tregs(Drashansky et al., 2019; Hasan et al., 2019). While effector T cells rely primarily on aerobic glycolysis for synthesis of metabolic precursors and their energetic needs, regulatory T cells are much more dependent upon mitochondrial oxidative phosphorylation to maintain both their metabolic needs and their regulatory function(Gerriets et al., 2015). In instances were Tregs are forced to become more glycolytic, their suppressive function is significantly impaired(Gerriets et al., 2016). As such, the loss mitochondrial function observed in Bcl11b-deficient T cells may be a driver of the impaired functionality of Bcll1b^-/-^ Tregs previously reported.

The functional intersection of IEI and IEM genes may provide important insight to both categories of diseases. IEM are often not fully examined for potential immune dysregulation and our data suggest that these phenotypes may be more frequent than currently appreciated. As such, patients with IEM may also have undiagnosed mild-to-severe immune deficiencies or inflammatory features that could impact their care. Notably, infection in patients with select IEM can often trigger a “metabolic crisis,” characterized an acute worsening of organ dysfunction(Tarasenko and McGuire, 2017). Furthermore, patients due to mutations in *ABCD1, GALC, IDUA, and ARSA* are frequently treated by bone marrow transplant. While this may serve as a functional enzyme replacement therapy, it may also point to immune-specific roles for these metabolic enzymes(Yu et al., 2022). Similarly, IEI patients may have previously undescribed metabolic defects that contribute to mechanisms of immune dysfunction or alter the function of other tissues. The large overlap of IEM and IEI that we describe here also reveals a potential for a new class of IEI based on metabolic dysfunction. IEI are classified based on immune phenotypes, but shared immunometabolic characteristics may provide new understanding of existing IEI and point to a considerable source of new IEI that are currently annotated as only IEM. These data are presented in an online Funtional ImmunoGenomics resource (FIGS; https://figs.app.vumc.org/).

## Limitations of the Study

All CRIPSR screens performed in this study used active Cas9 transgenic mice, which results significant disruption of the sgRNA gene target, resulting in near complete loss of protein product. In contrast, a proportion of variants described as IEM or IEI result in hypomorphic, but still present, protein products. Similarly, many variants are pathogenic in patients even at heterozygosity. Notably, analysis of *BCL11B* IEI variants shows that heterozygous pathogenic variants result in a dominant negative form of BCL11B. Thus, while this study uses the Inborn Errors to select genes for CRISPR screens, it does not directly test the effect of patient variants in T cell function or metabolism. Furthermore, due to the technical challenges to efficiently introduce sgRNAs into non-dividing cells, all CRISPR screens were performed in activated CD4^+^ T cells. As such, loss of protein 3 days post activation, at earliest. With this approach, the effect of protein loss in T cell development, early activation, or polarization initialization were not determined from these screens. Pooled CRISPR screens are performed by analyzing relative sgRNA abundance in each population compared to another population; it does not directly measure any given readout. As such, direct measurements of individual gene knockouts are necessary to confirm any screen result. Finally, a significant number of additional gene variants have been characterized as IEM (detailed at IEMBase.org) or IEI (published in IUIS’s 2022 report) since the generation of these initial pooled CRISPR libraries and were not evaluated in these screens.

## Supporting information

Graphic Abstract

## Acknowledgements

The authors thank members of the J.C. Rathmell and W.K. Rathmell laboratories and Drs. Ruben Martinez-Barricarte and James Connelly for helpful discussions.

## Author contributions

A.R.P. conceptualized the study, designed all experiments, conducted experiments and analyses, and wrote the first draft of the manuscript. K.K.S., C.C., E.L.F. performed some experiments. A.S. designed the initial CRISPR system used for screening. G.A.N. analyzed the RNAseq data and constructed the FIGS website. G.L.R. and C.B performed structured illuminated microscopy and subsequent analysis under the supervision of V.G. J.G.M. provided intellectual contributions. J.C.R. supervised the project, research design, and writing of the manuscript.

## Data and materials availability

The deposited data for RNA sequencing experiments are available under the NCBI GEO accession number GSE223334. Data for all CRISPR screen results are available for download on the Funtional ImmunoGenomics resource (FIGS; https://figs.app.vumc.org/). All materials using or generated in this study are available to researchers following appropriate standard material transfer agreement.

