## Supplementary figures and images for "Functional Overlap of Inborn Errors of Immunity and Metabolism Genes Define T Cell Immunometabolic Vulnerabilities"

### Graphic Abstract

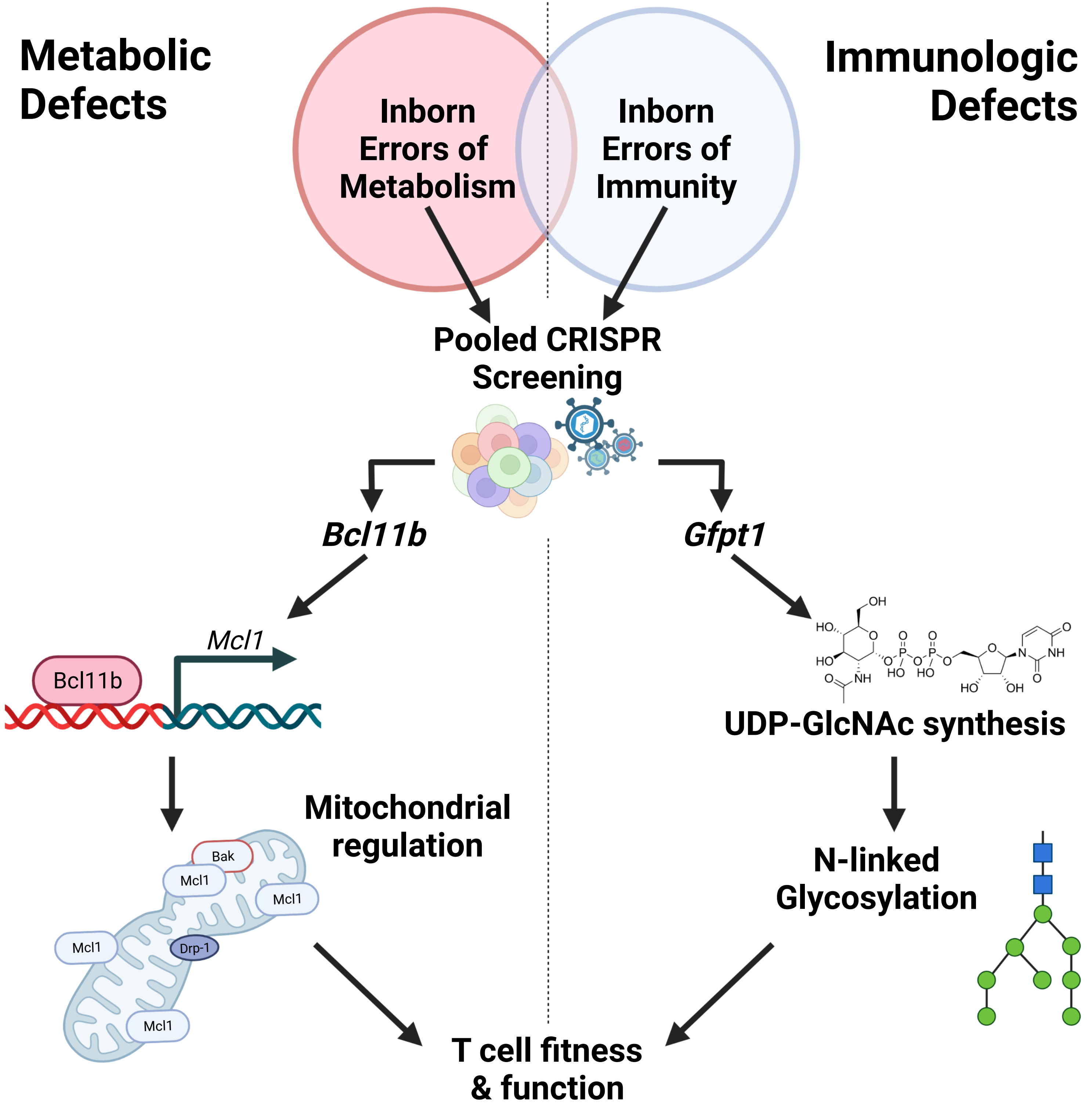

### Supplemental Figures 1-10

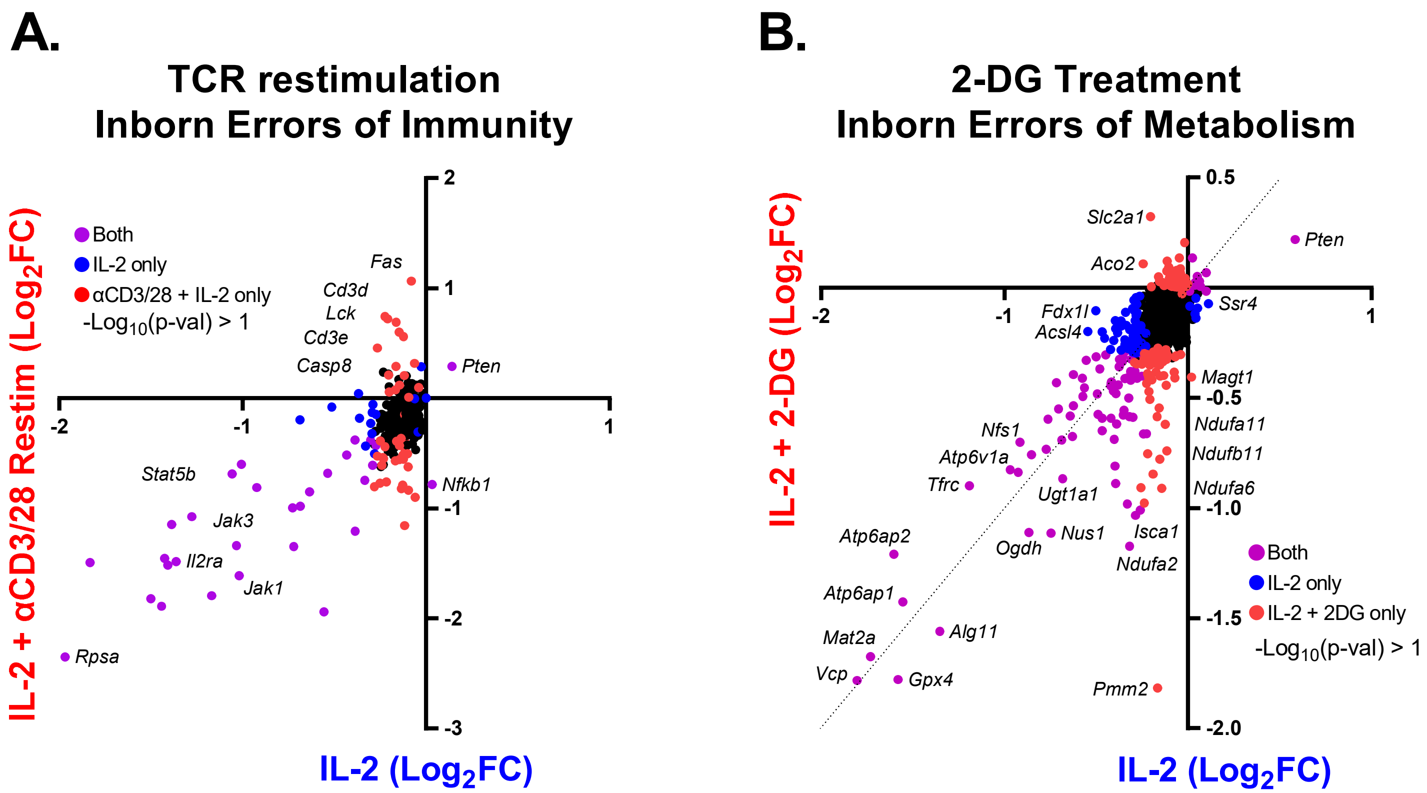

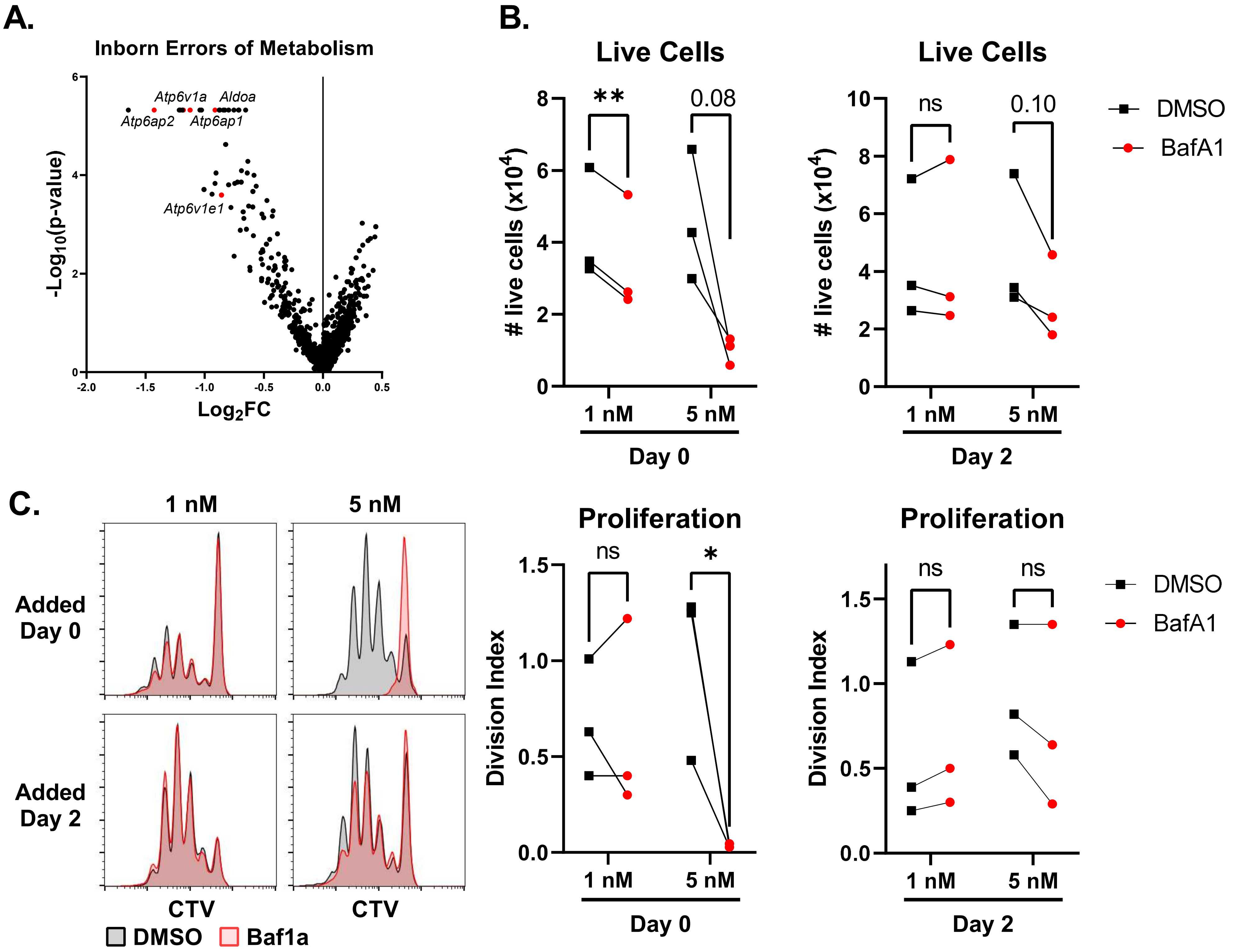

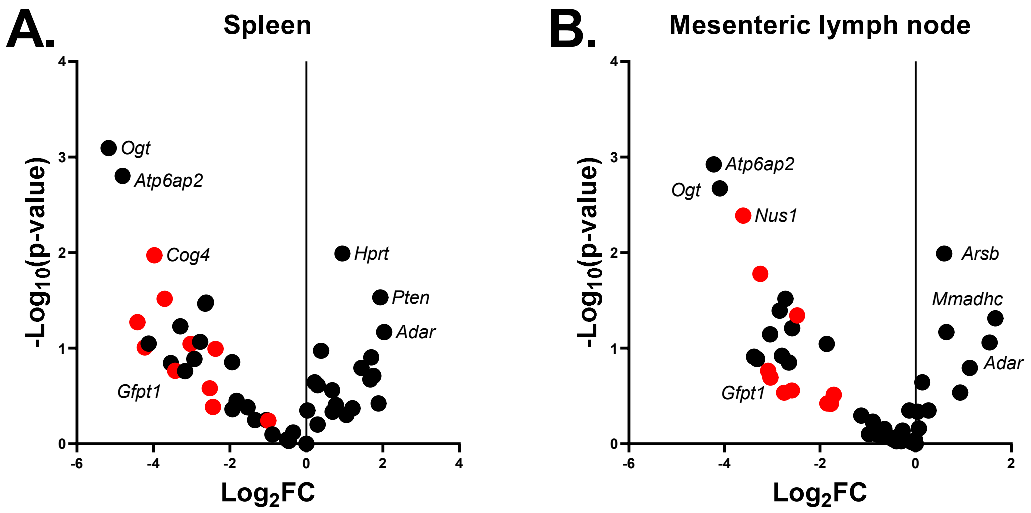

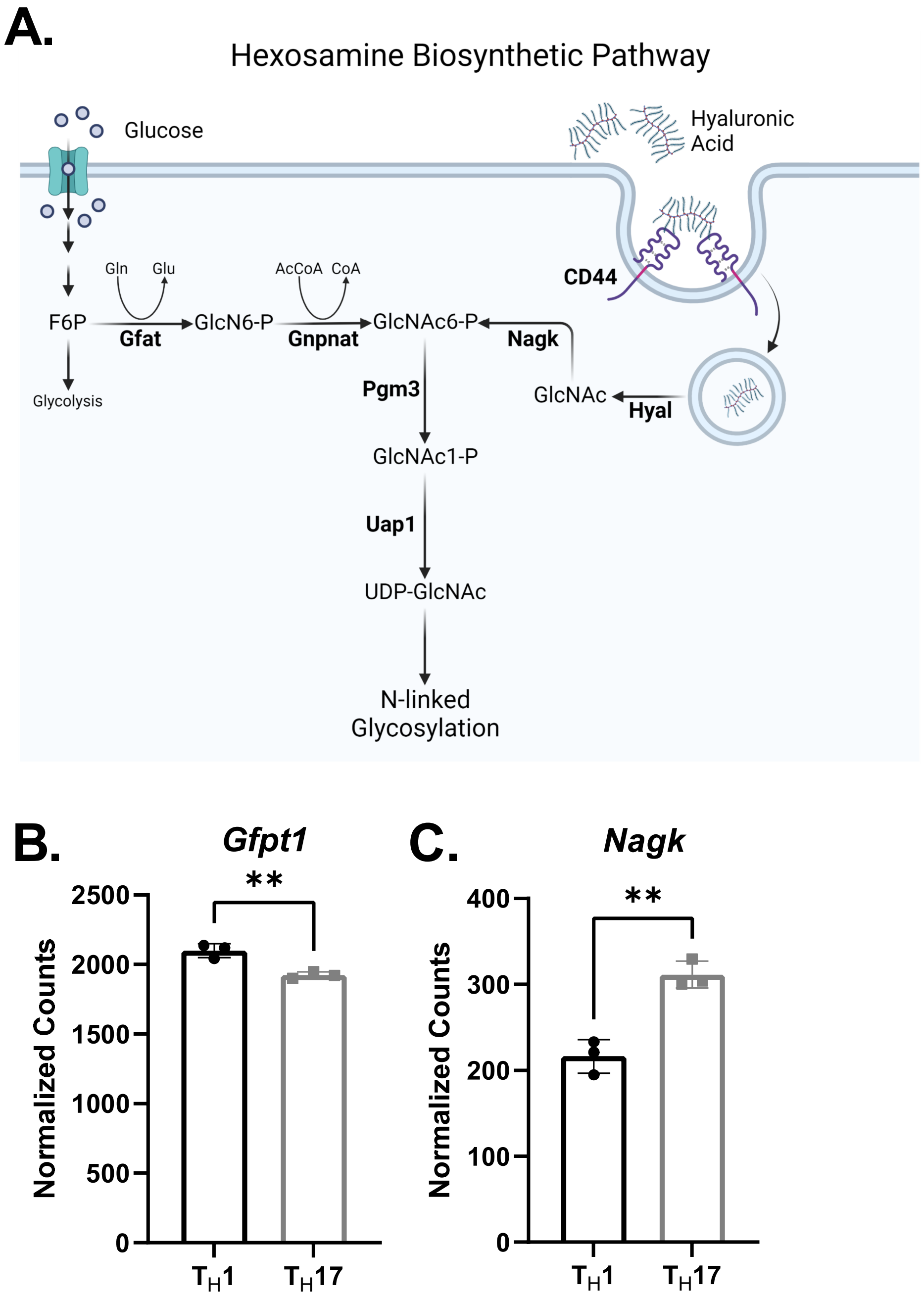

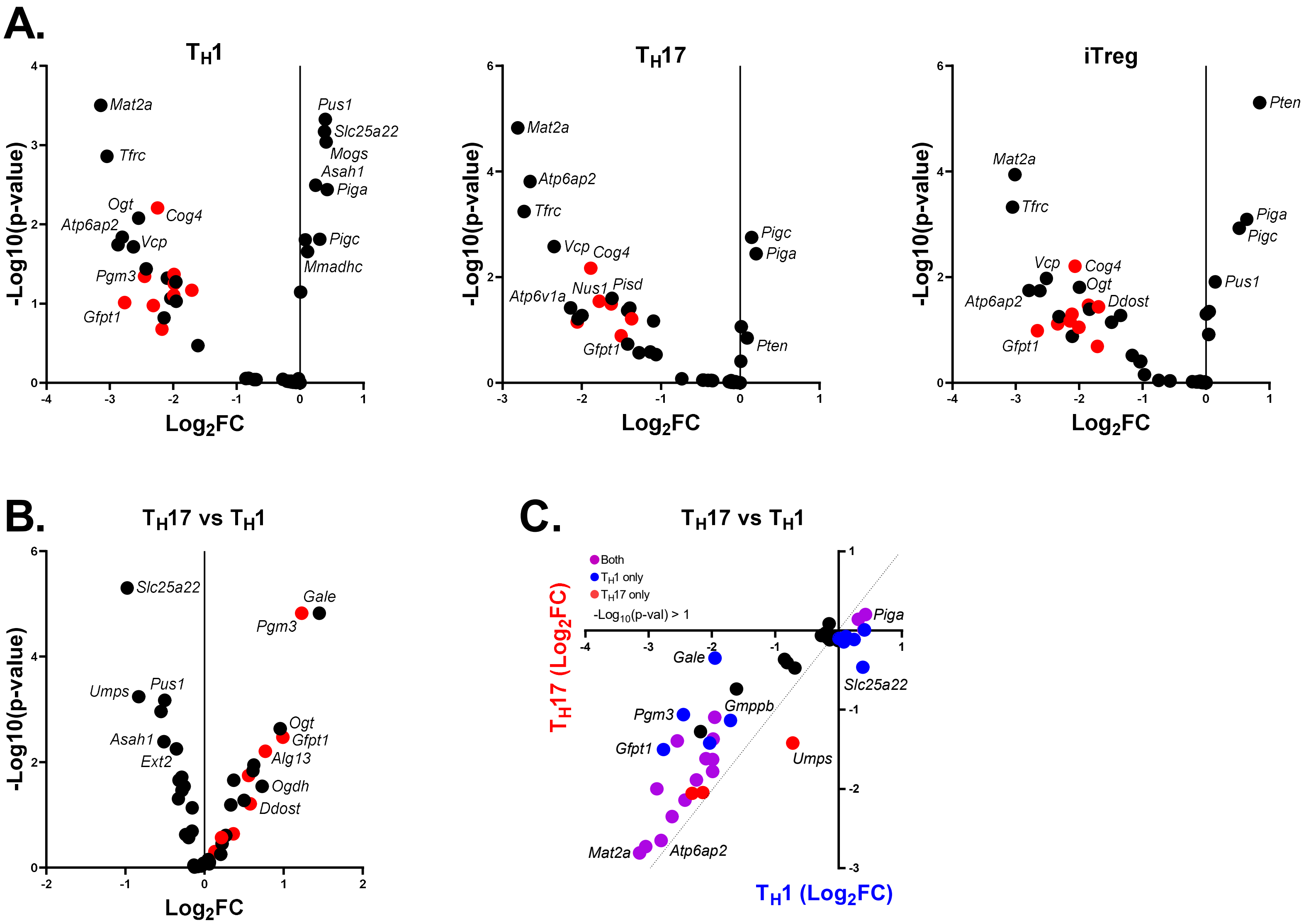

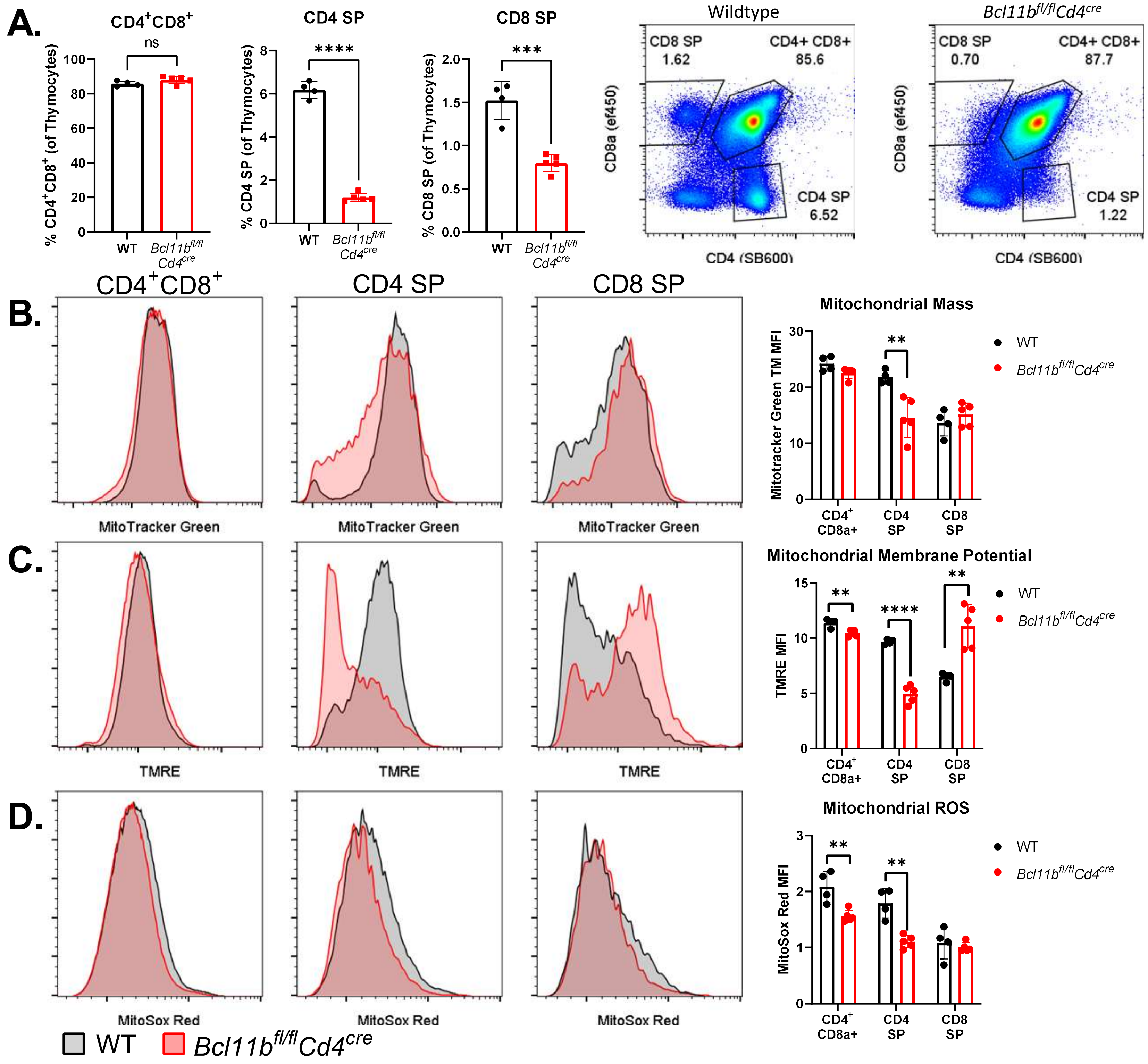

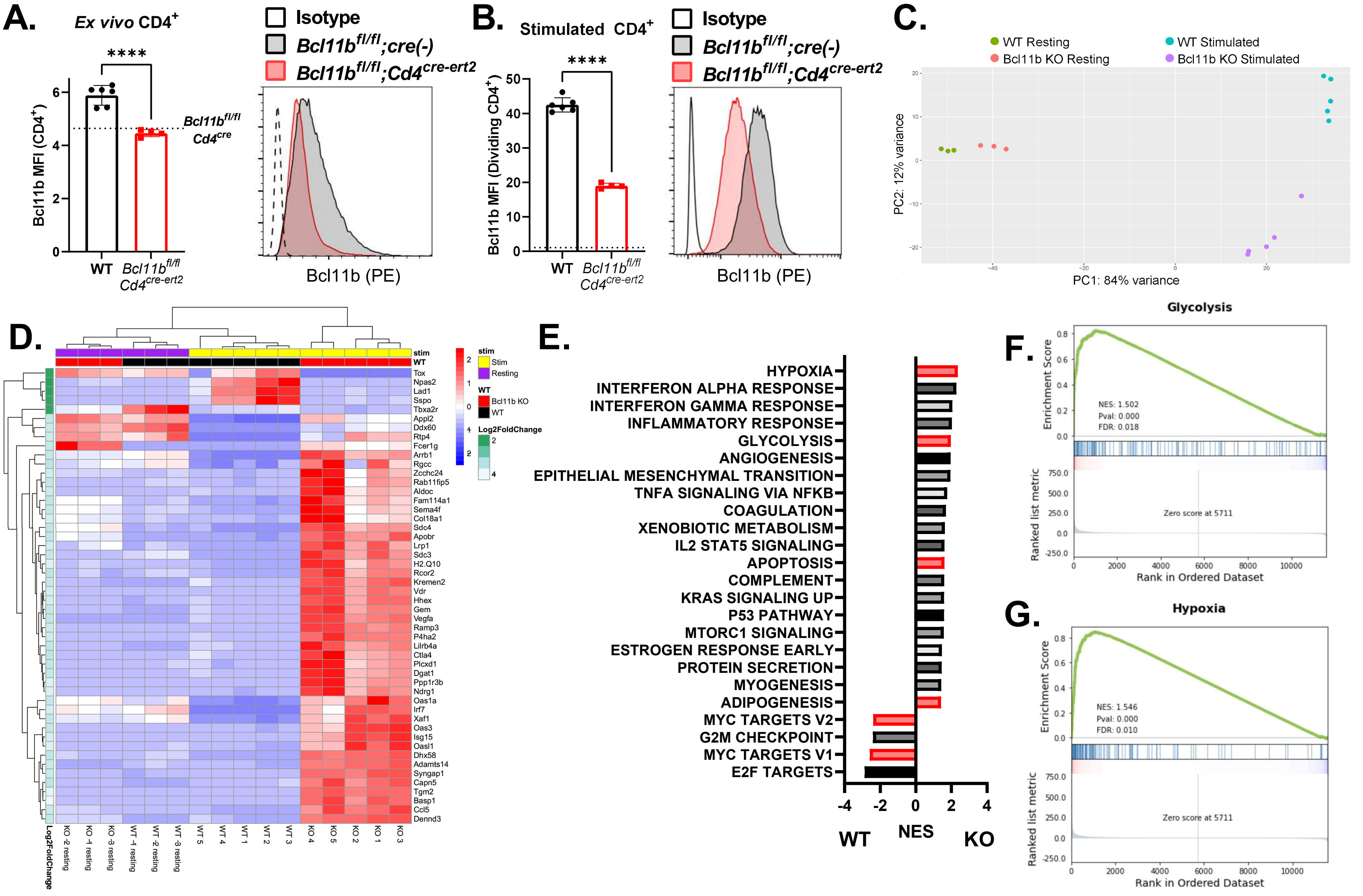

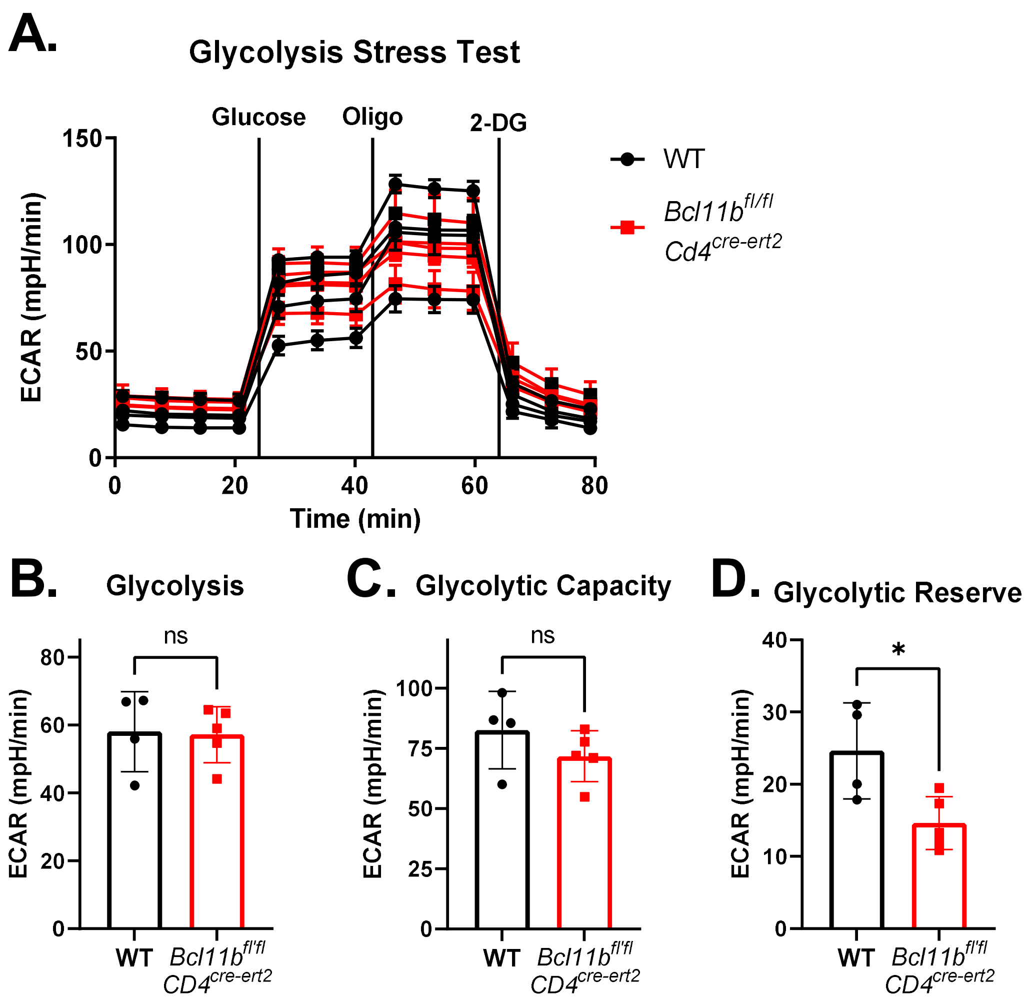

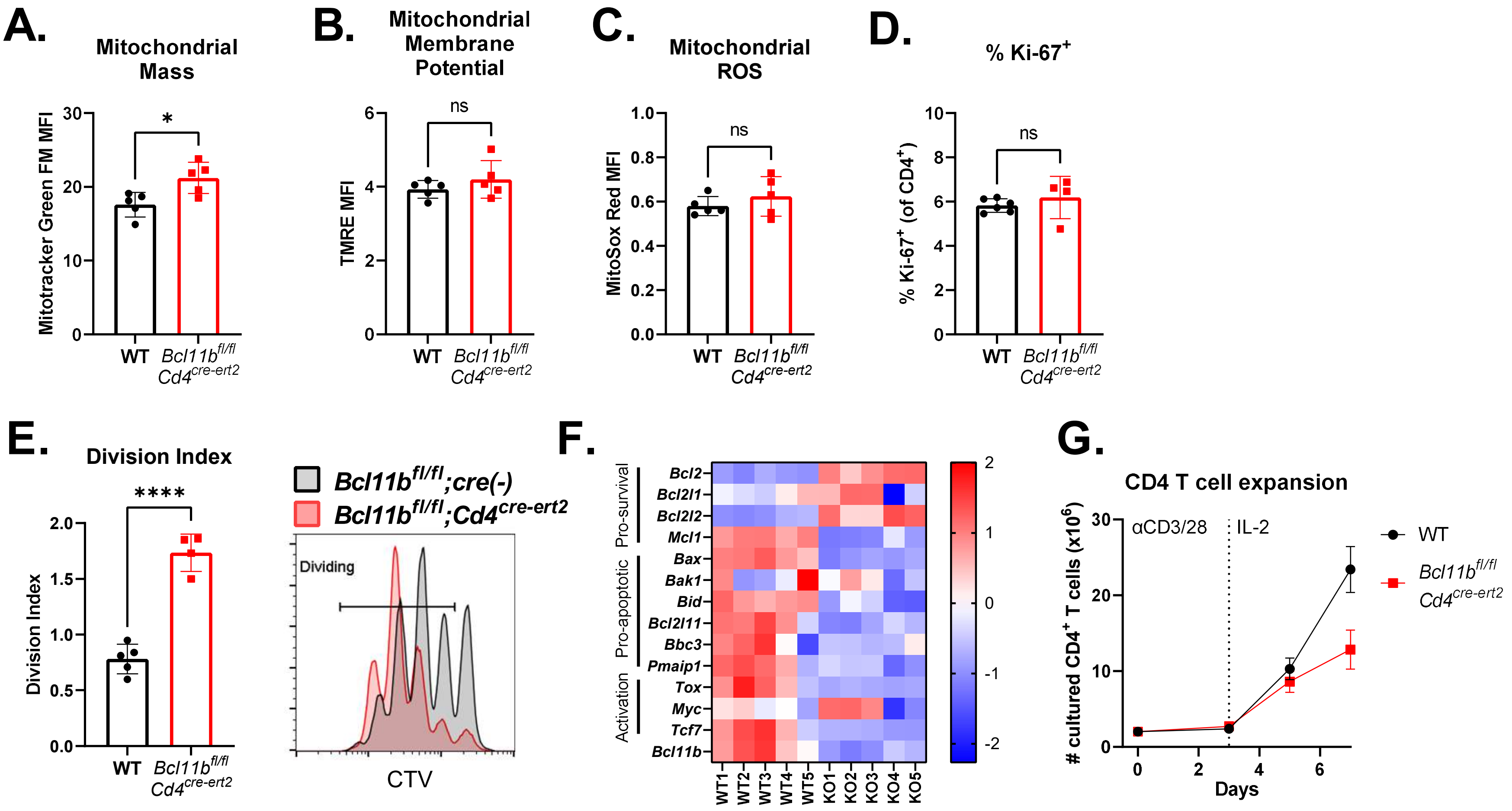

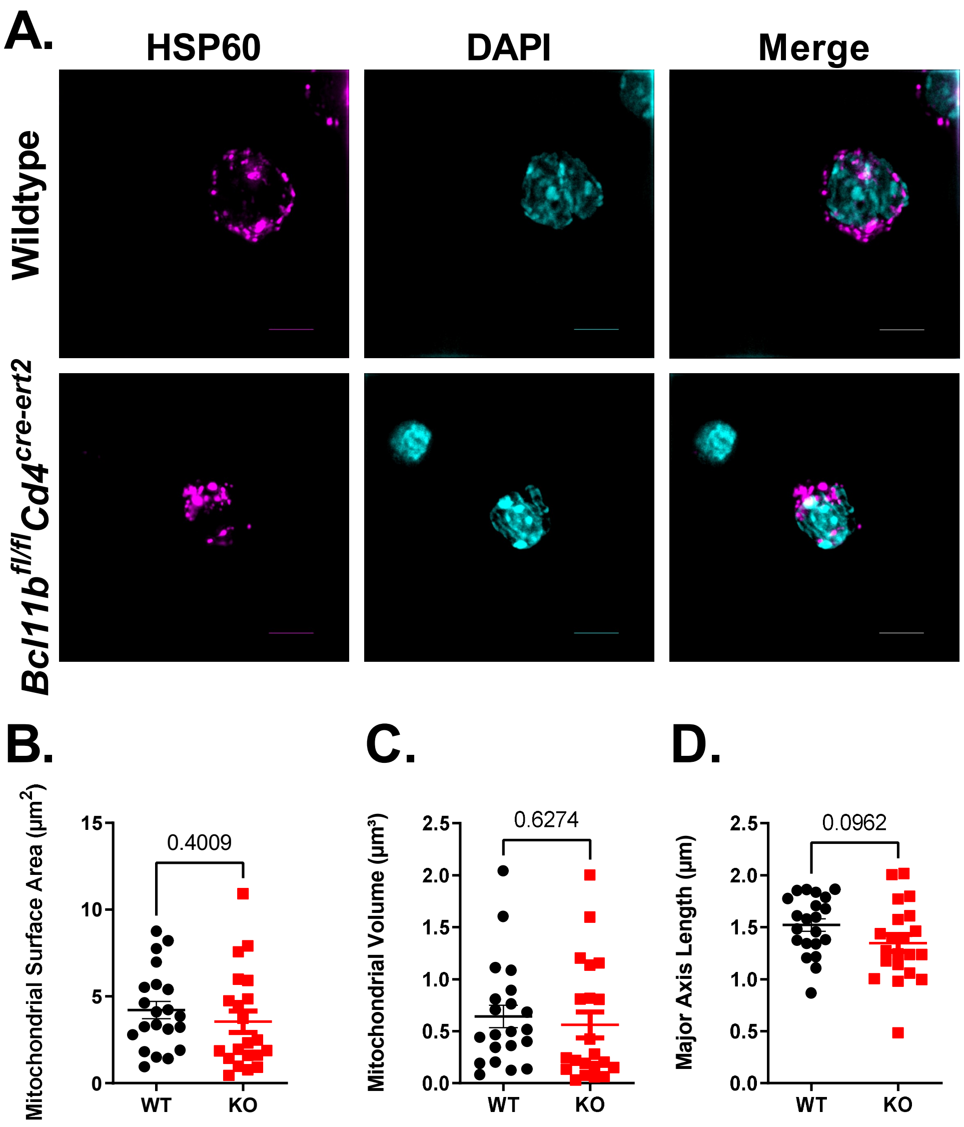
