## Supplementary material for "Functional Overlap of Inborn Errors of Immunity and Metabolism Genes Define T Cell Immunometabolic Vulnerabilities": Methods Text

**STAR METHODS**

**Key Resources Table**

| **Reagent or Resource** | **Source** | **Identifier** |
| --- | --- | --- |
| **Antibodies** | | |
| Anti-IFNγ | Thermo Fisher Scientific | Cat# 16-7311-38, RRID:AB_2637490 |
| Anti-IL4 functional | Thermo Fisher Scientific | Cat# 16-7041-95, RRID:AB_2573101 |
| Mouse Anti-CD3e | Thermo Fisher Scientific | Cat# 16-0031-86, RRID:AB_468849 |
| Mouse Anti-CD28 | Thermo Fisher Scientific | Cat# 16-0281-86, RRID:AB_468923 |
| Anti-CD4 SB600 | Thermo Fisher Scientific | Cat# 63-0041-82, RRID:AB_2717017 |
| Anti-CD8a e450 | Invitrogen | Cat# 48-0081-82, RRID:AB_1272198 |
| Anti-CD3 APC | Biolegend | Cat# 100236, RRID:AB_2561456 |
| Anti-CD45.2 BV510 | Biolegend | Cat# 109838, RRID:AB_2650900 |
| Anti-CD44 PE-Cy7 | Biolegend | Cat# 103030, RRID:AB_830787 |
| Anti-CD25 APC | Thermo Fisher Scientific | Cat# 17-0251-81, RRID:AB_469365 |
| Anti-IL23R PE | BioLegend | Cat# 150903, RRID:AB_2572188 |
| Anti-IFNγ APC | BD Biosciences | Cat#554413; RRID:AB_398551 |
| Anti-IL17A PE | Thermo Fisher Scientific | Cat#12-7177-81; RRID:AB_763582 |
| Anti-Mcl1 PE | Invitrogen | Cat# 12-9047-42, RRID:AB_2762598 |
| Anti-Ki-67 e450 | Invitrogen | Cat# 48-5698-82, RRID:AB_11149124 |
| Anti-FoxP3 e450 | Thermo Fisher Scientific | Cat# 48-5773-82, RRID:AB_1518812 |
| Anti-T-bet APC | Thermo Fisher Scientific | Cat# 17-5825-82, RRID:AB_2744712 |
| Anti-RORγt PE | Thermo Fisher Scientific | Cat# 12-6988-82, RRID:AB_1834470 |
| Isotype PE control | Thermo Fisher Scientific | Cat#12-4714-42; RRID:AB_1944423 |
| Anti-Bcl11b | Invitrogen | Cat# PA5-83952, RRID:AB_2791104 |
| Rabbit IgG Isotype | Cell Signaling Technology | Cat# 3900, RRID:AB_1550038 |
| mIgG1 Isotype Control PE | Invitrogen | Cat# 12-4714-42, RRID:AB_1944423 |
| Rat IgG2a Isotype APC | Invitrogen | Cat# 17-4321-81, RRID:AB_470181 |
| Anti-Rabbit IgG PE | Invitrogen | Cat# 12-4739-81, RRID:AB_1210761 |
| Anti-Rabbit IgG AF488 | Invitrogen | Cat# A11034, RRID:AB_2576217 |
| Anti-Hsp60 | Cell Signaling Technology | Cat# 12165, RRID:AB_2636980 |
| Anti-Gfpt1 | Proteintech | Cat# 14132-1-AP, RRID:AB_2110155 |
| Anti-Nagk | Proteintech | Cat# 15051-1-AP, RRID:AB_2152368 |
| Anti-pAMPK (T172) | Cell Signaling Technology | Cat# 2535, RRID:AB_331250 |
| Anti-APMK | Cell Signaling Technology | Cat# 2793, RRID:AB_915794 |
| Anti-pS6 (S235/236) | Cell Signaling Technology | Cat# 4858, RRID:AB_916156 |
| Anti-S6 | Cell Signaling Technology | Cat# 2317, RRID:AB_2238583 |
| Anti-cMyc | Cell Signaling Technology | Cat# 9402, RRID:AB_2151827 |
| Anti-Mcl1 | Cell Signaling Technology | Cat# 5453, RRID:AB_10694494 |
| Anti-ß-actin | Cell Signaling Technology | Cat# 4970, RRID:AB_2223172 |
| Anti-ß-actin | Cell Signaling Technology | Cat# 3700, RRID:AB_2242334 |
| Anti-mouse IgG-HRP | Promega | Cat# W402B, RRID:AB_430834 |
| Anti-rabbit IgG-HRP | Promega | Cat# W401B, RRID:AB_430833 |
| **Experimental Models: Organisms/Strains** | | |
| Mouse: C57BL/6J | Jackson Laboratory | JAX: 000664, RRID:IMSR_JAX:000664 |
| Mouse: *Rag1-/-* (B6.129S7-Rag1tm1Mom/J) | Jackson Laboratory | JAX: 002216, RRID:IMSR_JAX:002216 |
| Mouse: Cas9 (B6J.129(Cg)-Gt(ROSA)26Sortm1.1(CAG-cas9*,-EGFP)Fezh/J) | Jackson Laboratory | JAX: 026179, RRID:IMSR_JAX:026179 |
| Mouse: Bcl11b^fl/fl^ (STOCK Bcl11btm1.1Leid/J) | Jackson Laboratory | JAX: 034469, RRID:IMSR_JAX:034469 |
| Mouse: CD4-Cre-ert2 (B6(129X1)-Tg(Cd4-cre/ERT2)11Gnri/J) | Jackson Laboratory | JAX: 022356, RRID:IMSR_JAX:022356 |
| Mouse: CD4-Cre (B6.Cg-Tg(Cd4-cre)1Cwi/BfluJ ) | Jackson Laboratory | JAX: 022071, RRID:IMSR_JAX:022071 |
| **Experimental Models: Cell Lines** | | |
| Plat-E retroviral packaging cell line | Cell Biolabs | Cat# RV-101, RRID:CVCL_B488 |
| **Chemicals, Peptides, and Recombinant Proteins** | | |
| Recombinant murine IL-7 | RND Biosystems | Cat# 407-ML-200 |
| Recombinant murine IL-12p70 | Thermo Fisher Scientific | Cat# 14-8121-62 |
| Recombinant murine IL-6 | Miltenyi Biotec | Cat# 130-096-683 |
| Recombinant murine IL-23 | Miltenyi Biotec | Cat# 130-096-676 |
| Recombinant murine IL-1ß | Miltenyi Biotec | Cat# 130-101-681 |
| Recombinant human IL-2 | NCI | Cat# Ro 23-6019 |
| Recombinant human TGFß1 | Peprotech | Cat# 100-21 |
| Mitotracker Green FM | Thermo Fisher Scientific | M7514 |
| Tetramethylrhodamine, Ethyl Ester, Perchlorate (TMRE) | Thermo Fisher Scientific | T669 |
| MitoSOX Mitochondrial Superoxide Indicators | Thermo Fisher Scientific | M36008 |
| 2-NBDG | Invitrogen | Cat# N13195 |
| C16-BODIPY | Thermo Fisher Scientific | Cat# D3821 |
| 2-deoxy-D-glycose (2-DG) | Cayman Chemical Company | Cat# 14325 |
| BafA1 | Millipore | Cat# 19-148 |
| S63845 | MedKoo Biosciences | Cat# 406849 |
| Tamoxifen | Sigma-Aldrich | Cat# T5648 |
| Corn Oil | Sigma Aldrich | Cat# C8267 |
| FBS | Avantor | Cat# 97068-085 |
| IMDM | Gibco | Cat# 12440-053 |
| RPMI 1640 | Corning | Cat# 10-040-CV |
| DMEM | Thermo Fisher Scientific | Cat# 11965-092 |
| HEPES | Thermo Fisher Scientific | Cat# 15630-080 |
| Penicillin/streptomycin | Thermo Fisher Scientific | Cat# 15140-122 |
| Puromycin | Santa Cruz Biotechnology | Cat# sc-205821 |
| Blasticidin | Thermo Fisher Scientific | Cat# A11139-03 |
| Glutamine | Thermo Fisher Scientific | Cat# 25030-149 |
| 2-mercaptoethanol | Thermo Fisher Scientific | Cat# 21985-023 |
| HBSS | Corning | Cat# 21-022-CV |
| EDTA | Thermo Fisher Scientific | Cat# 50-255-956 |
| DNAse I | Sigma Aldrich | Cat# DN25 |
| Collagenase T4 | Worthington Biochemical | Cat# CLS-4 |
| PMA | Sigma-Aldrich | Cat# P8139 |
| Ionomycin | Sigma-Aldrich | Cat# IO634 |
| GolgiPlug | BD Biosciences | Cat# 555029 |
| Seahorse XF96 FluxPak | Agilent | Cat# 102416-100 |
| Seahorse XF RPMI | Agilent | Cat# 103567-100 |
| Seahorse XF Glucose | Agilent | Cat# 103577-100 |
| Seahorse XF Pyruvate | Agilent | Cat# 103578-100 |
| Seahorse XF Calibrant | Agilent | Cat# 100840-000 |
| Oligomycin | Cayman Chemical Company | Cat# 11342 |
| FCCP | Cayman Chemical Company | Cat# 15218 |
| Antimycin A1 | Cayman Chemical Company | Cat# 19433 |
| Rotenone | Cayman Chemical Company | Cat# 13995 |
| Cell-Tak | Corning | Cat# 354240 |
| Poly-lysine | Millipore Sigma | Cat# A-003-E |
| Retronectin | Takara Bio | Cat# T100A |
| ACK Lysing Buffer | Thermo Fisher Scientific | Cat# A1049201 |
| Ghost Dye Red 780 Viability Dye | Cell Signaling Technology | Cat# 18452 |
| RIPA Buffer | Thermo Fisher Scientific | Cat# 89900 |
| Halt Protease and Phosphatase Inhibitor | Thermo Fisher Scientific | Cat# 78442 |
| Bio-Rad Protein Assay Dye Reagent Concentrate | Bio-Rad | Cat# 5000006 |
| ECL Western Substrate | Thermo Fisher Scientific | Cat# 32106 |
| West Femto Maximum Sensitivity Substrate | Thermo Fisher Scientific | Cat# 34095 |
| BbsI-HF | New England Biolabs | Cat# R3539 |
| Prolong Gold Antifade + DAPI Reagent | Cell Signaling Technology | Cat# P36931 |
| Lympholyte-M cell separation media | Cedarlane | Cat# CL5030 |
| **Critical Commercial Assays** | | |
| CD4^+^ T cell Isolation Kit, mouse | StemCell Technologies | Cat# 19852 |
| CD4^+^ T Cell Isolation Kit, mouse | Miltenyi Biotec | Cat# 130-104-454 |
| Naïve CD4^+^ T Cell Isolation Kit, mouse | Miltenyi Biotec | Cat# 130-104-453 |
| CellTrace Violet Cell Proliferation Kit | Thermo Fisher Scientific | Cat# C34557 |
| Fixation/Permeabilization Solution Kit | BD Biosciences | Cat# 554714 |
| Foxp3/Transcription Factor Staining Buffer Set | Thermo Fisher Scientific | Cat# 00-5523-00 |
| BCA Protein Assay Kit | Thermo Fisher Scientific | Cat# 23225 |
| KAPA Mouse Gentoyping Kits | Roche Diagnostics | Cat# 07961804001 |
| QIAquick Gel Extraction Kit | Qiagen | Cat# 28706 |
| GeneJET Plasmid Maxiprep Kit | Thermo Fisher Scientific | Cat# K0491 |
| Herculase II Fusion DNA Polymerase | Agilent | Cat# 600675 |
| Gibson Assembly Master Mix | NEB | Cat# E2611 |
| jetPRIME DNA and siRNA transfection reagent | VWR | Cat# 89129-922 |
| RNeasy Mini Kit | Qiagen | Cat# 74106 |
| **Bacterial and Virus Strains** | | |
| ElectroMAX DH10B Cells | Thermo Fisher Scientific | Cat#: 18290015 |
| **Oligonucleotides** | | |
| Array primers:  Forward – TAACTTGAAAGTATTTCGATTTCTTGGCTTTATATATCTTGTGGAAAGGACGAAACACCG  Reverse - ACTTTTTCAAGTTGATAACGGACTAGCCTTATTTTAACTTGCTATTTCTAGCTCTAAAAC | (Shalem *et al.,* 2014) | n/a |
| Adapter primers:  Forward – AATGGACTATCATATGCTTACCGTAACTTGAAAGTATTTCG  Reverse - ACGGCATCGCAGCTTGGATACA | Modified from (Shalem *et al.,* 2014) | n/a |
| Sequencing primers:  Forward –  AATGATACGGCGACCACCGAGATCTACACTCTTTCCCTACACGACGCTCTTCCGATCTNNNNNNNNNTCTTGTGGAAAGGACGAAACACCG  Reverse - CAAGCAGAAGACGGCATACGAGATXXXXXXXXGTGACTGGAGTTCAGACGTGTGCTCTTCCGATCTACGGCATCGCAGCTTGGATACA | Modified from (Shalem *et al.,* 2014) | n/a |
| Inborn Errors of Immunity (IEI) sgRNA library | This paper | n/a |
| Inborn Errors of Metabolism (IEM) sgRNA library | This paper | n/a |
| Inborn Errors of Metabolism (IEM) – Small #1 sgRNA library | This paper | n/a |
| **Recombinant DNA** | | |
| pMx-U6-empty-GFP plasmid | (Taffalini *et al*., 2009) | n/a |
| pMx-U6-empty-BFP plasmid | This paper | n/a |
| **Software and Algorithms** | | |
| Prism v8 | GraphPad Software | https://www.graphpad.com; RRID: SCR_002798 |
| FlowJo v10 | FlowJo | https://www.flowjo.com; RRID: SCRJD08520 |
| MaGECK | (Li *et al.,* 2014) | n/a |
| WebGestalt | WebGestalt: WEB-based GEne SeT AnaLysis Toolkit | <https://www.webgestalt.com>; RRID:SCR_006786 |
| BioRender | BioRender | <https://www.BioRender.com>; RRID:SCR_018361 |

**Lead Contact and Materials Availability**

Further information and requests for resources and reagents should be directed to and will be fulfilled by the Lead Contact, Dr. Jeffrey Rathmell. CRISPR sgRNA libraries and screening results are available online at the Functional ImmunoGenomics reSource (FIGS; [https://figs.app.vumc.org/](https://nam12.safelinks.protection.outlook.com/?url=https%3A%2F%2Ffigs.app.vumc.org%2F&data=05%7C01%7Candrew.patterson%40vumc.org%7C219dcbb35f3f4f5aaba208daf4b6dad8%7Cef57503014244ed8b83c12c533d879ab%7C0%7C0%7C638091363429706989%7CUnknown%7CTWFpbGZsb3d8eyJWIjoiMC4wLjAwMDAiLCJQIjoiV2luMzIiLCJBTiI6Ik1haWwiLCJXVCI6Mn0%3D%7C3000%7C%7C%7C&sdata=WqmZAJ6hAqeuFQ5Xi3ZafFAPTNNzpnzEsaMQOuU6U2Y%3D&reserved=0))

**Experimental Model and Subject Details**

*Mice****:*** All experiments were performed at Vanderbilt University Medical Center animal facility in accordance with Institutional Animal Care and Utilization Committee (IACUC)-approved protocols and conformed to all relevant regulatory standards. Mice were housed in pathogen-free facilities in ventilated cages with ad libitum food and water and at most 5 animals per cage. Cas9 transgenic (026179) and *Rag1^-/-^* (002216) were obtained from Jackson Laboratory and maintained in VUMC animal facilities. *Bcl11b^fl/fl^* (034469) mice were obtained from Jackson Laboratory and crossed to CD4^cre^ or CD4^cre-ert2^ mice (also from Jackson Laboratory) to obtain Bcl11b^fl/fl^CD4^cre^ constitutive conditional knockouts, Bcl11b^fl/fl^CD4^cre-ert2^ inducible conditional knockouts, and WT littermates. Animals were genotyped for floxed and Cre alleles. To induce deletion of *Bcl11b* in peripheral CD4^+^ T cells of Bcl11b^fl/fl^CD4^cre-ert2^ mice, 6-10 week-old mice were administered 4x 200 µl doses of 20 mg/mL Tamoxifen in corn oil intraperitoneally over 8 days. Peripheral CD4^+^ T cells were isolated 14 days after the first injection. Eight- to sixteen-week-old male and female mice were used for all animal experiments. For induced colitis models, mice were euthanized if human endpoint was reached (weight loss >20%).

**Method Details**

*In vitro mouse CD4^+^ T cell isolation, activation, and differentiation:* Primary mouse bulk CD4^+^ T cells were isolated from the pooled spleen and peripheral lymph nodes (cervical, brachial, axillary, and inguinal) of 6–16-week-old mice using StemCell negative isolation kits. In experiments evaluating polarization, naïve (CD44-low) CD4^+^ T cells were isolated using Miltenyi negative isolation kits. Cells were cultured at 37°C with 5% CO_2_ in Iscove modified Dulbecco medium (IMDM) containing 10% FBS, 100U/mL penicillin/streptomycin, 2mM L-glutamine, and 50 µM 2-Mercaptoethanol (BME). Cells were stimulated with plate-bound αCD3 (3 µg/mL) and αCD28 (2 µg/mL) antibodies for 3 days at 1x10^6^ cells/mL for 3 days unless otherwise specified. In polarization experiments, subset specific cytokines were added as follows: T_H_1: IL-12p70 (10ng/mL), IL-2 (100U/mL), anti-IL-4 (10µg/mL), anti-IFNγ (1µg/mL); T_H_17: IL-6 (50ng/mL), TGFβ (1ng/mL), IL-23 (10ng/mL), IL-1β (10ng/mL), anti-IL4 (10µg/mL), anti-IFNγ (10µg/mL); iTreg: TGFβ (1.5ng/mL), IL-2 (100U/mL), anti-IL4 (10µg/mL), anti-IFNγ (10µg/mL). In cultures lasting more than 3 days, T_H_17-polarized cells were maintained on 1 µg/mL αCD3 and αCD28, while TCR stimulation was removed in other conditions. Media was refreshed and cells split regularly to prevent media exhaustion and overcrowding. Quantification of proliferation during the first 3 days of activation was evaluated by staining cells with CellTrace Violet (CTV) prior to activation. Expansion of cells at later time points were quantified by live cell count determined by Bio-Rad TC20 Cell Counter using Trypan Blue. Where indicated, 2-DG was added to cell cultures at 2 mM, S63845 at 500 nM, and Baf1A at 1 or 5 nM.

*Flow Cytometry and Cell Sorting:* CD4^+^ T cells *ex vivo* and *in vitro* were stained following standard protocols. Briefly, cells were stained with a fixable viability dye and surface antibodies. For intracellular and transcription factor stains, cells were fixed and permeabilized using BD Fixation/Permeabilization Solution Kit. For cells expression a BFP or GFP reporter, an additional fixation step with 1% PFA was performed to prevent loss of signal. For cytokine panels, cells were stimulated with 50 ng/mL 12-myristate 13-acetate (PMA) and 500 ng/mL ionomycin in the presence of GolgiPlug for five hours prior to processing as other intracellular stains. To evaluate T cell mitochondria by flow cytometry, T cells were cultured in standard complete media with 200 nM MitoTracker Green FM (mitochondrial mass), 150 nM TMRE (mitochondrial potential), or 500 nM MitoSox (mitochondrial reactive oxygen species) for 30 min at 37°C and 5% CO_2_ prior to antibody staining. Stained cells were measured using a Miltenyi MACSQuant Analyzer 16 Flow Cytometer and results analyzed using FlowJo v10.6.1. In noted experiments, viable transduced cells were sorted using a FACS Aria III Cell Sorter in the Vanderbilt Flow Cytometry Shared Resource. The Vanderbilt Flow Cytometry Shared Resource is supported by the Vanderbilt Ingram Cancer Center (P30 CA068485) and the Vanderbilt Digestive Disease Research Center (DK058404).

*Pooled CRISPR Library Design and Generation:* The Inborn Errors or Immunity (IEI) sgRNA library was curated by referencing the 2017 IUIS report on the Inborn Errors of Immunity (IEI)^1^. This library includes sgRNAs targeting 312 genes linked with IEIs and 30 nontargeting control (NTC) sgRNAs. The Inborn Errors of Metabolism (IEM) sgRNA library was designed using genes associated with IEM in IEMBase as of January 2019. This library included sgRNAs targeting 1038 IEM-associated genes and 42 NTCs. Genes without homologs in mice, long noncoding RNAs, mitochondrial DNA-encoded genes, and genes linked with specifically GOF mutations were excluded from both libraries. A smaller, targeted IEM library was developed for *in vivo* experiments, consisting of sgRNAs for 48 IEM genes and 8 NTCs. Specific sgRNA sequences in each library were selected by referencing the Mouse CRISPR Knockout Pooled Library (Brie) (Addgene, Pooled Library #73632)^2^. For select IEI and IEM genes not found in the Brie Library, sgRNA sequences from the Gecko v2 Library (Addgene, Pooled Library #1000000052) were used^3^. Four sgRNA sequences for each gene and the nontargeting controls, flanked by the following adapter sequences, were purchased as an oligo pool from Twist Bioscience: GGAAAGGACGAAACACCGXXXXXXXXXXXXXXXXXXXXGTTTTAGAGCTAGAAATAGCAAGTTAAAATAAGGC. The library was further prepared for transduction as previously described^3,4^. Briefly, additional sequences were attached by PCR using Array primers and Herculase II Fusion DNA Polymerase. After purification by gel extraction using QIAquick Gel Extraction Kit, the fragment was cloned into the retroviral expression vector pMx-U6-sgRNA-GFP or pMx-U6-sgRNA-BFP using Gibson Assembly Master Mix. The resultant plasmid pool was amplified by electroporation into ElectroMAX DH10B Cells and plated on ampicillin plates to obtain enough colonies for 50-fold coverage of the library. Plasmids were isolated using GeneJET Plasmid Maxiprep Kit. Pooled CRISPR libraries were transfected into the Platinum-E (PlatE) retroviral packaging cell line using Polyplus jetPRIME DNA and siRNA transfection reagent. Media was changed after 24 hours; viral supernatant was collected after an additional 48 hours.

*CRISPR Screening in CD4^+^ T cells:* CD4^+^ T cells were isolated from the spleen and lymph nodes of Cas9 transgenic mice as described above and stimulated with 3 µg/ml αCD3 and 2 µg/mL αCD28 in RPMI 1640 supplemented with 10 mM HEPES, 50 μM 2-mercaptoethanol, 100 U/mL penicillin/streptomycin, and 2mM glutamine or in polarizing conditions as described above. 48 hours post T cell activation, viral supernatant was spun onto retronectin-treated non-tissue culture plates at 2000 x g from 2 hours at 32°C. Activated T cells were then transferred to these plates and spun for an additional 15 minutes and place back into the incubator. At this time, non-polarized cultures were supplemented with 100 U/mL rhIL2. For screens in T cell polarizing conditions, media was replaced according to polarizing conditions previously described. On day 3 post T cell activation, a sample of cells consisting of ≥1000-fold representation of the library was collected. Cells were expanded in culture for an additional 4 days in 100 U/mL rhIL-2 or under polarizing conditions where noted. In select experiments, cells were treated with 2 mM 2-DG to inhibit glycolysis for the last 48 hours of culture or restimulated with 3 µg/mL plate-bound αCD3 and 2 µg/mL αCD28 for the final 24 hours of culture. Prior to collection of DNA, live cells were isolated via Lympholyte-M (Cedarlane) gradient to avoid contribution of noise from dead cells. In screens evaluating mitochondrial readouts, cells were stained with MitoTracker Green FM, TMRE, or MitoSox as previously described and the top and bottom quintiles of transduced cells for each readout sorted by FACS. In screens evaluating effects on nutrient uptake, transduced cells were starved in Krebs buffer for 1 hour at 37°C before incubation with 50 µM 2-NBDG or C16-BODIPY for 45 minutes at 37°C. The top and bottom quintiles for each readout among viable, transduced cells was then isolated by FACS. Kapa Express Extract Kit. sgRNA sequences were amplified by two rounds of PCR with two technical replicates: first round with adapter primers and second round with barcoded Illumina sequencing primers. The amplicons were then purified by gel extraction, combined at equimolar, and sequenced on the Illumina NovaSeq platform at the Vanderbilt University Medical Center’s Vanderbilt Technologies for Advanced Genomics (VANTAGE) core. At least a 1000-fold representation of the library was maintained throughout entire process to ensure robust coverage and accuracy of results. FASTQ files were analyzed using the Model-based Analysis of Genome-wide CRISPR/Cas9 Knockout (MAGeCK) method for statistical analysis^5^. In screens evaluating expansion and survival, sgRNA abundance was compared between the final, Day 7 sample and the “baseline” Day 3 sample. In analysis of these samples, L_2_FC was normalized to the NTC average to determine if gene loss resulted in an advantage or disadvantage. In screens evaluating the effect on mitochondrial readouts or nutrient uptake, the sgRNA abundance in the top quintile was compared to that in the bottom quintile.

Overenrichment analysis (ORA) of genes depleted in CRIPSR screens was performed using Web-Based gene set analysis toolkit (WebGestalt) to identify pathways critical for CD4^+^ T cell expansion and survival. Depleted genes (-log10(p-value)>1) were mapped against the KEGG database to identify pathways overrepresented in among depleted genes.

*Gfpt1 CRISPR KO:* Single sgRNAs targeting *Gfpt1* and nontargeting controls (NTCs) were cloned into the pMx-U6-sgRNA-GFP or pMx-U6-sgRNA-BFP retroviral vectors as described above in the “Pooled CRISPR Library Design and Generation” section. *Gfpt1* KO CD4^+^ T cells were generated using Gfpt1 and NTC retroviral vectors using the same transduction method detailed in the “CRISPR Screening in CD4^+^ T cells” method. After evaluation of deletion efficiency, Gfpt1-sgRNA sequence GGAGAGAGGAGCCTTAACTG and NTC sequence AAAACGGCTCGATCGGTGAT were used in further single gRNA KO experiments.

*Induction of colitis via CD4^+^ T cell transfer:* To evaluate the intrinsic requirement for *Gfpt1* in CD4^+^ T cells during colitis development, naïve CD4^+^ T cells were stimulated in T_H­_17 polarizing conditions and transduced with either a NTC (GFP reporter) or sgRNA targeting *Gfpt1* (BFP reporter). Transduced cells were isolated by FACS and mixed at a 1:1 ratio. 5x10^5^ total CD4^+^ T cells were adoptively transferred into *Rag1^-/-^* KO mice to induce colitis development as previously described^6^. After colitis development, CD4^+^ T cells from the spleen, mesenteric lymph node, and lamina propria were analyzed to determine the relative frequency of NTC and *Gfpt1* KO cells and their phenotype. To isolate CD4^+^ T cells from the lamina propria, colons were washed thoroughly with PBS, cut into small pieces, and incubated in a predigestion solution of HBSS supplemented with 5 mM EDTA and 20 mM HEPES on a shaker at 37°C. Colon fragments were then digested with Collagenase T3 and DNAseI in RMPI using a gentleMACS dissociator. Isolated cells were then analyzed by flow cytometry.

For *in vivo* CRISPR screens, 5x10^5^ T_H_17 polarized cells transduced with the small target IEM library were adoptively transferred into *Rag1^-/-^* mice to induce colitis. After colitis became apparent, CD4^+^ T cells were isolated from the spleen, mesenteric lymph node, and lamina propria, pooled by organ, and lysed for genomic DNA. sgRNAs were then amplified, sequenced, and analyzed as described in the “CRISPR Screening of CD4^+^ T cells” section and compared to the input sample.

*Immunobloting:* Protein lysates from CD4^+^ T cells were prepared according to standard methods from resting cells or cells stimulated with 3 µg/mL αCD3 and 2 µg/mL αCD28 for 3 days +/- polarizing conditions. Lysates were separated using 10% Bis-Tris Gels, transferred to nitrocellose, and immunoblotted with primary antibodies for Gfat, Nagk, phospho-AMPK (T172), total AMPK, phospho-S6 (S235/236), total-S6, c-Myc, Mcl1, and β-actin. Band intensity was quantified using Image Studio Lite v5.2.

*Extracellular Flux Analyses:* Extracellular flux analysis was performed with the Seahorse XFe96 Analyzer (Agilent). Plates were coated with Cell-Tak solution (Corning) for 30 minutes at room temperature before seeding the cells. 150,000 viable cells were seeded per well and a minimum of five technical replicates were seeded for each sample. Final cell counts of each well were acquired by bright field imaging using a Cytation5 imager for normalization. The Glycolysis Stress Test was performed according to the Agilent protocol recommendations. The Mito Stress Test was performed with oligomycin A at 1.5μM, FCCP at 1.5μM, and rotenone/antimycin A at 0.5μM final concentrations.

*RNAseq of Bcl11b^KO^ T cells:* CD4^+^ T cells from *Bcl11b^fl/fl^Cd4^cre-ert2^* mice and WT littermates were isolated and stimulated after induction of *Bcl11b* deletion and stimulated 3 days with αCD3/αCD28 as described above. Cells were washed with cold PBS and RNA extraction was conducted with DNAse treatment according to the RNeasy Plus Mini kit (Qiagen). mRNA enrichment and cDNA library preparation were performed using the stranded mRNA (polyA-selected) library preparation kit. Sequencing was performed with Paired-End 150bp on the Illumina NovaSeq6000 targeting 50 million reads per sample at Vanderbilt’s VANTAGE core. Alignment percentages for all samples were above 99%. RNA was aligned to the GRCm39 reference genome using **Hisat2**. The resulting reads from these alignment files were then assigned to genes using **featureCounts** to gene annotations from gencode.vM30, giving a raw count matrix. Using the **BioMart** package in **R** the matrix's ensembl IDs were converted to gene names; the **DESeq2** package was then used to find differentially expressed genes between the WT and KO mice from the Stimulated groups. Differential expression analysis criteria was set at an FDR <= 0.05. A heatmap of the normalized expression levels of the top 50 most differentially expressed genes (Fig. S5C) and select group of genes of interest (Fig. S7F) were generated. A GSEA of the HALLMARK gene sets with log2foldchange as our ranking metric was performed using the **fgsea** and **msigdbr** packages (Fig. S5D). The HALLMARK Glycolysis (Fig. S5E) and Hypoxia (Fig. S5F) genesets were subsequently pulled out and made into enrichment plots using the python package **GSEApy**.

*Mitochondrial imaging, quantification, and analysis:* Cells were seeded in poly-lysine and allow to adhere for 30 minutes. Cells were then washed in PBS and fixed in 4% paraformaldehyde for 20 minutes at room temperature followed by permeabilization in 1% Triton-X-100 for 10 minutes at room temperature. After blocking in 10% BSA, immunostaining was performed with anti-HSP60 (1:200; Cell Signaling Technologies; 12165S) and anti-rabbit IgG Alexa Fluor 488 (1:200; Invitrogen A11034).

Super-resolution mitochondrial images were acquired using a Nikon SIM microscope equipped with a 1.49 NA 100x Oil objective and Andor DU-897 EMCCD camera. Quantification of mitochondrial morphology was performed in NIS-Elements (Nikon) as described previously (Rasmussen and Taneja et al., 2020 (PMID: 32283523), Romero-Morales et al., 2022 PMID: 35792828))^7,8^. Mitochondria were segmented in 3D and skeletonized using the resulting binary 3D mask. All mitochondria within each cell were averaged resulting in one data point per cell. GraphPad Prism was used to perform student’s t-test for statistical analysis.

**Statistical analysis***.*

Statistical analyses were performed with Prism software (v9). Statistically significant results are labelled (* p < 0.05, ** p < 0.01, *** p < 0.001, **** p ≤ 0.001). For a comparison of two groups, student’s t-test was performed. For more than two groups, One-way ANOVA was performed. Figures with data points connected by lines are indicative of paired analyses, whereas all other statistical tests were unpaired. Error bars show mean ± standard deviation unless otherwise indicated. All experiments were carried out in at least triplicate biological replicates unless otherwise stated. FACs plots shown are representative and correspond to bar blots describing data of *n* = 4-5 biological replicates.

**Figure Generation.**

Figures were assembled using GraphPad Prism. Schematics of CRISPR screening protocols and the hexosamine biosynthetic pathway were generated using BioRender under license.
