## Supplemental Tables for "Functional Overlap of Inborn Errors of Immunity and Metabolism Genes Define T Cell Immunometabolic Vulnerabilities"

**Supplemental Table 1. IEI genes and sgRNAs in the Inborn Errors of Immunity Genes Library.**

| **Gene Target** | **sgRNA Target Sequence** |
| --- | --- |
| Acp5 | ATACCAGGGGATGTTGCGAA |
| Acp5 | TCCACGTACAAACATAACTG |
| Acp5 | TACCTGGAACCTCTTGTCGC |
| Acp5 | ACAGCCACAAATCTCAGGGT |
| Actb | AATGCCTGTGGTACGACCAG |
| Actb | ATGGAGGGGAATACAGCCCG |
| Actb | GGACTCCTATGTGGGTGACG |
| Actb | CAGCACAGGGTGCTCCTCAG |
| Ada | GTTGTGGATCTTGTGAACCA |
| Ada | ATTCATCGGACCGTCCACGC |
| Ada | CTTCATCTCCACAAACTCGT |
| Ada | GCTGCGCAACATTATCGGCA |
| Adam17 | GGTGTGTGGCAACTCCAGGG |
| Adam17 | ACACGTCGTGGGATAATGCA |
| Adam17 | CATCGACGTACGGCACACAC |
| Adam17 | GCCCCAAATGAGGACCAAGG |
| Cfd | TCTCACGTGGGGACCCAACG |
| Cfd | CCCCGAGGCCGGATTCTGGG |
| Cfd | TTGACACTCTGAGTTGATGC |
| Cfd | CCCCTGAACCCTACAAGCGA |
| Aicda | GTAGGAACAACAATTCCACG |
| Aicda | TTCACAGAAGTAGAGGCGCG |
| Aicda | ACCAGGTGACGCGGTAACAC |
| Aicda | TGAGACCTACCTCTGCTACG |
| Aire | TGTGCCGTGTGCCACGACGG |
| Aire | CTCTCCAGGAATTCAGACCA |
| Aire | ACAGAACCTGTCCCAGCCTG |
| Aire | GGTAGAGATGAGCAGAAAGT |
| Ak2 | TGAAGGCGACAATGGATGCA |
| Ak2 | GGTAGGACCGGCCACTCTTG |
| Ak2 | TCCGAACCGGAGATTCCGAA |
| Ak2 | TCAGCCAGTTTGGGTGCCTG |
| Ap3b1 | TGTGGCCAGTAAAAACATCG |
| Ap3b1 | GCTGCCTAATAAATCGTGTG |
| Ap3b1 | ATATGCTAACAAGATACGCT |
| Ap3b1 | TTGGCACATCTCACCCAAGT |
| Ap3d1 | CCAACGCAATGGTATCTGTG |
| Ap3d1 | AGCACATCACCAACTTCGAG |
| Ap3d1 | TCTTCACAGTAAAGTACCTG |
| Ap3d1 | TGTCCTCATCGCTCTCCGTG |
| Xiap | TTTCAGACACCATATACCCG |
| Xiap | AGCACTAGCTAACTCTCTGG |
| Xiap | CTTGGGAACAGCATGCGAAG |
| Xiap | ATGGACATCCTCAGTTAACA |
| Arpc1b | CACAATGCGGTTACTCTCAG |
| Arpc1b | CAATGAGAACAAGTTCGCCG |
| Arpc1b | CCAGTCCAGGCTGAGCACTG |
| Arpc1b | GAAGCGAGCACTCACGTGTA |
| Rab27a | TGGTTAAGCTACGAAACCTA |
| Rab27a | AGTGTACTGGTAGAGTACAC |
| Rab27a | AACCCAGATATAGTGCTGTG |
| Rab27a | CCTGAAATCAATGCCCACTG |
| Atm | TAAGTCATATAGGAAGCCGA |
| Atm | GAGTATAAATAACATCGCGA |
| Atm | AAGACTTGAACACCGGACAA |
| Atm | TGCAAGATACACATGAATCG |
| B2m | ATTTGGATTTCAATGTGAGG |
| B2m | ACTCACTCTGGATAGCATAC |
| B2m | TGAGTATACTTGAATTTGAG |
| B2m | TCGGCTTCCCATTCTCCGGT |
| Bach2 | TGGACAGACGAAAGATGACT |
| Bach2 | AATTACGGACAGCCCCACGT |
| Bach2 | CTCCTCGTATTCCTACGCAG |
| Bach2 | TCTCTGTTCGGTATAACGAA |
| Bcl10 | CTCCGGGTGGTACATGACAG |
| Bcl10 | GATTCAGAAGATAACGGATG |
| Bcl10 | ATAAAACTGGAGCACCTCAA |
| Bcl10 | CCTGGTGGAATCCATCCGCA |
| Blm | TTACCTGGAACATTTCAACG |
| Blm | GGTGGGTAAACATTCCTCAG |
| Blm | CCTGCAAGTGGATTTAACGA |
| Blm | CCTTCACCGACTTACACCTG |
| Btk | AATCCGGTACAATAGTGACC |
| Btk | TATGAATATGACTTTGAACG |
| Btk | TGGAGGAGAGCAACCTACCG |
| Btk | AGATTTAGCAAACACAGACA |
| Serping1 | GCTCTGAGATGCATTCACAT |
| Serping1 | ACAATAACAAATGACACCAT |
| Serping1 | AGAACTCATCAACACCTGGG |
| Serping1 | GGACACAGGCAAAATCCTTG |
| C1qa | CACAGATGAAGCGACCCGTG |
| C1qa | CCCAATGACGCTTGGCAACG |
| C1qa | TTCAGCCACTGTCCATACTA |
| C1qa | AGGCAATCCAGGCAATATCA |
| C1qb | AGGCACTCCAGGGATAAAGG |
| C1qb | TGACCTGGTTCGGTCGCAAG |
| C1qb | GAATCGCCTTTGGGACCGCG |
| C1qb | CAGGTGAACTTGCCGTTGCG |
| C1qc | GGGTGACTGTGAATACCGAC |
| C1qc | CTGTGGTCACCAACCCTCAG |
| C1qc | ATCATGCCCGTCCTTCCCAG |
| C1qc | GCTCCCCCGGAGGCCCCCTG |
| C2 | GTTCTAGGAAAGTCCAACAT |
| C2 | GTGTGATGTGAGCTAGACCT |
| C2 | AGTCAATACCATATTTGAGG |
| C2 | TTGGGGCGACAGTACCGCAC |
| Ciita | AGCTCGACTAAGGCTCCGGG |
| Ciita | AGGTCCTTGATTATATCGTG |
| Ciita | TCCAGTGTCCTAATCTACCA |
| Ciita | AGCAGGCCAAGACTTACATG |
| C9 | CTACAACGGACTCTGTGACC |
| C9 | TTTCCCATTAAACTGTCCGA |
| C9 | ATTCTGCAGTCTATCGGTAT |
| C9 | TTGAGAGGAGTGGTCCAACT |
| Hyou1 | ACAGGCGGATAACCCTCATG |
| Hyou1 | ACATCGTACTCACTTGCCCA |
| Hyou1 | TGGAATTGATATCTTTCCGG |
| Hyou1 | TGGCGTGCTCAGTTTAGACA |
| Casp8 | GATTATGAAAGATCAAGCAC |
| Casp8 | CTTCCTAGACTGCAACCGAG |
| Casp8 | ATGATCAGACAGTATCCCCG |
| Casp8 | CAAGAAGCAGGAGACCATCG |
| Ctla4 | TGTGATGGTGAATATTCACA |
| Ctla4 | GGACTGAGAGCTGTTGACAC |
| Ctla4 | ACAGGTGACCCAACCTTCAG |
| Ctla4 | TGCCCACAAAGTATGGCGGT |
| Cd19 | GAATGACTGACCCCGCCAGG |
| Cd19 | AATGTCTCAGACCATATGGG |
| Cd19 | GGCACCTATTATTGTCTCCG |
| Cd19 | TTTAGCCCACACATACAGCT |
| Ms4a1 | GTTACAGTACTGTGTAGATG |
| Ms4a1 | TACCATACACTCAAACAGAT |
| Ms4a1 | CAGTCGTAGATATCAACATA |
| Ms4a1 | CCACACAAAGCTTCTTCATG |
| Cd3d | TTACACAGATATATCCCTCG |
| Cd3d | TCCATCTAGATGCATGACGC |
| Cd3d | AAGAATAAAACACTCAACTT |
| Cd3d | GATACAAGTGACCGAATATG |
| Cd3e | AGGGCACGTCAACTCTACAC |
| Cd3e | TTCTCGGAAGTCGAGGACAG |
| Cd3e | TACTTGTACCTGAAAGCTCG |
| Cd3e | TCAGAAGCATGATAAGCACC |
| Cd3g | TGACACTGATACGTGCCTCG |
| Cd3g | TTCTGTAATACACTTGCAGG |
| Cd3g | AACTGCATTGAGCTAAACAT |
| Cd3g | GTACAAGTGGATGGCAGCCG |
| Cd247 | GCTCGGGATCCAGAGATGGG |
| Cd247 | GCTCAATCTAGGGCGAAGAG |
| Cd247 | CTCCTGGGAACCGCACGTGG |
| Cd247 | CATTGTATACGCCTTCCTGG |
| Cd79a | GGAACCCTAATATCACATGG |
| Cd79a | CCTACTCACTGCGCACGCGG |
| Cd79a | CGAAGTAAACAAGAACCACA |
| Cd79a | AGGCGTATGACAAGAAGAGG |
| Cd81 | GCAACCACAGAGCTACACCT |
| Cd81 | GGGCTTCGTAAACAAAGACC |
| Cd81 | ATCCATCACAGCTTGCTGAA |
| Cd81 | GGCAAACAGGATCACAAGGC |
| Cd8a | TGGGTGAGTCGATTATCCTG |
| Cd8a | ATCCCACAACAAGATAACGT |
| Cd8a | GTGTTGGGGTCCGTTTCGCA |
| Cd8a | GGACGCCGAACTTGGTCAGA |
| Cfh | ATTTACGCATAACTCCACCA |
| Cfh | CAAGTGTTCGGTATCCAGGG |
| Cfh | GAAATTGATTACCGTGAATG |
| Cfh | CATATTTCACCACTTCACCA |
| Cfi | CTTGTGGATTACCTTCACAA |
| Cfi | TGTTATTACACAGGTTGCCG |
| Cfi | AAGGTCGACGCAGGCCACGT |
| Cfi | TGCCTGCATGTACATTGCCG |
| Cftr | GCCGTGTGACTGACATACGT |
| Cftr | TTCTAACTGAGACCTTACGC |
| Cftr | GTGGCGATCATGTTGCTGCG |
| Cftr | TATGGAGAGTAAAATATCGT |
| Coro1a | TGACACGAACTCGCTTGTCA |
| Coro1a | ACCAGGCGATGTCTAGCACA |
| Coro1a | AGAAGGGAAGATTCTAACCA |
| Coro1a | GCCACAGAGGTAGACAATGT |
| Tpp1 | TGAGTTTCATCGCTATGTAG |
| Tpp1 | TTATGGTAGAAGGTTACCTG |
| Tpp1 | AACCTGACAGCCAAAGATGT |
| Tpp1 | GATCGAGGCCAGTCTAGATG |
| Copa | CTTGCGGTTACAGATCACAA |
| Copa | CTCTCTCTGACACATCACCT |
| Copa | CACCAGAAATATCCCAAACG |
| Copa | GAATGTCTAGGTACTCCACA |
| Cr2 | ATGTTGACCAGTTTGTTGCG |
| Cr2 | CTATACATTGCACCCCTGAG |
| Cr2 | AGACGGATTTCTATAAACCA |
| Cr2 | ATGCAATGCTCATGGCACAT |
| Csf2ra | AGGACGCGGTGACGTCACGT |
| Csf2ra | CCTACTTGGTCGTGACCGGT |
| Csf2ra | CTAGCGTCACTAACCCAGAA |
| Csf2ra | TGACATCCAGCGTGACACCG |
| Csf2rb | TACACTTGGAAGACTGACTG |
| Csf2rb | TGATGGAAAATCGTGTATAG |
| Csf2rb | GAAGAAATCGGACAGCTGGG |
| Csf2rb | TGGAGACTGTAGGCATCCTG |
| Csf3r | TCAGCTCTACAGAATTACAG |
| Csf3r | TGAGGCAGGATAGGTTTGAG |
| Csf3r | TGAGCTGCGTGGTGTTGCAA |
| Csf3r | CTTTGCCACACAATCCGGGA |
| Ctsc | CTGCAAGATACAACCTCCTG |
| Ctsc | GGCAGTAACTGATAGCTGTG |
| Ctsc | TCACAACCACAACTTTGTGA |
| Ctsc | CTGTATTTCATCAGTCATCG |
| Cyba | AGTAGAAGGATACATAGAGT |
| Cyba | AATGACTTACCATCGCTCCA |
| Cyba | GGACGTAGTAATTCCTGGTG |
| Cyba | CAGATAGATCACACTGGCAA |
| Cybb | CCTCTACCAAAACCATTCGG |
| Cybb | TTTACCAGACGAATTGTACG |
| Cybb | ATTCTAACTTGGATACCTTG |
| Cybb | TTATACTCGAAAACTCCTTG |
| Cd55 | CTCTTATACGTATAGCCAGG |
| Cd55 | CTGCTGTCCCCAACTGTACG |
| Cd55 | TGTGACAGAACAGAAAGTAG |
| Cd55 | CGAAAACAACCTCCACTCCC |
| Dnase1l3 | TCTCGGGAGTTGTGTGCAAG |
| Dnase1l3 | TCACACTCACTTGTAGACGA |
| Dnase1l3 | TTCATGGGTGATTTCAACGC |
| Dnase1l3 | CACTACCATGACTATCAGGA |
| Dnase2a | TACACCGTCTTGCCAACCGT |
| Dnase2a | AGGCCAGGTGTATGCACCAG |
| Dnase2a | CGAAGCCCTGAGCTGCTATG |
| Dnase2a | GAAATTACCTGACCTAGAGA |
| Dnmt3b | TGGTAGCCGGAAACTCCACA |
| Dnmt3b | CAGCCTTCTGAATTACACGC |
| Dnmt3b | GGAGGGTATGGATACCACAC |
| Dnmt3b | CAGTAGGCTTGAAGCCACCG |
| Fadd | TAGATCGTGTCGGCGCAGCG |
| Fadd | AAGCTGGAGCGCGTGCAGAG |
| Fadd | TTCGTTTGCTCACGCGCTCG |
| Fadd | GCGCCTGGACGACTTCGAGG |
| Fas | CAGTTAAGAGTTCATACTCA |
| Fas | TATTTATATATCGAAAGTAC |
| Fas | CATTTGCATACTCACACGAC |
| Fas | GAGGACTGCAAAATGAATGG |
| Fasl | AGGACCACAACACAAATCTG |
| Fasl | CTTCACTCCAGAGATCAGAG |
| Fasl | CCTCTGAAAAAAAAGAGCCG |
| Fasl | GGAACTGGCAGAACTCCGTG |
| Fcgr3 | TGGTGACACTGATGTGCGAA |
| Fcgr3 | ATGCACACTCTGGAAGCCAA |
| Fcgr3 | TGGTGAAACTGGACCCCCCA |
| Fcgr3 | TGCTGCTCCAGACCCCTCAG |
| Fpr1 | AGAAGGTAATCATCGTACCC |
| Fpr1 | TGGTGACAGTGTGTTTCATG |
| Fpr1 | GGCAATGTAAAATGGCAAAG |
| Fpr1 | GATGCAGAACACAAATACAG |
| G6pdx | AGAGGTGGAAACTGACAACG |
| G6pdx | TGCCCGCTCACGACTCACAG |
| G6pdx | ATGACCCCACAGTACCCCAT |
| G6pdx | AGAGATGGTCCAGAATCTCA |
| Slc37a4 | CTACGTTGACCAGACCAACC |
| Slc37a4 | TCTTTACTCCGAAGACCACG |
| Slc37a4 | CACAAACTTGCTGATGGCGT |
| Slc37a4 | CAGAGCGATCTCATCCACCA |
| Gata2 | CCTGGGCTGTGCAACAAGTG |
| Gata2 | GGCCGGGAGTGTGTCAACTG |
| Gata2 | ACAGCTGCTGCCTCCCGACG |
| Gata2 | GGCACATAGGAGGGATAGGT |
| Gfi1 | CAAATGCATCAAATGCAGCA |
| Gfi1 | CCCCGACTCTCAGCTTACCG |
| Gfi1 | CTGCACGCGGACAAGAGCGT |
| Gfi1 | CTACGGCGACTTCGCGCCTG |
| Ostm1 | CATCAGCCGAAACATCGGGG |
| Ostm1 | TTGCCTAACAAACAATGGTG |
| Ostm1 | AGCGCTGCGCACCATACAGG |
| Ostm1 | CAGAATGCAGATAGTTCTCA |
| Hc | GCAGATGACTCCCATTATCG |
| Hc | AGACAAACCTGTTTACACGC |
| Hc | ACCCTAAGGGAATTCGTGGT |
| Hc | GAACCTCCCATCAAATGTGA |
| Hells | TCTGCGGTACCGAATTTCAG |
| Hells | ACGGTCATTAAAACTTACAG |
| Hells | GCTGTATCATGGAACCCGGG |
| Hells | ACAACTACTTCTCGCTTAGG |
| Foxn1 | CCAGGTACTTACGTTCTGTG |
| Foxn1 | GAACAGTACATATGTTGCAA |
| Foxn1 | CACCTCACTATCCCTATCAG |
| Foxn1 | CCACAGACAGCCCTTTCGAG |
| Hmox1 | TCGTGCTCGAATGAACACTC |
| Hmox1 | TCAGGACCTGACCCCCTGAG |
| Hmox1 | ACGCTTTACATAGTGCTGTG |
| Hmox1 | TTCCTTGTACCATATCTACA |
| Irf8 | AGTTTACCGAATTGTCCCCG |
| Irf8 | TCGACAGCAGCATGTACCCG |
| Irf8 | GCGTAACCTCGTCTTCCACG |
| Irf8 | CAAGCAGGATTACAATCAGG |
| Ifnar2 | CAAAGACGAAAATCTGACGA |
| Ifnar2 | GCCATCGTCATAGTGCACAG |
| Ifnar2 | TAACCTGGATAATCCCTGAA |
| Ifnar2 | ACTGTGGAATTACGATTATG |
| Ifngr1 | TATGTGGAGCATAACCGGAG |
| Ifngr1 | GGTATTCCCAGCATACGACA |
| Ifngr1 | TTCAGGGTGAAATACGAGGA |
| Ifngr1 | TATACCAATACGCAAATACC |
| Ifngr2 | AGGGAACCTCACTTCCAAGT |
| Ifngr2 | TGGACCTCCGAAAAACATCT |
| Ifngr2 | TCCCTTTGATGTGTTCCACG |
| Ifngr2 | TGATGAGCAGATTCTAACTT |
| Cd79b | TGACCTGGTTCCGAAAGCGA |
| Cd79b | TCCCCCAGGATTCAGCACGT |
| Cd79b | CATAATGTCACCGACAGCTG |
| Cd79b | CGAGGTTTGCAGCCAAAAAG |
| Igll1 | CTGTTCCAGATCATCCCACG |
| Igll1 | GCTCACCAAACACACTAGTG |
| Igll1 | CCAAAGACATACCAAAACTG |
| Igll1 | GGAGAACTTCACACTGCCTG |
| Ikbkb | TCACACATACCCCGTGACGG |
| Ikbkb | CAAGATCCATGTCCAACGTG |
| Ikbkb | ATGTGGCACCCTCGGCAAAG |
| Ikbkb | GAAGCCAGTGATGCACTCGA |
| Ikbkg | AAGGATCGGCAAGCTTTAGA |
| Ikbkg | AGGCTGCCTTGCGAATGGAG |
| Ikbkg | TGGGTGAAGAATCTTCTCTG |
| Ikbkg | GCTCAGGTGACATCATTGCT |
| Il10 | GCTAACCGACTCCTTAATGC |
| Il10 | AACTGCACCCACTTCCCAGT |
| Il10 | AAGGAGCATTTGAATTCCCT |
| Il10 | ACTGGCATGAGGATCAGCAG |
| Il10ra | GCAGTGTTTACTTATCACGA |
| Il10ra | GTGGGGACAACACGGACAGT |
| Il10ra | GGTGAACGTTGTGAGATCAC |
| Il10ra | TCTGGCTTCAAACCACACAT |
| Il10rb | TGGCGGATGAACATTCGGAG |
| Il10rb | ATTTCAAGAACATTCTACAG |
| Il10rb | CGGAGGACCTCAGAGTCGTA |
| Il10rb | AGAGAAGTCGCACTGAGTCG |
| Il12rb1 | GTCGGTGAGGAACCAAACCG |
| Il12rb1 | CCTCCGAACCATACCCACAC |
| Il12rb1 | TGAGAAGACATCGTTCCCAG |
| Il12rb1 | TCAAGGTGTCACAATCACAC |
| Il17ra | ACTGAAGTAGCAAACAACGT |
| Il17ra | ATGAGGCCATACACCCACAG |
| Il17ra | GAAGGTCTGGATCGTCTACT |
| Il17ra | GACCTGGAGATGTTTGAACC |
| Irak1 | ACAGAAGGGCCAGCAAAACG |
| Irak1 | TGGGCAAGAAGCCATAAACA |
| Irak1 | GGTGCGTGAGGATGTGAACG |
| Irak1 | CCCACCGAACTGGCACCAGT |
| Il1rn | TTCTCCAGAAAAGATAGACA |
| Il1rn | GTGTTCTTGGGCATCCACGG |
| Il1rn | CTGCCTCTGAATGAAACAGA |
| Il1rn | CTTGATATCATCTCCAGACT |
| Il2ra | GTGTCTGTATGACCCACCCG |
| Il2ra | ATCTTGCAGATGCTAATAGC |
| Il2ra | GAGAGGTTTCCGAAGACTAA |
| Il2ra | GAATCTTCATGTTTCCAAGG |
| Il2rg | TTCCACAGATTGGGTTATAG |
| Il2rg | GGAGCAACAGAGATCGAAGC |
| Il2rg | CCCGATTACCAAGATTCTGT |
| Il2rg | CATACCTATAGTGCAGCGTG |
| Il7r | GGGAGACTAGGCCATACGAC |
| Il7r | AGACCTAGAAGATGCAGACG |
| Il7r | CAGAACCCAAGAATCAAGGT |
| Il7r | CCTTTGAAGTAATCGTTATG |
| Itch | TCCCGCACATAGGCTATCTG |
| Itch | AAAACATAATTTACTCGTAG |
| Itch | GCCAAGCTCCCCTACCACCT |
| Itch | ACAACACTCGGATTACTCAG |
| Itgb2 | GAGTATAGGCAAATCCCGTG |
| Itgb2 | TTGGCTGGCGCAATGTCACG |
| Itgb2 | ATCCTGAGTTCGACCAACGG |
| Itgb2 | TGGAGTAGGAGAGATCCATG |
| Itk | CAGCCCCAAGCGCTACTACG |
| Itk | GTAAGCCTTCTCATACCCCG |
| Itk | TCAGGAACCTGAAGAAACCC |
| Itk | TTGCTCCAGACTGTGAGAGT |
| Jak1 | CGATGCCATTCGAATGACAG |
| Jak1 | TGAATAAATCCATCAGACAG |
| Jak1 | TCCGAACCGAATCATCACTG |
| Jak1 | AAACATATAGTGTACCTCTA |
| Jak3 | CAGACATCGGAACTGCATGG |
| Jak3 | AGAAAAGTCCAATTTGATCG |
| Jak3 | GGGCTTTGGAGCCACCACGT |
| Jak3 | CACCACCGAGACCTTCCGTG |
| Rpsa | GAAGCGGCCAGCGATCGGAG |
| Rpsa | CTTACAGGGAGCTCACTCAG |
| Rpsa | GACGTCAGCAGGATTCTCGA |
| Rpsa | AGGGAAGTACTCCGCATGCG |
| Lck | AAGATCCGTAACCTAGACAA |
| Lck | GTACTACAACGGACACACGA |
| Lck | CCGGGAAAGCGAAAGCACTG |
| Lck | GCTTTATGCAGTGGTCACCC |
| Lig1 | GACCTCACTGACACCCCGAG |
| Lig1 | GACAATTCAAGACACTCTTG |
| Lig1 | TCTTCTCATAGAATTTCCCA |
| Lig1 | AGAGTGATTCTCCAGTGAAG |
| Psmb8 | CCGGAGCTCGCACTTCCCCG |
| Psmb8 | ACATGATGCTGCAGTACCGG |
| Psmb8 | CTCGCCTTCAAGTTCCAGCA |
| Psmb8 | AGGTTGTATTATCTTCGGAA |
| Blnk | CTATGCAGGTCGAAACAGTG |
| Blnk | AAGATAATCGATCCAGCCAG |
| Blnk | GTCTGTGACTAGACCCTCGG |
| Blnk | GATATTAAGAACAATGAAGG |
| Lyst | GGAGCCCTGAAAGATCGCGG |
| Lyst | GGGATGAGTCTTACCCACGT |
| Lyst | TAGGAAGTGGTGAACCACGT |
| Lyst | CATACCCCGAGTTTAAGCAA |
| Masp2 | GGCCTGTGATATAGTCCACA |
| Masp2 | CCTTTGCCACTGACGAGTCG |
| Masp2 | CTGCGAGTATGACTTTGTCA |
| Masp2 | TGTGGCAATAATGGTCACAA |
| Mcm4 | TTGTTGACAAGGTTCAACCA |
| Mcm4 | ACCCGGGTGGAGATAGATCG |
| Mcm4 | GGTTATACCAACCTTTGACA |
| Mcm4 | CTCAGGGTCTTTCATCACAT |
| Cd46 | CTGTGAGCCAAATCATACAT |
| Cd46 | TCCATAGCTTCAAATGGCCG |
| Cd46 | GGCCTCAGCATAGACGCCCA |
| Cd46 | ACAGTTGTGATCCTACCCCA |
| Msh6 | AGTTTCGTGATACCAAACAG |
| Msh6 | CATCAGTGACCGTCTAGATG |
| Msh6 | TAAGTAAAGACACGGCTGAG |
| Msh6 | GAGGCAAAGGATCTCAACGG |
| Msn | CGCTTGTTAATCCGAAGCCG |
| Msn | CCGGGCCAAGTTCTACCCAG |
| Msn | GAAGCAACTTATCTCCAGCC |
| Msn | CAGACTAAGAAGGCTCAGCA |
| Mvk | AGCGTCAATTTACCCAACAT |
| Mvk | GTGGTCGGAACTTCCCCCCG |
| Mvk | CAAGGTCCCGCGGAGTACCA |
| Mvk | TCTGAAGTCAATCAACAAGT |
| Myd88 | GGTTCAAGAACAGCGATAGG |
| Myd88 | AAGGAGCTGAAGTCGCGCAT |
| Myd88 | CCTGTCCTCAGGACAAACGC |
| Myd88 | GCATCCAACAAACTGCGAGT |
| Ncf1 | TCTTCACGGGCAGTCCCATG |
| Ncf1 | GATCCGGATCCCAACTACGC |
| Ncf1 | GCTACGCACTGGCTGTCAGT |
| Ncf1 | CGTTGCCCATCAAACCACCT |
| Ncf2 | GCACAAAGCCAAACAATACG |
| Ncf2 | TGGAATATCGGATTCTGGAG |
| Ncf2 | TCTCACCTGTGGCTGCAGAG |
| Ncf2 | TGAAGCAAATCCTCGAGTGG |
| Ncf4 | AGAAGAAGATCCTGACGTCG |
| Ncf4 | CGTAGAACTGGCGATAGCGG |
| Ncf4 | CGATCCATGATGGCCCCTTG |
| Ncf4 | ACCTCGATGACAAAAACCTG |
| Nfkb1 | TGTGAAGGCCCATCACACGG |
| Nfkb1 | GGAGTCACGAAATCCAACGC |
| Nfkb1 | TTTCGACTACGCAGTGACGG |
| Nfkb1 | ATGACAGAGGCGTGTATTAG |
| Nfkb2 | ACTGAGCGTGATAAATGACG |
| Nfkb2 | CGGAACACAATGGCATACTG |
| Nfkb2 | CCCACGCTGCTGGATCGGCA |
| Nfkb2 | ACCCGGATAGCAGCCCATTG |
| Pepd | CACAAATCGGATCTCCAGCG |
| Pepd | CTGCTATGGTGTCATCGATG |
| Pepd | GCCCTGCAACACGACAGCTG |
| Pepd | CTTGCTAATGCCCTCGAAGG |
| Cfp | GTTACACATTTGGTTCCGAG |
| Cfp | TATGAGCATAAGGCCTGCAG |
| Cfp | GATTATCACATACTCGTTGA |
| Cfp | CCTACTTGGGAGAGACATCA |
| Prf1 | GTTCGTGCCAGGTGTATGGA |
| Prf1 | TGCCACAGGTAGGCGCTGTG |
| Prf1 | TCAATAACGACTGGCGTGTG |
| Prf1 | GGTAGGAGACTGCCTGAACG |
| Pik3r1 | GAGCTTTATAAGGAGAGGCG |
| Pik3r1 | TCCATTAACCTTCAACTCTG |
| Pik3r1 | TGGCTACAATGAAACCACTG |
| Pik3r1 | CTGGAAATCTGAAAAGCACG |
| Prkcd | AGAAGGTGGCGATAAACTCG |
| Prkcd | TTATAAACCTTGAATCGGTG |
| Prkcd | AGCCCACCATGTATCCTGAG |
| Prkcd | CGATGATGTAGAGTGTACCA |
| Pms2 | TCTCAGGAAACCATAAACTG |
| Pms2 | GTAAGCTGCACTAATCAGCT |
| Pms2 | AGTTTCAGACAATGGATGTG |
| Pms2 | ACTAAAGAGATCAAGTCTAG |
| Pnp | AGATGCTGTGTGATGATGCA |
| Pnp | TCAGTGCCTGGAAACAAATG |
| Pnp | TGTGGCCAGAACCCTCTCCG |
| Pnp | CCTCAAGTGGCAGTGATCTG |
| Pola1 | CTAACGTTTACCATTTCACG |
| Pola1 | AAGAAAGCAACTTACGCTGG |
| Pola1 | AAAGAAAAAAAGATCTACTG |
| Pola1 | TGTACAGAACCATCAACATG |
| Pole | CATTATGGTCACCTACAATG |
| Pole | GATGCTGAGACCTACGTCGG |
| Pole | CATTGACCTCAGAATCCATG |
| Pole | AGGCTGGAAGGATCATAGCA |
| Pole2 | GATCGAACGATCTGTCGTGG |
| Pole2 | CCACAGTGGCTTATACACCG |
| Pole2 | TCAAGCGCTTTAATACCTTG |
| Pole2 | ATAGCGTTCAAGATACAGCT |
| Prkdc | AGAGCCAATTCAGTGACCCG |
| Prkdc | TGGCCCTTGTAAGTAGACGA |
| Prkdc | CATGCAGGGTAAGTAATCGT |
| Prkdc | ACAGAGGATGCTCAAAAATG |
| Psen1 | TTTCAACCAGCATACGAAGT |
| Psen1 | TGCTACTGTAACGTAGTCCA |
| Psen1 | CTGAGCCAATATCTAATGGG |
| Psen1 | AGTCAGCTTCTATACCCGGA |
| Pstpip1 | CTGGTTACAGTGCACCCACA |
| Pstpip1 | TGTTTGTAGAAAGAGTGTAC |
| Pstpip1 | ATTGGCACTCACACGCTCGA |
| Pstpip1 | GTCGCTCTACAAGAAGACCA |
| Pten | CCTCCAATTCAGGACCCACG |
| Pten | TGTGCATATTTATTGCATCG |
| Pten | ACTATTCCAATGTTCAGTGG |
| Pten | GGTTTGATAAGTTCTAGCTG |
| Ptprc | GTAGCAGAAATCTTATATCG |
| Ptprc | TTTCACAATGGAGTGTACGA |
| Ptprc | TTGTCAAGCTAAGGCGACAG |
| Ptprc | ACCACAACGAAGCAAACATG |
| Rac2 | GAGAAGACACGTCTTGCCCA |
| Rac2 | GCTGGACCTTCGCGATGACA |
| Rac2 | AAGAAGCCACTCACACAGTG |
| Rac2 | GGCACGGACATTCTCATAGG |
| Rag1 | ACACCAAAGCAGAGTCGTAG |
| Rag1 | TGAAACGATTCCCACAGATG |
| Rag1 | TCCCGCGCAAGATTGCAATG |
| Rag1 | TGGGAAGTAGACCTGACTGT |
| Rag2 | CATCAATATATCATTCACGG |
| Rag2 | ATTGACGTGGTGTATAGTCG |
| Rag2 | TAACTTGTATAGAATAAGAG |
| Rag2 | CATACCAGGAGACAATAAGC |
| Ranbp2 | AAATTATTCTCGTCACAAAG |
| Ranbp2 | TGCTAATGTAACTCCCACCA |
| Ranbp2 | ATGTTGTTAAACTTAAGTCG |
| Ranbp2 | ACATGCAATGCACCTAGAGA |
| Rasgrp1 | ACGTACAGATATCCGTCGGA |
| Rasgrp1 | TAAGAACTATGATCTCGACC |
| Rasgrp1 | ACAGTTGGTTATTCCGACAC |
| Rasgrp1 | TGACCTTATTGATCTCATGT |
| Relb | GTTCAAAACGCCACCCTACG |
| Relb | TACACCCACATAGCCTCGTG |
| Relb | GCGGATTTGCCGAATCAACA |
| Relb | GCCTCCTATCGGGACCAGCA |
| Rfxank | GTTCTACTTACAGTCCAGGG |
| Rfxank | ATCAACAAACCGGATGAGCG |
| Rfxank | GCACTGTCACTTGCCAGTAT |
| Rfxank | AGTTCGCTTCCTGCTAGACT |
| Rorc | CTTGAGTATAGTCCAGAACG |
| Rorc | GTCATCTGGGATCCACTACG |
| Rorc | TCTGGGGCACTGCAGAAACT |
| Rorc | GACAAGCAGAGGCCTCGGGT |
| Sema3e | ACACGATCTACACCCGAGTG |
| Sema3e | TGTGCCAGCAAAGTAAACGG |
| Sema3e | AAGATAATTACCAACTAGCG |
| Sema3e | TGTGATACACACATACAGCA |
| Foxp3 | CATACCTGATGCATGAAGTG |
| Foxp3 | TCTACCCACAGGGATCAATG |
| Foxp3 | AGGTCGGGACCTGCGAAGTG |
| Foxp3 | GCAAGAGCTCTTGTCCATTG |
| Sh2d1a | AGAAGCTCTTACTCGCTACC |
| Sh2d1a | GATGCAGTGACTGTGTACCA |
| Sh2d1a | AACAGGTTCTTGGAGTGCCG |
| Sh2d1a | CACACAGGCAGTACACGCCA |
| Clpb | AGGACCGCGTTCCGACGAGG |
| Clpb | GGAGAACGGCTGGTACGATG |
| Clpb | CGTTGTCACCGGAGACCGCG |
| Clpb | CTCTCGAGTGACTAGGACTG |
| Stat1 | GGATAGACGCCCAGCCACTG |
| Stat1 | TGTGATGTTAGATAAACAGA |
| Stat1 | TTAATGACGAGCTCGTGGAG |
| Stat1 | GAAAAGCAAGCGTAATCTCC |
| Stat3 | CCAACAAATTAAGAAACTGG |
| Stat3 | CTGCTTCTCTGTCACTACGG |
| Stat3 | GTTTACCACGAAAGTCAGGT |
| Stat3 | CAAAGAGTCACATGCCACGT |
| Stat5b | CTGATTCGCAGTGATTACAG |
| Stat5b | TACAGCGAACCAGCTCCATG |
| Stat5b | GGCGTTGTCCCAGAGGACAG |
| Stat5b | AGTGGATCGAAAGCCAAGCC |
| Stim1 | TGAGGATAAGCTTATCAGCG |
| Stim1 | GAATACAGGAGCTAGCTCCG |
| Stim1 | GAGCCGTCAAAAATATGCTG |
| Stim1 | CAGCAGATCGAGATCCTCTG |
| Stxbp2 | CGTGTAATAGTGGGGACACG |
| Stxbp2 | CATCCATCCGATACCGCAAG |
| Stxbp2 | TATTATAGGTACCTTCTCCG |
| Stxbp2 | AAGGTTGTAGGTGCTATGTG |
| Tap1 | ACTAATGGACTCGCACACGT |
| Tap1 | GTCTCTAGCAAAGTCCACGC |
| Tap1 | TGCCACATAACTGATAGCGA |
| Tap1 | TGGACATGAGCCATATGTTG |
| Tap2 | AGAAGCCACTCGGACTACTG |
| Tap2 | TTACACGACCCGAATAGCGA |
| Tap2 | GCTGTGGGGACTGCTAAAAG |
| Tap2 | CATGCACACATACCTGATGG |
| Tapbp | TGGTGTTAGAGACACTCTGG |
| Tapbp | CTTTCCCAGCTGGACTCGAG |
| Tapbp | CGCCACTCCAGCCCAAAGGG |
| Tapbp | AGCCGTGAAGCCTTCTCAGG |
| Tbx1 | ACATAGACAACATGGAATCG |
| Tbx1 | CTTCCGGGATTGCGACCCGG |
| Tbx1 | ACACTACCACCCGGACTCGC |
| Tbx1 | GCGCACGGATCGTAGCGCGG |
| Tcf3 | ATTATTGCTGGAGTGATCCG |
| Tcf3 | GCTCCTAGGAACGTGGAAGG |
| Tcf3 | GAAGAGCCGTCACCTAGCAT |
| Tcf3 | CTGCAAACTGGGTTCCCCCG |
| Tcn2 | GAAGCGGCTCCATGACAGCG |
| Tcn2 | TCTGAGACCACGAATCACCA |
| Tcn2 | GAGACTAGCAATACCGCAGG |
| Tcn2 | GAATATCTATAGCACCCCAC |
| Tert | CCTCCAGCCTAACTTGACTG |
| Tert | CACAGAGGGCCAGATATCCG |
| Tert | TCAGCATGCTCAACTATGAG |
| Tert | AAGTCCTGCTCGGTCCCCCG |
| Thbd | TACCTACAACACCCCGTTCG |
| Thbd | GTGTGAGACAGGCTACCAGT |
| Thbd | CTGTGAAGTAAAACTCACAG |
| Thbd | TCGCAGTTAGATCCGAAACA |
| Tnfaip3 | GCAGCTTGTCAGTACATGTG |
| Tnfaip3 | ATATCCATGAGTGATAGCTG |
| Tnfaip3 | AGCCCCGAGGAAACCGCTGG |
| Tnfaip3 | AGGACTTTGCTACGACACTC |
| Tnfrsf11a | ACCAGCACAACGGTCCCCTG |
| Tnfrsf11a | ACACTGAGGAGACCACCCAA |
| Tnfrsf11a | GTTTAAGCCAGTGTTTCACC |
| Tnfrsf11a | AGACGCAAGGAGACCTCTCG |
| Tnfrsf1a | AGTTGCAAGACATGTCGGAA |
| Tnfrsf1a | AGACCTAGCAAGATAACCAG |
| Tnfrsf1a | GATGGGGATACATCCATCAG |
| Tnfrsf1a | GGATCCCGTGCCTGTCAAAG |
| Cd40 | ATTCGCCTGAGTCACATGGG |
| Cd40 | GGGATGACAGACGGTATCAG |
| Cd40 | AGTCAGACTAATGTCATCTG |
| Cd40 | CTGCACCAGCAAGGATTGCG |
| Cd27 | TCTCTCCAGACTACCACACC |
| Cd27 | TGCTGCATACCTGTGCCATG |
| Cd27 | AGACAAACACTACTGGACTG |
| Cd27 | CTCAGGTACATTCTTTGTGA |
| Tnfsf11 | GCCTCGATCGTGGTACCAAG |
| Tnfsf11 | GACCCTCGTGTGGGACGCCG |
| Tnfsf11 | GTTAAGCAACGGAAAACTAA |
| Tnfsf11 | AGATTTGCAGGACTCGACTC |
| Tnfsf12 | GGCTGACCACGACCAGCAGG |
| Tnfsf12 | GCCCAGGCTCAGCACCAGCG |
| Tnfsf12 | CAACGCTGTCTGCCCAGGTG |
| Tnfsf12 | GGAACTGAATCCCCAGACAG |
| Cd40lg | TATTTCAAAACAGGTCGAAG |
| Cd40lg | AAGCTAAAGAGATGCAACAA |
| Cd40lg | TTATACCATGAAAAGCAACT |
| Cd40lg | TGAACTGTGAGGAGATGAGA |
| Cd70 | TTGGGAAGGTCCTTCACACA |
| Cd70 | AGCTGTAACTCAGCTGTGTG |
| Cd70 | AAGGACCCCACACTGCGCTG |
| Cd70 | CATCTGCGTATCCATCAAGA |
| Traf3 | GCTGGGGGCATTGACACACT |
| Traf3 | CAGGTTCACGTGCTGTACCG |
| Traf3 | AGTGACTGCACGTGGCCTCG |
| Traf3 | GCTTTGAGATCGAGATTGAG |
| Trex1 | TTTCCTCGAACCATTCCCTG |
| Trex1 | ACACAGAAGGTACCATCTAG |
| Trex1 | AGCTTGTCCACCACACGGGG |
| Trex1 | GGAGCAGAGGAAAGTCATAG |
| Tfrc | CTACACGCTTACAATAGCCC |
| Tfrc | GAATACATACACTCCTCGTG |
| Tfrc | GGGCTCCTACTACAACATAA |
| Tfrc | AACCCTCGGGAGACTCCACT |
| Tnfrsf4 | GAGCCGCTGTGATCATACCA |
| Tnfrsf4 | TCACACTTGGAGTTACAGCA |
| Tnfrsf4 | TTCCAGATAAGGTACAACTG |
| Tnfrsf4 | GTAGACCAGGCACCCAACCT |
| Ung | ACGGACCTAATCAAGCTCAC |
| Ung | TTGTCAGGGTGGGCCCGACA |
| Ung | CCAACCCCGACTCTGACTCC |
| Ung | CCACAAGGTCTATCCGCCCC |
| Vps45 | AAAGCTGATAACGAATCATG |
| Vps45 | ACGAACTCTTTGAATTCCGG |
| Vps45 | TGTCTCAAAGCACGTGACAG |
| Vps45 | TTGTTTCCTTCGACCCACAA |
| Was | CAACTGATAAGAAACGCTCA |
| Was | CCACCAGCACCAATCAATGA |
| Was | TCACCTGTAGGCCATAAAGG |
| Was | GCAGGTGAACAACCTAGACC |
| Wdr1 | GTAGCCAAGTATGCACCCAG |
| Wdr1 | GGCCCTACCGACTAGCAACA |
| Wdr1 | GTGTGCGATTCTCTCCTGAT |
| Wdr1 | ATGGCTCCCAGAATAGATGT |
| Zap70 | CAACGGCACGTACGCCATCG |
| Zap70 | GAAGCGAGAGAATCTCCTCG |
| Zap70 | TCGACAACCCCTACATCGTG |
| Zap70 | CGCGCACCATAGCATCACGC |
| Ikzf1 | GATGGCCTGGTCCATCACGT |
| Ikzf1 | GTTGGTAAGCCTCACAAATG |
| Ikzf1 | CAGAACTCCAAGAGTGATCG |
| Ikzf1 | AGAAACTAACCACAACGAGA |
| Hax1 | CAGAACTATTCTCACCTCCG |
| Hax1 | AATTCACACACCAGGAGAGT |
| Hax1 | CTGAAACCGAATTCCTCTGG |
| Hax1 | GCTCCAGATTGGGGGTCGCA |
| Sh3bp2 | TCAGCAAGAAGCACCGAACA |
| Sh3bp2 | GAGGTGAACGAGTGGGCACG |
| Sh3bp2 | ACCTGGAGCCTGATTCCCCG |
| Sh3bp2 | TGGGTACATTGCTCATTGGG |
| Srp54a | CTTATAGAGAAGTTGAAGCA |
| Srp54a | AAAGCGTGGACACCGACTAA |
| Srp54a | TATGTGATGGATGCATCCAT |
| Srp54a | AAACTCGACGGTCATGCGAA |
| Rbck1 | TGCTTCATACCAGCCTGACG |
| Rbck1 | AGTACGCCCGGATATGACAG |
| Rbck1 | TGCATTCACACGGCATTCGG |
| Rbck1 | ACCCGAGGTCTCCCCAACAC |
| Usp18 | CATCATGAACACTTGAAGCA |
| Usp18 | ACAGCTCTCGCAGCACATGT |
| Usp18 | TGTACAGCCCACGCAAATCA |
| Usp18 | CAGGCACTGAACGAGCTCCG |
| Clcn7 | GGAAAGACGAATCAACCACA |
| Clcn7 | ATAGCTGCAGGGATTTCACA |
| Clcn7 | AATCGGACAGATGAACAACG |
| Clcn7 | ACTCTGCCTTCGTACTCGTG |
| Tcirg1 | ACACAAGTGCCTCATCGCGG |
| Tcirg1 | TCTCCCGAAAGCTGGCAATG |
| Tcirg1 | CAGCCACACTCCAACCTGAG |
| Tcirg1 | CGCTACAGGGAAGTTAACCC |
| Nbn | TATTCCTACATCAACAACGC |
| Nbn | TGGAGAAAGCTGATAGCTCG |
| Nbn | ATAGCTGGTTCATCAATGGG |
| Nbn | GAGAATTACTGTAATCCGCA |
| Tinf2 | CAAAGGCGTGCCATAAAGAG |
| Tinf2 | CCGGTAGCAAACCAAGCCGG |
| Tinf2 | CCAAAGGGGCCAGATTAAAG |
| Tinf2 | AGTGCTCTACGCGGCGACGG |
| Elane | CCGGCCACCAACAATCTCTG |
| Elane | CTCACAGGCCGTTCACACAG |
| Elane | GAACGACATTGTGATTATCC |
| Elane | ATGACCTCCACGCCTCTGCA |
| Cfhr1 | TTGCCTTCTTAGATCCAACA |
| Cfhr1 | TGCATATACTGGTAACGACA |
| Cfhr1 | ACTGGTGGAAATGCATTTGG |
| Cfhr1 | AGAAGGTGACATTGTACAAG |
| Ctps | ATTGGCCATTAACCACAAGC |
| Ctps | ATACCAGTACGTCATTAACA |
| Ctps | GCCCACAAGAGCGATCGAGC |
| Ctps | TTAATACCCGTAGACGAAGA |
| Slc46a1 | AGAGCTAACATCTGCCACAG |
| Slc46a1 | GGGCAATGGATCGATGATGG |
| Slc46a1 | TGGACCAGAAGAGTCCCACC |
| Slc46a1 | GAACTGTGGGAACCAAAGCG |
| Nhp2 | GCAGAAGCAGATTCGTCGCG |
| Nhp2 | CCATCTTCCAGTTCTGTGCG |
| Nhp2 | AGGAGATACATTGCCGATTG |
| Nhp2 | GGCCGCTCCCGAAGAGTCCG |
| Map3k14 | GCAAAGCCCGAAAAAAACGT |
| Map3k14 | CGTGGTTTAGACATTGCAAG |
| Map3k14 | CCCCGAAACTGAGGACAACG |
| Map3k14 | TGGGCCAGTTGGCTTAGATG |
| Rfx5 | ACATAATGACCGTTCTCGAG |
| Rfx5 | ATGGGTGTGATAAGTGATCG |
| Rfx5 | AGAGCGTCTATGATGCCTAT |
| Rfx5 | TCTACCTTCAGCTCCCATCG |
| Irf7 | CTTGCGCCAAGACAATTCAG |
| Irf7 | GGCAGGTTAACTCCACTAGG |
| Irf7 | TGTGCGGCCCTTGTACATGA |
| Irf7 | GGAGCAAGACCGTGTTTACG |
| Irf3 | CCAGTGGTGCCTACACCCCG |
| Irf3 | AGAAACAATAGCCAGATCTG |
| Irf3 | GGCTGGACGAGAGCCGAACG |
| Irf3 | CTGGCGGCCTCGGTAGAAGG |
| Icos | TGGTCTTGGTGAGTTCGCAG |
| Icos | TATGCAAATATCCTCCACTA |
| Icos | GCAGAAGTAATAGCTTCCCT |
| Icos | AAATGAAAACATCCTATGAT |
| Smarcal1 | CTCCCAGGTGAAGCGCACAG |
| Smarcal1 | TTTGGGTTATAAATCCAGCG |
| Smarcal1 | TCATTGCAGATATCAAGACC |
| Smarcal1 | GCTTGGCATCCACTCCGGAA |
| Atp6ap1 | GAGGATTTCACAGCATACGG |
| Atp6ap1 | GATATGACCCTCATGTGTGT |
| Atp6ap1 | TAGCTAGATCCACATGCAAG |
| Atp6ap1 | GTGTCATTGTAACTCACAGG |
| Unc93b1 | CTGGGAACGCTACTACACGC |
| Unc93b1 | CCGCAGTTGGACGAACTCGT |
| Unc93b1 | AGGTGAAGTATGGCAACATG |
| Unc93b1 | GCACACGCGAAGCTCAACTG |
| Nfat5 | TCAGCCATTTACGTACACTC |
| Nfat5 | AGTATCCGGTTAAAAGTGAG |
| Nfat5 | GCCGTGGGGGTAAGTAACAG |
| Nfat5 | AAGACCAACTTCTATAACAG |
| Il1f5 | GTCATCCTGGGCGTTCAAGG |
| Il1f5 | AACATCATGGAGCTCTACCT |
| Il1f5 | AGGGCCAATTCTGAAACTTG |
| Il1f5 | TGTTGTCCCAAATCGGGCAC |
| Mefv | TCTGATAACCTACTACGGGG |
| Mefv | AAGGAGGGTACCTTCACAAG |
| Mefv | ACCAAAAGGAAGGATCAGAG |
| Mefv | TCTGAATGGAAGGACTACGG |
| Extl3 | GCCCAAGCCTCGCGTCACAG |
| Extl3 | TCAGACATAGCATGGACAAG |
| Extl3 | CCACACAGTGCCCACTCAGT |
| Extl3 | ATTGCGGAGGTATTTAGGTG |
| Tyk2 | GTGTGGTTACGGCGACAGAG |
| Tyk2 | TAGACCCCCGCATGATGACG |
| Tyk2 | CTTCCAGGACATTTCCCACG |
| Tyk2 | ATCCACATCGCACACAAAGT |
| Samhd1 | ATCCTTACATTATGTCGATG |
| Samhd1 | GCTTGATATAGCGAAGTCGC |
| Samhd1 | CTTGGGCTGCCATCGCAGCG |
| Samhd1 | TTAGGATCTTACCTAGGTCG |
| Adar | ACTCCAACAAGCCGCCTACG |
| Adar | AGAGGTAACCCCAGTAACAG |
| Adar | TTCTTGTAGGGTGAACACCG |
| Adar | TGTATCCAGGAATTCCCTAG |
| Tbk1 | TGCCGTTTAGACCCTTCGAG |
| Tbk1 | CTTCTCGCTACAACACATGA |
| Tbk1 | CAACATCATGCGCGTCATAG |
| Tbk1 | CGGGAACAACTCAATACCGT |
| Mogs | TCTAGGTCATTCTTCCCACG |
| Mogs | TCGGCAGCATATCCACGATG |
| Mogs | TGCCGAAATAGACGTGTGGG |
| Mogs | GAGGTCCTACTACCAGAGAT |
| Tnfrsf13b | GTACTACGACCATCTCCTGG |
| Tnfrsf13b | GAAGTCAGGTCAGACAACTC |
| Tnfrsf13b | GTGGCTCTCCTCTACGCCTG |
| Tnfrsf13b | ATCAGTACTGGGACTCCTCA |
| Bcl11b | GTTGTGCAAATGTAGCTGGA |
| Bcl11b | CTGAGAGCCCGTCGTCTGAG |
| Bcl11b | GCGGGAAGTTCATCTGACAC |
| Bcl11b | CAGAGGTGAAGTAATCACGG |
| Stk4 | AATACACCGAGATATCAAGG |
| Stk4 | TATCTGATATCATTCGGCTA |
| Stk4 | TGCTGATATCCATCCAATGA |
| Stk4 | TTCAGGAGCCATCCAAAACG |
| Ncstn | CCAGCAGAACCATGTAAGGG |
| Ncstn | GTGCTATCAAGATCACAACC |
| Ncstn | AGGGACAGAACTCACCCGTG |
| Ncstn | GGTGACTCACCGATTGTGGA |
| Il21r | GCACGTGTACCATATATGTG |
| Il21r | AGGACGCTATGATATCTCCT |
| Il21r | GCCACCTCACCACAGCATAG |
| Il21r | GGGCTGGAAGAAACTCTCAG |
| Il21 | GGTGACGAAGTCTAATCAGG |
| Il21 | GAAAATCTATGAAAATGACT |
| Il21 | AAACTCAAGCCATCAAACCC |
| Il21 | GACATTCATCATTGACCTCG |
| Lpin2 | GGATACTCACGTTTCTAATG |
| Lpin2 | GGTTATATATCCGGATCACG |
| Lpin2 | TGCAGGTCACCAAAAGAGAA |
| Lpin2 | AACCTGGATCCGTGTCACTG |
| Nop10 | CGATCGCGTTTATACGCTGA |
| Nop10 | CAATATTACCTCAACGAGCA |
| Nop10 | GGAGAACCGAGCAGGATGGG |
| Nop10 | GAGCAGAAATTTGACCCTAT |
| Psenen | CGAGAATGATCACCCAGAAG |
| Psenen | CCGGAAGTACTATCTTGGTA |
| Psenen | CTTGGAGCGGGTATCCAATG |
| Psenen | ATGTTGACTAACCAAAGAAA |
| Ndnl2 | CACGTATTGCAGGCGCTCGG |
| Ndnl2 | CAACACCATCACCGAGACTG |
| Ndnl2 | AGAAGAAGATCCCGATCAAG |
| Ndnl2 | CCCTGTCGGCCCCGACCGTG |
| Sbds | CCAGATCCGACTGACCAATG |
| Sbds | CCTTCATGGCTCTCTCGATG |
| Sbds | ACTGAAATCTGCAAGCAGGT |
| Sbds | TAGGGATATCGCCACCATTG |
| Taz | TTTGGAGAAGCTTAACCATG |
| Taz | GGTCATCCATGCAAGACTGG |
| Taz | AGCAGCTCACCTCGACACAC |
| Taz | TATGAGCTCATTGAGAACCG |
| Cdca7 | TCTACTGGTCGGATTATATG |
| Cdca7 | GCTGTCACAACTGTCATCGG |
| Cdca7 | TGTGGCTACCAGAAGGAACC |
| Cdca7 | GTCGTCCAGAGAACAAGCCA |
| Magt1 | TGTGTGTGCGATCGCAGCGG |
| Magt1 | CAGCTGAGCAGATTGCCCGG |
| Magt1 | TTATGCTGGACCCCTAATGT |
| Magt1 | GTGGAGCTTTAACAAGACGA |
| Rnaseh2b | CTCCTAGGTCAACCAAACTG |
| Rnaseh2b | TCAGCCCTTGGACCAAGTCG |
| Rnaseh2b | CAGCCTTTAGGAGATAGTGA |
| Rnaseh2b | TGGCCAGCTTTACGAACAGG |
| Jagn1 | GCACTACCAGATGAGGTACG |
| Jagn1 | GTACCTCATCTGGTAGTGCA |
| Jagn1 | AGTGCATGGCGACGCGCTCC |
| Jagn1 | CCGTCGGTGCCGGCCGCTCG |
| Ino80 | GGGTTGCGGAATATCCTCAC |
| Ino80 | AGAAAATGAATTGTCTGACA |
| Ino80 | TGCCCATCAATGCATGAAGG |
| Ino80 | GTGACGTGGAGATTCTTCAG |
| Rnaseh2c | CGTGAAGAAGCGATCTACCG |
| Rnaseh2c | CGGCTGACTAGAACGTCGCA |
| Rnaseh2c | AGGGTTTGCGGGATTCGTGA |
| Rnaseh2c | ACCGTCTGCATCGTGGCGGA |
| G6pc3 | TAGGCCGACTGCCAATAGGA |
| G6pc3 | TTCCCGGGCTAGAGAATATG |
| G6pc3 | GGGGCTCATTAGCCAGCCAA |
| G6pc3 | TAAAGAGAGTCCAATACATG |
| Ctc1 | CTGGTCGAATAACCCGCCTG |
| Ctc1 | CTTGTAGATGAGATCCTCCA |
| Ctc1 | AATTGCAAGTAACCAAGACC |
| Ctc1 | CTTGGAACTATGGGGTACAC |
| Gins1 | CCGGCGCGCGGTGTAACTCG |
| Gins1 | TGCTTCGGATTAGAGCACTC |
| Gins1 | GTTGTTGTTCACAGACGGAG |
| Gins1 | AGTCTCTTGCTACTTACATG |
| C8g | TGGGACTCACAGCTTTCGGA |
| C8g | CTGGGGAGCTGCGTGCAATG |
| C8g | CCTCACCTTGGAACAAGAAG |
| C8g | CTGAGCACTGAAATTGACCT |
| Rnaseh2a | TGTCACACGATACAGCTGCG |
| Rnaseh2a | GTAACAGATGGCGTAGACCA |
| Rnaseh2a | AGACCTTGACAGAGAACGAG |
| Rnaseh2a | GGCTCCTTGAGACACACAGC |
| Trnt1 | CATGAATTAAGAATAGCAGG |
| Trnt1 | ATTCGCATGATCAACAACAA |
| Trnt1 | AACCTGTTCTCACCTATGTG |
| Trnt1 | GGCAGAATTGTCGACAGACC |
| Rnf168 | GGTGTGCCCCAGTTAACTGG |
| Rnf168 | CCCGGGCCACAGTTATGTGT |
| Rnf168 | AGCATGGGTTACAGAGCGTG |
| Rnf168 | AATCTCTGGACAAGAATCAA |
| Unc13d | AATCTGGTACAGGACGTCAT |
| Unc13d | ACAGAGACCTACCCAGACCG |
| Unc13d | TGGCTGGCTGAAACCAGCGG |
| Unc13d | GCAGCCCTGTGTCCCGACGT |
| Nbas | AACAAAACAGCATATTCGTG |
| Nbas | ATAGCAGTATTTATTCCATG |
| Nbas | ATTGATTGATGTCAATTGGT |
| Nbas | TTTGCCGAATGACATGACAT |
| Slc29a3 | AATGATGGCCATGCACGCGA |
| Slc29a3 | GGGAAACTGCGCAGAACCCG |
| Slc29a3 | CAAGGAAGACTGCTGCCATG |
| Slc29a3 | ACCAGAAAACACTCGAACTG |
| Ifih1 | TGTGGGTTTGACATAGCGCG |
| Ifih1 | CGTAGACGACATATTACCAG |
| Ifih1 | TGGGTTCCAAAATCTGACAT |
| Ifih1 | GATTGATGCATATAGCCACC |
| Snx10 | GCAGAGCCATCTGAACTCCG |
| Snx10 | CATGTCGACCAACGCCGCCA |
| Snx10 | AAACATCTTGTGTACGAAGA |
| Snx10 | AAAGCAAGAGTGCATTCTGC |
| Tnfrsf13c | CAGACTCACTAGACCCACCA |
| Tnfrsf13c | GACACGCAGTTTCTCACCAG |
| Tnfrsf13c | CTCCGCGCTGAGACCCGACG |
| Tnfrsf13c | GCACTCGGTCTGATTGCACT |
| Spink5 | CGGGAAAGTGACCCAATCCG |
| Spink5 | AGGGAGAGTGACCCTGTACG |
| Spink5 | CGAGAAAATGACCCTGTGCG |
| Spink5 | TGGAAGACTTGGATGTACAC |
| Tmem173 | GAAGGCCAAACATCCAACTG |
| Tmem173 | TATCTCGGAATCGAATGTTG |
| Tmem173 | AGTATGACCAGGCCAGCCCG |
| Tmem173 | CAGTAGTCCAAGTTCGTGCG |
| Parn | AAAACGGTCTCAAGCCAATG |
| Parn | GGTCATACTTAAAAGCACAA |
| Parn | AAGACATATAGTTATCAGCA |
| Parn | CCAACTCACCGAATTAGCAA |
| Stx11 | TCGAACACTATGTCCTCCCG |
| Stx11 | AGCCATGTACGAGTACAACC |
| Stx11 | CATGTCGGGCGAGCAGATTG |
| Stx11 | AGCTGCTTCTGATAGACGTG |
| Rhoh | GGTACTGATGTGCTACTCTG |
| Rhoh | CCCACGGTGTACGAGAATAC |
| Rhoh | ACAACCAGCACCGGGGTACA |
| Rhoh | CAGGGGCCGGATACTTCTGA |
| Dock8 | ACTTCCGGTTTGCGACACCG |
| Dock8 | CCGGAACACGTAGTGCACGT |
| Dock8 | AGCACGTTTAAGGGATGACG |
| Dock8 | TAGCGAAGTTCAGTCGCTGG |
| Ercc6l2 | CTGTTACAGACCAACTCACG |
| Ercc6l2 | CGTACCAAGACTCTTATCAA |
| Ercc6l2 | GGTACTTGCGAGATTACCAA |
| Ercc6l2 | AAGGATGAATTGGATACCTG |
| Dnajc21 | TAGGTACGATAACCACCGAG |
| Dnajc21 | CTCCCAGAGCGACTATGACA |
| Dnajc21 | TAAACGAGATAAGAGAGTGC |
| Dnajc21 | TGGATCAGATGAAAACGAAG |
| Lrba | GACCGTCCCAATGAACTCAG |
| Lrba | GTGTGGCGAGTGGACGAAGA |
| Lrba | ATAGTTACCATGTACCACTG |
| Lrba | CAGATGCAGTAGACCAACAA |
| Lamtor2 | GTTCCCGTTCCTATCATACG |
| Lamtor2 | GCTGGCCTACTCCGGTTATG |
| Lamtor2 | CCACCCTCGTAATGGCTACA |
| Lamtor2 | CACAGATGCCCGGGTCACTG |
| Smarcd2 | GAGATAATGCGGGAACTGCG |
| Smarcd2 | GCATCGGATGCCCACCACAC |
| Smarcd2 | CCCAACACACTCACTCGCTG |
| Smarcd2 | CGGAGCAGCTGTGCCAAATG |
| C230052I12Rik | CTAGCCACAATATGTCCTAG |
| C230052I12Rik | TGAGGCAGGATGCTTCCGAC |
| C230052I12Rik | CAGTAAACTTCTGGACTGCT |
| C230052I12Rik | TATGATCCCTTGAAGATGTG |
| Usb1 | ACACGGTATGTAGATGTGGG |
| Usb1 | TCCTCGATCCTCACCAACGG |
| Usb1 | GGTACAAACATCTGAGCTCG |
| Usb1 | GACGACAGTGCAAAGCATGG |
| Traf3ip2 | GTCCTGCAGGTAACACGAGG |
| Traf3ip2 | GTAGTACTGACAGTTCCATG |
| Traf3ip2 | CCTGCGAGCTAAAGTCCTGG |
| Traf3ip2 | CCAAAGCATTAGGTAAACTT |
| Ticam1 | AAGATGCCATCGATCACTCG |
| Ticam1 | TCTGGAACGCTAATTTCGTG |
| Ticam1 | GATATCAAGGGGGACCCCAG |
| Ticam1 | CTTCACACCATGGAACCCAT |
| Fermt3 | GGAGGTGCATGACCTGACAA |
| Fermt3 | CTGTCACACTCCGAGTCACG |
| Fermt3 | GGGCTACCGCCAGTACTGGG |
| Fermt3 | CTCACCCACATCCCCGCTCA |
| Mthfd1 | ACACCAACGATAGATTCCTG |
| Mthfd1 | CACTATGAATCCGTGCACAG |
| Mthfd1 | GATTGCCGGAAGGCACGCGG |
| Mthfd1 | GGTAGCGTCCAGTAAGAAAG |
| Obfc1 | GACGCAGTTTATAACCCCAG |
| Obfc1 | CAACGGGCATCCAATAAGGC |
| Obfc1 | GGAGATATCATCCGAGTCCG |
| Obfc1 | TCGCAGTGCTCAGAATAGCT |
| Card11 | GCAGCATAACTGTGTTCATG |
| Card11 | ACTATGGAGTCATTTCCCGG |
| Card11 | CACACACTTCCTGATGAACG |
| Card11 | GCGACCTCCAACTCGAGGTG |
| Sp110 | AAATGACCTGGAAATGGCCA |
| Sp110 | GGTCAAATTCCAAAAGAAGA |
| Sp110 | TCCTGAATGCAGCCAAGGAG |
| Sp110 | AGCTCAAGGCAGTGAGCAGG |
| Orai1 | GATCGGCCAGAGTTACTCCG |
| Orai1 | TGGCGATGCATGCGCTCGTG |
| Orai1 | AAGACGATGAGCAACCCTGG |
| Orai1 | AATCCGGAGCTTCCCGTGAG |
| Pgm3 | TACGGCCTCACATAACCCTG |
| Pgm3 | AACTTCTTCAAGGTACCGCG |
| Pgm3 | TACACCATGTAGTGCAACTG |
| Pgm3 | CTGCTATGACATACCCTGTG |
| C7 | CATACTGATCGATTAACCGT |
| C7 | ATCAACACCAAAAGTTTCGG |
| C7 | CAATCCTGCAAACCTGAACG |
| C7 | GTGTCTCCACAACAATAGCG |
| C8b | AAAGACGCCATGGAGCAAGG |
| C8b | ATTGTGTGACTTGTCCGACA |
| C8b | TTGACTGTGAGCTATCCACC |
| C8b | CAAGGACAGAGCGCGCGTGG |
| Cebpe | CAGTACCAAGTGGCACACTG |
| Cebpe | GCAGGTAGTGAGGAAACGAG |
| Cebpe | CAGACTCGATGTAGGCGGAG |
| Cebpe | CTTACCTTGAGGACACGCAA |
| Tirap | CTGTCTGTGAACCATCATAG |
| Tirap | CGAAACAATGGCGCCACCCG |
| Tirap | TGCAGCTCGAGGGTGAGCTA |
| Tirap | CAAGTAGGAGACCAGCTCCT |
| Dclre1b | TGCTAGACAAACCCACCGTG |
| Dclre1b | TCCACTGAAGCATGGCAGAG |
| Dclre1b | AGAGAAACATGACAGAACCA |
| Dclre1b | TGAATGGTCAGAGTACGGGA |
| Tlr3 | CTTAGATGAAATCCCAGTCG |
| Tlr3 | GTTGTAGGAAAGATCGAGCT |
| Tlr3 | GAATGGTCAAGTTACGAAGA |
| Tlr3 | ATCTACAAAGTTGGGAACGG |
| Card14 | CCACAGCCGCATGAAACGTG |
| Card14 | TGTTCTACCTTAGAGACTCG |
| Card14 | CACTCTGGAGAATACAACGC |
| Card14 | AGAACTAAATCGGCTTAAGG |
| Rfxap | TCCAAGACCTGCACGTACGA |
| Rfxap | GCACCGCAACAAGATGTACA |
| Rfxap | CTGGACACATCGGACCCGGC |
| Rfxap | GTCACCGGAGGGGCCGTCCG |
| Il17rc | CTGTGGAACGATGACAACAT |
| Il17rc | AGCAATACTTACCCCAGAGG |
| Il17rc | GGAGCCACAAGATTTCCAGT |
| Il17rc | ATCTAGCTGCCATACCCCTG |
| Wipf1 | GGAACAGAATGCCTCCCCCG |
| Wipf1 | CCGTCGAGTCTGCACAACCG |
| Wipf1 | GGGGCTTGTCATCCAAGGCT |
| Wipf1 | GCTGAGGTCCACCGCCAACA |
| Wrap53 | GGTCATTACCACTTACTAGG |
| Wrap53 | AGAAACGAATCTCCCCGAGT |
| Wrap53 | GAAGAGTTGGGAACCATCCG |
| Wrap53 | GACACAACCAAGCTAGCCAC |
| Tmc6 | GGAGTCATGTCGCTCCAGAG |
| Tmc6 | TTTCCAAGGTCGCAGCCGTG |
| Tmc6 | GACCAGAAAACGTAGGCACG |
| Tmc6 | TACCGGGTTGGCAGTACCAA |
| Tmc8 | GGAGCTCTCTACGAGATTGG |
| Tmc8 | CAACGCTTGCGACTACCAGG |
| Tmc8 | CTTCTACGGTGCCTACCGAG |
| Tmc8 | CATCCGCACAGGCGTCCGGG |
| Ttc37 | ACAAACAGATCACTTCTAGT |
| Ttc37 | CGCCTGTGTTCAGACAGGCG |
| Ttc37 | TGACGTGTGTAAGAAACTCG |
| Ttc37 | GAGGCGTACTTAAGCAGAGG |
| Mkl1 | AGACAGTTCCTCCTTCGACG |
| Mkl1 | GTGATGAGAATTCCACACCT |
| Mkl1 | CTGCCCCCAAGCCTAGCCAA |
| Mkl1 | GTCAGGATGCACATTCTGGA |
| Ttc7 | TGGGATCGATGACATATCCG |
| Ttc7 | TCCAAGACCAATTACTACCG |
| Ttc7 | GCAACACACCTGTCGGACGA |
| Ttc7 | CTGCCCCAAAGACAACATAG |
| Dclre1c | TTTATTCACCAACCTAAGCG |
| Dclre1c | CAGGGAAACATACACGACTT |
| Dclre1c | GCATCAAGCCATCTACCATG |
| Dclre1c | TCAACTAAAGATATCTGCGT |
| Slc35c1 | AGGGCACCCCTACGTACTTG |
| Slc35c1 | AGGTTAGGCGCCAGATACTG |
| Slc35c1 | CTCACCAATGATGACGCCGC |
| Slc35c1 | GCAGTGAGGTCACCAGGCAT |
| C8a | TCCTGCTATAGGTACTACTG |
| C8a | GGAGATTGAAGTATCCGCCA |
| C8a | CATTGTACACAGTTTCACAC |
| C8a | CTTGCCTTAGACAAGGTGTG |
| Rltpr | GCCTGACTTACCATGAAGGG |
| Rltpr | CAACCAAGTAGACTCTACTT |
| Rltpr | AGACCACCCTGGATACCACA |
| Rltpr | ACATGCGCCTGTCAATCACT |
| Malt1 | GTCCTATGCCTCACTACCAG |
| Malt1 | ATATGAGATGTGTAACGCTG |
| Malt1 | CCACTGGCTAAATTCAAAAG |
| Malt1 | AGCTTGGACCGCGCTCCGGA |
| Dkc1 | TTTACTGCAGCAATAAGTGG |
| Dkc1 | TCTCTACCCGAAGTATTCGT |
| Dkc1 | TACGCATAGTGTCCGAATGT |
| Dkc1 | ATGATGTACTCGATGCTCAG |
| Ap1s3 | CCACACTCCCTGACAAGGAG |
| Ap1s3 | TCCAGACCGTCCTCTCTCGT |
| Ap1s3 | TCAGTCGACAAGGGAAGCTG |
| Ap1s3 | TAATATCCAGCTCACAGACC |
| Il17f | GACTTACTTGTAATCCCATG |
| Il17f | TGGGAACTGTCCTCCCCTGG |
| Il17f | AGCGGTTCTGGAATTCACGT |
| Il17f | GGGCCTCAGCGATCTCTGAG |
| Nod2 | CCGACCCATCGTAAGTACTG |
| Nod2 | AGATGCCGACACCATACTGG |
| Nod2 | GCAGAGTCTGGACTGACGTG |
| Nod2 | GTTGTAGAGTCTCCTCACAA |
| Irak4 | GTAGCTATCAAAAAGCCGTC |
| Irak4 | CTGTGTGAACAACACCATCG |
| Irak4 | TGTAAGCATACACTAAGCAC |
| Irak4 | CAAGGTGCAAGGTTGCTCAG |
| Zbtb24 | GGAAATCGAAGTTACCCGTG |
| Zbtb24 | TGAGAAATACTCGCTACTGG |
| Zbtb24 | ACACGTCCATAAACTTTCGG |
| Zbtb24 | GATCCTCTGAAACGGAAACG |
| Rnf31 | GATGGATTGAGTTTCCCCGA |
| Rnf31 | GAACTATGAGTTGTTGGACG |
| Rnf31 | CTACCTCAACACCCTATCCA |
| Rnf31 | GGAGGAACCAAGGTGTTGTG |
| Rtel1 | ACACGGGGATCCATGCGTGG |
| Rtel1 | TATGCCGACATACCGGTAGG |
| Rtel1 | GTTGTCATACACCTTAAGGT |
| Rtel1 | GAGGCGTCACCAAACCTGGA |
| Irf2bp2 | CGCACGCGTGCTGAAGTCGG |
| Irf2bp2 | GCGGCAGAACCGTTGACCAG |
| Irf2bp2 | TCTCGATGACGAACTCCACG |
| Irf2bp2 | GGCCGACAGCTTATCCAGCG |
| Gimap5 | GTCACCAGTAACAACACATG |
| Gimap5 | AGCTAGATTCTTGTACACAG |
| Gimap5 | ACTGGAATGCTGGTCGTCGG |
| Gimap5 | CTGAAGATGCCATGGCTGTG |
| Lig4 | TTAGACGTCCTAATTGTGGG |
| Lig4 | GCAACTCAACTGCATCATTG |
| Lig4 | GAACGCATTGTGAATAAATG |
| Lig4 | CATCCGTGCGAGCTCCACTG |
| Chd7 | GACATGCCCATAAACGAACG |
| Chd7 | AAACGGTTTAAGTCAAAACA |
| Chd7 | CCTACCGGAATGATTTAGCA |
| Chd7 | TCGATGACCACACAGCGCCA |
| Ccbe1 | GACCTACCGAGAGGAACCCG |
| Ccbe1 | CAGCAGTGCACGGATAACTT |
| Ccbe1 | ACAGAGAGTGGTATTGCTGG |
| Ccbe1 | GAACTGGGCAAGTATGTCAA |
| Fat4 | GACATCGTGGACGATCGAGG |
| Fat4 | CAGTTATCTCATCACTACCG |
| Fat4 | GGGATGTCGAAAGAGTACAC |
| Fat4 | ATTAGATCCTATGTCCGCGT |
| Card9 | GATGTACAAGGACCGTATCG |
| Card9 | GTGACTTTCCGGTATAACTG |
| Card9 | TAGATAGGGTGTGATCCGGG |
| Card9 | CCAACCTGGTCATCCGCAAG |
| Plekhm1 | TGACTTGTAGGAGAGTTCGA |
| Plekhm1 | TGATGAGGAACGCACCTGTG |
| Plekhm1 | GCTGGTAGCTAGGCTATGGA |
| Plekhm1 | ACTCACCGGACTCGTAGGAG |
| Kmt2d | AAATGGCTGTTGATCCCATG |
| Kmt2d | GTTCACCATTAATACCCCCA |
| Kmt2d | TCGGGCCGGACTAACATCCG |
| Kmt2d | TGGGGATGGACAGCCCGACG |
| Otulin | GGAACTTCACAGCTTCGTAG |
| Otulin | TGATAACTACTGTGCACTGA |
| Otulin | AACAGAACCCAGGTTAAGTG |
| Otulin | AGTATACCTGGATCAAGCAG |
| Cfhr2 | TTGTCCCCTTAGACTCAACA |
| Cfhr2 | GAATGGAGACTCTACATACT |
| Cfhr2 | ACTTATTTCCTATAGACACT |
| Cfhr2 | ATACCATTTAGAATATAAGG |
| Nlrp1b | CTAAATGACCTGTGTGACGA |
| Nlrp1b | AGATGCTAAAGAGCACCCTA |
| Nlrp1b | AGAAGATCATTCCTTATGTG |
| Nlrp1b | CTGTAAGCAAGGGTTAGCAG |
| Vps13b | TGTCCATACTACCCAAATCG |
| Vps13b | TAAACACTGCAATACAAGCG |
| Vps13b | GAAACCTCTTCCCGATACAG |
| Vps13b | TGGCAGTAGTCCATGTACTG |
| Isg15 | GTCCGTGACTAACTCCATGA |
| Isg15 | CAGCAGCACAGTGATGCTAG |
| Isg15 | GCATCCTGGTGAGGAACGAA |
| Isg15 | TGGAAAGGGTAAGACCGTCC |
| Epg5 | ACAGCCGACTCGTTGTAACA |
| Epg5 | TCGAGCCAGAAGAACCAATG |
| Epg5 | TGGGTACCATACCCATATTG |
| Epg5 | GAAACGCTGTCTTACACAAG |
| C1ra | TTTGCCAGAATGATGGCACA |
| C1ra | TGTGCAGGTATATATCCCTG |
| C1ra | CCACACAGACTTCTCCAATG |
| C1ra | TGGTTGTCTCCAAGTCACTG |
| C1s2 | AATTCCTCATGTCATCATGG |
| C1s2 | TTCTCTTAGATAATCTCAGG |
| C1s2 | TCTTTCAAACTGATCTAATG |
| C1s2 | AACACACAGTTAAACTGTCA |
| C4b | GCACAAGATGCCTCTTAGTG |
| C4b | CAGGCACAGCCCCTCAACTG |
| C4b | GTAAGCCACAAAGTAGAACG |
| C4b | TGAAACAAAGGACCATGCTG |
| Cd59b | TATTATGAGCCGATTAGACG |
| Cd59b | ATGCTACAACTGTTTAGACC |
| Cd59b | GGCAAGTGTATCAACAGTGT |
| Cd59b | TTGATACACTTGCCTTCCTG |
| Lat | ACTCACGAGGTGGCTTGATG |
| Lat | GCATCCGATGGGAACCCCCA |
| Lat | ATACTCACGGGATGGGGAGC |
| Lat | CTCTGTGGAAGTGCTGTCAT |
| Nhej1 | CCCTATGCATAGAGCTCGGC |
| Nhej1 | ACCAGCCGAGCTCTATGCAT |
| Nhej1 | GGCTGGTTATCAGTTCCTCG |
| Nhej1 | TCACCAACAGCACACGCCTC |
| Dock2 | AAGAAGTGACAACCACGCTC |
| Dock2 | CAGCATCTCACGCTACAGAT |
| Dock2 | GTGACAGTTTATGCTTTATG |
| Dock2 | TACATCCTTTATCCATCTCA |
| Mysm1 | ATCACTATCTTGATTCAACG |
| Mysm1 | AAAATTCTGGGTTAATCAAA |
| Mysm1 | TATCATTGAGAAAATGCTGT |
| Mysm1 | TTATCTAATAAATCACTTCC |
| Kdm6a | TTGGATAATCTTCCAATAAG |
| Kdm6a | TAGCATTATCTGCATACCAG |
| Kdm6a | GAAACCTCACGAACCCGAAA |
| Kdm6a | CGCCAGGATGAAGGCCCTGC |
| Tpp2 | CACACCAAGCAGTCATATAC |
| Tpp2 | ATTGATATCATTGATACAAC |
| Tpp2 | ATAGGCCAATAAACTAATCA |
| Tpp2 | TGGTTATGACTTCTATCCAA |
| Il12b | CCTGCCCATTGAACTGGCGT |
| Il12b | CCATGAGCACGTGAACCGTC |
| Il12b | CATGTCACTGCCCGAGAGTC |
| Il12b | GACTGGACTCCCGATGCCCC |
| Stat2 | TGAGATTGAAAATCGAATCC |
| Stat2 | TCCACAACTGCTTCGGGGGC |
| Stat2 | CTGAGCTGTAGTGGTCCCAC |
| Stat2 | AGTTCTTGGTGAGATCCATC |
| C3 | GATGACGACTGTCTTGCCCA |
| C3 | ACAAAGGCAAGATGCCGTGT |
| C3 | GAGCGAAGAGACCATCGTAC |
| C3 | GACAGTCGTCATCCTCATTG |
| C6 | TAGTAGTGAACGATTACTAT |
| C6 | CGATAAGCTTTGTATCAAGC |
| C6 | TGAAACACGGTATGGATTAC |
| C6 | GGTTGCCCCCCAAACTGACT |
| Cfb | TTCGAGTCTGCACGGGGTAT |
| Cfb | AATACGCTGCCCACGACCGC |
| Cfb | GCTGGATACTGTCCCAATCC |
| Cfb | AGCCACGCAGGACAAGTCCC |
| BRDN0000737505 | AAAAAGTCCGCGATTACGTC |
| BRDN0000737693 | AAAACGGCTCGATCGGTGAT |
| BRDN0000737637 | AAAACGTAATTATACCGAGC |
| BRDN0000738185 | AAAATTGCACCTTCCCGGCC |
| BRDN0000737801 | AAACCCCCGCGCGGAGCGTC |
| BRDN0000737467 | AAACCTAGCGTAGATTCGGC |
| BRDN0000737848 | AAACGAGGCTGTTCGTACAC |
| BRDN0000737609 | AAACTCATACGTAGCGAATC |
| BRDN0000737434 | AAACTCCCGTGTCAACCGAT |
| BRDN0000738254 | AAAGACGTGCATTCAGCGAG |
| BRDN0000737777 | AACATGTTAAGTCGCGTTAT |
| BRDN0000737611 | AACCAGCATTTGACCGCGCT |
| BRDN0000737528 | AACCCCGGCTGTCATCGCCG |
| BRDN0000738228 | AACCCGCCGGAACAATCAGC |
| BRDN0000737727 | AACCGGCTGCGCGTTTGCAA |
| BRDN0000737483 | AACCGTACTGCGAGGAGCAT |
| BRDN0000737872 | AACCTCGTCTCATGTACGAA |
| BRDN0000737516 | AACGCCCCGGATTTCGTTGA |
| BRDN0000737844 | AACGGCTGCGCCCGCGGCAA |
| BRDN0000737412 | AACGGGCGCAATACCCTTTT |
| BRDN0000737631 | AACGGTAGCGTACCCGTGAA |
| BRDN0000737750 | AACGGTCAAATCCGTGAGGG |
| BRDN0000737875 | AACGTCACCAACCTCGATCC |
| BRDN0000738229 | AACGTTATAGCTTCGTCTCT |
| BRDN0000737806 | AACTAACTCACTACGCACGA |
| BRDN0000738366 | AACTCCTCATCGTACGCTAA |
| BRDN0000737593 | AACTCGCGTGGGAAGTCCGG |
| BRDN0000738128 | AACTTATACGTAATCTGATC |
| BRDN0000738307 | AAGACTCCTACGTATCGAGC |
| BRDN0000737391 | AAGCACAAGAACGGTCCGCC |

**Supplemental Table 2. IEM genes and sgRNAs in the Inborn Errors of Metabolism Genes Library.**

| **Gene Target** | **sgRNA Target Sequence** |
| --- | --- |
| Abca1 | ACATGTCATCAACATAACAG |
| Abca1 | GTGGACCCGTACTCTCGCAG |
| Abca1 | CAAGCTGTCAAGCAACACTG |
| Abca1 | GGTATACACAGAGCCATTTG |
| Abcb7 | GATTAAATACTAACGCACTG |
| Abcb7 | CGCTGGGACGAACTCCATCG |
| Abcb7 | ACAGTTACTAGATGCTACAA |
| Abcb7 | TCACAGTTGCAGTTACACGG |
| Acadl | AGGCGATCGAGCTTCACGGT |
| Acadl | AGAGCGTACTCCAATTGCAC |
| Acadl | AACATCGCAGAGAAACATGG |
| Acadl | CCGGAAAATGTCATGCTCCG |
| Acadm | AAGATGTGGATAACCAACGG |
| Acadm | CCCGGAATATGACAAAAGCG |
| Acadm | TCGAACACAACACTCGAAAG |
| Acadm | AGCAGGTTTCAAGATCGCAA |
| Acadvl | CAAAGCATCTAACACGTCAG |
| Acadvl | CAACAACGGAAGATTCGGGA |
| Acadvl | GCTACATCGGATCCACTCGA |
| Acadvl | CCAGCGACTTTATGCCAGGG |
| Acads | GGTGACTCATGGGTCCTCAA |
| Acads | GCTCATGATAACTCCCGTGG |
| Acads | GATGGGCTTCAAAATAGCCA |
| Acads | GCTCACCCATCTTCTTAACC |
| Slc33a1 | GTGACTTACCTAAAGCCCCG |
| Slc33a1 | ATGTTATCCCGGGAAAACGT |
| Slc33a1 | GAAAGGGTAACGATTCCCCT |
| Slc33a1 | AAATATTGATGGCAGAACAC |
| Aco2 | AGATCCGTGCCACTATTGAG |
| Aco2 | GCGTTTACGGCCCGACCGGG |
| Aco2 | TGGGTGAGGTCCAACCAGAG |
| Aco2 | GATTGAAATTAACCTCAATG |
| Acox1 | CGATCCAGACTTCCAACATG |
| Acox1 | AATGTCGGATGGCTTGCGGT |
| Acox1 | CCTCACAGCACTGTATCGAA |
| Acox1 | TCTCTTCATAACCAAACTTG |
| Aspa | ATTCATTACCAATCCAAGGG |
| Aspa | AGTGCAACCCATGTTAGAAG |
| Aspa | ACCAACAGGATACTTGGCAA |
| Aspa | CTTGCCATATGAAGTGAGAA |
| Ada | GTTGTGGATCTTGTGAACCA |
| Ada | ATTCATCGGACCGTCCACGC |
| Ada | CTTCATCTCCACAAACTCGT |
| Ada | GCTGCGCAACATTATCGGCA |
| Adk | TGTGCTGCGTGTATCACTGG |
| Adk | CACTTTCAATACCGACTCTG |
| Adk | ATAAGGCATGACGTCCATCA |
| Adk | CATTGGGATAGATAAGTTTG |
| Adsl | AAAGTGTGTGAAACCTAACG |
| Adsl | CCAGTGCGATGCCGTACAAG |
| Adsl | GCTGCGGGCATTATTCATCT |
| Adsl | TCACAGGACAGACGTACACA |
| Aga | CCAGATCCCTCAAAATACTG |
| Aga | AGGCCAAGTGTTGACGACCA |
| Aga | TGGTGGACATTGCTATCTGG |
| Aga | ATGGATTACAACCATGCCTA |
| Gla | CTAAACTCACGTAATTTGCG |
| Gla | CATAGGATCCAAAACTCCCG |
| Gla | TCATACAGGTTATAAGTACA |
| Gla | CATTGGACAATGGCTTGGCG |
| Agxt | AGCCCCTGTCAGTTCCTACG |
| Agxt | AGAACATTATACACTGCAGG |
| Agxt | ACATACTTGGCCAAATTGTG |
| Agxt | GTAGATAGGAACCCCGCCCA |
| Aicda | GTAGGAACAACAATTCCACG |
| Aicda | TTCACAGAAGTAGAGGCGCG |
| Aicda | ACCAGGTGACGCGGTAACAC |
| Aicda | TGAGACCTACCTCTGCTACG |
| Ak1 | CTACACCCACCTGTCTACTG |
| Ak1 | GATCGACGGCTACCCGAGGG |
| Ak1 | GTCAGCTCTGGATCGGAGAG |
| Ak1 | CTTTGGCTAACATAGCATCT |
| Ak2 | TGAAGGCGACAATGGATGCA |
| Ak2 | GGTAGGACCGGCCACTCTTG |
| Ak2 | TCCGAACCGGAGATTCCGAA |
| Ak2 | TCAGCCAGTTTGGGTGCCTG |
| Alpl | GGATAACGAGATGCCACCAG |
| Alpl | CTAGGAGGCAGGATTGACCA |
| Alpl | ACCTAAGAGGTAGTCCACCC |
| Alpl | GTTGCATCGCGTGCGCTCTG |
| Akt2 | CATACTCCATCACAAAGCAT |
| Akt2 | TACAGAGAAATTGTTCAGGG |
| Akt2 | TGTGGTGTACCGTGACATCA |
| Akt2 | AGTGGACCACAGTCATCGAG |
| Alas2 | ATCCAAGGCATTCGCAACAG |
| Alas2 | ACATCACACAATTCCTCCAG |
| Alas2 | AATACTAAATAGGAACTGGT |
| Alas2 | AGGTACCAGCAAGTTTCATG |
| Aldh3a2 | CTGTATGCGATTGTTAATGG |
| Aldh3a2 | GGAAATCTTAGCAGCCATCG |
| Aldh3a2 | AGTCTCTGTCAATGTAACAA |
| Aldh3a2 | TCTGGAGGGAGGCTTCGCAC |
| Aldoa | AATGGCGAGACAACTACCCA |
| Aldoa | CCTTGCCCGGAGCCACAATG |
| Aldoa | CCACGAGACACTGTACCAGA |
| Aldoa | GCCAGCATCTGCCAGCAGGT |
| Gfer | AGAATTGGGTCGCCACACCT |
| Gfer | ACGACCTGGTGACTGACGCG |
| Gfer | CCTGCGTGGACTTCAAGTCG |
| Gfer | TTCCCGTGCTCCAAACCTTG |
| Ampd3 | TACCTGGTTCGCATGCACGG |
| Ampd3 | GTCCTCGGGGACAGTAAACA |
| Ampd3 | TGATCCGGGAGAAGTATGCA |
| Ampd3 | AGTAGCTCCGGGAACCAGAG |
| Mat1a | ACAGGTATGGTGCTACTGTG |
| Mat1a | CTATGCTACTGATGAGACCG |
| Mat1a | GTGTTGCACAGAGATGACGA |
| Mat1a | GACACATTGGGCAATATCTG |
| Slc25a4 | CAGTTTGACCCTCTCGATCG |
| Slc25a4 | GGAAGATCCCTTGCCCACGT |
| Slc25a4 | TTGTGTCGTGAGAATCCCCA |
| Slc25a4 | GGGGAAGTACCGGATCACGT |
| Ap1s1 | CCAAGAGCTCTACGTATCGG |
| Ap1s1 | GAGCTCTTGGACAAGTACTT |
| Ap1s1 | TCAGCCGGCAGGGAAAACTT |
| Ap1s1 | TGGCCACCTCAGACAAGGAG |
| Atg5 | AAGAGTCAGCTATTTGACGT |
| Atg5 | AAATGTACTGTGATGTTCCA |
| Atg5 | CCTTCTACACTGTCCATCCA |
| Atg5 | AAGAAAAACTCACCATTTCA |
| Apoa1 | GGAGCTCTACCGCCAGAAGG |
| Apoa1 | CAAGGAGGAGGATTCAAACT |
| Apoa1 | GTCCCAATGGGACAAAGTGA |
| Apoa1 | CTGGAAAACTGGGACACTCT |
| Apoc2 | AAGTTACTGGACCTCTGCCA |
| Apoc2 | AGAGGTCCAGTAACTTAAGA |
| Apoc2 | GGGAACCAGGAAGATGACTC |
| Apoc2 | GAAGACATACCCGATCAGCA |
| Apoc3 | GCAGGAGTCCGATATAGCTG |
| Apoc3 | CATGGAACAAGCCTCCAAGA |
| Apoc3 | GGATGCGCTAAGTAGCGTGC |
| Apoc3 | GCCCCGGACGCTCCTCACTG |
| Aprt | GAGTCCGGGTCTTTCAAGAG |
| Aprt | CTAACAGGTCTAGACTCCAG |
| Aprt | TGTGTGCTCATCCGGAAACA |
| Aprt | CGCACCTGAACAGCACGCCC |
| Aqp7 | GCCACTCACCATCATGACAT |
| Aqp7 | AGTCACTGCGGCATTCATGT |
| Aqp7 | ACACCCCAAGGACGGTAACA |
| Aqp7 | TTCCTGAATACATGACACTG |
| Ar | GGTGGAAAGTAATAGTCGAT |
| Ar | ACCAGGATACCACACTTCGG |
| Ar | GACTTGGGTAGTCTACATGG |
| Ar | GCTCCTGGGAGGTCCACCCG |
| Arg1 | AATAAACTTACTGTTCCCCA |
| Arg1 | AGTATGACGTGAGAGACCAC |
| Arg1 | AAATGACACATAGGTCAGGG |
| Arg1 | AGATGTACCAGGATTCTCCT |
| Arsb | GGTGGGCAGACTAGGTCTGG |
| Arsb | GAATGTCTGCCGACACGCCG |
| Arsb | AGCACAGACGTATTTATGCA |
| Arsb | GCCAGCACGAAGACCACATG |
| Arsa | AAGCACGTTAGGTTCTGACA |
| Arsa | AGGAGTCCCCAAATGGCCCA |
| Arsa | GTGACCTGGCCAGTAGACCA |
| Arsa | TCGGGAGAAAGACACATACC |
| Asah1 | GCAAGGTGTACGTTACCTAG |
| Asah1 | TAACATTTATAACATACCGC |
| Asah1 | AGTGATAAACCCTACCCACT |
| Asah1 | GAATATAAATAATAACACTT |
| Ass1 | TAACATCTTACCTTAATCTG |
| Ass1 | TGGGATCCTGGAAAACCCCA |
| Ass1 | AAGGCACTTCCTCACCAGGT |
| Ass1 | TGCCTTCACCTGTAGCAACA |
| Fxyd2 | ACCGGCTTACCGTACTCGAA |
| Fxyd2 | TATGAAACCGTCCGCAAAGG |
| Fxyd2 | GATGAGGAGGCCCACGACGA |
| Fxyd2 | CCCACTTACTGAGAATGATG |
| Atp6v1a | TGACTGCTGATATCCGACAG |
| Atp6v1a | AGTCGGCCATCATACTGACG |
| Atp6v1a | ATGTTGCCCCCACGTAACAG |
| Atp6v1a | CTTACGGGAAAAGGGCATCG |
| Atp6v1e1 | CCAACTTGATGAATCAAGCA |
| Atp6v1e1 | TTTGTAGGATCTGCTAAATG |
| Atp6v1e1 | TCAGTCTTTGCGTTTGCACA |
| Atp6v1e1 | AGGCATCCAAGCATACCTGA |
| Atp7a | TCTATAGGGCAAAACCTCCG |
| Atp7a | ACACGGTATTGGTTAAGACA |
| Atp7a | CAATTATAATCCACACCAAG |
| Atp7a | AGGGATCTTCTACTGCTCTG |
| Atp7b | AAATATAGGCCAACTCCCGG |
| Atp7b | AAAATGGCTCCCGACACTAG |
| Atp7b | AGTTGTGTCATTACCCACAT |
| Atp7b | AGACCATCAATCCCATGACG |
| Slc7a2 | AGGACGTCACTATTCCGATG |
| Slc7a2 | GAACGGAACAAGCATCTACG |
| Slc7a2 | GTATCTATACACTTACGTCA |
| Slc7a2 | CCGAGACAACATATTTGGCG |
| Auh | CAATGTCATTGATCACCGAT |
| Auh | GGTTTAGTTATCAAAAGCGG |
| Auh | GAGGAGGGGCTACAGCTCGG |
| Auh | CCCTCCCTAACCAGGCATTG |
| Baat | AGGACCAGAAAAGGCCCATG |
| Baat | AGGCCCATTGATAAGTACAG |
| Baat | GTAAAGGAAAGCCGCATCCG |
| Baat | AAGGCACACCACCTGAAAGG |
| Bcat2 | ACGGAACGAGCCTCTACGTG |
| Bcat2 | CAGGAACTATGGACCCACTG |
| Bcat2 | GTGGAGTGGAATAACAAGGC |
| Bcat2 | GCACAGAATGACGTACAGGA |
| Bckdha | CATGACCAACTATGGCGAGG |
| Bckdha | CAGCGAAATTGAAACCGGCG |
| Bckdha | CTTGTCTACAAACTCTGCAG |
| Bckdha | CTGGCCACGCAGATCCCTCA |
| Bckdk | ACAGCACGAGCTATACATCC |
| Bckdk | AGTCCGCATCAACGGACATG |
| Bckdk | GCGACCGGAATAGAGCATCA |
| Bckdk | TGTCTTCATGTAGCGCCAAG |
| Glb1 | GCGTAGGTAGTCGTAATCGC |
| Glb1 | CTTCCCGAGATGTATCGGAA |
| Glb1 | GGGTACTCACAAGTTCACGT |
| Glb1 | GCAGGACCTGTACGCCACAG |
| Blk | TTATTACAAAAATAACATGA |
| Blk | ACTTGAATAGTGCTGCACCA |
| Blk | CGTGAAAGATATCACCACCC |
| Blk | AAGGTCCCTGTCATTCACAG |
| C1qbp | ATCAGTCAAGAATTCAACGA |
| C1qbp | CGTACGCTGAGCAAACCGAA |
| C1qbp | GTTGAAGTTACCAAGACTGA |
| C1qbp | TCCTCCTCACCATCAAATGT |
| Cat | AACTTAATATCATGACCGCG |
| Cat | TGGCATTGAAAAGATCTCGG |
| Cat | GGAGTATCTGGTGATATCGT |
| Cat | AGTGACCGAGGGATTCCCGA |
| Cbs | ACACTACGATGACACCGCCG |
| Cbs | TCGCCATGCCACTCCCACGT |
| Cbs | GTGAGTTCTTCAATGCGGGT |
| Cbs | ATAATGTGGGGAGTCCGCCA |
| Scarb2 | ATAGAACAGGCCGAAATTCG |
| Scarb2 | AAATAAAACTGGATGTACAC |
| Scarb2 | AATATGATTAACGGGACAGA |
| Scarb2 | ATAATGACACGTACCAACAG |
| Chkb | ATGGTCCCAAACAACCATCG |
| Chkb | GGTGCTGCTACGACTCTACG |
| Chkb | CGTGGTTCGGTAGTGAGCAT |
| Chkb | CAGAGCCAATCCCCAATCGG |
| Tpp1 | TGAGTTTCATCGCTATGTAG |
| Tpp1 | TTATGGTAGAAGGTTACCTG |
| Tpp1 | AACCTGACAGCCAAAGATGT |
| Tpp1 | GATCGAGGCCAGTCTAGATG |
| Cln3 | ACACTCCGACTATCCAACCG |
| Cln3 | AATCGAGATGAGCTGTTGTG |
| Cln3 | GGACCCTGGAGGGGAAAACG |
| Cln3 | CAGGAGTTTGATGACAAGGG |
| Abcc2 | AAGACCCTGACTCATATCCG |
| Abcc2 | GCATTCGGAAGAAAGAACTC |
| Abcc2 | ACTCGGATCTTGGTTACACA |
| Abcc2 | GTTTAAGACGATCATGACAA |
| Cox6a1 | TGAGGGTAGGCAACGAACGG |
| Cox6a1 | CCGTGGGCGCCACTCGACAT |
| Cox6a1 | CGCCGCAGCTCGGATGTGGA |
| Cox6a1 | TCAACGTGTTCCTCAAGTCG |
| Cox8a | GCCTGACCGGCTCGGCCCGG |
| Cox8a | TTCGAGTGGACCTGAGCCCG |
| Cox8a | TCTGTGTAGGATATCACCAT |
| Cox8a | GCTCCCGCGCCGGCTTCGAG |
| Cp | CATATAAGCATCAATTAGGG |
| Cp | GCTGTGAGGAGCGACCTGGT |
| Cp | ATGAAAAGTGTAGATCCTAG |
| Cp | GCTGAACAAATACCACACGA |
| Cpox | TGGGCGCATAAGGATTCTTG |
| Cpox | GAAGCAGCGAACCAAATGAG |
| Cpox | ACCGACATACTTGAACCAAG |
| Cpox | CGGACCCGCCTCTAGCCCAG |
| Cpt1a | CACATTGTCGTGTACCACAG |
| Cpt1a | CATACTGCTGTATCGTCGCA |
| Cpt1a | ACCTTGGACCCAAATTGCAG |
| Cpt1a | ACGTTGGACGAATCGGAACA |
| Cpt2 | TCACTGGTCAAATAAGCCAG |
| Cpt2 | TCGGGAAGTCATCTAAGCAG |
| Cpt2 | AAATATTGGGACATATCCAG |
| Cpt2 | TTAAATACATATCAAACCAG |
| Crat | ACCACCCACGCATATAACCG |
| Crat | GCAGGCCAGATGCTACATGG |
| Crat | TCCACAAGTGCAACTATGGG |
| Crat | TTTGCTGCCAAACTCATCGA |
| Pcyt1a | AATACGTATCTAATTGTGGG |
| Pcyt1a | ATGCAAGACGGAACCTACAG |
| Pcyt1a | GCACCACCTCGTCCACGTAG |
| Pcyt1a | CTTTAGTAAGCCCTATGTCA |
| Ctsc | CTGCAAGATACAACCTCCTG |
| Ctsc | GGCAGTAACTGATAGCTGTG |
| Ctsc | TCACAACCACAACTTTGTGA |
| Ctsc | CTGTATTTCATCAGTCATCG |
| Ctsd | TATCCGTCGGACTATGACGG |
| Ctsd | TGACTCCAAGTACTACCACG |
| Ctsd | GACTGTGAAACACTGCGGCG |
| Ctsd | ACGTCCTTTGACATCCACTA |
| Ctsk | AAGTTGTATGTATAACGCCA |
| Ctsk | CACTCTCTATACCCCAGAGT |
| Ctsk | CAATACGTGCAGCAGAACGG |
| Ctsk | AAGCCCAACAGGAACCACAC |
| Cyb561 | ATGCCTATGACCATGCAGAG |
| Cyb561 | GCACGACCAAGATCTTGCAT |
| Cyb561 | TGCTGCACTGCCGTACTATG |
| Cyb561 | GTTCAGGAACTCACCCACCA |
| Cycs | TCTTCCGCCCGAACAGACCG |
| Cycs | TTCTGTTTAGGCATCACCTG |
| Cycs | GTTCGGGCGGAAGACAGGCC |
| Cycs | CTGGGGAGAGGATACCCTGA |
| Cyp11a1 | AGAGTATCGACGCATCCTTG |
| Cyp11a1 | GTATTATCAGAGGCCCATTG |
| Cyp11a1 | GGACCTAGGACTGCTAGTAG |
| Cyp11a1 | CCTTACACTCAAAGGAAAAG |
| Cyp11b2 | TGGAGAGTATGCTCCCCCGT |
| Cyp11b2 | GACCGTGTCAACGCTCCCAG |
| Cyp11b2 | AGGTGCTTCAAAATGCTCGT |
| Cyp11b2 | TTCCGCCACAATGCCACTGT |
| Cyp17a1 | GGAGCTACTACTATCCGCAA |
| Cyp17a1 | AAACGGTAGACTACCCACCA |
| Cyp17a1 | TAAAATTCGAGAAAAAACAC |
| Cyp17a1 | TACCATACAGACCTTTACAG |
| Cyp19a1 | AGTGACCGACATGGTGTCAG |
| Cyp19a1 | AAGGGCGAATTGTTCTCCAA |
| Cyp19a1 | GAGAGCTCGTCTTCAATACC |
| Cyp19a1 | CCATCAAGCAGCATTTGGAC |
| Cyp24a1 | TTCGTTGCGATGGTCCCGAT |
| Cyp24a1 | CTTGCTGATAAATATCACAA |
| Cyp24a1 | AGCGCCTCAACACCAAAGTG |
| Cyp24a1 | GCTGGACAAGAAAATCAATG |
| Cyp27b1 | GAGAGCGTATTGGATACCTG |
| Cyp27b1 | TGCGACGACTAAGGCGCCAG |
| Cyp27b1 | CTCAGGTGCATGGCGCTGCG |
| Cyp27b1 | TTAGCAATCCGCAAGCACGC |
| Cyp7a1 | GGTGAACCTCCTTTGGACAA |
| Cyp7a1 | TTTGATTTAGGAAGGCCCGG |
| Cyp7a1 | AGTATTTCCGGCACTAGTGG |
| Cyp7a1 | AGCACAAGAACCTGTACATG |
| Cyp7b1 | GTATGTCAGATACTAAGTAT |
| Cyp7b1 | ACTGTCAATTTCGTCACGCA |
| Cyp7b1 | TTTATCAAGGGTGGTTCACG |
| Cyp7b1 | AGCAGAGCTTCTTACCCACA |
| Slc6a3 | TAGATGATGAAGATCAACCC |
| Slc6a3 | GCTCGTCAGGGAGTTAATGG |
| Slc6a3 | CAGGGAGGGTGACTCCACGC |
| Slc6a3 | TTACTCAAAATACTCAGCAG |
| Dbh | AGTGATATAGCACCAGTACG |
| Dbh | AACTTCCAGTCGGAGAAACG |
| Dbh | AGAGCTCTAAAATCCCTTCG |
| Dbh | CGTGCCGGCAGTAGTTGAGT |
| Dbt | TAGATGATATCGCTTATGTG |
| Dbt | TCATCACAATACCCGAAGTG |
| Dbt | CTGTGAAGTTCAAAGTGACA |
| Dbt | TGGAGGCTTTGCTATCGGCG |
| Pcbd1 | AGCTGGACCACCATCCCGAG |
| Pcbd1 | ACTGCTTGAAGATAGCATCT |
| Pcbd1 | GGCACACAGGCTGAGCGCCG |
| Pcbd1 | TGCCAAACCTGAGGGCTGTG |
| Ddc | CCTAGGTGGTCGCTACACTG |
| Ddc | TAACCCAGCTCTTTCTACAG |
| Ddc | CCGGTATCTTCTGAATGGTG |
| Ddc | AGCCAGTAGGGCCACCAAGG |
| Ddost | GAACTCCCCGTACTTAATGA |
| Ddost | CCCAGATAAACCAATCACCC |
| Ddost | AGGCAACTATGAACTAGCTG |
| Ddost | GGTCAGAAACATCATAGTTG |
| Dgat1 | AGTGGTTTCAGCAATTATCG |
| Dgat1 | AAAGCGCTTTCGTATTCGGG |
| Dgat1 | ATACCCGGGACAAAGACGGG |
| Dgat1 | GCTCACCAATAATCACGCAT |
| Slc25a1 | ATGAACGAGCGAACCCACCG |
| Slc25a1 | GAATAATCTCTCTAACCCCG |
| Slc25a1 | ACTGCGACTGTACTGAAGCA |
| Slc25a1 | CTTCACGTATTCGGTCGGGA |
| Dhcr7 | CTGTAGCTATATCCAATGCG |
| Dhcr7 | TGTTCTTCAATGGACGACCA |
| Dhcr7 | GCCCTTGATCATTGCGAACG |
| Dhcr7 | AGCCTAGGTACCACCCAAAG |
| Dhfr | GACATGGTTTGGATAGTCGG |
| Dhfr | AACCTCAGAGAACCACCACG |
| Dhfr | TCGCCGTGTCCCAAAATATG |
| Dhfr | CAGCCCGGCCAATACCTGAG |
| Dld | GGTGGAACATGCTTGAACGT |
| Dld | GCAGTAAAAGCATTAACAGG |
| Dld | AGAGAAGCTGGTTGTTATTG |
| Dld | CAAAAACATCCTTGTAGCTA |
| Dpagt1 | CTTTGGCAATACAACCATCG |
| Dpagt1 | TCCTGTACTACGTCTACATG |
| Dpagt1 | CATGCCCGCAAAGTAACAGA |
| Dpagt1 | AGCACCGCTGATCACTCCCT |
| Dpm1 | ATGGAATCAAACACGCCACG |
| Dpm1 | TCTCTGGAACTCGATACAAG |
| Dpm1 | TGATTAATTTGGCAGCCGTG |
| Dpm1 | GAGAAGCCCCTTCTACCTCT |
| Dpm2 | TGATCAAGCTAACGGCGACA |
| Dpm2 | CTTCACCTACTACACCACTT |
| Dpm2 | ACCGGGACAGACCAAGCAGT |
| Dpm2 | ACATGCTGGCTGTCAATGAA |
| Slc26a2 | GAGCCGACACCATGACTCCG |
| Slc26a2 | ATGGCCGGAGAGCTTTCCGT |
| Slc26a2 | ACTGTGCCTTATGATTGGTG |
| Slc26a2 | TCCAAAATGAGAAGCCAATG |
| Ebp | GAAAGGCAATCACTACCCAT |
| Ebp | GGGGCCTAATTGTGATCACG |
| Ebp | CCAGGATATGCGAAGTCGGG |
| Ebp | AAGCTGTAGGACAAAGCGGA |
| Egf | TCACCGTAACAGATATGACA |
| Egf | AGACTGTTCTGGACGGACGT |
| Egf | TGTTCATCGCCTGACAATGG |
| Egf | GTGTCACAAAGGATGGATGG |
| Eno3 | AGCCGGTTACCCGGACAAGG |
| Eno3 | GAGACAAAGCACGATACCTG |
| Eno3 | CTTCCAGAAACTAAGTGTTG |
| Eno3 | CTGCAAGATCTGCGATGTGT |
| Ephx1 | CAGAAGTTCTACATTCAAGG |
| Ephx1 | TGGAAGCGACTGCCCTCCAA |
| Ephx1 | CCCCACCACCCATCTTCAAG |
| Ephx1 | GGAGGAATGAGTTTGACTGG |
| Epm2a | TGTACCAGAACGTGTCAACG |
| Epm2a | CCGTTGCTGCACATATAATG |
| Epm2a | GCCCGGCCTGTGGCTCGCCG |
| Epm2a | GTCACATGTTCCAGTTGGCG |
| Esr1 | GGCATACGGAAAGACCGCCG |
| Esr1 | CACTGTGTTCAACTACCCCG |
| Esr1 | TATTCAGAATAGATCATGGG |
| Esr1 | AGAGGCATAGTCATTGCACA |
| Esr2 | TGGCGCTTGGACTAGTAACA |
| Esr2 | TTCGTGACGGCTCTCTACAT |
| Esr2 | GAAGTAGGAATGGTCAAGTG |
| Esr2 | CGGCTCACTAGCACATTGGG |
| Ext1 | CTTATATCACGTCCATAACG |
| Ext1 | GATTGTATTAACTACACTAG |
| Ext1 | CTGACTACACTGAGGACGTG |
| Ext1 | CAAAGGCAAGAAGTGCCGCA |
| Ext2 | TGTTTCGATGTCTACCGCTG |
| Ext2 | CTGGAGGACTCAATGGAGTG |
| Ext2 | TCCTGCAGAACATCCCACAG |
| Ext2 | AGGGCAGTGTTGTAATCTGG |
| Fah | CCCCGATTCGAAGACCCATG |
| Fah | CCGATGGCTACACCAATCCG |
| Fah | ACATCAACATGTCTTCGATG |
| Fah | AATGGGAGTACGTCCCACTT |
| Fbp1 | CATGGCAAGGACCAACATGG |
| Fbp1 | CACAAGAACACAGGTAGCGT |
| Fbp1 | TGGCTCAACCAATGTGACTG |
| Fbp1 | AACATCTACAGCCTTAATGA |
| Fdxr | CCAGAAAACAGACATCACAG |
| Fdxr | TCCATTGTACCAGCCCACAA |
| Fdxr | GCTTCTCGTAGATGTCTACG |
| Fdxr | CAGATGTCCCCCGTCCAAGG |
| Fech | CATTTACAGATACTATAACG |
| Fech | CGTACTTACAGACATCGGCA |
| Fech | GGCATGATGTACCTTCTCCG |
| Fech | GGCATATTGATGTTAAACAT |
| Lpin1 | GGGGGTACGTTGAAATCGAG |
| Lpin1 | CAGTCGCCAACAATGGCCCG |
| Lpin1 | GACCACAGAGAGATCACCAA |
| Lpin1 | CCATTTAAAACAGGCCACTG |
| Folr1 | GACAATTTACACGACCAGGT |
| Folr1 | AGTTCGGGGAACACTCATAG |
| Folr1 | CCAGTTGAACCGGTACAGGT |
| Folr1 | CTCCACCTACTCCTTACCCG |
| Fxn | AGGGAACCGATCGTAACCTG |
| Fxn | CGAAGCGCGTACTCACGGCG |
| Fxn | GCGCGGGTTCCCGACCCAGG |
| Fxn | TGGGGACATTGGACAACCCA |
| Ftcd | CAGGTTAAGAGCAATACGGT |
| Ftcd | CCGAACAAAACCGCGCAGTG |
| Ftcd | GCACTGTCTACACTTTCGTG |
| Ftcd | CACAGCTCAAACAGGCCGAG |
| Fth1 | CACCATAGACAGATAGACGT |
| Fth1 | AGTGCGCCAGAACTACCACC |
| Fth1 | GGAGAGCGGGCTGAATGCAA |
| Fth1 | TCTTCAGAGCCACATCATCT |
| Ganab | GATAAAGTTAGTCTCGCGCT |
| Ganab | CAGACCCATATAGAGCCATG |
| Ganab | AAGTCCAAACCTACAGACGT |
| Ganab | ACTCGGTATCGAGGCCGCCG |
| G6pc | GGTGTTTGAACGTCATCTTG |
| G6pc | GTTGTCCAAACAGAATCCTG |
| G6pc | GGTCAGCAATCACAGACACA |
| G6pc | ACAAGACTCCAGCCACGACC |
| Slc37a4 | CTACGTTGACCAGACCAACC |
| Slc37a4 | TCTTTACTCCGAAGACCACG |
| Slc37a4 | CACAAACTTGCTGATGGCGT |
| Slc37a4 | CAGAGCGATCTCATCCACCA |
| Gaa | ACCATCCCCACTTTACAGCG |
| Gaa | GAGTTACAGGCCCTACGACG |
| Gaa | CTAACCTGGAGGTCAACGGG |
| Gaa | ACCTGAGCTCTACAGAGTCG |
| Gabra1 | CAGAGAGCCAGCCCGTTCAG |
| Gabra1 | CAGTGTCACTTACCATATCG |
| Gabra1 | AGAATAACTGTCATTATGCA |
| Gabra1 | ACGCAGGAGCTTATTAGGCA |
| Gabra6 | GGAAGTTAACCAATCTCATG |
| Gabra6 | GGGACTTCTACTGAGTAAAG |
| Gabra6 | TCTACTCTGAAAATGTCAGT |
| Gabra6 | TTAAGCTCAGAATCTCAGCA |
| Gabrb1 | TCTGTCCACAGGAATCACTA |
| Gabrb1 | CACTTCAGATGGCTATACCA |
| Gabrb1 | ATAGTCGTGGATATGCCCCT |
| Gabrb1 | CAAGAAATCATTTGTGCATG |
| Gabrb2 | ATGATTCGATTGCATCCCGA |
| Gabrb2 | TAACCAGCGACATATTACTA |
| Gabrb2 | CTTGACTTTGGACAACCGAG |
| Gabrb2 | GATGAGTTTATAATCTACGA |
| Gabrb3 | CCTCACGCTTGACAATCGAG |
| Gabrb3 | CCTGGTAGATGGCTACACTA |
| Gabrb3 | CGCCTGAGACCCGACTTCGG |
| Gabrb3 | CGAAAACTCAATGAAAGTCG |
| Gabrd | TGTGGCGCTTGCCCTAGAGG |
| Gabrd | GTAACTGGTGATAGTGAACT |
| Gabrd | AACCATACCAACGAGACCTT |
| Gabrd | TATCCGCCTACAGCCTGATG |
| Gabrg2 | AACAAACTTCGACCTGACAT |
| Gabrg2 | GGTTGAATAGCAATATGGTG |
| Gabrg2 | TCCTGCTATCGCTCTACCCA |
| Gabrg2 | TACAACTGGAGAACTCCAGG |
| Gad1 | CTATTCCATAAAGAAAGCCG |
| Gad1 | GACATTTGATCGCTCCACCA |
| Gad1 | GCGGTTGCATTGACATAAAG |
| Gad1 | AGATGAGAGAGATCGTTGGA |
| Galc | AGTTGCCTTATGGACGAAGT |
| Galc | CTAGATGAGAATTATTTCCG |
| Galc | GTCGGAGTCGTCTAGCACGT |
| Galc | CATAGCGAGCGATAATCTCT |
| B4galnt1 | TCTAGCAGATCGAGTCTCGG |
| B4galnt1 | TGACCGTAGGGTAAAAGCGT |
| B4galnt1 | GTTCGCAGGTCGGAACCTGG |
| B4galnt1 | TGCAGTTGTGAATCCAAGGG |
| Galnt3 | TACGGAAGCAACCATAACCG |
| Galnt3 | AGCTGCCCCTAGCAACCGCG |
| Galnt3 | AAAAACCGGCTTCAACTCCG |
| Galnt3 | GTGGACGGTCCTAAGCAGCG |
| Galt | AGAAAGGTACCATCATAGTG |
| Galt | GCAGCGTCACATCCGACCAG |
| Galt | GGGTTGAGTGGGTCGTGGCG |
| Galt | GGACGAGTGGGTGTTAGTGT |
| Gamt | AGACGCGTCATAGGCCGCGG |
| Gamt | CTATATGCATGCGCTAGCGG |
| Gamt | GATTATTGAGTGCAATGATG |
| Gamt | ATCAAAGTGACCGTCAGGCA |
| Gata1 | GTCACCGGCAGTGCTTACGG |
| Gata1 | AGTATGGAGGGAATTCCTGG |
| Gata1 | GAGCAACGGCTACTCCACTG |
| Gata1 | GGCCTCAGCTTCTCTGTAGT |
| Gba | CGTTACGAGAGCACTCGACG |
| Gba | GGATAACTGGAAGTCGTTAG |
| Gba | GACTGGCAAAGAGTGAAATG |
| Gba | CGGAGAATGAACCTACAGCA |
| Gch1 | GCACCAATGGGTTCTCCGAG |
| Gch1 | CCCTTCCTACAAATGGAACA |
| Gch1 | GGTGAACCTCCCCAAACTGG |
| Gch1 | GTACTTCACCAAGGGATACC |
| Gdap1 | TCCTTGTGGTTGCATAAGCG |
| Gdap1 | AAGATAATCAATGATCTGAG |
| Gdap1 | TGTGTAAGCATCCATTGGCA |
| Gdap1 | AGAAAACCCTGACTTACAAG |
| Gpd1 | GATGGGGTACCACAAAAACC |
| Gpd1 | ACCCAACTTTCGCATCACTG |
| Gpd1 | GTCTTCCTCAAACACCCACA |
| Gpd1 | GTTGGAGCTCACCACATTGG |
| Gfpt1 | TGTGGCACAAGTTACCACGC |
| Gfpt1 | AAGCTGCGGTCTTTCCCGTG |
| Gfpt1 | GGAGAGAGGAGCCTTAACTG |
| Gfpt1 | TCTGTTGTGAACACAATGAG |
| Ggps1 | GAGCCACTGGGAAACCACGT |
| Ggps1 | AGCCCAAGTGTGTCAAGCAG |
| Ggps1 | GGTGAGAAGCAAACTTTCAC |
| Ggps1 | TGTCCCTCCAGTAAATATCG |
| B4galt1 | GGCCAGAGAGGTAATAGACG |
| B4galt1 | CAGGGCTGGAGTCGAGACCC |
| B4galt1 | GCTCAATATTGGCTTTCAAG |
| B4galt1 | TGATGTGGACCTCATTCCGA |
| Ggt1 | ACTGACGTATCACCGTATCG |
| Ggt1 | CGTACAGGGTCGCATCACCG |
| Ggt1 | GAATTCAGGCTCATAGTAGG |
| Ggt1 | CGACCACGTGTACTCCAGGG |
| Gif | ACGTGAGAGCCATAACGGTG |
| Gif | CCTGGCCGGCGCCTACAACG |
| Gif | TACTCAACGAGATTAAGCAA |
| Gif | TATGTACAACAAGATTCCTG |
| Gclc | TGTGCCGGTCCTTGACTGCG |
| Gclc | CAATATGAGGAAACGCCGGA |
| Gclc | AGAAACATCCGGCATCGGAG |
| Gclc | TGTAGATGATAGAACACGGG |
| Galk1 | CTGGAAAGCCCACCCCCCAG |
| Galk1 | GGGCCGCGTCAACCTCATCG |
| Galk1 | AGTTGGTCCATGATGCCACA |
| Galk1 | ACAGTGGGCCAATTATGTCA |
| Glul | GATTACGGGGACAAATGCGG |
| Glul | TGGAAGGCCAACCAAATGGG |
| Glul | CTTGCCCAGAGTTACCTGAG |
| Glul | TATTTCTAGAGACCAACTTG |
| Glra1 | CCACTTCCACGAAATCACCA |
| Glra1 | TTCCATCGCTGAGACAACCA |
| Glra1 | ATGCTGCACCAGCTCGTGTG |
| Glra1 | TCCTGGATAAGCTCATGGGG |
| Glrb | AGGTTACTACACTTGTGTGG |
| Glrb | GGAAACAGAGTTAAGTCTAG |
| Glrb | CACAGCGTACTCCACGAGGG |
| Glrb | CCACCATGTATAAGTGCTTG |
| Slc6a9 | ATACCTCTGCTATCGCAACG |
| Slc6a9 | GTAGTACATGATACCCGTGA |
| Slc6a9 | ATGGTGGTGTCCACATACAT |
| Slc6a9 | TGTGCTACCAGCGTCTACGC |
| Gm2a | GATCCAACCTGACCCCATTG |
| Gm2a | ATTCCTTGTGTAGAACAGCT |
| Gm2a | CCACCAAGTTGGGAAAGCTA |
| Gm2a | GTGACAGGTGGAGCTCACCG |
| Bscl2 | GTTTCATGTTATACCAGAGG |
| Bscl2 | CCTGGGCCCACAGTAAGGCG |
| Bscl2 | CAAAGGATCAGACAAAGACG |
| Bscl2 | AATGTCTCACTGGCTAAGAG |
| Gnmt | TGTGTGGCAGCTGTACATCG |
| Gnmt | AGAACATTGCAAGCATGGTG |
| Gnmt | GAGCCATCCTTTGACAATTG |
| Gnmt | GGGCTTCAGCGTGATGAGCG |
| Gnpat | CCTTCGGCTTAGGAACTCCG |
| Gnpat | TACACCCCGCCCTTGTATAG |
| Gnpat | CTCACCAAACTTAGGAGTCA |
| Gnpat | ATGAGCCACAAACTGCGCAT |
| Got2 | TGGAGGTCCCATTTCAACAT |
| Got2 | TTTCTGCCCAAACCATCCTG |
| Got2 | CATCCTCCTCACCTTCACCA |
| Got2 | AGCTCACCTTCCGGACACTG |
| Gpaa1 | GTGGAGGCACTAACCCTACG |
| Gpaa1 | CCGCAGGATGCCATACACGT |
| Gpaa1 | GTCCCTCCACTGTCACATCG |
| Gpaa1 | AGAGTCCGATGGAGACAAAG |
| Pigq | CAGGATATGCTCAATAAAGG |
| Pigq | TGACACAGTGGCACGCAGCG |
| Pigq | GAAGGCCAGTCAAGTACCAG |
| Pigq | CAATAACCGAATTGGACAGC |
| Gsr | GGCTATGCAACATTCGCAGA |
| Gsr | CGGCCCCCACGATGACGCTG |
| Gsr | GCTGTGAGGGTAAATTCAGT |
| Gsr | AACATCTGGAATCATGGTCG |
| Gria4 | TGCATACATTGGTGTCAGCG |
| Gria4 | GCATGTCAGTGCGATATGTG |
| Gria4 | ACGTAGAGTTAATTACACAA |
| Gria4 | TGCACCTCTGACAATCACGT |
| Grin1 | AACATCACTGATCCACCGCG |
| Grin1 | GTGGACATCTGGTATCCTCG |
| Grin1 | CTGTCCTATGACAACAAGCG |
| Grin1 | AACCAGGCCAATAAGCGACA |
| Grin2a | ATGGTAAAAGAAGGCCCATG |
| Grin2a | AGAAGAAATCGTAGCCGGTG |
| Grin2a | ATCTTGACAAACTTCCGACA |
| Grin2a | TGTGTGCGACCTCATGTCCG |
| Grin2b | TATCCTACGCTTGCTCCGAA |
| Grin2b | GGCACCGGTTGTAACCCACA |
| Grin2b | ACATCATGGAAGAATACGAC |
| Grin2b | TGACTGGCTACGGCTACACA |
| Nr3c1 | CATTATGGGGTGCTGACGTG |
| Nr3c1 | AGCTTGCCTGGCAATAAACC |
| Nr3c1 | AAAGCCGTTTCACTGTCCAT |
| Nr3c1 | AAAACTGGAATAGGTGCCAA |
| Grm1 | GGGTGACAAATAGCGTCACG |
| Grm1 | TAGTGTACAGCTGACACTAG |
| Grm1 | ACTCTCCCACGACGCCAAGG |
| Grm1 | AGATAAGATTAATGCGGACC |
| Grn | CCTATCCAAGAACTACACCA |
| Grn | CCCTGCACAAAAGACCAACA |
| Grn | GACACTGGACAGCACCCAAG |
| Grn | CACCTAGTGAAGTCATCACA |
| Gss | GTTCTCTGGACCAAAACCGA |
| Gss | AGTCAGTATAATTCACAGGT |
| Gss | ACAGCTGGCTGGGACTAAGA |
| Gss | TCAGATTACATGTTCCAGTG |
| Gstz1 | GTCAGAAATCATGCGCACGA |
| Gstz1 | AAGAGACTCGGCCTATCCCA |
| Gstz1 | ATTTCTGACCTCATCGCTAG |
| Gstz1 | TAATGACTCACCGTTAAAGC |
| Gys1 | TTGGGACACCTGCAACATCG |
| Gys1 | CAACGCCCAAAATACACCTG |
| Gys1 | TGTTGTGGGTGCACACTGGT |
| Gys1 | ATCCCAGATCACCGCAATCG |
| Hadh | CCTTTCAACCAGCACCGATG |
| Hadh | AGCAAATCGGTCTTGTCTGG |
| Hadh | AAGCATGTGACCGTCATCGG |
| Hadh | TGGCCATACAGTAGTATTGG |
| Hsd17b10 | TGTTAATGATAACTCCACGT |
| Hsd17b10 | GCTACGGCCAAAAGACTGGT |
| Hsd17b10 | TTGGTCGCGGTAGTAACTGG |
| Hsd17b10 | GTCACGACTCACATTTGCTG |
| Hal | ATGCTGTCCGACCTCACCGT |
| Hal | AAAAGATACAGGATTGTCTG |
| Hal | GGACAATGGTGGCTTCACTT |
| Hal | TCTTAGCCAAAGGTTACAGT |
| Hao1 | CAGCATGCCAATATGTGTTG |
| Hao1 | ACTGTGGACACCCCTTACCT |
| Hao1 | GCAACGTTGCGAAGCATCCG |
| Hao1 | GTATGACTATTACAGGTCTG |
| Hccs | CAAAGTAAATGGCTGATCTG |
| Hccs | TAGCTCTAGCACCAGTAACA |
| Hccs | TCCAGAACATCTGTTCAGAA |
| Hccs | ATCAGAATAATGAGCAAGCG |
| Hcfc1 | GGTACCCTTCACAACCAATG |
| Hcfc1 | ACAGCAGTATGTGACTCCCG |
| Hcfc1 | TGGAAGGGCTAGTTGCACAA |
| Hcfc1 | CCCAAGGTTAGCCACAACAG |
| Hexa | ACGCCCCGGTGAGGGAATCG |
| Hexa | AGGAGGTCATTGAATACGCA |
| Hexa | AGTGAAGCTCTCATATGGGA |
| Hexa | AGCAAGGTGTTAATAACCCA |
| Hexb | GCTCAGCTCGAAATCTAGCA |
| Hexb | TACACCAAACGATGTCCGGA |
| Hexb | GCGGAGATGTACAACAGCCG |
| Hexb | TAACGCTCCCCAAACGCTGT |
| Hfe | TGCTCCACGTACCCTTACTG |
| Hfe | TTCCTCCCGCACTCACGCGG |
| Hfe | GATCCGTGCCAAACAGAACA |
| Hfe | CATGAAGACAACAGTACCAG |
| Hgd | GAAGATCGTCGAGTGCCAGG |
| Hgd | CCCTTTGAGTCCATCGACCA |
| Hgd | GGAGACATCAAGTCTAACAA |
| Hgd | GATGTCTTTGAGGAGACCAG |
| Hk1 | CCGACAATCCAAAATAGACG |
| Hk1 | CGTAGCCGCCATTGAAACGT |
| Hk1 | GGATCTTTACCAGTAGGACT |
| Hk1 | CTCCCGGGATTATAACCCAA |
| Hmbs | AAGATGAGGGTGATTCGAGT |
| Hmbs | CCTGGTCGTTCACTCCCTGA |
| Hmbs | AGAGAAGAGCCTGTTTACCA |
| Hmbs | CTCAGTTGCTATGTCCACCA |
| Hmgcl | CACCTCAGCAACTTTAGCCG |
| Hmgcl | CTTGGGAGAAACAAAGCTGG |
| Hmgcl | CTTGGGGACACCATCGGCGT |
| Hmgcl | GGGGTGGGTACAATACTCTG |
| Hmgcs2 | GACATTGCAGTCTACCCGAG |
| Hmgcs2 | TTTATTACGAACCTTGCTCG |
| Hmgcs2 | GAGGGATTGTAGAAAACCTG |
| Hmgcs2 | TCATATACTGCACATCATCG |
| Hmox1 | TCGTGCTCGAATGAACACTC |
| Hmox1 | TCAGGACCTGACCCCCTGAG |
| Hmox1 | ACGCTTTACATAGTGCTGTG |
| Hmox1 | TTCCTTGTACCATATCTACA |
| Hnf4a | GCAATGACTACATCGTCCCT |
| Hnf4a | AATGTGCAGGTGTTGACCAT |
| Hnf4a | AGCGGGACCGGATCAGCACG |
| Hnf4a | GGCTCCGTAGTGTTTGCCGG |
| Hpd | GAAAGACATCGCATTCGAGG |
| Hpd | CATTGTGGTGACCAACTACG |
| Hpd | AGATACCACACACACCCTGG |
| Hpd | ATGATCTCAAGGTTACATCT |
| Hpgd | GCTGCTAACCTCATGAAAAG |
| Hpgd | TACATGAGTAAGCAAAACGG |
| Hpgd | CATAGGCAAAGCCTTCGCTG |
| Hpgd | GCTGACCAGAAACAACTGAG |
| Lipc | TTATCATGATCATCCACGGG |
| Lipc | AGGAGAAAGGCGCTCGTTGG |
| Lipc | CATAACCCAGAGTGTTGCAA |
| Lipc | GGTGTAGTGCTGGTATGCCA |
| Hsd11b1 | CCTGGCAGTCAATACCACAT |
| Hsd11b1 | ACAGCGAGGTCTGAGTGATG |
| Hsd11b1 | AAAGAAAGTGATTGTCACTG |
| Hsd11b1 | GTAGGGAGCAATCATAGGCT |
| Hsd11b2 | CCACTCTTGCGTCACTCGAG |
| Hsd11b2 | GGTGATGAGCACCGCGCGAG |
| Hsd11b2 | GCTAAGAAACTGGATGCCAT |
| Hsd11b2 | CGCTGGCCTCAATATCGTAG |
| Hsd17b3 | CAACATTACCTCCGTAGTCA |
| Hsd17b3 | AAGCCTATTCATTTGAGGTG |
| Hsd17b3 | GCTGAGTACAGGCTGTAAAG |
| Hsd17b3 | TTTCTGCCAGTCAACAACGT |
| Hsd17b4 | TCTAACATAGGCTCTTCACG |
| Hsd17b4 | ACCAAACCGTACCAGTCACG |
| Hsd17b4 | TGGGCGCCATCGTCAGAAAG |
| Hsd17b4 | TGCAGTGAACGACTTAGGAG |
| Hsd3b2 | CTTCAGACCAGAAACAAGGA |
| Hsd3b2 | GATTGTCTTGAATGGCCATG |
| Hsd3b2 | ATATGTGGGCAATGTAGCCT |
| Hsd3b2 | ATTATTATGTTAGAAATGAG |
| Hspd1 | CGGAGAAGCTCTAAGCACGC |
| Hspd1 | TCTTGAACTAGGTGTGATGT |
| Hspd1 | AGGGACAATGGACTGAACAC |
| Hspd1 | TGACTTAGGAAAAGTTGGGG |
| Hspa9 | CAACTTGCCATACCTTACCA |
| Hspa9 | CTCAACCCAAGCATCACCAT |
| Hspa9 | GCCCTCCATAACAGCCACAC |
| Hspa9 | GTGAATCATTGAAATAAGCA |
| Ndst1 | CATCCAGGTTGACATACTTG |
| Ndst1 | CTATGTGACACGGCCCAGTG |
| Ndst1 | TTCTACAACGAGTACCCTGG |
| Ndst1 | CCCCACAGATATGGGCTATG |
| Hyal1 | TTGGCCAGAAACTTTAGTGG |
| Hyal1 | CCCCTACTATACACCCACAG |
| Hyal1 | CTGCAGGCAAATAAATACTG |
| Hyal1 | TGAGAAGTCAGGTTCAGGCA |
| Ids | GTCTCCATAACAGCCCAGGG |
| Ids | CATTCTCCTTGAAGTATTGG |
| Ids | TCCCTTGGAAAACATAACCC |
| Ids | TCCCTATGGACCAATTCCTG |
| Idua | TACCCCTATTTACAATGACG |
| Idua | GTTGGACAGCAATCATACAG |
| Idua | GGTCCAAGAATGCATCCAAG |
| Idua | ACTATGATGCCTGCTCTGAG |
| Inppl1 | AGAATCCGGTCACACCACGA |
| Inppl1 | CCTGGATATCCATGTCTAAG |
| Inppl1 | TCACACTGGACGTACTGACG |
| Inppl1 | GCAGGGCACAAACAAGACCC |
| Insr | CGAGGATTACCTGCACAACG |
| Insr | GTGATACCAGAGCATAGGAG |
| Insr | CACACTGCACCTCTCATCTG |
| Insr | TATAGCCAGACGGGCACTCG |
| Stt3a | AAGGTGGTACGTAACGATGG |
| Stt3a | TACCATGTAGAAATAAGCGA |
| Stt3a | GTTTACCTGGAAACCAACAA |
| Stt3a | ACTTTAATTATCGGACTACC |
| Itpa | ATAGCAGTACTCACATGTAG |
| Itpa | TGATAAGTGCCTGAGTACCA |
| Itpa | AAACGCCAAGAAGCTGGAGG |
| Itpa | TGGCCTCTGAAGAGAAGCAC |
| Itpr1 | CTGTGAAATATGCCCGACTG |
| Itpr1 | ACACGAACGATGTCATCGAG |
| Itpr1 | AAACGTGTCAATCTCTGCCG |
| Itpr1 | GGAAGCTGAGAACTCCACAG |
| Itpr2 | TCTGATACAGGATTCTACGG |
| Itpr2 | CAAGTGCCTTAACCGCACGT |
| Itpr2 | AATGCCGTAATCAACACCAG |
| Itpr2 | TGTTGTGAGACTCTTTCATG |
| Kcna1 | CGACACAATGGCAATAACCC |
| Kcna1 | TTCTGAACACCCTTACCAAG |
| Kcna1 | GGACTGGTAGTAATAAAGGA |
| Kcna1 | CGCAGCATTCGTGGTCGTCG |
| Kcnj10 | AGGTCGTCCATAGATCCTTG |
| Kcnj10 | TGGCCCCAGGAATACGCCGG |
| Kcnj10 | GCAACCCGGATCATAAGGCA |
| Kcnj10 | CTGCGCAATAAGAAGCACGA |
| Kcnj11 | CCAGGTACCGTACTCGAGAG |
| Kcnj11 | GAGTGGATGCTTGTGACGCA |
| Kcnj11 | CGGCGGGCGCATGGTGACAG |
| Kcnj11 | GAGGCACAACTTCGCCCTCG |
| Khk | TCCGATGCATTCCGGCCCTG |
| Khk | AACTGGAAGAGCTCCTCACG |
| Khk | TGTGGTGGACAAATACCCAG |
| Khk | AAGAGCCCATGAAGGCACAG |
| Lamp2 | GAGTGTAGTTGTAGTCGACG |
| Lamp2 | CCTGACAAGGCGACACACGA |
| Lamp2 | CACTTAAAGATGACATCCAA |
| Lamp2 | GCTGCAGCTGAACATCACTG |
| Lcat | GATGTGCTACCGTAAGACAG |
| Lcat | CCTGTCCTAGGTGACAACCA |
| Lcat | GTGCAGAATCTGGTTAACAA |
| Lcat | GCGGGGGGAAGAGCACATTG |
| Ldha | CAAGCTGGTCATTATCACCG |
| Ldha | GTTGCAATCTGGATTCAGCG |
| Ldha | GGAGAACATGGCGACTCCAG |
| Ldha | GTCATGGAAGACAAACTCAA |
| Ldhb | AAATTGTGGCCGATAAAGGT |
| Ldhb | GTCTTCCAACACATCCACCA |
| Ldhb | GCTCGCCCAGGATCCATCCG |
| Ldhb | GGGCTGTACTTGACGATCTG |
| Cog1 | GGGCTGGAACTCGATAATGG |
| Cog1 | TGAGGATTGGAAATCTCGAG |
| Cog1 | GTGGACATGCTTGTTCGATG |
| Cog1 | CAGCAGCAGCTTAATCTGGG |
| Ldlr | AAAATGCATCGCTAGCAAGT |
| Ldlr | GGTGTCGTAGGACAAGTTAG |
| Ldlr | GCAGACTGGTGTACTCGCTG |
| Ldlr | TGACCGTGAACATGACTGCA |
| Lfng | GGCGATGAAGACGTCGCGAG |
| Lfng | AGACGCGGATCCACCGCCCG |
| Lfng | CACACCCAAGACGTGTACAT |
| Lfng | GGAGAGGCTATGCACCTCGC |
| Lipa | ACCGAGATAATCATGCGCTG |
| Lipa | GTATTCACCGAATCCCTCGT |
| Lipa | ACTTCAGCATCGCACTCTGA |
| Lipa | TGTAGTTAATTGAAGCAGGT |
| Lipe | GAGTATGTCACGCTACACAA |
| Lipe | CACTTAGAGAGTACGCTCAG |
| Lipe | TGCGGTTAGAAGCCACATAG |
| Lipe | AGAGCGGATATGCCTTGCAG |
| Phyh | CTCATTATAACGATCCCTGG |
| Phyh | CCCAAAGGATACAATCGTTG |
| Phyh | AAACAATTAGGTTGCTAGGT |
| Phyh | GAGCACATTGACAGAAACAA |
| Lpl | GAAAAACGTACCGTCTGCTG |
| Lpl | CCATCCATGGATCACCACGA |
| Lpl | TGGATTCCAATACTTCGACC |
| Lpl | TGACACTGGATAATGTTGCT |
| Ltc4s | CTTGCAACAGAACTCCCACG |
| Ltc4s | AGACGCGCTCGAACTCGGGA |
| Ltc4s | CGTATCCCTGGAAATAGCGG |
| Ltc4s | TACAGGTGATCTCTGCACGA |
| Alad | AAATGGAGCATTCCTAGCAG |
| Alad | ACGGACAGCCTCAATAGTTG |
| Alad | GCTCCGTCAGACATGATGGA |
| Alad | AAAACATGGACTTGGCAACA |
| Amacr | GCAGGTCATCGATTCAAGCA |
| Amacr | GGACAAAACATCTTAGATGG |
| Amacr | GCGGCGCATGTGCGCACGCG |
| Amacr | AAGCCAAATAGTTGATGTCA |
| Man2b1 | ATGTTGTAAATGACTACGCG |
| Man2b1 | TGAGTTCAATGCAAAAACGT |
| Man2b1 | CTTGATAGTCAATGCGCCCA |
| Man2b1 | AGCTCCCAGAGGTAACACGT |
| Man2b2 | CAATGTCTACACTACCGTGG |
| Man2b2 | TGGTGGCCAACGTTAAACAG |
| Man2b2 | CAGGCCTTGTGTCTACCGAG |
| Man2b2 | TCAAGTCATGCACGCCCGCG |
| Maoa | TCTTGGCAGTCAAAACCGGT |
| Maoa | TAGGAACGGAAATTTGTAGG |
| Maoa | GTTGTAATCCAAATATGCCA |
| Maoa | ATGGCAAGCAAGACATGCTG |
| Mc2r | ATGGGTTATCTTAAGCCTCG |
| Mc2r | ACAATCGGAGTTATTTCTTG |
| Mc2r | GGCGCATGGTCACAATGCTA |
| Mc2r | AGATAGAGCCCAGCAAAGAG |
| Rdh11 | CCGGCAAGCTAAATACACAC |
| Rdh11 | GCACCTTCTCATCAACAATG |
| Rdh11 | GGAACAGTCAGGTCTTCGTA |
| Rdh11 | TCCACTTCCATAACCTGCAG |
| Mocs2 | ATCATCCCGCCAATCAGTGG |
| Mocs2 | GAGACACTGCACCACACAGA |
| Mocs2 | GAATACTGCTATGTGTCTCA |
| Mocs2 | ATATGAAGCGTATGTACCGA |
| Mdh2 | TCACGCACCTGAGAGATCAG |
| Mdh2 | CGATATCGTAGAGGGTCAGG |
| Mdh2 | GTTCGCTCTGACGATGTCAA |
| Mdh2 | GTTGGCAATGATGCAAACCA |
| Mpv17 | CATTGAGTATCCCGACCAGT |
| Mpv17 | TGTACCAGCCTCCGACGACA |
| Mpv17 | ACAGGATCACTGATGGGCGT |
| Mpv17 | GGGATACCATGGTCAGAGTG |
| Mthfr | AGACCCTGTAGGTGACCACT |
| Mthfr | GACTGGGATGAGTTTCCTAA |
| Mthfr | AGGTAACATCTACGAAGAGG |
| Mthfr | CGAAGCTCTCTGCATCGGGG |
| Mtm1 | AGGGAACCACCAAAAGAGCA |
| Mtm1 | TTACCACTCATACCGACAAG |
| Mtm1 | ATGGGAGGCGCGACAAGTAG |
| Mtm1 | TCTGACCGGTGCCATTCAAG |
| Mttp | TGAGCGGTCTGGATTTACAA |
| Mttp | TGATCAAGTGATCCAAGTCA |
| Mttp | GATATACCACCAGAATCGTA |
| Mttp | ATCCTTTGCAGACACGCTCG |
| Mut | TTTGACTTGGCAACACATCG |
| Mut | TTATATGGCACACCCCAGAA |
| Mut | AGAGTTCGTATGCAAAGACT |
| Mut | ATCATGTACGACCCTCCCCA |
| Mvk | AGCGTCAATTTACCCAACAT |
| Mvk | GTGGTCGGAACTTCCCCCCG |
| Mvk | CAAGGTCCCGCGGAGTACCA |
| Mvk | TCTGAAGTCAATCAACAAGT |
| Naga | GCTAGGAAGGCAATCCCGTG |
| Naga | TGTTGAGGTATACATAGCCC |
| Naga | TCTCACAAACCAGTCCAGGA |
| Naga | TGTTCAGGTGAACTACACCG |
| Ndufa2 | GCAGGGATTTCATCGTGCAA |
| Ndufa2 | TCTGATCCGCGAATGCTCGG |
| Ndufa2 | ATTCGCGGATCAGAATGGGC |
| Ndufa2 | GGCTGCCGCTGCTAGCCGAG |
| Ndufa4 | AAAGACCGTGAACTTACGCT |
| Ndufa4 | AGCACTGTATGTGATGCGCT |
| Ndufa4 | CACTGTTTAATCCAGATGTC |
| Ndufa4 | TCTTCGTATTTATTGGAGCA |
| Ndufs4 | CCTGGATGGAACTCTACAGA |
| Ndufs4 | AGAGCACATCAAAACCAGAA |
| Ndufs4 | GTTGATGCCCAACCCATCAA |
| Ndufs4 | GTTGTCTGCCAGCTTCCAAG |
| Neu1 | ACAGCCTTCATCGTAGACGA |
| Neu1 | TTGGTTTGGAGTAAGGACGA |
| Neu1 | TGTGTGTGGACACGGGACGC |
| Neu1 | GAAAAAATCTGCATCCGATG |
| Neurod1 | GAGCTGTCCATGGTGCCGTA |
| Neurod1 | CCGGCGACCAAATTGGTAGT |
| Neurod1 | AGCAAGGTACCACCTTGCGC |
| Neurod1 | GTGTCTCAGTTCTCAGGACG |
| Nfs1 | CTAGTGAAAATGATCTCCCG |
| Nfs1 | TCACAGTCATAACGGAGACC |
| Nfs1 | AGGGCCTCCACACGTACCCG |
| Nfs1 | CCTCTGTTAGTCCTTACCAG |
| Npc1 | GGGGAAGGTGATCACAAGCG |
| Npc1 | CATCATGTGGGTCACCTACG |
| Npc1 | CGGTTCGTAGATATGAACAC |
| Npc1 | CCATCCCTACCTGAAAACAC |
| Slc11a2 | ATGTCACCGTCAGTATCCCA |
| Slc11a2 | AAACACAAAAGTGTCTGCGA |
| Slc11a2 | TGAGAAAATCCCCATTCCTG |
| Slc11a2 | CCTTGACTAAGGCAGAATGC |
| Nsdhl | TAGGCTTCATGGCGTAAGGG |
| Nsdhl | CTATAAAGAACTGCACCCGG |
| Nsdhl | CTTGGGCCGAAAATGCCATG |
| Nsdhl | CCTGTGTACCCACAGAAATG |
| Oat | CCGTTCAAAAATGTACTCGG |
| Oat | TGTACTCCTCGTATTCACCA |
| Oat | ATCATAACTGGTCGGATCTG |
| Oat | ACCCTGATGTACCTCCAGTG |
| Ogdh | TTGGCCCACTCATAGATACG |
| Ogdh | GACTAGTTCGAACTATGTGG |
| Ogdh | GTAAGTGGAAGACCTTGTCA |
| Ogdh | AAAGCTGAACAGTTCTACTG |
| Slc25a15 | ACCTTTCCAGACCTCTACCG |
| Slc25a15 | GCCCATGGTAGAAGCCCAAG |
| Slc25a15 | GCACAGCATGCGTACTGACT |
| Slc25a15 | GGACAGCACTTACTTCTGAC |
| Otc | CAGTCCATTGACAATTGGGA |
| Otc | CCTTCAAGCAGCTACTCCAA |
| Otc | TAGAAAGGGTCACACTTCTG |
| Otc | AAATTCAGGATCAAGCAGAA |
| Pah | CCTCTTCTGGAAAAGTACTG |
| Pah | CACTTACCTCAAATAAGCGC |
| Pah | GCAGCATCATCAAGAGCCTG |
| Pah | TCCTCGGGTGGAATACACAG |
| Pax4 | TTGGAACCCAAGTGTATTGG |
| Pax4 | AGGGTACTCATCCTTTAGCT |
| Pax4 | ACTCAACTCAGATCACCAGG |
| Pax4 | GTCTCTACAGAGTTTCAGCG |
| Pck1 | ACTGACAGACTCGCCCTATG |
| Pck1 | GTGGCCGAGACTAGCGATGG |
| Pck1 | CCTTTGGAAGCGGATATGGT |
| Pck1 | TCGCAGATGTGGATATACTC |
| Pdha1 | AGAACAACCGCTATGGCATG |
| Pdha1 | ATCACTGCCTATCGAGCACA |
| Pdha1 | GCGCCGGATGGAGCTAAAGG |
| Pdha1 | TGTTTGACATTATACGGCGA |
| Enpp1 | TACAACGCAAGTTGCCACTG |
| Enpp1 | GGTGACCGCTAATCATCAGG |
| Enpp1 | ATGTGAAAGCATCGATACCC |
| Enpp1 | AACGTCTTGGTAGGGTACAT |
| Pdx1 | GACCCGTACTGCCTACACCC |
| Pdx1 | ACTGCCAGCTCCACCCGGCG |
| Pdx1 | AATCCACCAAAGCTCACGCG |
| Pdx1 | CCATTCGGGAAAGGTCCGGG |
| Pepd | CACAAATCGGATCTCCAGCG |
| Pepd | CTGCTATGGTGTCATCGATG |
| Pepd | GCCCTGCAACACGACAGCTG |
| Pepd | CTTGCTAATGCCCTCGAAGG |
| Pex11b | CTCTACAAGTGCTACGCCTG |
| Pex11b | CAAAGTACAAGGCTCGGTTG |
| Pex11b | GGCGAATCTCATAAGCATCA |
| Pex11b | ATCTCAGAACGACATCTGAC |
| Pex16 | GCTTCGAAAAAAGTTGCCTG |
| Pex16 | TGAATCGGAGAAGCGACCTA |
| Pex16 | TGAGAGCAATGACGAGCCAA |
| Pex16 | GCCAGGAGCCATCATATGTG |
| Pex7 | GATGCCGTAGTGCTGCGCCG |
| Pex7 | ATGGGATCAAACTGTCAAAG |
| Pex7 | CATCACCACTACAGGTGACA |
| Pex7 | CTATAAAGAGCACACGCAGG |
| Pfkm | CCTCACGGTAGAGCGAACAG |
| Pfkm | GCGCCTTGGATATGACACCC |
| Pfkm | TTAGACCAAAGACGTGACCA |
| Pfkm | CATAGACACGCTCTCCCACG |
| Pgk1 | TAAGGTGCTCAACAACATGG |
| Pgk1 | TCAAGAACAGAACATCCCTG |
| Pgk1 | GGACTGCACACCGAGCCCAT |
| Pgk1 | CTTCCTCTACATGAAAGCGG |
| Pgr | CTCTGGCCGACTCATGAGCG |
| Pgr | AGGAGGAGTCGCAGCCAACG |
| Pgr | GGCGCGAACGAACCCTGCGT |
| Pgr | CAGTCTGGGAAGTCACCGCA |
| Abcb4 | GATGACATCTGCGTTCCGGA |
| Abcb4 | AGGACCGTGATAGCTTTCGG |
| Abcb4 | TAGCGAAAGCATCAATACAG |
| Abcb4 | TACTACTATTCGGGACTAGG |
| Phka1 | GTAGAAAACCTACGATTCAG |
| Phka1 | AGTGTAGTGAAGTTAATGAG |
| Phka1 | GAGATAGGGAAAGTGATCGT |
| Phka1 | CTAAACTAGCTCCTACCTCA |
| Piga | TCTTCCACGCCAAGACAATG |
| Piga | ATGGGTGCAGGTCCTATCGT |
| Piga | CAGACTGTGAAAGAGAGTCG |
| Piga | CCATGCTTATGGAAATCGAA |
| Pik3r1 | GAGCTTTATAAGGAGAGGCG |
| Pik3r1 | TCCATTAACCTTCAACTCTG |
| Pik3r1 | TGGCTACAATGAAACCACTG |
| Pik3r1 | CTGGAAATCTGAAAAGCACG |
| Pip5k1c | TGGTGGCAAGAACATCCGCG |
| Pip5k1c | CCGACGCATCCACGCCTCGG |
| Pip5k1c | TTACCAAATAGTCATCTGGA |
| Pip5k1c | GCCATGGAGTCTATCCAGGG |
| Pklr | TGTACGAAAAGCCAGTGATG |
| Pklr | GGGTTCACTCCAGACCTGTG |
| Pklr | GGGCGATGCAAAGACAGTGT |
| Pklr | CTGGTGACCGAAGTGGAACA |
| Plcb1 | CAAATACTTACGAATCCACG |
| Plcb1 | CCTGCACAGGCAATATCCGG |
| Plcb1 | GGAACAGCGCATGATAACTG |
| Plcb1 | TCCCAACTTTCTTATCAGAG |
| Plcb3 | GAAGGTGGGCATCTACGTCG |
| Plcb3 | CATACCGGGTATTTGTCGAG |
| Plcb3 | TTGATAAAATATGCGCTCAG |
| Plcb3 | CAGTGAGGTCAATGCCACGG |
| Plcb4 | TGAAGTAATGAGCCAGCGGG |
| Plcb4 | TAAGAAGATCGGGACATACG |
| Plcb4 | CAAGTACGGATGGATGTTCG |
| Plcb4 | CTTGGCGAATACTGTTGATG |
| Plcd1 | ATTTGCCAGAGACATACCCG |
| Plcd1 | GGGAAGAAGACACTAAGTAG |
| Plcd1 | AGGTGGATGACAGCTACGCC |
| Plcd1 | GTAGCGCTCAATGAGAGAGA |
| Pnp | AGATGCTGTGTGATGATGCA |
| Pnp | TCAGTGCCTGGAAACAAATG |
| Pnp | TGTGGCCAGAACCCTCTCCG |
| Pnp | CCTCAAGTGGCAGTGATCTG |
| Polg | TGCTCATAAACGTATCAGGT |
| Polg | CGGCGGGGAAATGCCCGACG |
| Polg | CCGTTGCCATGGTGATACGT |
| Polg | AGATCCTGGCCCGCCCAGCG |
| Por | TGGCTCCCAGACGGGAACCG |
| Por | ATGTCTCTAAACAATCTCGA |
| Por | AAGAGGATTTCATCACATGG |
| Por | TCCAAGACTACCCGTCCCTG |
| Ctsa | GTGGTGCTTTGGCTTAACGG |
| Ctsa | GGACTCGATATACAGCACGT |
| Ctsa | CTCCGGCTACCTCAGAGCAT |
| Ctsa | CATGACCAGTACAGCCAAGG |
| Ppox | CCAGCCGGCCTAATCCCTCG |
| Ppox | CCGGGCCTGACGAATTAGTG |
| Ppox | GTACTCAGGATTCGAGCTAG |
| Ppox | TAATAGTGAGGTGTTACCTG |
| Inpp5k | TTGTGCAGGAGAGTATAACG |
| Inpp5k | ACAGCGTACATGTTGTGACG |
| Inpp5k | AGGATGCGGTCAGTCCACGC |
| Inpp5k | TACCTTGACAAAGTTCAGTG |
| Ppt1 | GGTACACACTCTCCTTGATA |
| Ppt1 | TGTTAATGTCCAAGTCAACA |
| Ppt1 | AGAGACAGGACGTAAATCCC |
| Ppt1 | CCATGCCAGATCACCAGCGG |
| Prkcsh | AAGACCAGGTAGAAACACTG |
| Prkcsh | CAGGGACAAGTACCGCTCTG |
| Prkcsh | GGCACAGACGAGTACAACAG |
| Prkcsh | AACTTGACGACAACATGGAT |
| Prodh | AGGAGGCGTATCGCAGCCGG |
| Prodh | CCTGCTGTCACGGTTCACTG |
| Prodh | CAGGATAAAGCCAACACCAA |
| Prodh | TCTCCCAGGAGCAAATAAGA |
| Prps1 | GATGACTGCAGTAACCCGGC |
| Prps1 | TGCAGATCATATTATCACCA |
| Prps1 | CATCTCCCACAAGTACCATG |
| Prps1 | ATGTCTACATTGTTCAAAGT |
| Psap | CTACGTGGACCAGTATTCCG |
| Psap | GTCAACCACCTCCTTGCACG |
| Psap | CCTCAGCTAACCTTAGGTTG |
| Psap | TCTGGCATAAAATCACATTG |
| Pten | CCTCCAATTCAGGACCCACG |
| Pten | TGTGCATATTTATTGCATCG |
| Pten | ACTATTCCAATGTTCAGTGG |
| Pten | GGTTTGATAAGTTCTAGCTG |
| Pts | GGGAAATGCAACAATCCGAA |
| Pts | CGAGGCGCGACAGTCGCGCG |
| Pts | CGGGCACAACTATAAAGGTG |
| Pts | GTGATCAAGAGGCTTCATGA |
| Pex19 | ACAGCACATCCTTAGACAGG |
| Pex19 | GCTGGCTTCCCAAGCTACTG |
| Pex19 | GATGGTCGGAGCATGTTCTG |
| Pex19 | TGGAACTGCTCCACCAGATG |
| Abcd3 | GCTCACACGGTACCTCTACG |
| Abcd3 | AACCAGGTACCCGACGACAG |
| Abcd3 | GTGAAATGACTAGATTGGCT |
| Abcd3 | AGAATGGGACGCTCATTGAG |
| Abcd4 | AGACTCCATAATACTGACTG |
| Abcd4 | GGACGACATTGATAATCCGT |
| Abcd4 | ATGTTGCCGGTTACACACAC |
| Abcd4 | CATCTTTGGATATTTCATCG |
| Pex2 | GTATGCTGTGTGCACCATTG |
| Pex2 | TCCCAAAAGACGCTAAATGA |
| Pex2 | GTTCTGGGGCTTGCAAAATA |
| Pex2 | ATGAAAGCACTGAGTAAACT |
| Pex5 | CTGGACTCACCATCGATCAG |
| Pex5 | TCTTGTAAACTGATCAACCC |
| Pex5 | TCGTGCGGCAGATTGGCGAG |
| Pex5 | AGAAGGGCTGCATCGACTGG |
| Pygm | GTAGCCGCCAACATTGACTG |
| Pygm | ACTTGGAGGACTTGAAACGT |
| Pygm | GGATCCAGCGTCCCACGAGG |
| Pygm | GCAGCCTATCTACGTCCCCA |
| Rbp3 | TAAGCAGGTACACTCCACGC |
| Rbp3 | TTACGAGCCCAGTACCCTCG |
| Rbp3 | CAGCACTGTATCTCTCCCCG |
| Rbp3 | CTGCAATCTAAGTTGGCCCA |
| Rbp4 | AACTTCGACAAGGCTCGTGT |
| Rbp4 | CTTGCAAAAAGAGACCCTCG |
| Rbp4 | AGGGACGAGTCCGTCTTCTG |
| Rbp4 | CCCACTACTCACTTCCTCGC |
| Rdh5 | TGATATCCAGTAGTGTTGTG |
| Rdh5 | ATGGAGACTTGTACTCCGAA |
| Rdh5 | AAATCATCCTGTGTTAGCCA |
| Rdh5 | AAGTGGGTGAAGACACGTGT |
| Rlbp1 | TCGGGACAAGTATGGTCGAG |
| Rlbp1 | ATCCGGCCTCGATAGTGCAG |
| Rlbp1 | CCGTGCCCGCAAGTTCGATG |
| Rlbp1 | ACCTCACCTTCTGCAAAGTG |
| Rnaseh1 | TGGGCGACAGACAAACCAGA |
| Rnaseh1 | AGAGTCAGTCGTTGTCTACA |
| Rnaseh1 | AGCAGGAAACCGGTCCACCT |
| Rnaseh1 | AGGAGCTCTTCAAGCCCGGA |
| Rpe65 | CAAAGAGCCCTGGCCCACAT |
| Rpe65 | TCGAGTCCAATGAAAGCATG |
| Rpe65 | CCTTGTAAATATCTACCCAG |
| Rpe65 | TCAAGCCATCTTATGTACAC |
| Rpia | ACTGCACGTGGGAACCCGAG |
| Rpia | CGTGGTTCTCCACCGCCGTG |
| Rpia | ATGATGTACCTGGAAAGATG |
| Rpia | TGACCTGGATCAACACCCAG |
| Nr1h4 | AAACGGGACATTGTTGTATG |
| Nr1h4 | TGATGGACATGTACATGCGC |
| Nr1h4 | TGTGACAAAGAAGCCGCGAA |
| Nr1h4 | AGTGTAAATCTAAACGGCTA |
| Scp2 | ACAGATCCCTTACTCCGCAG |
| Scp2 | AAGTGGGTCATCAACCCTAG |
| Scp2 | CTCCTCGCTGGACAGAATCG |
| Scp2 | GCTGTATTCATCTTGGAACT |
| Sord | CAGTACTCATCTACTTCTCG |
| Sord | GTTACTAAAGATGCACTCGG |
| Sord | TGTGCTTGTAGAATCGGCAG |
| Sord | TCTTACCAGCGCCACACACA |
| Selenbp1 | AGTCATGGTCAGCACCTTGG |
| Selenbp1 | CATCTCCTCCCGCATCTACG |
| Selenbp1 | TGCAGGAAGCGGATCTCCAG |
| Selenbp1 | CCAGCCTCGACACAATGTCA |
| Sgpl1 | ATATAAAATCCCACTCCATG |
| Sgpl1 | AGGCTTATGGAGAATTCACG |
| Sgpl1 | ATTGCACCAAATATGAGCCC |
| Sgpl1 | GGGAACGGAAAGCATCCTGA |
| St3gal3 | GCTGGACAAACCCTAGGCAC |
| St3gal3 | CCTACGCATCACCTACCCTG |
| St3gal3 | ACGACTATGACATTGTGATC |
| St3gal3 | GATCCTAGCCCACTTGCGAA |
| St3gal5 | GTTCGGGCAGCATATCCAAG |
| St3gal5 | TAGTATTCAACGTCCGACAG |
| St3gal5 | CGCCCTCAACCAGTTCGATG |
| St3gal5 | AGCTGAGAAGTGATTGCTCA |
| Clpb | AGGACCGCGTTCCGACGAGG |
| Clpb | GGAGAACGGCTGGTACGATG |
| Clpb | CGTTGTCACCGGAGACCGCG |
| Clpb | CTCTCGAGTGACTAGGACTG |
| Slc10a1 | TACAGCAAAGGAATCTACGA |
| Slc10a1 | TTAACCCTCGGTCCTACCTG |
| Slc10a1 | AGGACGTAGGGTACATAGTG |
| Slc10a1 | GAGGGGCATGATACCGTACT |
| Slc10a2 | CTATTGGATAGATGGCGACA |
| Slc10a2 | GTTGCTCTCAGGTACTACGC |
| Slc10a2 | TAGGACATATAAAGAGACCA |
| Slc10a2 | GCTCACCATCCTCTTAGCCA |
| Slc12a3 | AACCTGGTACCCGACTGGAG |
| Slc12a3 | CGGTTACAACACCATAGACG |
| Slc12a3 | TGTGGTCTTCCACCTCGTTG |
| Slc12a3 | CACAGGCTAGCCCTTCGCAG |
| Slc16a1 | ACTACTAAGAAAGACCAAAG |
| Slc16a1 | CACCAGCGATCATTACTGGA |
| Slc16a1 | GACTTGCAGCCAACACCAAG |
| Slc16a1 | AGGCCCTATTGGTCTCATCA |
| Slc1a1 | CGACTCACCTAGTACCACGG |
| Slc1a1 | TAGGATTACAGCAATGACGG |
| Slc1a1 | ATCATGCTGGATACGATCAG |
| Slc1a1 | TCACCTGATCAGGTCCAACA |
| Slc1a2 | CATGTTGATAGCCTTCCCGG |
| Slc1a2 | CCATAGCTCTCGTGCCTAGG |
| Slc1a2 | TAATTGCCCATAGGTCTGAT |
| Slc1a2 | GTTCATGGTTTCATTCAACA |
| Slc1a3 | GTATAAAATGAGCTACCGGG |
| Slc1a3 | GACTCTGACCCGGATCCGGG |
| Slc1a3 | GAGGCCGACAATGACTGTCA |
| Slc1a3 | AGGCTTCTACCAGATTGGGA |
| Slc22a5 | TTTATGATCTGATCCGAACA |
| Slc22a5 | GGGTCAGATCTCCAACTACG |
| Slc22a5 | CACAAGGCAACGGTGCTCCG |
| Slc22a5 | CACACCCACGAAAAACAAGG |
| Slc22a12 | GGGCCTGGGAGTTACATACC |
| Slc22a12 | ACGGTAGGCAAGCTGGACCA |
| Slc22a12 | GAGGTGCTATTGTCCAGGAG |
| Slc22a12 | CATCACCAAAGGGCTACCCT |
| Slc2a1 | CCTGCTCATCAATCGTAACG |
| Slc2a1 | TCAGCATGGAGTTCCGCCTG |
| Slc2a1 | GTGTCACCTACAGCTCTACG |
| Slc2a1 | CAAACATGGAACCACCGCTA |
| Slc2a2 | AGAGGGCTCCAGTCAATGAG |
| Slc2a2 | TTACCGACAGCCCATCCTCG |
| Slc2a2 | GGACTGGTTCCAATGTACAT |
| Slc2a2 | TGTGATCAATGCACCTCAAG |
| Slc3a1 | CCATATACCAGATCTACCCG |
| Slc3a1 | ATCCTTGGTTCCAATCGAGT |
| Slc3a1 | AGAGGAGCCTCACCTAAAGG |
| Slc3a1 | TGGCAAGCCATAGTACATCA |
| Slc5a1 | AGGAAGAATGCTACACACCG |
| Slc5a1 | GAGACATGTTCTTGGCCGAG |
| Slc5a1 | CATCGCCTACCCCACGCTCG |
| Slc5a1 | CCGGCCACCACACCATACTT |
| Slc7a5 | GCCCTCCTCGCAGTACATCG |
| Slc7a5 | ACCCCTACTTACGCACGCAG |
| Slc7a5 | AGCGGCCTCTTCGCCTACGG |
| Slc7a5 | GTAGCAGAGTGCGCCCACGA |
| Slc7a7 | AAGAGATCAGGAACCCCGAG |
| Slc7a7 | CAGCGCCAACACCTTAGCAT |
| Slc7a7 | GGCCCCGGATTTCTTAATGG |
| Slc7a7 | GAGACACACGCCATTAAGCA |
| Smpd1 | AGGCTTTCTAGCGTTCGCAA |
| Smpd1 | GAAGAGGACACGGCTGACAG |
| Smpd1 | AGTCTCGCCAAGATCAGCTG |
| Smpd1 | GCTGACTGGCACACATCTAG |
| Sms | ACTTACTAACATCCCCGCTG |
| Sms | TTACCACCCATAGTTCGCGG |
| Sms | CTTACACGAACAAGAATGGC |
| Sms | AAACAAGAAACTGACAGCGT |
| Spr | GCGTGCGCTTACGAGCATCA |
| Spr | GCGCAGTACGCGAGCTCCCG |
| Spr | AGCCTCGGTGCCCAGATCGG |
| Spr | TACAGACCCCAGCCTTTGTA |
| Sptlc2 | GTTGTGTTTGAAGATTCGAA |
| Sptlc2 | TGAGAGCAATCACTTCAGGA |
| Sptlc2 | AATCTCGAAGATATCCAAAG |
| Sptlc2 | ACAACTATCTTGGATTTGCG |
| Scarb1 | GGGGCCGTGAAGCGATACGT |
| Scarb1 | TGCGGTTCATAAAAGCACGC |
| Scarb1 | GATGAACAACTCGAATTCTG |
| Scarb1 | GAGGATTCGGGTGTCATGAA |
| Ssr4 | TAGTAAGAAGGGGTGATCTG |
| Ssr4 | AAACAATTTCCTGTAACCCG |
| Ssr4 | TACAGAGACTGTATTCATCG |
| Ssr4 | CACCTTCCTTAGGAGGCTAT |
| Star | AGAACTTGTGGACCGCATGG |
| Star | CGAACTTGACCCATCCACCC |
| Star | GAAGCTCCTATAGACATATG |
| Star | CGCACGCTCACGAAGTCTCG |
| Stra6 | TCTTCAAGCACTACACCGAG |
| Stra6 | CAAGTTGTACTGGATACCAA |
| Stra6 | CCAGACCTGAACACCAAAGT |
| Stra6 | GAAGCATCACCTATGGACTG |
| Sucla2 | CAGAGCGTAACATACTGTCA |
| Sucla2 | TGTGCACTTCCTATCAGTAC |
| Sucla2 | GTAGAAGATTCTGACGGAAA |
| Sucla2 | TCAAGTATTCATGCAGCGAA |
| Abcc8 | ATGGCTGCTAAATGCCACGG |
| Abcc8 | CAGACCAACGAGATGCTCCG |
| Abcc8 | TGGACAATGAAGATCATCGG |
| Abcc8 | GAAACTGGGATTAACCTGAG |
| Surf1 | TGTTTCTATAGGTCCAACGT |
| Surf1 | ATGGAAAGGAGTAACTACAT |
| Surf1 | GTGGTCGCAATGGGCCTACG |
| Surf1 | TTATGTACAACTCTTTAGAG |
| Taldo1 | GCGCATCCTTGATTGGCATG |
| Taldo1 | CTTCTTTGTAAAGCTCGATG |
| Taldo1 | TTCTGAATTCAGGCCTCAAG |
| Taldo1 | GCTTGTATTCATCGATGGCT |
| Tbxas1 | TCACAGGCTTGGCTGATGAG |
| Tbxas1 | CCACACTTACCATTTCAGGA |
| Tbxas1 | ACAGAGGCCCGTATCGCTCT |
| Tbxas1 | CTGCTGTTACACCATAGATG |
| Hnf1a | CCTATAACGGACCTCCACCG |
| Hnf1a | GACGTACCAGGTGTACAGAG |
| Hnf1a | TGCCAACTAAGAAGGGGCGT |
| Hnf1a | CGGACAGTCTGCAACCAGTG |
| Hnf1b | CAGCTTCACCCCGAAATTCG |
| Hnf1b | CTGGTTCGCAAACCGCCGGA |
| Hnf1b | GCCGCAACCGGTTTAAATGG |
| Hnf1b | CTTGGTACGTCAGAAAGCAA |
| Tcn2 | GAAGCGGCTCCATGACAGCG |
| Tcn2 | TCTGAGACCACGAATCACCA |
| Tcn2 | GAGACTAGCAATACCGCAGG |
| Tcn2 | GAATATCTATAGCACCCCAC |
| Tfam | TAAATGTTATATGCTGAACG |
| Tfam | GGAGCGTGCTAAAAGCACTG |
| Tfam | CTAACTCCAAGTCAGCTGAT |
| Tfam | GCTGTTCTGTGGAAAATCGA |
| Th | GGGTGAGCCAATTCCCCACG |
| Th | ACTGTGTGCACTGAAACACA |
| Th | CCCCAAGGTTCATTGGACGG |
| Th | GTGCGCTTCGAGGTGCCCAG |
| Tkt | CGTGGACGGACACAGCGTGG |
| Tkt | CAGCGCTGCAGCATGATGTG |
| Tkt | CCCACAGATAGCCACCCGGA |
| Tkt | CTCCGAGGGCTCCGTCTGGG |
| Tmem165 | GGCATTAGAATGCTTCGGGA |
| Tmem165 | ACATAGTATGTATACACCCT |
| Tmem165 | GCAGCTGCCCAGACGAATCT |
| Tmem165 | GTCATCATAGTGTCCGAACT |
| Tpi1 | TGAAGGTCAGTACAAACGCA |
| Tpi1 | AAGTCGATGTAAGCGGTGGG |
| Tpi1 | AGTGAGCCACGCCCTAGCAG |
| Tpi1 | CCAACGAAGAACTTCCTGGT |
| Tpmt | AAAGAACCAAGTACTAACCC |
| Tpmt | TTCTGCAGGTTCGCAGATCG |
| Tpmt | GCATTAGTGGCTATCAATCC |
| Tpmt | AAAACACTCGCAGTCCACTC |
| Trex1 | TTTCCTCGAACCATTCCCTG |
| Trex1 | ACACAGAAGGTACCATCTAG |
| Trex1 | AGCTTGTCCACCACACGGGG |
| Trex1 | GGAGCAGAGGAAAGTCATAG |
| Tfrc | CTACACGCTTACAATAGCCC |
| Tfrc | GAATACATACACTCCTCGTG |
| Tfrc | GGGCTCCTACTACAACATAA |
| Tfrc | AACCCTCGGGAGACTCCACT |
| Cmpk2 | TCGGCCGCGGCGCTTCACTG |
| Cmpk2 | GGTGTTCCAAGACCGGGACG |
| Cmpk2 | AGTCGCGCCGAGTGTCCGGG |
| Cmpk2 | AGTTGACCAGTGCCCAAAGG |
| Tyr | AGAAATTCGAGAACTAACTG |
| Tyr | TTTATGCGATGGAACACCTG |
| Tyr | ATGTTGATATCATTAAACAT |
| Tyr | ACCCCTTTGAAGGGGAACTG |
| Ucp2 | TCTGGGTACCATCCTAACCA |
| Ucp2 | GTCAAACAGTTCTACACCAA |
| Ucp2 | AGACCATTGCACGAGAGGAA |
| Ucp2 | CGGACCTTGGCGGTATCCAG |
| Slc35a2 | TCACCCGCTGTAGTGGACCC |
| Slc35a2 | CTGCTCTTCGCACAAAAGAG |
| Slc35a2 | CTGCAAGGTATAGATGAGAG |
| Slc35a2 | GGCCACTGGATCAGAACCCG |
| Ugcg | CGGCTACATACGGCAGCCCG |
| Ugcg | TGGCCAAAGCAATAGCCGAC |
| Ugcg | AAGGATGTGCTAGATCAGGC |
| Ugcg | CAGCCGTATAGCAAGCTCCC |
| Umps | GAGCAGATAACTGTCGCCAG |
| Umps | CCGCAGGTCGATGTAGACTG |
| Umps | AGAGCGTGCACACGGCGTGG |
| Umps | TCTGTCTGCCGATGTGTCGG |
| Ung | ACGGACCTAATCAAGCTCAC |
| Ung | TTGTCAGGGTGGGCCCGACA |
| Ung | CCAACCCCGACTCTGACTCC |
| Ung | CCACAAGGTCTATCCGCCCC |
| Uqcrq | GATCCCTACAGCGTTTGTAG |
| Uqcrq | CTCAAAGGGCGACAAGCTGT |
| Uqcrq | GCAGGATGCGCTCGCGAGTG |
| Uqcrq | TTTGCTGAAATAGCTTGGGA |
| Urod | CCATCACCCTTACTCGACAA |
| Urod | GTGGACCCTAATGACATACA |
| Urod | ACTGCCCTACATTCGTGATG |
| Urod | CAGCACCTGCCGATCTCCCG |
| Uros | TCTCCTTTGATAGTTCCACA |
| Uros | TGACAGCACAGGAATCAGTG |
| Uros | GCCAAGTCTGTGTACGTGGT |
| Uros | AGACATGCATGCTTTCCATG |
| Usp9x | GCAGATATGGAAACTCGAAG |
| Usp9x | ATGGGTATAGTATGACACAC |
| Usp9x | GAAGTTGATTGATTAAGTCA |
| Usp9x | TATCCAAACACATCATCCCT |
| Vdr | TGGAGATTGCCGCATCACCA |
| Vdr | AGCGTTGAAGTGGAAGCCCG |
| Vdr | TTCTTCATTCAGATCCATCG |
| Vdr | TTCGTGCAGACGTAAGTACA |
| Xdh | CCAGCATGCAACGTACAGGG |
| Xdh | CATACTCATGACAATACCAG |
| Xdh | TCAAAACGCAACGTCTTCCG |
| Xdh | GGAATTCCACCATCCCACCA |
| Nt5e | TCATGAATTTGATAACGGTG |
| Nt5e | TGAATAAGATCATCGCCCTG |
| Nt5e | TATGCCTTTGGCAAATACCT |
| Nt5e | CCTGAAGCGGCACGTCTGAG |
| Papss2 | GAATATCCGCCGGATCGCGG |
| Papss2 | AGTCGATCAAATCCGAGCTG |
| Papss2 | CTTGATGTACGAAGGTCGGA |
| Papss2 | GGTGCTAGAGAGAATAAGGT |
| Slco2a1 | CCCACGGATGATCGGCATAG |
| Slco2a1 | ACTCGGGGGATGGTTTGCAG |
| Slco2a1 | CTTCGTGGACTACGGCAGAG |
| Slco2a1 | CTCTGCAAAGTCGTCCACAT |
| Slc35a1 | GAACACTCAGCAAATTACAG |
| Slc35a1 | ATAGCACCAAAGCCTAACAA |
| Slc35a1 | TGCACAGCATACACTAGTGA |
| Slc35a1 | TCTTAAAGCTACGGTGTAAG |
| Mpdu1 | TTCCTGGTCATGCACTACAG |
| Mpdu1 | CTAGCTCCAGCATTACTGAC |
| Mpdu1 | ACTACAGCCAGAGGCGTGAG |
| Mpdu1 | ACGACCAGCTCTTCGTGCAA |
| Rbck1 | TGCTTCATACCAGCCTGACG |
| Rbck1 | AGTACGCCCGGATATGACAG |
| Rbck1 | TGCATTCACACGGCATTCGG |
| Rbck1 | ACCCGAGGTCTCCCCAACAC |
| Btd | CTTGCTGTTCATAGACGTCG |
| Btd | GTACCAAAAGGACTTGAGAG |
| Btd | ATACCAGTTTAACACAAATG |
| Btd | CCGCGAGGCTGAGTACTACG |
| Slc27a5 | GTGGGCTTAATGAACTATGT |
| Slc27a5 | TACCTCTGTACCATACGATA |
| Slc27a5 | GTAACAGTGATCTTGTATGT |
| Slc27a5 | CCTTTGTGGATGCTTTAGAG |
| Txnrd2 | TATCCAGTCCAATTCCAGTG |
| Txnrd2 | TGAGCACACAGTTCGCGGTG |
| Txnrd2 | GGTGGCACCTGTGTCAACGT |
| Txnrd2 | CGGTGGCCTAGCTTGTGCCA |
| Slc27a4 | CAAACGGATAGGGTACACAA |
| Slc27a4 | TCTACACATCGGGCACCACG |
| Slc27a4 | TGACTTCAGGAAACATCGTG |
| Slc27a4 | ACAGACCCACAAACTCATTG |
| Cln8 | GCTCATCTCTAGGAGCAACG |
| Cln8 | GTGTTGGTTTCACATCACCA |
| Cln8 | AAAGGTGCGGAAGAACAGGT |
| Cln8 | CTTTGTCGGCATAGAGCACG |
| Mecr | CGTGGCGGTACCAAGCCTCG |
| Mecr | AAGGATCTGACGTCCACGTG |
| Mecr | ATCCAGAATGCATCCAACAG |
| Mecr | AGCACTGATTGGAATCCCTA |
| Aifm1 | CTGCCTAATATTGAGAACGG |
| Aifm1 | ACCATGGAAAAAGTCAAACG |
| Aifm1 | GTCAATTACAGTTATCGGCG |
| Aifm1 | AAGTCTGTCTGCCATCGATA |
| Sgsh | CTGACCTAGAAGGCTCACGT |
| Sgsh | CGTTACGGAAGATAAGGCTG |
| Sgsh | TGTACCGGGCCAGTACAGGT |
| Sgsh | TCCTGAGGGTCGTAGATCTG |
| Asns | GCATGCCATCTATGACAGCG |
| Asns | GAATGCAGCCGATAAGAGTG |
| Asns | GCTGTGTGTTCAGAAGCTAA |
| Asns | ATTTGAATATCAGACCAATG |
| Sec23b | GCAACACCAGTGGACCGCAG |
| Sec23b | AATGGCGTTTGGTGCTACGT |
| Sec23b | AAAAGTGATGAGACCAACCA |
| Sec23b | GTGTTGGTGGCACAAGTCAG |
| Dguok | AGAAAGGGTCTGGAATGTGT |
| Dguok | TTTCATGAGTAACTTCACAA |
| Dguok | CGTGGACGCGCCACACGCCA |
| Dguok | TCCTGCAGGAGTTCGCAAAC |
| Slc25a10 | CATGCGGGACTACATGACCA |
| Slc25a10 | TACACGGTACAGACCATCCA |
| Slc25a10 | GTCAGAGAGTAGGTCATCTG |
| Slc25a10 | GACATTGACCAAATCTGCTG |
| Yme1l1 | CAGGACGTATTAAGGCACTA |
| Yme1l1 | TTAACTATTACCTCACTACT |
| Yme1l1 | TCATCGATGAATATAACACA |
| Yme1l1 | CTTGGAGGTAAACTTCCCAA |
| Pign | CCAGCTCCCCAAGTAACGAA |
| Pign | GTGTTGTTAAGATAACCCAC |
| Pign | TGCCACAAAACTGGATACGT |
| Pign | AAGTCAATAGTGATTCAACC |
| Pdhx | CTTTAGTGAAGATCCCGCGA |
| Pdhx | GTTAGACGAGATCTGGTCAA |
| Pdhx | TGTCTCCTACGATGGAGCAA |
| Pdhx | TCCTGGGCAACCGAATGCAG |
| Abcg5 | GCTGATAGAGCAGCCCCACG |
| Abcg5 | AGCTTGCCGAACATCCCAGG |
| Abcg5 | CAATCATTTGGTCCGCCACG |
| Abcg5 | GGATTGGAATGTTCAGGACA |
| Abcb11 | TACGCACCATGCCTTCGCAG |
| Abcb11 | ATGAAGGCGAGTACACACCA |
| Abcb11 | CCACTGTTCGAATAGATGAG |
| Abcb11 | AAGGTTGTGGGTAATCACTG |
| Naglu | TGCACACATTCTGGTAATAG |
| Naglu | GAGATCGATACGTACTTCAC |
| Naglu | AGGAAGAGGTTCCCGATGAG |
| Naglu | GCCCTCGAGGATGTGAACCG |
| Abcc6 | GTGACCGCAACTCCTCGGCG |
| Abcc6 | CCTCTGTGGAGGATCCACCA |
| Abcc6 | AACAGGGTGTAGGTACAGAG |
| Abcc6 | TGCACTGCATCGTTCAGGTG |
| Tango2 | GTTTGCTACTATGGAAACCG |
| Tango2 | GGCTGGGCATCAGCACACGT |
| Tango2 | CCTGAAGAAGGTCTCTACAG |
| Tango2 | TAGATGCCTTACCTCTACCA |
| Vkorc1 | GTACGCACTGCACGTGAAGG |
| Vkorc1 | GTGAGAGGGCTAAGCCAGCG |
| Vkorc1 | GGTGGAGCACATGCTAGGAG |
| Vkorc1 | CCATGTCTGCACGCACCGAG |
| Gfm1 | GAACGGGTGCTATACTACAC |
| Gfm1 | GAAGCACCATCTACAACACG |
| Gfm1 | AGCAATGAGGCCTTCTAACA |
| Gfm1 | TTACTCTGTAGCGAGCAATC |
| Tpk1 | CTTGCCTGAATTCGTCAGTG |
| Tpk1 | GAACTTGAAATACTGCCTTG |
| Tpk1 | GACACTTGGTAAAGTCAGTG |
| Tpk1 | CACTTATATGATCTCACTGA |
| Dnajc12 | GAGCCCGCTATGACCATTGG |
| Dnajc12 | GAGCTCATCGCATCCCAGCA |
| Dnajc12 | TGTCAATGCCGTTCGAGCAG |
| Dnajc12 | AATTTCTGAAAAGTCTCCAC |
| Angptl3 | TGCTCTGCCGTTTATAACAG |
| Angptl3 | TACACTACAAGTTAAAAACG |
| Angptl3 | CATGGACATTAATTCAACAC |
| Angptl3 | GAAGACAGCCCTTCAACACA |
| Aass | GAAACTTCTCTTAATTCGTG |
| Aass | GCAAATTATTCACGACAGGT |
| Aass | TAGGCAGTGATTACGTCCAA |
| Aass | GGATAGTGGCTTTCGGACAG |
| Slc7a9 | GAGCACTTACCAACCATCGT |
| Slc7a9 | CAAAGGCCTCCATCAGATAG |
| Slc7a9 | GGACTGCAAGAGCTCCGTTG |
| Slc7a9 | GCTGGCCAACACAGAATCCG |
| Hacd1 | GCAACTCACCGATCAGACAA |
| Hacd1 | CTGGCCCATTTAATGAAGTG |
| Hacd1 | GATCCAGAATGAAGAGAGCG |
| Hacd1 | CTTCTACAATATCGCCATGA |
| Ttpa | GAGGTGGAAACTCAACGCAA |
| Ttpa | CAGATCCAGATCGAAATCCC |
| Ttpa | GAAGTCCAAGGATACTTCTA |
| Ttpa | TTTATTTGTTGTAGCATACT |
| Mrps7 | GGAATATTACCGCAAGCCGG |
| Mrps7 | GGCCGCGACTGAAACCAGCT |
| Mrps7 | CCCCTACAGGATCTTCCACG |
| Mrps7 | GAGGCAACAAAGTTCTGGCC |
| Tfr2 | GAGGTCGCTCCAGTACAACG |
| Tfr2 | AGTGCGTGTCAGTCCACACG |
| Tfr2 | TGGTGTACGCCCACTACGGG |
| Tfr2 | ACCCCCAGTGAAGATTAGCA |
| Pnpla6 | CTCCGTAGTGTCATCCAACG |
| Pnpla6 | GCCATGTCAAAATCAGAACG |
| Pnpla6 | AGGAGACTCCGACCCTACAA |
| Pnpla6 | GGCCTGGCGCAGACAACGGA |
| Atp8a2 | GCAAGCCCTCTTCATAAACT |
| Atp8a2 | GGCCCTTATCCTATTGAAGG |
| Atp8a2 | ACAACTTCGACTCTACTGCA |
| Atp8a2 | CAGTGTTAAGAAATGGCATG |
| Polg2 | GAAAGAACCTAGCCTCACAG |
| Polg2 | TGTGGGAGTAAACCATACCA |
| Polg2 | TGACGCCCCCGAGCATGCGG |
| Polg2 | GCAATTAACATAGTGCTCCA |
| Hs6st1 | GGCGCACGTTCTGCACTAGG |
| Hs6st1 | ATGGCGACATGTACAGCGTG |
| Hs6st1 | CTCATCCTTTACCAGTACGC |
| Hs6st1 | CCTATAACCTGGCTAACAAC |
| Acsl4 | GTCCAGGGATACGTTCACAC |
| Acsl4 | GCCCATATCCCTGACCAATG |
| Acsl4 | CAATAGAGCAGAGTACCCTG |
| Acsl4 | GGAACAGCGGCCATAAGTGT |
| Gne | ACGTCCAACTCAAAGAACGC |
| Gne | TTGCAGCTCAAAGATATATG |
| Gne | GCTCCACACGATTGTTAGAG |
| Gne | CCTCTTGTTAAACGAGATCA |
| Slc25a13 | GGGGCGACTCCCAGTAACTG |
| Slc25a13 | CAGATTTATATGAGCCGAGG |
| Slc25a13 | ACAAGGCATCCGGAGCACAC |
| Slc25a13 | TACAAGATCGATAGGATACA |
| Galns | AGCAGTAGCACGATGTTGGG |
| Galns | CCCAATTAACCGGAAGACTG |
| Galns | GGCGTAAACTGGTGCATGAG |
| Galns | GAGAGACCCCAAATTTAGAC |
| Nus1 | GTGGTCGTAGACGCTAATGT |
| Nus1 | TCCAGGTGCCGAAGCGAACG |
| Nus1 | ACAGCACCTTCACTGCCGAA |
| Nus1 | CCAGCGCAGCCGAGGATGGG |
| Coq5 | CTTCGGGTTTGAGACCGTGT |
| Coq5 | GTTGTGCCTGAACGTAACTA |
| Coq5 | ATGAATGACATGATGAGTCT |
| Coq5 | GCCAGGATCTATGACACCCG |
| Hgsnat | CCTGCAGGTTAACTCCACCT |
| Hgsnat | ATGACTTCTATCCTGCAACG |
| Hgsnat | GATGTGTGGACACATTTAGG |
| Hgsnat | GTTGGGAGTGACATACTTCG |
| Slc46a1 | AGAGCTAACATCTGCCACAG |
| Slc46a1 | GGGCAATGGATCGATGATGG |
| Slc46a1 | TGGACCAGAAGAGTCCCACC |
| Slc46a1 | GAACTGTGGGAACCAAAGCG |
| Coa3 | GATACAAATAGCTAACACCA |
| Coa3 | CTCAGCGCATCGACCCGTCG |
| Coa3 | GGGTCGATGCGCTGAGCGAA |
| Coa3 | CATAAACTGCAACTGCGCTG |
| Trmt10c | CTTACGGTATCTGTATGGGA |
| Trmt10c | TTATTATGACAAGACAAAGA |
| Trmt10c | TTAGTTCTCTGGTTGAGGCG |
| Trmt10c | TAGGCCATGTCAAAAACCAA |
| Slc52a2 | TGTTGTGGGTTCAGATGTCG |
| Slc52a2 | ATGGGAGACACCTCGATCGG |
| Slc52a2 | GGCCTCTCTGTGGAACCACG |
| Slc52a2 | AAGACCGTAAAAAGGGGGGT |
| Sco1 | AAATCAATACCCATTGACCC |
| Sco1 | AAAAATGATTGAAGTCGTGG |
| Sco1 | TTGGGAAGCCTTTACTAGGG |
| Sco1 | CACTCACCCCAGGCTTCGGG |
| Pla2g6 | AAATCCATGGCCTATATGCG |
| Pla2g6 | GGCACCTGCATTAGCCCCGT |
| Pla2g6 | AGTCTCCCCAAAGTCGTTTG |
| Pla2g6 | CCATTGGGCCAAGAACGCCG |
| Chst3 | CCACGAACGAGGAACCCGTG |
| Chst3 | AACCTCGGGTCCTCAACTAG |
| Chst3 | ACTCAGTTCCTGTTCCGCCG |
| Chst3 | AAACTATGACCACAAATGCC |
| Fut8 | TGAAACAGTAGACCACGTGA |
| Fut8 | AAATGACAAAAACATTCAAG |
| Fut8 | CCAGAAGGCCCCATTGACCA |
| Fut8 | AATCAAGTATTTGACAAACT |
| Gria3 | TGTGACGAAAGATGTATGCA |
| Gria3 | GTACAATGTCAGTAAAACCC |
| Gria3 | TACCTCTATGACACAGAACG |
| Gria3 | AGGAATCCAAGTGGTCTACG |
| Porcn | ACAACTTCCACCATGGACCG |
| Porcn | GTGACATGGCACAAGATGCG |
| Porcn | CCACCTTCTTCAGCCATCGG |
| Porcn | GAAGGAGACAGCACTCTCGT |
| Clpp | GCGCTTATGACATATACTCG |
| Clpp | CCTGCAGATTGACGACAGTG |
| Clpp | CATATGTATATCAACAGCCC |
| Clpp | CAACACACCACGTGCAGATG |
| Slc40a1 | CAGGGTACGCCTACACTCAG |
| Slc40a1 | CCTTTGGATTGTGATCGCAG |
| Slc40a1 | TCATCAGGATGATTCCGCAG |
| Slc40a1 | CCCATCCATCTCGGAAAGTG |
| Pmm2 | ATTCAATGAAAGTTCCCCTG |
| Pmm2 | AAGACCAAAATTGGAGTGGT |
| Pmm2 | GCTTGGTAGCGTACAAAGAT |
| Pmm2 | AAAAGTTCGTAGCAGACCTG |
| Sult2b1 | GGGAGACCACGACATCGCGG |
| Sult2b1 | GGGCTCCGATCGGATCCACG |
| Sult2b1 | AATGTTTCCGAAATGAGGTG |
| Sult2b1 | AGGTGAGAGCTCATAATGCG |
| Cd320 | TGGTCTCAGAACAGGCCTAG |
| Cd320 | GCAACCACTGATGTTGTCAC |
| Cd320 | CCATCACAGCGCCACGTGTG |
| Cd320 | ACCTTCCAGTGTCTTACCAG |
| Elovl1 | GGCTATTGGAAAAGTCTATG |
| Elovl1 | GGGCAGACAATCCATAGTAG |
| Elovl1 | TCCAAAGCTACCCTCTGATG |
| Elovl1 | CGATAGGATGAAGTACACAT |
| Ndufa1 | CATCCACAAATTCACCAACG |
| Ndufa1 | TGTACGCAGTGGACACCCCG |
| Ndufa1 | GAATTTGTGGATGTACGCAG |
| Ndufa1 | CGCGTTCCATCAGATACCAC |
| Atp6ap1 | GAGGATTTCACAGCATACGG |
| Atp6ap1 | GATATGACCCTCATGTGTGT |
| Atp6ap1 | TAGCTAGATCCACATGCAAG |
| Atp6ap1 | GTGTCATTGTAACTCACAGG |
| Extl3 | GCCCAAGCCTCGCGTCACAG |
| Extl3 | TCAGACATAGCATGGACAAG |
| Extl3 | CCACACAGTGCCCACTCAGT |
| Extl3 | ATTGCGGAGGTATTTAGGTG |
| Wdr45 | GACACTCGGGACAACCCCAA |
| Wdr45 | AGGCGTCCGGATCTACAATG |
| Wdr45 | AAGCTGGTAGAGCTTCGAAG |
| Wdr45 | GTGTGGAAGTCTGCAACTTG |
| Atp8b1 | AATGCCACACCGTCCTACCG |
| Atp8b1 | AGCGTTGGATAAAGTGTACG |
| Atp8b1 | GCTGATAAAATCCTGTTACG |
| Atp8b1 | GATTTGCCATACGGTCATGG |
| Mpc1 | ACTTCCGGGACTATCTCATG |
| Mpc1 | GGCGGACTATGTCCGGAGCA |
| Mpc1 | AAATCTCCAGAGATTATCAG |
| Mpc1 | TGGGGCCCAGTTGCCAACTG |
| Slc1a4 | GTCTGCAACCGATTACACAG |
| Slc1a4 | TAGAGCCACTCCTAACACCA |
| Slc1a4 | GATGCCACCCAGACGCCCGA |
| Slc1a4 | ACCCACCAACACTCCCGACA |
| Pgam2 | GGATGTTACGGACCAAATGT |
| Pgam2 | AACCAAGAGAACCGTTTCTG |
| Pgam2 | GCTTCAAGCCTGCATAGCGG |
| Pgam2 | GTTTGACATCTGCTACACGT |
| Samhd1 | ATCCTTACATTATGTCGATG |
| Samhd1 | GCTTGATATAGCGAAGTCGC |
| Samhd1 | CTTGGGCTGCCATCGCAGCG |
| Samhd1 | TTAGGATCTTACCTAGGTCG |
| Pdss1 | AGCCTTTAGACCGATTATTG |
| Pdss1 | GAGGCGGCACGTGTTCCGAG |
| Pdss1 | TATACTATGCATAGTTGGGG |
| Pdss1 | GTTATTGAAGATTTGGTGCG |
| Dgke | TCTCGTAGTGGAACTAACAT |
| Dgke | GATCATGCTCAAGAACGACA |
| Dgke | GTCCACGACGAGTGCATGCG |
| Dgke | ACAAGAAAAATACATTCCAG |
| Pigp | TCTCTTGTGCAGATACTGGG |
| Pigp | TGCAGTACTTTATCTTGTGT |
| Pigp | AAGGAGTTTAACCAAGATTC |
| Pigp | ATGAATGGAGTCAAGTGGAG |
| Prodh2 | TAGTCCATGAGTGACCAAAG |
| Prodh2 | CCTGTGCCACTCGGTGCAAG |
| Prodh2 | GGACTTATCCCGAGCCCTCG |
| Prodh2 | GTTGGCAGTACCCACCGAGG |
| Pex14 | CTTCACTGGGATCTGCCAGG |
| Pex14 | ATGCTCTCCTGGTCGCAAGT |
| Pex14 | GGAACAGGTGACTTACTGTA |
| Pex14 | ATACTTACCAGTGGCTCTCG |
| Mrpl12 | TCTTTGGGAGCATTATCCAG |
| Mrpl12 | ATACCTTCAGGAGTTCGTTG |
| Mrpl12 | TGCCCAGCGTCTGTGCAGCG |
| Mrpl12 | GCCCCCCGGAGTCCAAGCCG |
| Ggcx | TGTGTGTATAAGAGGTCCCG |
| Ggcx | GCCTGCACGATGTCCACACG |
| Ggcx | AGAACTGTGTAGTTCCAAAG |
| Ggcx | GCTCGCCCGTGAGGCCGTCG |
| Timm22 | GCTGGCATTGATACCAACGT |
| Timm22 | AGAAGAAGGCTGTACTGCAA |
| Timm22 | TGGGCGACAAGCGTCAGCCC |
| Timm22 | CCTCTAGGGTTTGTCTTGGG |
| Ivd | AATATCGAGTGGGCCCGCCG |
| Ivd | CTCCTGGCTCAGGACGTTAG |
| Ivd | GGTAATGGAAGAGATATCCC |
| Ivd | TTCTTGGAGGTACTAAAACC |
| Pus1 | CCTGCAAAGAGGGTCAAGGG |
| Pus1 | CTTACCCAGAATCCGAATGT |
| Pus1 | TACTCGGGCAAGGGCTACCA |
| Pus1 | TCCAGGCCCTCACGTACGAA |
| Adar | ACTCCAACAAGCCGCCTACG |
| Adar | AGAGGTAACCCCAGTAACAG |
| Adar | TTCTTGTAGGGTGAACACCG |
| Adar | TGTATCCAGGAATTCCCTAG |
| Suclg1 | GAGTACGGCACCAAACTCGT |
| Suclg1 | AACAGAATGGGATACGACAC |
| Suclg1 | TGACACGCCAGGGAACGACG |
| Suclg1 | CATTAATGAAGCAATCGACG |
| Aldh18a1 | CAAGTCTAGAGTGGGCCTAG |
| Aldh18a1 | CGATGGGGACGATGTTCATG |
| Aldh18a1 | CGTCCGAGAGGACAATCAAG |
| Aldh18a1 | TGAGGGGTACCGTGATAAAG |
| Ctsf | ACAATTACGGCCGTGCTGCG |
| Ctsf | AAAGACAGTCAATCGCCACT |
| Ctsf | ACCAGGGGATCATTGCAAGG |
| Ctsf | CCTGAATCCCCTCTTACAGA |
| Tbk1 | TGCCGTTTAGACCCTTCGAG |
| Tbk1 | CTTCTCGCTACAACACATGA |
| Tbk1 | CAACATCATGCGCGTCATAG |
| Tbk1 | CGGGAACAACTCAATACCGT |
| Pex3 | AAACAAGCTGGAAATATGGG |
| Pex3 | TCATTAAACAAGCTGTACAA |
| Pex3 | CATGCTGCCGACACTGAGAG |
| Pex3 | TACTGCTGCTGTACATCCGG |
| Txn2 | AGACCACCAGCATTGTACTG |
| Txn2 | AGGGAAGCCCACACACCCTG |
| Txn2 | GCGGTCCTAGGATCTTGCAG |
| Txn2 | AAAGGTCGTCAAACAGACTC |
| Mlycd | GACTTCGTGAGCTTCTACGG |
| Mlycd | TGGAAGAGGCCGCGATACCG |
| Mlycd | CTCCGACTGAAACCGAGGAG |
| Mlycd | CCGCACAGCCGACGTCCCCG |
| Tdo2 | GATAGCTCGGATGCATCGTG |
| Tdo2 | TGATGAATAGGTGCTCGTCA |
| Tdo2 | AATCCATTTGGCTCTAAACC |
| Tdo2 | CACTATCGTGATAACTTTGG |
| Alg2 | CAGAAAGACCATGCGCACGT |
| Alg2 | GCTCTCGGTGCAATGCGCAG |
| Alg2 | TCTCTATCAACCGATACGAA |
| Alg2 | ACTGGCCAGACGGCGTAAGA |
| Mocs1 | CACTTGGATTCCGAACACGG |
| Mocs1 | TAGGTCTCGAGGGCTCCCCG |
| Mocs1 | CGATGTCCACCACATCCGGT |
| Mocs1 | AGGCCAGTATTGCATGCCCG |
| Nfu1 | ACTGTACCTGTGTTACACAA |
| Nfu1 | GAAACAGTACCTGGCCAGAG |
| Nfu1 | GTGTAGTCCGGCTGTGCAGG |
| Nfu1 | GCGCTACCTGCTTCTCCCGG |
| Dhodh | TTGATCCAGAGTCGGCGCAC |
| Dhodh | GTAGAAATGGTCGTCCCCCG |
| Dhodh | ATAAATTCCGAAATCCAGTA |
| Dhodh | GGTATGGATTCAACAGCCAC |
| Stap1 | GTTGGCCAGTAAGGCACACG |
| Stap1 | GGAGCCAGTACAAGACTATG |
| Stap1 | CTTCATTCTTACAGTAACAG |
| Stap1 | GGAGTACAAACACTATTGGA |
| Aldh1a3 | CACCAGGCATGAGCCCATCG |
| Aldh1a3 | TGCATCCAGCCGGCGCCACG |
| Aldh1a3 | CCACCCGGCAAAATATCTGA |
| Aldh1a3 | AACACTAGAGAAAATATGTG |
| Slc25a20 | ATAGGGGTGACTCCAATGAT |
| Slc25a20 | TCCAAGGTCCCAGAGTACAT |
| Slc25a20 | TCAGGGGAGAACAAGTACAG |
| Slc25a20 | TCCCAGCTGTAAACAGCTGT |
| Srd5a3 | ACTGCTTGGTCTTCCCGTAG |
| Srd5a3 | AGCACTCAAAGAGTCTCCGA |
| Srd5a3 | ATCTACTGATACAAGCCCGG |
| Srd5a3 | TCTACGTCATCTCAGTTGTG |
| Mogs | TCTAGGTCATTCTTCCCACG |
| Mogs | TCGGCAGCATATCCACGATG |
| Mogs | TGCCGAAATAGACGTGTGGG |
| Mogs | GAGGTCCTACTACCAGAGAT |
| Tk2 | CTCCAATACAACAGACGTCG |
| Tk2 | TTGAGGGCAATATTGCAAGT |
| Tk2 | AGAATCGCGTAGTCAACCTC |
| Tk2 | TACCATGATGCCAGCCGATG |
| Cldn10 | GCCCGAGATGGAGACTACGA |
| Cldn10 | CAGACCATCCAACGCCAGCA |
| Cldn10 | TAGAGGACTAATGATCGCTG |
| Cldn10 | ATCTGCGTTACCGATTCCAC |
| Chst11 | AGCCAACGAAGCCCACGTGT |
| Chst11 | GCAGACACCAGCCTCTCGAA |
| Chst11 | AGCAGATGTCCACACCGAAG |
| Chst11 | ACCTCCTTCCACAAGCGCTA |
| Treh | AGATTAGGGAGGACCTCTCG |
| Treh | AGCTACGGACATATCCCCAA |
| Treh | AGGGGACCGAGAGACTCTGT |
| Treh | AGGAACGCGTTCAGATCAGC |
| Hibadh | AGCTCCACAGTATACCACGT |
| Hibadh | GCTGCCCTCCAGTATGAATG |
| Hibadh | CGTGTTCCCTGATGTATGCA |
| Hibadh | GGAAACCTTACATTTATGGT |
| Sumf1 | GCAGCCGCACGAGCCCGCAA |
| Sumf1 | CTCGACTGGCTATTTGACAG |
| Sumf1 | AGTGAAAACGCATATCCACC |
| Sumf1 | AGCTAATTGGAGACACCCAG |
| Ngly1 | GAGCTTCAAACCCTAATGCA |
| Ngly1 | AGAGACTAGATCTAGAGATG |
| Ngly1 | GCTACACTGTGATGCATGTG |
| Ngly1 | GCTGGGTTAATACGGCACTC |
| Thap11 | AGCCTGGTTGCTGTCTACGG |
| Thap11 | GCTGCTACAACAATTCACAC |
| Thap11 | GGGACACGTTCTTGAGCCAG |
| Thap11 | CCACGCAGCGGGAAAATGGT |
| C1galt1c1 | ATATGGACACAAATGACATG |
| C1galt1c1 | TGTGCCTTGATCACTATGCT |
| C1galt1c1 | CAAGAACTATACAGTATACC |
| C1galt1c1 | CAATAACAGCGAAAGTAGTG |
| Bco1 | GACATGATGGAAGACCACCA |
| Bco1 | GAAGAGTCCCGTGAAGCACG |
| Bco1 | TAACATGGGCACATCCGTCG |
| Bco1 | GGGCACAGCAAACCTCCTGA |
| Slc29a1 | GCTGATGCAGAAACGAGTTG |
| Slc29a1 | GCATGATTGATCAGTGTCCG |
| Slc29a1 | TACACAGCCCCCATCATGAG |
| Slc29a1 | GGCCAAAATGACAACTGCAC |
| Inpp5e | GAGGTCCCTACGGATAAACA |
| Inpp5e | AGTGATCGTCACCAGCCAAG |
| Inpp5e | CACCGAGGCTGACTACACTC |
| Inpp5e | GGAGATACCTAAGTCCCGAA |
| Mrps22 | CCGATGCTCTGCTCGCGACG |
| Mrps22 | ATTTCTTTAGGCTACTAGAC |
| Mrps22 | AATAAATGACGTATTAGCTG |
| Mrps22 | TGGTATATACTCACCCGATG |
| Mrps23 | TTATATATGTCATACCACAG |
| Mrps23 | GCTTTGCCGTATCGCAAGCG |
| Mrps23 | ACTTCAAGTCTACCTGTCAG |
| Mrps23 | AGCTTTCTGACCAGATCCAT |
| Htra2 | AAACGGATCAGGATTCGTAG |
| Htra2 | GCCTCATAAGTATCCCCGCT |
| Htra2 | GTACAATTTCATCGCAGATG |
| Htra2 | CGGAGGTCAGGAGCTAACAG |
| Dpys | GTGTGCGTGTCGATGCCCCC |
| Dpys | GTGGCGATTTGACCACAACA |
| Dpys | CGAAGGTGGTAGCCGATGCG |
| Dpys | GAAAGGCAGCTCCCTCATCG |
| Lpin2 | GGATACTCACGTTTCTAATG |
| Lpin2 | GGTTATATATCCGGATCACG |
| Lpin2 | TGCAGGTCACCAAAAGAGAA |
| Lpin2 | AACCTGGATCCGTGTCACTG |
| Cubn | GTGTATCTGGAACATTCGCG |
| Cubn | AGTTATCAACTTCACCCACG |
| Cubn | ACAGCTCCGAATGCTACTGG |
| Cubn | CCACCTGTGTGAACACTATG |
| Sdhc | TACTTGTAGATAGTCAAATG |
| Sdhc | TCTGGAATAGCCTTGAGTGG |
| Sdhc | TTTGGGAACCACAGCTAAGG |
| Sdhc | CAGGAAGCAGCAGTGCCGAC |
| Ethe1 | GGGCTCAAGCTGTTGTACGC |
| Ethe1 | AACTCACAAAGCGTCCAAAG |
| Ethe1 | TCCACACCCTCGGATCAGCA |
| Ethe1 | GGGTGACCGGGAGTCAAGAG |
| Sdhaf2 | CTCTATGAGAGCAGAAAGAG |
| Sdhaf2 | CGTTAATCAGGCGATCATAG |
| Sdhaf2 | AATGATGTCACACTAGGCAA |
| Sdhaf2 | GCAGATACTCTTTAGCAAAC |
| Rmnd1 | CGGGTCCGGGATGCTTTCAG |
| Rmnd1 | GAAGTGACAAGCCTGCCTAG |
| Rmnd1 | GAAAGTACAACATGCCACAA |
| Rmnd1 | GGACAATCAAAACTGCACAG |
| Ndufa9 | GGTTAACAACGTATCGACCC |
| Ndufa9 | CCAGGTCACCCATCAGACGA |
| Ndufa9 | GATGCCTGAGCTATTGCTCG |
| Ndufa9 | GTCATACCTCACGGGAAAGG |
| Apoa5 | CAAACTCACACGTAAGGCGA |
| Apoa5 | TCACCAAGCGTTCTGCGTAA |
| Apoa5 | GCAGTTGAAACCCTACACGG |
| Apoa5 | AAAAGCTGGGACCCTTGAGA |
| Cox7b | TTATCTGCTCACCTTGGAGA |
| Cox7b | GGGAATGCTATATTAGCAGG |
| Cox7b | AACTAGGTGCCCTCTTCTGG |
| Cox7b | TAGCATTCCCATATTTGTCA |
| Acer3 | ACAGAACTCAGCGACGAACA |
| Acer3 | ATACAGTTTAACAGTAACTA |
| Acer3 | TCAGGTCATGTATGGAATGT |
| Acer3 | TGAATTGCACCAAAAATTGG |
| Ndufb9 | TGTGGATACACCATGACTCG |
| Ndufb9 | GTAGCACTCATATCTCTCGA |
| Ndufb9 | AGAATGAGAAGGATATGATG |
| Ndufb9 | AGGATACATTGCTTTCTCAG |
| Msmo1 | TAGTATGTTCCACAAATCAG |
| Msmo1 | AAAGTTCCAGATCGCAACCT |
| Msmo1 | GATAAACCAGAAACCTTCGA |
| Msmo1 | ACAGGTACTTAACTTTGGCA |
| Mrps16 | TGCTCACAACAAGTGCCCCA |
| Mrps16 | GCCACCCAAAGCAAGGCGTA |
| Mrps16 | AAAACTAGTTGCCCTCAACC |
| Mrps16 | GGCCGATTTGTGGAGCAGTT |
| Tmem126a | TTTGGCTATACCCGTCAATG |
| Tmem126a | AACTAGTGCACCCCGTGTTG |
| Tmem126a | TAAAACATACCTGTACTCAA |
| Tmem126a | AACAACTTCCAGAATCAGAC |
| Cox20 | CGCCCCGGAGCCCCACGAGA |
| Cox20 | TAGCGAAATTATACCTACAG |
| Cox20 | AGTATAGATTCTCGAGCACA |
| Cox20 | ATTAGAAGATCATGTGATGT |
| Cox14 | CACAGTGAGGAGCATCATCG |
| Cox14 | GGTAACCCCCATACACAGTG |
| Cox14 | GCGCAGCTGGAGGTAACGGT |
| Cox14 | GCCAAGCAGCTAGCCGATAT |
| Iscu | TATGAAAACCCTCGGAACGT |
| Iscu | CCAGCTCCTTAGCCACAGAG |
| Iscu | TTCATGACGTCACCACATGC |
| Iscu | GGCCAGGCTCTACCACAAGA |
| Sar1b | AAGAATTAGAATAGGCACGT |
| Sar1b | ACTTACTGGGATGTAGCGTT |
| Sar1b | TCCCGGCATTATCCAATCCA |
| Sar1b | CAATGCCATTGATAGCAGGA |
| Tsfm | CAGATACTCACCAATCGCTA |
| Tsfm | CCATGGCTCACGTTTCACGC |
| Tsfm | ACTGTCAGAACCTGACGGAT |
| Tsfm | AGGAGCTCCTTATGAAACTG |
| Cyc1 | TAGCTCGAACGATGTAGCTG |
| Cyc1 | GGTGGGAGTGTGCTACACGG |
| Cyc1 | AGCTACCCATGGTCTCATCG |
| Cyc1 | GGTCCCGGCAGCTTCCATTG |
| Pam16 | TCACCTTCTGGACCTCCTCG |
| Pam16 | CCTGGCCCAGATCATTGTGA |
| Pam16 | AGGCAGCCGCTGACGCTCGA |
| Pam16 | CCGTGTATCCACAGCCAGCC |
| Nmnat1 | AAAGGCATTATCTCACCGGT |
| Nmnat1 | GAACAGCCTGAGGTGCATGT |
| Nmnat1 | GTTCTGCCATGATGATTCGG |
| Nmnat1 | GGAGGACATCACGCAAATCG |
| Ndufb3 | GCTGGACATGGACATGAACA |
| Ndufb3 | AACTTCCAGATTACAGACAG |
| Ndufb3 | GACTCATCTTACCGAGCCCA |
| Ndufb3 | CATGGACATGAACATGGACA |
| Timm50 | GACGCCACCAGATATATGGA |
| Timm50 | GGGTCCACACTATCAATGAG |
| Timm50 | ACTGGTCCTGGAGCTTACCG |
| Timm50 | ATTCCTGATGAATTCGACAG |
| Pgm1 | CATTACCGATGGACGCGCTG |
| Pgm1 | ATCATCTCTCCCCACGATCG |
| Pgm1 | TGGGGGTTATATCAGAGAAG |
| Pgm1 | AGGCCAACTGCACAAACTCG |
| Alg14 | TGTCCAATGCCTATTCACCA |
| Alg14 | CCGAATTCCGAGAAGCCGGG |
| Alg14 | AAGAGTCTGAGAGACTCTCG |
| Alg14 | GCTTTATTCGGAGAACCAGT |
| Taz | TTTGGAGAAGCTTAACCATG |
| Taz | GGTCATCCATGCAAGACTGG |
| Taz | AGCAGCTCACCTCGACACAC |
| Taz | TATGAGCTCATTGAGAACCG |
| Etfdh | GGTACGGATTCTGATAGTCG |
| Etfdh | GAGTAGATCACACTGTTGGT |
| Etfdh | GGCGGGAAGAGGATAGCCTA |
| Etfdh | GGCAATTACATCGTACGCCT |
| Pnpla2 | GGCAGGAGGCCACGCCAATG |
| Pnpla2 | TGTTCTTGCAAAGCGCTATG |
| Pnpla2 | AGCAGGTGCCAACATTATTG |
| Pnpla2 | CCTGTTTGCACATCTCTCGG |
| Pmpca | AGAGCTCACACACATCATGG |
| Pmpca | AGAAATTGAGATGACGAGGA |
| Pmpca | TGAAACCAAAGTTACCACTC |
| Pmpca | TCAGTGGCACAGTACACTGG |
| Acadsb | GAGGATTTGAGGACACGAGG |
| Acadsb | CATAGCGAAAGAGTCGCTAC |
| Acadsb | AAGTTGAAGCACAATATGGA |
| Acadsb | TTCAAATGTTAACTGACAGG |
| Pccb | GGAAATTGAAGAATTCACGC |
| Pccb | GGAGTCCTTGGCTGGCTACG |
| Pccb | GCTTGTGCTGCGCGTCGATG |
| Pccb | TGTGAAGTCTGTTACCAATG |
| Sdhd | GCAGAGAGGACATACAGTGG |
| Sdhd | AGGACCAGCCTACCCAAGGA |
| Sdhd | TGCAGTGGCCAAGGAGCTCG |
| Sdhd | AGGGATTCAAGTACCCAGCA |
| Sdha | GTCAGTTACCTCAACCACAG |
| Sdha | TTCTACTCAATACCCAGTGG |
| Sdha | TGCACAGTGCAATGACACCA |
| Sdha | ACTGTGCATTACAACATGGG |
| Acad8 | TGGCTCCCAATATGGCGGAG |
| Acad8 | CCAACAGGATTGGGACCGAG |
| Acad8 | TCTCCATGGTACAGAGCGGT |
| Acad8 | GTGGATGCTTATATAGGCAG |
| Trit1 | GCCACTTGTAGTGATTCTCG |
| Trit1 | TGAGCGCTTGGATAAAAGAG |
| Trit1 | CCATAAACGGCTAAGCCAGG |
| Trit1 | TGTGACCAGCTACACCGTGG |
| Plin5 | AGGGACTTAGACTCACACTG |
| Plin5 | GCAGAGCAAACACCGTACCC |
| Plin5 | GAATGTGGTGAATCGAGTGG |
| Plin5 | ATGGTGGACCTGGCCCAAAG |
| Uqcrc2 | AACAAGCCGATTCTTGACAG |
| Uqcrc2 | ACAGTACAAAGGATTAGCCA |
| Uqcrc2 | CATCTTTCAAGATAACCCGT |
| Uqcrc2 | GCAAAAGCCAAATACCGTGG |
| Oxct1 | GAAGCGTTTATCACTCCGAA |
| Oxct1 | CATTGCCAGCAAGCCACGAG |
| Oxct1 | AGCTGCAGGAACTACCGTGG |
| Oxct1 | TCTAGGGCACACTTGCCGAG |
| Magt1 | TGTGTGTGCGATCGCAGCGG |
| Magt1 | CAGCTGAGCAGATTGCCCGG |
| Magt1 | TTATGCTGGACCCCTAATGT |
| Magt1 | GTGGAGCTTTAACAAGACGA |
| Gatm | ATCAAAGACTACTTCCATCG |
| Gatm | ATTCGTTGTAAGAGGAGACA |
| Gatm | ACTTGAGTGACCAGTCGATG |
| Gatm | ACAACCATCAGGATGTCTCG |
| Trak1 | GTGGAAGTAATTCTGGACGG |
| Trak1 | TCCGCTCTTAGCTTGTAATG |
| Trak1 | TTAGGCGGCTATCCCTACGC |
| Trak1 | TTACCATGTTTAGTGCGCGG |
| Mmachc | CAAACCAGCCCCCAAATCGG |
| Mmachc | CCAAACACTGAGAGACCCGG |
| Mmachc | GAAGTTTATCCCTTCCAGGT |
| Mmachc | ACTATGAGGTACACCCCAAT |
| Ndufa6 | GCACCTCCCGATACCAAGCG |
| Ndufa6 | ATATCACGGTGAAACAAGGA |
| Ndufa6 | ACTGAAAATGGGCTTCACCG |
| Ndufa6 | ACCTGAACGAGGCCAAGCGG |
| Rnaseh2b | CTCCTAGGTCAACCAAACTG |
| Rnaseh2b | TCAGCCCTTGGACCAAGTCG |
| Rnaseh2b | CAGCCTTTAGGAGATAGTGA |
| Rnaseh2b | TGGCCAGCTTTACGAACAGG |
| Lipt2 | TACACGGGCGGGCTACGCGG |
| Lipt2 | ACCCGCTTTGGTCCCCGACA |
| Lipt2 | CCGAAGTTCGCACAGTCGCA |
| Lipt2 | CAGACGCCAGTGTAGGGCGG |
| Pdzk1ip1 | CGGCGAAGACGATTGCAACA |
| Pdzk1ip1 | AGAACACAGCGACAGCAATG |
| Pdzk1ip1 | GCACCTGCCAGCTGTCAACA |
| Pdzk1ip1 | CAACCACTTCTGGTGCCAGG |
| Ndufa13 | GGTTCCGCTTGTAGTCGATG |
| Ndufa13 | ACTGACCCGACAGTCCCCGG |
| Ndufa13 | GATTCTCCGGGAAAACCTGG |
| Ndufa13 | GTGGAACCAGGAGCGCAGGT |
| Ndufb8 | GGGGTCCTAGGATATGACCC |
| Ndufb8 | CAAGAAGTATAACATGCGAG |
| Ndufb8 | CATGTACATCAGGAATCGTG |
| Ndufb8 | CAACCGATCACAGCATGAGA |
| Uqcc2 | CCAGTGGACGAGACCAAACG |
| Uqcc2 | TTCAAACTACTACAAGCACA |
| Uqcc2 | GCTCTCGTACATCTGATCAC |
| Uqcc2 | CACAGAGCTTAAGGAAACGC |
| Ndufa10 | TCGATATAGATGACTGCGTG |
| Ndufa10 | CCACTAAACTCTATGTCGAG |
| Ndufa10 | TGGGAAAAACAAGCTCGCAA |
| Ndufa10 | CTGGAGGCAATGTACAACCA |
| Slc25a19 | ACGGACCATGTATAAGACCG |
| Slc25a19 | TCAGCGCACTTTGTGTGCGG |
| Slc25a19 | GACCCCAATGCCAAATACCA |
| Slc25a19 | AGAACTGCAGGCCCGCGTAG |
| Pigc | TGTACTGACAGACTCCGTCA |
| Pigc | GAATAAAACATACCCAACCA |
| Pigc | TCCGGAAAAACATCTATGCC |
| Pigc | GGCCGACCTGAAGAGTACTC |
| Far1 | CTCTCGAACAGGCCTTCAGA |
| Far1 | ACATAGACAGAATTCACCCT |
| Far1 | CGTCGATCGTCGGCGCCAGT |
| Far1 | TCTCGTTCCTGTAGATGTAG |
| Dhdds | GGATGTCGGGATGAGGAGAG |
| Dhdds | CTATGCCAAGAAGTGTCAGG |
| Dhdds | CTTCAAACGTTCCAAGAGTG |
| Dhdds | GCAGCAGATGCAGATCACCC |
| Hoga1 | GCGGCCACGATAGTAACAAG |
| Hoga1 | ACAGTGTCCCAGCCAATACG |
| Hoga1 | TGGGTATGGCCTGGCGCACG |
| Hoga1 | CTTCCCTCTTACCTCGAAAG |
| Mtpap | TTTGCCAAATGTGTTCACTG |
| Mtpap | CAGTTGTCCAGGGACTATTG |
| Mtpap | TGTTAGCAGTCAGATCACAC |
| Mtpap | ATGTATCAAAACAGTCCGCT |
| Pnpla8 | GAAGGCGTACAAGCCTTAGT |
| Pnpla8 | AGTAATACAGAGCTTTGGGT |
| Pnpla8 | CTTAGCACTTCTGCTCCCAA |
| Pnpla8 | CATGCCGCTGGATGAATGTG |
| Slc25a46 | GTGAATTTACACCTTTACCG |
| Slc25a46 | TGAGATGGTAATGCCGAGCA |
| Slc25a46 | GCGGAACCCTCGAGCGTCGG |
| Slc25a46 | AATTGGACGAGTGATAGGCT |
| Abhd5 | AGTTTGGATCAGGGCCCTAG |
| Abhd5 | TTGAAGATCTAAGCACCGAT |
| Abhd5 | CGGCAGCCAAGAACCCTCCC |
| Abhd5 | GAGCCTGTGCGCATATCCAA |
| Abcg8 | GCGCCGACGAGTGAGCATTG |
| Abcg8 | GGTTGCTATAGCGAGGACAA |
| Abcg8 | AGTCTGTACTTCACCTACAG |
| Abcg8 | TGTGATCACGTCGAGTAGTG |
| Agpat2 | TGATTAGAGATGATGACACA |
| Agpat2 | GGGTACACGCAACGACAATG |
| Agpat2 | AGCGTGAGCTAATGTTCACA |
| Agpat2 | GCCGCACCGTGGATAACATG |
| Uqcrb | ATGCCTCATAGTCAGGTCCA |
| Uqcrb | GCTGCAGGATTCAATAAACT |
| Uqcrb | TAGTTTCAGCATCAAGCAAG |
| Uqcrb | AGCCATAAGAAGGCTTCCTG |
| Cog6 | AGTCGTTGGGCATATCACCG |
| Cog6 | TTGTGCAGCTTACGCGACAG |
| Cog6 | CGGACTCGAAGAAATTTACG |
| Cog6 | GGTCAATCACTATACCTCAG |
| Slc39a8 | TCAGCTGCTGTAAGATCGCG |
| Slc39a8 | AGGGGGTTAAAATCAATCCC |
| Slc39a8 | CGTTAGGCTCAGTGACAGCG |
| Slc39a8 | CGGCGCCAACCGGAGCCTGT |
| Pigm | GTTTGTACGACTTCCTGAGG |
| Pigm | CCGCGCGCTTCGTCACGGAA |
| Pigm | CGCGAAGCCATAGAACACCG |
| Pigm | AGTACGGAGAGAAATTATGG |
| Alg13 | GATCTTGTCATTAGCCACGC |
| Alg13 | GTGAATGACTCAGTACGGAA |
| Alg13 | GGTTGTAACCCAGACTCTCG |
| Alg13 | TTTCGACGAGCTCGTCGCAC |
| Slc25a26 | GGCATTCAAGGACTGTACCG |
| Slc25a26 | AACACCCTTACCGTTAGGAA |
| Slc25a26 | CCAGCCTTGTTAAATCCCTG |
| Slc25a26 | ACTTTCAAGGATTCCCACAA |
| Sdhb | TGCGCCATGAACATCAACGG |
| Sdhb | ACAGTATCTGCAGTCCATCG |
| Sdhb | ACCTCGAATGCAGACGTACG |
| Sdhb | TAGAAGTTACTCAAATCCTG |
| Dnajc19 | AATTTCTTTCAGGCATTCGG |
| Dnajc19 | TTAGCCCTACTGCCAATAAA |
| Dnajc19 | TTTACAAGCCATGAAGCATG |
| Dnajc19 | CCAGCACAGTGGTAGCAGTC |
| Jagn1 | GCACTACCAGATGAGGTACG |
| Jagn1 | GTACCTCATCTGGTAGTGCA |
| Jagn1 | AGTGCATGGCGACGCGCTCC |
| Jagn1 | CCGTCGGTGCCGGCCGCTCG |
| Dcxr | TATGGTTGGTCAGTGCACGT |
| Dcxr | GGCGGTGAGTCGGACGCGAG |
| Dcxr | GCCAAGGGCATGATAGCTCG |
| Dcxr | CCTAAGCAATGTGGGACCCG |
| Coa6 | CAGCCCCTTCCATGAAGGAA |
| Coa6 | GCGGGACTTACCCACTGCTG |
| Coa6 | CGTCCAGGCAGCGCCAGTAC |
| Coa6 | GCGCTGCCTGGACGACAACG |
| Coq9 | CTCAGGTACACAGACCAGAG |
| Coq9 | GTGCCTTCCACACTGCCGTG |
| Coq9 | GGGGCCAGTGCTCAATGTAG |
| Coq9 | GTTGAGGCGAGCATTGCACT |
| Npc2 | GTCAACATCACCTTTACCAG |
| Npc2 | TAAGGTGGGAGTTATAAAGG |
| Npc2 | CCCCTGCACTTCAAGGACTG |
| Npc2 | GTCAGGCTCAGGAATAGGGA |
| Atad1 | TACGCACATGCATATTAAGA |
| Atad1 | TAGATGCCATTGACCCCACC |
| Atad1 | AAGGCTGTAGCTTTATGGCA |
| Atad1 | CACTTATCAGTCAGTGTCGA |
| Apopt1 | TGAGCGCCCTACGGATCACA |
| Apopt1 | TGTGAAAATGAACGGGACGA |
| Apopt1 | CAGTCTTGCCATGATTGGAT |
| Apopt1 | AGAATGGAATCAACAGTTCT |
| Gcsh | GCGGCGGCGCTCGGTACACA |
| Gcsh | AGAGGAAGGTATTGGAACGG |
| Gcsh | AATTCACAGAGAAACATGAG |
| Gcsh | GGGACAAAATTGAAAAAACA |
| Rnaseh2c | CGTGAAGAAGCGATCTACCG |
| Rnaseh2c | CGGCTGACTAGAACGTCGCA |
| Rnaseh2c | AGGGTTTGCGGGATTCGTGA |
| Rnaseh2c | ACCGTCTGCATCGTGGCGGA |
| Pdhb | TCTTAATTGTAGGTTAGCAG |
| Pdhb | GGGCACAGGCTGAAGGCCAG |
| Pdhb | TGAAGCTATTAATCAAGGTA |
| Pdhb | GCCATTCGTGATAATAACCC |
| Slc25a22 | GTGTCTTAGCCAGGTCGATG |
| Slc25a22 | ATACATGCCGAAGTAGCCCT |
| Slc25a22 | TGATTCAGGTTGGCAAACAG |
| Slc25a22 | GACACCAGCTCTCTAAGGAT |
| Pomgnt1 | GCCTTCGTGGGACGAAAAGG |
| Pomgnt1 | CACTCGGAGAGCAATCAGCG |
| Pomgnt1 | GCTGTGTGTTCATACCCCTA |
| Pomgnt1 | GGGCTGAACTCAATAGGCGT |
| Mto1 | TTCGACGTGGTAGTTATCGG |
| Mto1 | AGTGTAACAGATAACCCTTG |
| Mto1 | ATGGACTCTTTAGCGAGTCG |
| Mto1 | ACTTTATAAACAGAACATGC |
| Stt3b | AGGGTACATATCTCGGTCAG |
| Stt3b | CGACAGCATGCAGACGACCG |
| Stt3b | GTCATCTATCTGACATACAC |
| Stt3b | TACAGCAAGAGAGTCTACAT |
| Sdhaf1 | CGGACCCCGCGGGCGCACGA |
| Sdhaf1 | GCAGCACGTCGGTTCGCGGA |
| Sdhaf1 | GTATCGAGTATCTGTATCGC |
| Sdhaf1 | AGGTACCCATGGCCGTGGCG |
| Ndufs3 | TTGTGGGTCACATCACTCCG |
| Ndufs3 | TACGTGCTGCCACAGCACGG |
| Ndufs3 | CAGCGTTGGGATGACTCCAT |
| Ndufs3 | GGGTGTCAGCTCATCTGCAT |
| G6pc3 | TAGGCCGACTGCCAATAGGA |
| G6pc3 | TTCCCGGGCTAGAGAATATG |
| G6pc3 | GGGGCTCATTAGCCAGCCAA |
| G6pc3 | TAAAGAGAGTCCAATACATG |
| Lmbrd1 | TTACAAACAGCAACGCAACG |
| Lmbrd1 | AAGGCACGTCTATCCCTCGC |
| Lmbrd1 | CATCATCTGTTTCAGGCGTA |
| Lmbrd1 | AACGGTATTCTCAATCTGTA |
| Slc39a13 | ATAAAGAAAGCGAGTCCTGG |
| Slc39a13 | TACACCTGTAACATCACCCC |
| Slc39a13 | CTCACCTTCTGACTGTAACA |
| Slc39a13 | CTGGGGCTATGGGTCATCGC |
| Gpihbp1 | GCAGGTATCAGTACACCACA |
| Gpihbp1 | CACTGTGCAATATTCCACCC |
| Gpihbp1 | CTGTCAAGTGCTTCACAGCG |
| Gpihbp1 | CCTGCTTCCAGGGATCATGT |
| Tmem126b | AAGATGGCACTTTATAAACA |
| Tmem126b | AAAGAGCTTGTAAGAAACGA |
| Tmem126b | GCACTAATTGGCATGGCATG |
| Tmem126b | CCGTCTAGGTGACATAAGCA |
| Ndufaf4 | TAACGTATACATTACCGGCA |
| Ndufaf4 | AATATCACGGACATTCCCAA |
| Ndufaf4 | AAGGAATTCAGACTGCCGAT |
| Ndufaf4 | CGCCTTAAGGAACTTCAACG |
| Dpm3 | AAGTTAACACAGTGGCTTTG |
| Dpm3 | GCGATAGCCCACCGTGCCCA |
| Dpm3 | GGCCACAGGACCTCTCGGCA |
| Dpm3 | ACCAACAGGTAGGCAGGCAG |
| Mocos | GCTTCAGCGCTATTACATCG |
| Mocos | GCTGTAAATGCGCACCACAG |
| Mocos | GGTGTGGAAGCTGACAGTCG |
| Mocos | CAAACTCACCCACGACACTG |
| Pmvk | AGCTGGTGAGTGACACACGG |
| Pmvk | CTACAAGGAGACCTATCGGA |
| Pmvk | GCTCTCTGGTCCACTCAAGG |
| Pmvk | TGTCTGTATCACAGCCCCAT |
| Elac2 | TGTGCTGCGCAGGTGAACGG |
| Elac2 | GGGAAGAGTATCACTTACGA |
| Elac2 | CTCAGCGTGCCAACAGTTCG |
| Elac2 | ATGCATTGGTCAAATGTTGG |
| Nadk2 | GCCTTACGGAGGTTCTCTCG |
| Nadk2 | CCTCCGAGAGCTCCGCGTAG |
| Nadk2 | ACTTCATTGAGCGCTCTCAC |
| Nadk2 | GGTGGTAGCCCCGCCGACGG |
| Elovl5 | CTGCCAGGGAACACGCAGCG |
| Elovl5 | AGATGTTGAGCATGGTAGCG |
| Elovl5 | GACCATAGTACGAGTACATG |
| Elovl5 | ATCCTGCAGTTGTATAACCT |
| Vars2 | ATCCCCGGGTCTGATCACGC |
| Vars2 | TCCAGATGACCCGAGATACA |
| Vars2 | CCAACCCACGTCACCTCTCG |
| Vars2 | TGCAGATGGGCAGTACCATG |
| Pink1 | CATGGTGGCTTCATACACAG |
| Pink1 | GTCGGACTTGAGATCCCGAT |
| Pink1 | CGAGGAACAGTGTGCGACCG |
| Pink1 | CCAGCCCAGCTACCGACTGG |
| Phkg2 | AATGTGCATCTCTCGCCGTG |
| Phkg2 | CGGTAGCTCGATGGACACAG |
| Phkg2 | CATGAAGCTAGAAGACTCGT |
| Phkg2 | GTAGCATCAGGATTTGGCGC |
| Isca1 | CTTGCTCACAGCCCGCACGG |
| Isca1 | GAGGTGACTCACCAGTGTGA |
| Isca1 | AAAGTTGGCGTGCGAACCAG |
| Isca1 | CAACAGTTACTTACATGCTC |
| Pycr2 | GGTGCCCGTGGCATACACGG |
| Pycr2 | GATGTGCCTCTCCTGCACGT |
| Pycr2 | CGTGGGCAGGTCCATATCCG |
| Pycr2 | GATGCTATCACCGGGCTCAG |
| Gmppa | GGAGCCAGATCAAGTCTGCA |
| Gmppa | TGGAAGGTTAAACTCCTGCT |
| Gmppa | AGGACACAATCCCTCAACTA |
| Gmppa | ATCAGTGACATCATCAACTG |
| Mrpl44 | AAACTGGGATTACCATGCGG |
| Mrpl44 | TAGACTCTCAGTGCCTTCGG |
| Mrpl44 | TTCTTCACTCCCCGAACCGG |
| Mrpl44 | TGAGGCAGTTGTCTGTCACG |
| Ndufaf5 | GCGACACACTCTATGAACTC |
| Ndufaf5 | TCAGGTAGTCAAATTTCATG |
| Ndufaf5 | ATCGAGCGGCAGTCGCAGGG |
| Ndufaf5 | TCAGCTAAAATATTCACAGT |
| Hfe2 | GGCTCACCGGACACGCCGCG |
| Hfe2 | GTTGGCTCCCGACGAAACCG |
| Hfe2 | CTATGCCATGCACCGCAGAG |
| Hfe2 | ACAAACTCGAGCCCCCAGGT |
| Afg3l2 | AAAAGCGATAATGAGTACGG |
| Afg3l2 | GTTCGTGTGACCTTTACACC |
| Afg3l2 | GCCTGCCAGGATGACCACAT |
| Afg3l2 | GTGAAGTTTAAAGATGTGGC |
| Mtfmt | GTGTACGAGTGGCCCGACGT |
| Mtfmt | GTGCCTTCGCTATCCCCCAA |
| Mtfmt | ACCGTGTGGATTATCGGAGC |
| Mtfmt | GTCAAAACTTGGTGCCAACA |
| Pitrm1 | TTGGGTGATAATGAGTAGCG |
| Pitrm1 | ATATAATGGCTATACACGGG |
| Pitrm1 | AGGGAATTCCATATAACATG |
| Pitrm1 | GGTGAATTCTCTCTCCCCGG |
| Ccdc115 | AGCTGACGCCCAGACCCCCG |
| Ccdc115 | ACAGCTGCTCAGTGACCTGG |
| Ccdc115 | CTAACTCTTTGGTCTTGGTG |
| Ccdc115 | GACCTGAGGCTCCATACGCG |
| Slc52a3 | GTGACCTCCTGGATACAGGG |
| Slc52a3 | TGTCTCCGTGACATTGACAC |
| Slc52a3 | GCTGGTGACTGAGTTGCCCG |
| Slc52a3 | GGGTCGGAAGCGGTGCATCA |
| Ndufaf1 | AGACGGCCGCCCATTACAAG |
| Ndufaf1 | GAACAGGCCACACCTACCCG |
| Ndufaf1 | TCTGTATCTCCGAGTTCGTG |
| Ndufaf1 | ACTTCTGATAAGACAATTGG |
| Cad | CGCAGGGGTACCCGACCGTG |
| Cad | AGGATTAGAACCTTTCGTGG |
| Cad | ATGGTGAGTGCCCACCACAA |
| Cad | CTCAGAAACTCTGTTACGGG |
| Rnaseh2a | TGTCACACGATACAGCTGCG |
| Rnaseh2a | GTAACAGATGGCGTAGACCA |
| Rnaseh2a | AGACCTTGACAGAGAACGAG |
| Rnaseh2a | GGCTCCTTGAGACACACAGC |
| Ndufa11 | CAACCCCGCAGATTCCACCC |
| Ndufa11 | TCTTACCATGAGGTCCCCGA |
| Ndufa11 | GGCCGGTACACATTCACTGC |
| Ndufa11 | AGCGGAGCCGATTATGCCTG |
| Coa7 | GTTGAATCCTGTCACAACGT |
| Coa7 | CTACCAATGCTACCGCGAGA |
| Coa7 | AACTGTGAGAAATACGGGCA |
| Coa7 | AGGATGGCCAGCCTGACCTG |
| Slc25a32 | CGGATCTTCACGAGGTCGAG |
| Slc25a32 | GGTTCTCGTACCGGACGTGG |
| Slc25a32 | GGAGTAACCCCGAATGTGTG |
| Slc25a32 | AGGTGTGCGTGGATTATACA |
| Agk | ATATTGCAGACGGATTATGA |
| Agk | CTCTGAAAGATCCCCACCGG |
| Agk | GAAATACCCTTTGCAAGCTG |
| Agk | AGTACTGAGAAGAACAGATG |
| Fars2 | CACACACATCAGCGCATCAG |
| Fars2 | ACCTTAAACAAGTACTGACT |
| Fars2 | TGGAGGAAGCTGGTCATAGA |
| Fars2 | ACTTGGTAAATCCTACCCTC |
| Trnt1 | CATGAATTAAGAATAGCAGG |
| Trnt1 | ATTCGCATGATCAACAACAA |
| Trnt1 | AACCTGTTCTCACCTATGTG |
| Trnt1 | GGCAGAATTGTCGACAGACC |
| Sdr9c7 | CACCGCCGAATATAGCCACA |
| Sdr9c7 | AGGTCAGCATCATTGAGCCA |
| Sdr9c7 | CCGACTTGGTGACATCTAGG |
| Sdr9c7 | GCCAAACAACTGGTTGATAG |
| Yars2 | CCACGAGCGCGATCACGTTG |
| Yars2 | TCCCCCCAGACGGTGTACTG |
| Yars2 | CTGGGCGGATCAGATCAGTT |
| Yars2 | AGAACTCCTTGAACAAGCCG |
| Taco1 | GCGATGCCTCGGGTTCCGCG |
| Taco1 | GCTGACTCGATGGTTGACTT |
| Taco1 | ATTGAGGCGTTATCAAACAG |
| Taco1 | CCACTTATTGTGTCCCGCCG |
| Pigw | TGTGGCTGTCAGCATAACTG |
| Pigw | ACAGTCAGAGTGATCACTAG |
| Pigw | TGTTGTATCAGATATACCAC |
| Pigw | GTTCCAGTGAATGCCGTACT |
| Gtpbp3 | GTGGAGTTTCACGTACACGG |
| Gtpbp3 | CCACAACATTTGCCCCCGAG |
| Gtpbp3 | GTCAATGTAGGCCTCCACAT |
| Gtpbp3 | AGTCTCCAGCACGTCACGGG |
| Cox10 | TCTGTCCCGGAAGCCAAATG |
| Cox10 | GGGAGTGAATCCACTCACAG |
| Cox10 | TATACAGGGATTGCCACACA |
| Cox10 | AGTTGGCAGCACAGGATGCG |
| Tmem70 | CGGGATCGCTTTCTGGCGTG |
| Tmem70 | GAAGGTACGGCAAGAATGCA |
| Tmem70 | GCTGATTTATACTGGAAACC |
| Tmem70 | GTAAGAAGGTGAAGCAGCGT |
| Mipep | TCCACGTTTGTATTCAACCT |
| Mipep | GATTGCCACTTCACCATCCG |
| Mipep | TGGGACCCTCCCTACTACAG |
| Mipep | CTGGGGCTTGACGTTGAAGG |
| Atp6ap2 | TTAGCATATTAAGATCGCCA |
| Atp6ap2 | TGAACTTGGGAAGCGTTATG |
| Atp6ap2 | CCGGTGGAATAGGTTACCCA |
| Atp6ap2 | TGGACAGTGCAGCTACGTCT |
| Wars2 | GAGCTGGGTGAACTTACAGG |
| Wars2 | CACACACGTTCCTGTCGGGG |
| Wars2 | CAGAAGCATGATGGGACCGT |
| Wars2 | TGGTGTAAATGCTGCAATCG |
| Kynu | GCGGATGGTAAAGCCACGAG |
| Kynu | CAAACGCCCTTGGATTGTAG |
| Kynu | TCTTTAAGCCTACTCCAAAG |
| Kynu | CAACACCCCAGTCATGTAAG |
| Slc29a3 | AATGATGGCCATGCACGCGA |
| Slc29a3 | GGGAAACTGCGCAGAACCCG |
| Slc29a3 | CAAGGAAGACTGCTGCCATG |
| Slc29a3 | ACCAGAAAACACTCGAACTG |
| Pdss2 | ATGATATTGGAATCTCGACC |
| Pdss2 | TGCGTGTCAGAACTACGACA |
| Pdss2 | CGGCATAACCTACAACTGCG |
| Pdss2 | AGCTCTTCTACAGAACACCA |
| Pex1 | TTGGTCCCAAGGAAATACGT |
| Pex1 | TATTACCAAAGGAAGCATCG |
| Pex1 | GCAGCTTCATACGAACTCAG |
| Pex1 | AGTGTTGTGAAGAATAAACT |
| Cyp2u1 | GCTCCTTAAATGGACCAAAG |
| Cyp2u1 | CAGGCGCAGCAGCTTCGACG |
| Cyp2u1 | GAAATGACGAAGCGTCGAGT |
| Cyp2u1 | AAGTGTTCAGTGACCGCCCG |
| Fuca1 | ACTGCCGAGCTGGTTCGATG |
| Fuca1 | GTTCGGCCCACTGATCCGGG |
| Fuca1 | CCAACTCACCGACCAAATCA |
| Fuca1 | GTGATAGTGAATGACCGGTG |
| Pnpt1 | AATGTGTCGTTAACCCAACA |
| Pnpt1 | ACCAACGGCATGAATTGGGA |
| Pnpt1 | GAAGAGATCTGACTTCACTT |
| Pnpt1 | TTGCCATGCTATCAAAGTTG |
| Ubiad1 | GGCTTCCCGAACGATCCTGG |
| Ubiad1 | CCAAATCGAACAACATCCTG |
| Ubiad1 | CCAGCGGGCCGAAAGTGATG |
| Ubiad1 | TCAGAGCGGACAGGTAGTAG |
| Tmprss6 | GGTTCGATGCCTACGCACTG |
| Tmprss6 | GGAAAGCAATAGACTCCCGT |
| Tmprss6 | AAGACCTCATGATCAAAGTG |
| Tmprss6 | CCTTGATCCCGTCACACGCA |
| Tars2 | CTTTAAGCTTCATCTGATCG |
| Tars2 | AAGAGTCTATAATGCCCTGG |
| Tars2 | CTACTGCTACAGTGTAACAC |
| Tars2 | TCCCCAAAGTAGAGTTACTG |
| Coq2 | ACCAGTCTGGAAAACAACCT |
| Coq2 | ATGCGCCTGGACAAGCCCAT |
| Coq2 | AGGGCCAATGGGAGCCCGCG |
| Coq2 | CTGTCCCCCAAGGAAGACGA |
| Cars2 | TTAGATTTGACATTATCCGG |
| Cars2 | TCAGATCATTGCTTTCATCG |
| Cars2 | GCACATCGAGTGCTCTACCA |
| Cars2 | ACAGCAGCGACTTCGCCCTG |
| Sars2 | CCGGTCCTATTACCTGCGTG |
| Sars2 | CTGTACGAACACGCACGAGA |
| Sars2 | TCGCTTCGAAGACCTTAACC |
| Sars2 | ACGGTCCCAGACCTTCTGAG |
| Trmu | AGGGGTATTTATGAAGAACT |
| Trmu | GTGATGGAGGCTATTCTCCG |
| Trmu | GCTTATTGCAGTTGATGTCG |
| Trmu | ATTCCAATACTCCTTCACAT |
| Slc39a4 | GGTCCTGAATACGGATAGTG |
| Slc39a4 | CAGTTGGGGAAGATCTACAC |
| Slc39a4 | CATGCAGCGTGATATTGGGA |
| Slc39a4 | TGGAGAGGGTCACACCCATG |
| Mccc1 | GTTATGATCAAAGCAGTCCG |
| Mccc1 | AACAGGAGGAAGTATTACCA |
| Mccc1 | CAGGAAGAAATCCCTCTGCA |
| Mccc1 | GCAGTACCTCAGCTTCGACA |
| Ddhd2 | AGAAAATGAGCAGATCGGAA |
| Ddhd2 | TATTGTATTACACAACCCAA |
| Ddhd2 | GTCCACAATAGTCTGACAGT |
| Ddhd2 | TACCACAGTGAATATCAACA |
| Pex13 | AGATGAGATAAGGACCGCCA |
| Pex13 | TGGGAGGAAGATCATCTACA |
| Pex13 | GGCCTGGGAAGAATAGGCGG |
| Pex13 | AGCCCTGAAACTGTTATAGA |
| Chst14 | GAAACACCAAGTCACTGCGG |
| Chst14 | CGGACCTTGAGGGCCGTGTG |
| Chst14 | GCTGATGTTCGCTGTAATCG |
| Chst14 | TGTTACGGTAAGCAGACAGG |
| Cep89 | AACTGTCTGCCCAGTCAACG |
| Cep89 | TGAGATGTTAGGCTACGGGG |
| Cep89 | ACACTCTGTTCTGAATCTAG |
| Cep89 | GTTTCACTTACCAACCAGGG |
| Mfsd8 | AGCAATGTATGATCGGACAA |
| Mfsd8 | TGGACCGAGAATAAAGCCCA |
| Mfsd8 | GGATATGTACGCCTGGACCC |
| Mfsd8 | GAGCATCGTGTGGATGACTT |
| Lrpprc | AGCTGATAGACTACTGTCGG |
| Lrpprc | GAATAGCCGAAGCATCTAAG |
| Lrpprc | CCGAACTGAGTCTCGCCGAG |
| Lrpprc | GAAAGACCTTCCGATCACAG |
| B3gat3 | GGACTCGCGGTGTCTCAGTG |
| B3gat3 | TCTACACTGGCTGCTAGTGG |
| B3gat3 | AATGACATAGATAGTAGGCA |
| B3gat3 | ACGGAGATCAGCTTGCAACT |
| Ttc19 | CAGGTCATAAGTGTACGTGA |
| Ttc19 | TCTGCATTTCAACTCTAGAG |
| Ttc19 | AAGAGTCTAAGCACATACCA |
| Ttc19 | GCAACGATGAGTTATCTGCT |
| Ndufv2 | ATTCAATACCTTGTTCATAG |
| Ndufv2 | AAGGTGTAGTAGTGCAGACC |
| Ndufv2 | TAGTAAAAAACTACCCAGAA |
| Ndufv2 | CTCTTACCCACTGAGCAGCG |
| Phykpl | AGACTAGCTCGACAGTACAC |
| Phykpl | CCAGAAGGAATGGGTCCATG |
| Phykpl | AGTACCTGTACGATGAGCAA |
| Phykpl | CGTGTTGAAGTATTCTACAC |
| Appl1 | CCGACTGTACCTATTAATTG |
| Appl1 | AACTTAATGAGTCAAGCCCG |
| Appl1 | GCCTGCATGTATCCCAGTAG |
| Appl1 | AATTGTATCAAGCTATGCAT |
| Glrx5 | GCACGGTGTTCGCGACTATG |
| Glrx5 | GCGTTGCTGAAGCCGCACTG |
| Glrx5 | AGGTGGTGGTGTTCCTCAAG |
| Glrx5 | GGAGTAGTCTTTAATACCTG |
| Pmpcb | GAAATATAACTAACCTCCGG |
| Pmpcb | GGACCGACTAAGAAGCACAC |
| Pmpcb | GAGACAGATCGTGTCTGGAT |
| Pmpcb | CTTCCAGTGAACTTGCAGGG |
| Slc25a42 | GCTGGGTGTCATTCCCTATG |
| Slc25a42 | AGGAATCACTCGCACCATGG |
| Slc25a42 | GTACGTGAGGGAAGCCGCAG |
| Slc25a42 | CTTCATCCGAATCTCGAGAG |
| Mcee | TGGAACTGCTTCATCCACTG |
| Mcee | TTCAAGAATCCAGTCCTGTG |
| Mcee | AACATCCCTGTAAAATGACG |
| Mcee | TGCAACTGAAGTTTGGACTC |
| Dnm1l | AATCGTGTTACAATACTCTG |
| Dnm1l | GCACAAATAAAGCAGGACGG |
| Dnm1l | GATGCCATGGATGTATTGAT |
| Dnm1l | GTGACCACACCAGTTCCTCT |
| Plce1 | CAGGTATTCATCGCGCAGCG |
| Plce1 | GTGACTCCTCGGATCCATGG |
| Plce1 | CACCGGCCATCAGCTCAAGG |
| Plce1 | GTGCAGTAATCTATTGTCCG |
| Abcb6 | CTCCAGAGGTCACATAACGC |
| Abcb6 | TGAAGTAGACCGCTATCGAG |
| Abcb6 | GTGCAGCAGTTTACGTCCCG |
| Abcb6 | TAGGTGCCAAACCAGTTGAG |
| Dmgdh | TTAGCCGGACTGTATAACCC |
| Dmgdh | AGGATCCTTCCAGGTCGCGG |
| Dmgdh | TGGGATGTGGAGACTCCACA |
| Dmgdh | ACAGCATCAAACTTTACGAG |
| Lonp1 | TCATTCGCAACTTGTCTCCG |
| Lonp1 | GGCAGCGACGAGACCTCCGA |
| Lonp1 | GCGCTTTATCAAGATCGTGG |
| Lonp1 | CAAGGCGATGATATCCCGAA |
| Opa1 | GCGCCTGCGAGAGCTCGACA |
| Opa1 | AGGTTGTACTGTTAGCCCCG |
| Opa1 | AGAGCGTGTCATCATCTCGC |
| Opa1 | AAGTGACAAGCATTACAGGA |
| Ehhadh | TTCATATGGATGCTTCACGG |
| Ehhadh | CCACATCATGAGGTTACTAG |
| Ehhadh | GTAAACCCATAGAACCCCGC |
| Ehhadh | TCACTATGGCTCTAACCGTA |
| Acbd5 | AGTGTATGAAACCAGATTTG |
| Acbd5 | CCAGGCCGTGAAAGTTTACA |
| Acbd5 | GGAGCAATTTGGACAAGAAG |
| Acbd5 | AAAGCTACCTTTATAGCCAT |
| Gbe1 | GGCTATGAACCACATCTAGG |
| Gbe1 | AATCCCCCCTGATACCTCAT |
| Gbe1 | AGACTCGATAACAATAGCAG |
| Gbe1 | GAACTATGATTGGATACACT |
| Gale | ATCTTCACCGATGCGCCCAG |
| Gale | AGAACTTGGACTTGCCGTAG |
| Gale | TTAACTCTATAGTAGTCCAG |
| Gale | CTGGGGGTTCCCGTACACGG |
| Isca2 | CTTGTCCCTAACTGCCGAGG |
| Isca2 | TCATTCCTAGGCGCTCGGGA |
| Isca2 | GTCCTCCTTGCCTGTCGTCG |
| Isca2 | CTCAGGCTGCAAGTAGAGGG |
| Slc6a19 | ATCGGTCAGAGGCTACGCAA |
| Slc6a19 | ACAGTCATCAAAGCGCTCAG |
| Slc6a19 | AGCTGAGCAACCCCAACACG |
| Slc6a19 | TCCGTGGCATCGAGACCACT |
| Pank2 | GTCCGCAGACTCGCCGACCG |
| Pank2 | GGATCCGTAAGCCACATTGG |
| Pank2 | AAAGCAGGCATGTCATGGGT |
| Pank2 | CGAGTAGCGCGGCGCCGTCG |
| Mgme1 | AAAGTGGAGAAACTCACAGG |
| Mgme1 | GGAGAAGAGCACCCCCAGTG |
| Mgme1 | CTAAACGAGAGTACTTCGCT |
| Mgme1 | CAAACAGTTCCATAAAGCCT |
| Pck2 | TGCGTATTATGACCCGCCTG |
| Pck2 | TGATTGTAACTCCTTCGCAG |
| Pck2 | AGGGTTTGGATGCTACGGCA |
| Pck2 | ATGGAAGCACATACATAATG |
| Shpk | CATGGTTTGAAATCCCAAAG |
| Shpk | GACCAGAATGCCGCTAGCTG |
| Shpk | GCAGGCTGTGAATGGATGGA |
| Shpk | GTAAGGGAAGAAGGCAACTG |
| Pomk | CCCATTCTGAAGTAGCCCGG |
| Pomk | GAGTGAGCACGTGGTCACGC |
| Pomk | CACATGACCCTCGTGCCCAG |
| Pomk | CGCCACGTTCATAAGGGCCA |
| Dhcr24 | ACCTGACCCATAGACACCAA |
| Dhcr24 | GTAAGACCTTCATGTGCACG |
| Dhcr24 | AGATGCTGCGCGTGTGTCGG |
| Dhcr24 | GTCTGCCAGGATCAGCTCGT |
| Atp13a2 | CGTCATCACAGGTACGACCG |
| Atp13a2 | CCAGCTACATGACACCCCGG |
| Atp13a2 | GGGACCCCGTGTGCTAGCAG |
| Atp13a2 | TGTGGTAGTGACATGGCCAG |
| Ppa2 | TGGTTGTGGAAATACCTCGG |
| Ppa2 | CCGAACATCTTCCCTCACAA |
| Ppa2 | TACAGGGACATGACGCGGCG |
| Ppa2 | TAAGAGCACCGACTGCTGTG |
| Ndufs7 | AAGCGGTCCATGTCATAGCG |
| Ndufs7 | CATCAGAGTGTAGCCACTGA |
| Ndufs7 | TGGCACGCTTACCAACAAGA |
| Ndufs7 | CCCGGCGTGCCCAGTTGATG |
| Secisbp2 | AGTACGACTACAGCCAAGCA |
| Secisbp2 | CTGTACAAATGGATAGTAGG |
| Secisbp2 | ACCGTCAGGACTTCATAACT |
| Secisbp2 | TGCCGAACAATATAACCCAT |
| Oplah | TGGTGCGGATTGTTCCTCGG |
| Oplah | TGACTGTCATCACACCGGTG |
| Oplah | CTGTGCTTCGTGAACCACAT |
| Oplah | GCTCCAAAGTCACCAGCACG |
| Lyrm7 | TCTTAACAGATGATGAAACT |
| Lyrm7 | AATGATAAGAGAGCATTGGA |
| Lyrm7 | AGCTCTTTAAAACACTGCAC |
| Lyrm7 | GATGAAACTAGGTTCTGATG |
| Ndufaf2 | CAGGAGACATCCCAACCGAG |
| Ndufaf2 | ATTCTTACCTCCATAGTGGG |
| Ndufaf2 | ACGTGGGCACGGACCATCTG |
| Ndufaf2 | TTGGAGCGCCTTGTCCAGGG |
| Gns | CCAAAGTGTTGTTAACGACG |
| Gns | TGCTGGGAAGTATTTAAACG |
| Gns | TCAGACAACGGCTACCACAC |
| Gns | CAGAGGAACCTACCGTCCCG |
| Fastkd2 | ATGTGAATGCGCTTCGAGCA |
| Fastkd2 | TGAAAGAGGTATCTAACAGA |
| Fastkd2 | TAGCATTAAGGCTGAACAAG |
| Fastkd2 | ATAAGCTTGTCCATCAAACC |
| Mff | GGATAAGCGACAAAATGCCA |
| Mff | GCAAGTCCCAGAGAGGATCG |
| Mff | AGGTATTAGTCAGCGAATGA |
| Mff | TGTTCGCCAAAATGGACAGT |
| Ispd | CGGCGGCCACGGTTCCAGGG |
| Ispd | TAAGCGCATCTCACTAGCTG |
| Ispd | TGGTCACTTAGACCACTCAC |
| Ispd | TCTTAGCAGCTAAGGAACAT |
| Cant1 | GGAGTGGGATAAAGACCACG |
| Cant1 | CAGCACCCACAAAGGACGTG |
| Cant1 | CCATCAGAAAGGATCACCCA |
| Cant1 | GGAGTGGACCACCACGACAG |
| Coa5 | CCGGTATTATGAGGACAAGC |
| Coa5 | ATCCGCTACCTGGAGCACAC |
| Coa5 | GCGGGCGTGAAGGAGGATCT |
| Coa5 | GCCGGAGGGCGGCGCGTGTG |
| Abhd12 | TGACTCAGTAGGAACACCAT |
| Abhd12 | GATAATCCTGTGTATATTTG |
| Abhd12 | GCCGCACAGCGCATCCGACG |
| Abhd12 | GCAGGTATAGTATGATGGCA |
| Grhpr | TGGAGCCCATTATGGATGTG |
| Grhpr | TCCAGAGATTTCTTTACACG |
| Grhpr | CTTTGGATGAAATCAAGAAG |
| Grhpr | AAGAAACTTCTGGATGCCGC |
| Cog2 | GACGGTGTGCCAACAAAGGT |
| Cog2 | TGCCACCATTGATAAGACAC |
| Cog2 | TCTTCTCGTAACTGTCCCAA |
| Cog2 | GTGATGCCAGCTATACGCTG |
| Trmt5 | AGAGGAAAATATGCTGACCA |
| Trmt5 | CAGCAGTAAGCATCCTGTAG |
| Trmt5 | ATTCCTCGAACGTCAGAGGG |
| Trmt5 | TTTCCTAAGTCTCGGCACCA |
| Lmf1 | GATCTGCCCTACGTTGACCA |
| Lmf1 | CTTCAGAATCATGCTTGGAG |
| Lmf1 | GGTGGCATTCAATCAGAACA |
| Lmf1 | CCTATGCACCAACCATTATG |
| Cln6 | CCTGCACGAACGCGACACCG |
| Cln6 | CTGACATAGACTATAGACCG |
| Cln6 | TGGATCGGGAATACCAGCTG |
| Cln6 | ATAGTAGTACAACAGCTCAA |
| Qrsl1 | ACGTCCATTGAATTCACCAG |
| Qrsl1 | ATCAGGGTGCCCTACTCATG |
| Qrsl1 | TTACGCAAAGCAGGTGAACG |
| Qrsl1 | GCTGTGTATAGGAATCCCAA |
| Mfsd2a | TCAGGGACGGAAAGTTCACA |
| Mfsd2a | GCAGCCGGTCAACTGGTACG |
| Mfsd2a | GAGGGACTTACTGTATGCCG |
| Mfsd2a | TCTCTACTTACCAGGGCATG |
| Alpi | AAACTAAGGTATTACCCATG |
| Alpi | AAGGCCAACTACAAGACCAT |
| Alpi | AGCCGGCACCTACGCACACA |
| Alpi | ACAGTAACCAGTCTGGAACC |
| Slc10a7 | CCGTCGGTCGGAGTGAACGG |
| Slc10a7 | GTTGAAGAATATCGTTGCGA |
| Slc10a7 | GAAGAAGCCACCATTTGGTG |
| Slc10a7 | AACTGCCTTGGTTAAAATCA |
| Nubpl | CCCAAAGCTCGGCACCCCCG |
| Nubpl | GTGGAGACAATCACAGCACC |
| Nubpl | TCAGCTGGACTATTTAGTTG |
| Nubpl | ACACCTTACAATACCTGCGG |
| Timmdc1 | ATCCGGATTCCAGGGAACCG |
| Timmdc1 | GGAAATTGACTATATCTACA |
| Timmdc1 | ATACCGGATGAAGCCCCTTG |
| Timmdc1 | CACCGGCAGCAAAGACTCGG |
| Ndufaf6 | AATTATGCTGAGAATACGCA |
| Ndufaf6 | ACACGTTCCCACGCATCCCG |
| Ndufaf6 | AACATAACCTAACTAAAAGG |
| Ndufaf6 | GCAGTTCTGGAAAAAAGCTG |
| Mrap | GAGAAAGAGGAGCACCACGA |
| Mrap | GTCCATGAACATATTGGCTG |
| Mrap | CAACCACAGGGCGATGACAA |
| Mrap | GGTGAGCGGGACAGAGGCGT |
| Mtmr2 | TGCTCGGCCAAGTGTCAATG |
| Mtmr2 | GCCTTCTGTACTCTAGAAGG |
| Mtmr2 | GTGGCATCCTTTAGGTCACG |
| Mtmr2 | GTAATGATTCTCTCATAACA |
| Agl | ACTGGTCTACCGGTACGGAG |
| Agl | AGAAATGGCCTTATCCGCAG |
| Agl | TGGTCATCACTTAAGTACGC |
| Agl | AACAGACTGGTGGATGCTCG |
| Vps33a | GCAAGAGGTTATCAAACACG |
| Vps33a | TAATTGCTGAGAATGTACTC |
| Vps33a | ACACAACGCTAAGACAGTCG |
| Vps33a | GCCTGTACCACGCAGCCAAG |
| Mboat7 | TGATCAGAGAATGCAAACTG |
| Mboat7 | ATAGCAGTCGATGTTACGGA |
| Mboat7 | AGCTATAGCTACTGTTACGT |
| Mboat7 | GCCTGGATTGCGGCCGAGTG |
| Mmab | CAGTGCATGCTACAGGACGT |
| Mmab | ATCAAGTATTTGAAGCCGTG |
| Mmab | GAAGCTCTTCGGCAAACATG |
| Mmab | TGTCCCCGTTTCCAGAGCCG |
| Rdh12 | GGAAAGGTTGACCACCCGTG |
| Rdh12 | CAATTTCCGCACTAGCACCT |
| Rdh12 | TGCTAGGAAGCGTTCAGCAA |
| Rdh12 | CCCAAAGATACTTACCCAGG |
| Sbf1 | ATTGTGCCCTCATCTAGCCG |
| Sbf1 | AGAGCAGGGCTACACTACAG |
| Sbf1 | ATTGTTGCTGATCTTGACGG |
| Sbf1 | TGTGTCGTGACAGGAGACTG |
| Mccc2 | TTACCAGTTATATGGCGACG |
| Mccc2 | GAGGAGCTTTGATGTCCGAG |
| Mccc2 | TGACTGTGAAAAAGCACGTG |
| Mccc2 | GGTGGTCGTCCAAAGCATAG |
| Cpt1c | CCACTCGGACAGGAGTATGT |
| Cpt1c | TCATACTGGGCGGAACACAA |
| Cpt1c | CATTTGCCAGCGCTGTAGCG |
| Cpt1c | TGGGAGAGTGAGTCCCTACA |
| Bola3 | GCGAATGTTTGCCTCTCAGA |
| Bola3 | GCTGCGCACAGTGCCACAGA |
| Bola3 | CCCGCGCAGCAGAGGCGCAG |
| Bola3 | GTATGAAATTAAAATCGAAT |
| Ak7 | TCTCCCTTAGGACGACAACG |
| Ak7 | GGAGGCGATTGTCGCACCCA |
| Ak7 | CTCCAAACACGGGTAGTGCG |
| Ak7 | GGAGAACTTTAACATCCGCT |
| Slc25a12 | ACCCGGATGCAAAACCAGCG |
| Slc25a12 | ATAGCACTTTAGCAGGCACG |
| Slc25a12 | CTCTTGATAGGAGATCAACC |
| Slc25a12 | TCACTGGAAGTCTTACCCTG |
| Pigt | GATCCGAATCCCAACGCGTG |
| Pigt | AAACAAGTCATAGACGGCAT |
| Pigt | GGTAACTGGTGTGGAACAAG |
| Pigt | TGAGAGTCCGGGAGAACATG |
| Mrps34 | TGGGTACCAAACGCCAATCG |
| Mrps34 | AGTCGGGCCTCACACGCGTG |
| Mrps34 | GCGGCTCTCGCGACGCACGT |
| Mrps34 | ACGCTGACCCGGCCGCACTC |
| Lrat | GAAATCAGCTCTTTCGTCCG |
| Lrat | TCCTAGTCAATCACCTAGAC |
| Lrat | CAAGGTGGCTAGCATCCGTG |
| Lrat | TCTAATGCCTGACATCCTGT |
| Lias | AGGTTGCCTTACCATGATCG |
| Lias | GCAGTCACCTCTGTAACTCG |
| Lias | ATAAACTGAAAAATACATTG |
| Lias | GGATTATGTTGTCCTGACGT |
| Slc19a3 | CTGTCGGAGTATCACACTGG |
| Slc19a3 | TGCCAATTCAAAGAATCTAG |
| Slc19a3 | ACCAGAGGGACCAGTAAACA |
| Slc19a3 | ACAACGTGTAGCATGATGAC |
| Ctns | AGGCATCATTACTGTCGACG |
| Ctns | TGGAGGGCCATGACACACGC |
| Ctns | TAACAGGTTATAGTGCCTCG |
| Ctns | CCCAAGGGTGATGTCGACGT |
| Elovl4 | TTCATGGGATCATACAACGC |
| Elovl4 | ACACCGTGGAGTTCTATCGC |
| Elovl4 | TGCAGTGGTGGTACACGTGA |
| Elovl4 | CGATACAAAATACCACCACA |
| Hamp | CACCTGTTGATGGAGATAGG |
| Hamp | AGATACCAATGCAGAAGAGA |
| Hamp | AGAGCTGCAGCCTTTGCACG |
| Hamp | AGAAAGCAGGGCAGACATTG |
| Cox4i2 | GGGCGTAGCAGTCAACGTAG |
| Cox4i2 | CCAGCGCTCCTATCCCATGC |
| Cox4i2 | GGACAGGACTCAGAACTAGA |
| Cox4i2 | CTTCTGCACAGAGCTCAGCG |
| Kars | CTCATCTTCTATGACCTGCG |
| Kars | GGAGCGTTGTGAAAGGAGCA |
| Kars | TGTATGTGATGATCTTAGAG |
| Kars | TGTGGCTATTGACTACCTTG |
| Acox2 | TAGGCCGAAGACGAGACTGG |
| Acox2 | GATGAGCAGATTGCTAAATG |
| Acox2 | CTTGAGTGACCAGTGTCGGT |
| Acox2 | ATGCCAAGATAATTGCTCTG |
| Echs1 | AGTCGGCAAATCGCTAGCAA |
| Echs1 | GACAGCCAGAAATCCTCCTG |
| Echs1 | GATGACCGGTTTCTTGACCC |
| Echs1 | GTGCATTGAGTGCTTTGGGG |
| Amn | GCAGACGTTCTCGCGCGACG |
| Amn | ACAGACGCAGCCCGACGCGT |
| Amn | CCAGAACCGGACCCCGTGCG |
| Amn | GCGCGATGGATCTTTCCGCG |
| Mrpl3 | GGAGACCAGTTGCTTAACAA |
| Mrpl3 | TGTGTTAAAATACACTCCCA |
| Mrpl3 | CCCACCTTCTCTCTAAGCCG |
| Mrpl3 | GAAAACCTTTACCAATACTG |
| Mcoln1 | GACATAGGCATACCGGCCCA |
| Mcoln1 | TTTGACAATAAAGCGCACAG |
| Mcoln1 | ATACCTTTGACATTGATCCA |
| Mcoln1 | TGGCCCGGAACTTGTCACAT |
| Nans | TATGTGACGTTCCAACACCT |
| Nans | TATACTTCGAAGCATTCATG |
| Nans | TCGTGCCCGGAATACCCGAT |
| Nans | GAGATCACCATAGGACGACC |
| Cnnm2 | ACCGTCGTGGTAAATCCAGG |
| Cnnm2 | CGGCGATTGAGAATGATGTG |
| Cnnm2 | TGCTAACCGGATAAGAAGCG |
| Cnnm2 | CTGCTTCATGATAACCGGCG |
| Srd5a2 | CCCAGCAGCACATTCCCCGG |
| Srd5a2 | ACAGACATGCGGTTTAGCGT |
| Srd5a2 | CATCCCTACCGACACCACAA |
| Srd5a2 | GGTATTCCGCGCAATAAACC |
| Sfxn4 | TTAAGCCAAAAGTTATTCGG |
| Sfxn4 | AGCACATGAGACATACCACT |
| Sfxn4 | TGCAGTCAGGAACTCACCAT |
| Sfxn4 | CCATCGAGGAATTGAGGTGA |
| Hadha | ACACTTTGTGCTTTACACCG |
| Hadha | TATTAATTATGGCGTCAAAG |
| Hadha | TTAAAGACACCACAGTGACG |
| Hadha | AAGGACAATTGAATACCTAG |
| Cog8 | CTACAAGACGTTCATTCGCG |
| Cog8 | ACACACGTGTCCATAAGCTG |
| Cog8 | TGGCGTGAGAAGCCCGACGT |
| Cog8 | AGGTGGCTACCTGGATAACG |
| B3galnt2 | TAGTGGCTAAATTTCCAGTG |
| B3galnt2 | ACACCTTCCATGAATTCATG |
| B3galnt2 | ACCCACCGTTGACTTAATGT |
| B3galnt2 | GGAACAGTTCATCTTACCAG |
| Kmo | GCTGTCGAAATGCATTCCGG |
| Kmo | GATGTGTACGAAGCTAGGGA |
| Kmo | GACAATTCCACCTAAGAATG |
| Kmo | TTGCCCAAAAAATGGGACAA |
| D2hgdh | GGCTCACAGAGCATACTCCG |
| D2hgdh | AGGCCTGGGTGGGCACACAA |
| D2hgdh | GGATACTGAATGCATGCAGT |
| D2hgdh | GACTGGCTGAAGACCGTCCG |
| Lbr | TTCAAGTGACCACTCCACAG |
| Lbr | GCGCCCTGGATTGATTGGAT |
| Lbr | GAAGTAAGTACCAACACAAG |
| Lbr | GAGCTTGAACGAAAAACCAG |
| Pomt1 | TGTGAACAGAGAGTCCAACC |
| Pomt1 | ATAGTCGGGTAAGTAGTCCC |
| Pomt1 | GGGGCACTCACGTGTTAAGG |
| Pomt1 | GTCCACGCGTGGAATCTGAT |
| Dpyd | CTATAACCTTCCGTGGCGAG |
| Dpyd | GAGGCACAGCTTATACTCGC |
| Dpyd | TTTGTCCAGGGCCATACAAG |
| Dpyd | AGAGCTAAGAGTCATTTCAT |
| Ldlrap1 | GCTCCACCAGCGTCATACCA |
| Ldlrap1 | TCGGTCAGGATGATCCCCCG |
| Ldlrap1 | TGCAGACAAGATGCACGACA |
| Ldlrap1 | CAGAGAACTGGACGGACACG |
| Pcsk9 | GCTGCCAGGAACCTACATTG |
| Pcsk9 | TGGAGCGAATTATCCCAGCA |
| Pcsk9 | AGGACCAGCCAGTTACCTTG |
| Pcsk9 | CATGCTTCATGTCACAGAGT |
| H6pd | TAGGCGTAGTAATCAGACAG |
| H6pd | CTTGAAGGAGACCATAGATG |
| H6pd | AGTTCCACGCCCATACCCAG |
| H6pd | ATGTCTGCGTAGGCAAAGGG |
| Psph | TTCGTACATTTCAGACACTG |
| Psph | CTTATGCCAGGAGTCAGATG |
| Psph | CCTGGACATTACGCTCCTGG |
| Psph | CAGCCTATTGGCAAACACAT |
| Acacb | AGAACAACGATATCGACACG |
| Acacb | CCAGTGTGCCGACCACATGG |
| Acacb | GTGGCCGTACATGTCGATGG |
| Acacb | TTCATTACGGAACATCTCGT |
| Eogt | GTCGACATAGCTGCAAACGG |
| Eogt | GAGGCTGGAGTAGTCATACA |
| Eogt | ACATCAAGAGAAACCACGAC |
| Eogt | AACAGCCCGTACCGCATGCG |
| Hsd3b7 | CCTTCACCATAAATGCCCGT |
| Hsd3b7 | TGCTTGGATGCACATACTGG |
| Hsd3b7 | ATCCACAAAGTCAACGTGCA |
| Hsd3b7 | ATGTGGTCATCCATACAGCT |
| Phkb | TCGGCCATAAAGTGTATGAG |
| Phkb | AAGCGGTTTCCAAGTAACTG |
| Phkb | CCGTGTTATAGATTATCTGG |
| Phkb | AACGCAGGGTAACTGATGCA |
| Cog4 | GGAAACTCACCAACCCATCG |
| Cog4 | GCTGTGTACGAACGACTCTG |
| Cog4 | GCACAAGTAACGATGAATGT |
| Cog4 | CAACAGAAAAAATTGAACCA |
| Lars2 | GGTTAGCCATGATAACGATG |
| Lars2 | TCTTAATGAACCACTGTCGG |
| Lars2 | CAAATGCATCCCACCCCATG |
| Lars2 | CATCTATGGCATCTCCCACG |
| Alg9 | ATGAGCGGATGGCATACACG |
| Alg9 | ATACAAAACAATGTTGAGAG |
| Alg9 | TCCCGTCATGGCGATCAGCG |
| Alg9 | TCAAGTGTCTGCTCTCAGCG |
| Slc6a8 | GGAAGCGCCACACGTTACCG |
| Slc6a8 | GCATCAGTGTGACAGCCCGT |
| Slc6a8 | GCACAACGAGGACCACGTAG |
| Slc6a8 | ACTCGATGACAGGGGACCGG |
| Upb1 | TCTCGTGTACAAAAAGCGAA |
| Upb1 | CTGTACATGAGCCAGTTGAG |
| Upb1 | GTAGAGAATTCGCTTCACCT |
| Upb1 | ACTGAAGGGATATGCCTTCG |
| Fig4 | CTTGGTGGTAATCGACGACA |
| Fig4 | AAATCCGAACAGAGTCATTG |
| Fig4 | GAAGGATTAATTACACAGGG |
| Fig4 | CACAGGTGGAATGAACTAGG |
| Pnpo | AATGCGCAAGAGTTACCGCG |
| Pnpo | ACTAACTACGAGAGCCGGAA |
| Pnpo | GGTAGCCACACACATAGCAT |
| Pnpo | CTGAACAGCCTCGTCAAACC |
| Pex12 | TTTCATCAAACCACCGCCAG |
| Pex12 | TTATGGCTTAAAGCGGATTG |
| Pex12 | GGAACATGGCAGATTTCCAG |
| Pex12 | GGCCATGTTAACAAAAGGGT |
| Plin1 | ACAGCCCTGCCCAACCCGAG |
| Plin1 | GTCATGGCTCTCATCCTGAG |
| Plin1 | ATACCTGACTCCTTGTCTGG |
| Plin1 | GGTGTGTCGAGAAAGAGTGT |
| Gck | TTCTGGGGTGGAACGCACGT |
| Gck | AAGGCACGAAGACATAGACA |
| Gck | CCATCCGGTCGTACTCCAGG |
| Gck | CCAGATGTATTCCATCCCCG |
| Synj1 | TATGTCAATCAAGGCAACGA |
| Synj1 | GGGACAAGGTTCAATGTCCG |
| Synj1 | TAAAATATATGCAGGCACCG |
| Synj1 | GAAGAGCATCCGAATTGCAA |
| Cyp27a1 | CTACACGGATGCCTTAAACG |
| Cyp27a1 | AGACCAAGTATGGTCCAATG |
| Cyp27a1 | GAGGACGCGTCCATTTGGGA |
| Cyp27a1 | GGTCCTTCCACTGATCCATG |
| Ndufb11 | TGTAATCGCCCCATCCGGTG |
| Ndufb11 | GGACGAAAACGTCTACGCGA |
| Ndufb11 | TATTCCAGACGTCCACCACA |
| Ndufb11 | GTCGCTGCCTGTCAGCAGCG |
| Gldc | CCACTGAAATCCATTTCGTG |
| Gldc | TTTACTCAACTACCAGACCA |
| Gldc | CTTGATCCAGGGATGCCACA |
| Gldc | AGGCCACAAACAGCTACCAG |
| Slc6a5 | GGGATGGTCACCGATAACAC |
| Slc6a5 | GTGTACGCATCACTGGCGAA |
| Slc6a5 | GCTTACCGGTATGGTAGTGG |
| Slc6a5 | GAGGACGCGAACGTGAGTGT |
| Aldh6a1 | TAGATGTGTCTGCCCACGCA |
| Aldh6a1 | TAAGTTACTTACCACTGATG |
| Aldh6a1 | TTCAACAGCCATCCTCGTAG |
| Aldh6a1 | AATGTTCAGTGTTCCGTCAG |
| Tecpr2 | AGCAGACATCCTGACCCATG |
| Tecpr2 | GTGGCAAAGTCACCATCAAG |
| Tecpr2 | TCTCGCCTGAGAAACACCTG |
| Tecpr2 | AGCAGATTGGATCCCAACCA |
| Nhlrc1 | GCACGACGTGAAGTACCCAC |
| Nhlrc1 | ACTGTGCCCCAAGACCGGAC |
| Nhlrc1 | GGAGATCACCCCTCACAATG |
| Nhlrc1 | TCCCTGAAGGGGTCCTTCGG |
| Tecr | TACGTCCAGTCGGCATACGG |
| Tecr | ACTCTACTTCCGGGACCTCG |
| Tecr | TGTCTGGACCCACTTACTAG |
| Tecr | ACTTTGAAGATGTTTCGCAA |
| Ppcs | ATAGAAATCTGACACTGCCG |
| Ppcs | ATGCAGATAATCCGCCAAAG |
| Ppcs | AACTTCAGTAGCGGGCGACG |
| Ppcs | GTTCTTGTACCGAGCGCGCT |
| Uqcc3 | AGATGCCGGAAAGGGACTCG |
| Uqcc3 | GGACGAAGCAGTCAGGACCA |
| Uqcc3 | GGTGGCTCGTAAAGCACTTG |
| Uqcc3 | GCAGTTGCAGTGCTAGGCGG |
| Psat1 | TGGAAGGAGTGCTGACTACG |
| Psat1 | TGCAAACGAGACTGTGCACG |
| Psat1 | CAATACAGAGAATCTTGTGA |
| Psat1 | CTTTGTAGTCAAGGACTGAT |
| Acaca | GCCATTCATTATCACTACGT |
| Acaca | AATGCATGCGATCTATCCGT |
| Acaca | TTGATTCATAGGTACCGAAG |
| Acaca | AAGCCCTTCGAACATACACC |
| Haao | TGGGTGCTAACTGCGTGCCA |
| Haao | AGAGGTTTGCCAACACCATG |
| Haao | GCAAACCGGAGGCTGAAAGG |
| Haao | CGGGACGTGCCTATACGGCA |
| Cth | TGCATGGATGAAGTGTATGG |
| Cth | GCAATGGAATTCTCGTGCCG |
| Cth | TTCCAAAACCAAATTGCTAG |
| Cth | TTTGTGGACAATTTGTGCGC |
| Mthfs | TTGCTCTGGAATTGGTACCG |
| Mthfs | CTTCTCGCACCTTCTGCGTG |
| Mthfs | GATGAGGTCAAGTCCACCTG |
| Mthfs | ATCTTGCATGCTTAGAAAGA |
| Grm6 | ATTCTCATTGAACATCACTG |
| Grm6 | GGAGTCAGGAGGCACCACTC |
| Grm6 | CCGGTGTGCGACTAGGCGCG |
| Grm6 | GTGGTCAGGCGACCCCCATG |
| Prkag2 | GGGACGAGGAAGCATAGATG |
| Prkag2 | AAGAACATAAGATTGAAACG |
| Prkag2 | CCACTGAAGCTCATCCGGCG |
| Prkag2 | GCGTTTATATGCGATTCATG |
| Atic | AGACGACTGTCACTCGAGCG |
| Atic | GAGATGTGTCTGAGCTAACA |
| Atic | AATTGCTTCATCGTATTGAG |
| Atic | TTCTTTCAAGCACGTCAGCC |
| Ogt | TTTGAGCCCAAATCATGCGG |
| Ogt | AGCATTATCGACATGCCTTG |
| Ogt | ACAGTGCACCACTAGTCCCA |
| Ogt | GAAAGTTTGTACCATCATCC |
| Mthfd1 | ACACCAACGATAGATTCCTG |
| Mthfd1 | CACTATGAATCCGTGCACAG |
| Mthfd1 | GATTGCCGGAAGGCACGCGG |
| Mthfd1 | GGTAGCGTCCAGTAAGAAAG |
| Zdhhc15 | TGGAACTTAGAGCGAACACT |
| Zdhhc15 | GAATCAAAAGGGCATACCAT |
| Zdhhc15 | GTGTACACAAGAACCGGGAG |
| Zdhhc15 | AGACTGTTGTAGCAATGTAC |
| Atad3a | AAGCAGGAGGCCATACGGCG |
| Atad3a | TTACCTGATAGACTCCAAGA |
| Atad3a | TGCCACATCAACTCACACTG |
| Atad3a | GCTGTCGAGCGAGCTTATCC |
| B4gat1 | CCGGGATTAATTATGCACTA |
| B4gat1 | GTAGACGCGATAATCGCCGC |
| B4gat1 | TTTACCTGGTCTACGGATCG |
| B4gat1 | CGAGGGGCACACGAGGTGCA |
| Rars2 | AAACCTGGTGGCATAGAGAG |
| Rars2 | TGTCACCCATTTCTAGTCGG |
| Rars2 | TCAGCTTAAGTGTGATACAG |
| Rars2 | ATTATCTTGATAGGTCTGCT |
| Slc30a9 | AAGGACTCGGTGTCATCGTG |
| Slc30a9 | ACTGAGACCGCTCTGGAACG |
| Slc30a9 | ATTATCTGATACTTGTAACC |
| Slc30a9 | CTACGAATGTCCAGAAAGGA |
| Mmadhc | TCTGTGAGGCAGTCCCATTG |
| Mmadhc | GTGTGGCCTGATGAAACTAT |
| Mmadhc | TCACTAAAGAACAAAATCCT |
| Mmadhc | TATGTAAATGAGTTTCAGGT |
| Mmaa | CGACCTACTCGAAATGTGAG |
| Mmaa | AGGACAGAGAGCTTGCCTAG |
| Mmaa | TCAGGCCCTCTCCTACCAGT |
| Mmaa | ACTTGAGTGAAGGAGCCATG |
| Trip11 | TACTGATCATAAACGAACCA |
| Trip11 | TTATTGAGTCAAGAAAAGGT |
| Trip11 | ATCCTGTGAGTAGGACGCGG |
| Trip11 | GAAGGAGAACCATGAACTGA |
| Acy1 | AATCTGACCAAGCTTGAAGG |
| Acy1 | TGTATCCTCAATGAAGCGTG |
| Acy1 | GACAGGTACCACATCTGTGT |
| Acy1 | AAAGACAGTAAAGGCATCCG |
| Cyb5a | GTCCAAAACATACATCATCG |
| Cyb5a | TCTGACCAAGTTTCTCGAAG |
| Cyb5a | CTTATGATGCAGGATCACCC |
| Cyb5a | GTCCTAAGAGAGCAAGCTGG |
| Ampd2 | CACTGCCTACGACTTAAGTG |
| Ampd2 | CAAATGCAAGGAGATCGCTG |
| Ampd2 | TCATAAGTTCGAGTCTCCAG |
| Ampd2 | ATATTGTAGAAGTCCCGGTG |
| Cyb5r3 | GTCATCATCGCTCGACACAG |
| Cyb5r3 | ACTTCACCGTCCTGACAACG |
| Cyb5r3 | GGAGACACCATTGAATTCCG |
| Cyb5r3 | AGGTCATCAGTCCTGACACT |
| Blvra | TCTGAAACAGACGAGAACTT |
| Blvra | TAAGAGAAAGCTCCCCAAAG |
| Blvra | ATGCCATGGGGTATTCCACG |
| Blvra | TATAGGCGACATCAACCTCT |
| Pgm3 | TACGGCCTCACATAACCCTG |
| Pgm3 | AACTTCTTCAAGGTACCGCG |
| Pgm3 | TACACCATGTAGTGCAACTG |
| Pgm3 | CTGCTATGACATACCCTGTG |
| Asl | CTCGTGTCAGCGCAACAGCG |
| Asl | TGGTTCCAATGAGCACCCGG |
| Asl | AGATGCAGCAGATACTGCAA |
| Asl | GCTGCAGGGAAGCTACACAC |
| Gusb | GTATCTTGGGTCCATCGCCG |
| Gusb | TGTTCGAATCCCGATAGGAA |
| Gusb | GGAGCGCGCACCTCCCGTAG |
| Gusb | GGTGTCAGTCTTGTAGACGA |
| Phka2 | GGGTCATAATGCAATCGGTG |
| Phka2 | ACAGTAGACCGAGTTCCACT |
| Phka2 | CACAGAAAGTATGTCCCGAG |
| Phka2 | CAAACAAATTGGTATCCCAG |
| Pygl | AAACCAACGGGATTACCCCG |
| Pygl | GGGCGAAGTAGTAGTCGCGG |
| Pygl | CTTACAAAATGCCTGCGATG |
| Pygl | AGGAGGCAAACGGATCAACA |
| Cyp11b1 | TTGCTATCCCATCCACCAAG |
| Cyp11b1 | ATAGTCCATAGAGAACTCCG |
| Cyp11b1 | CAAAGAAAGTCATTACCAAG |
| Cyp11b1 | GGCTCAGATCACGGCCAATG |
| Mpi | TGGATGGGGACACACCCCCG |
| Mpi | CGAAACATCCGATATCACCT |
| Mpi | AGGGTGTGCCTGGATAGACA |
| Mpi | CAGGTAGGCATGGGGTACGT |
| Manba | GTCTGGTTCCCATGTCCACG |
| Manba | AACCAACGGGATGAAAACCA |
| Manba | GTTCACGTCTTTGACTACGT |
| Manba | TTGAAGTAGAAACTCAGACC |
| Fdps | AGATACACTGAAAAGAGGTG |
| Fdps | CCAACTCACCATGTACATGG |
| Fdps | ACAGATCTGCTGGTATCAGA |
| Fdps | TTGCCCGGCTCAAGGAGGTG |
| Cox6b1 | AAGGCAATGACGGCCAAGGG |
| Cox6b1 | AACTGTTGGCAGAACTACCT |
| Cox6b1 | AGAACCAGACTAAGAACTGT |
| Cox6b1 | ACGCCGGTACCACTCACACA |
| Qdpr | TGTCATCTTAACAACCACGC |
| Qdpr | CTTACCAGGAGTCCCATCCA |
| Qdpr | GGTGGATGCAATTCTCTGTG |
| Qdpr | CACGCAGTGTTACCCAGTTG |
| Pigh | CTTGTCCATCTCTATGAAGG |
| Pigh | GGTCAATCTTCACAAAATGG |
| Pigh | TGATGAAGATGGTGGCAGAG |
| Pigh | TGAGTAGTAGCGGCACCGCA |
| Acat1 | GTCCCATACGTAATGAGCAG |
| Acat1 | AGTGCTATAAGAACTCCCAT |
| Acat1 | AGGCAAGCAACACTGGGCGC |
| Acat1 | GCCTCTCAAAGTCTTATGTG |
| Acat2 | CAAACCTGATGCGTTCGCTG |
| Acat2 | CTGAGAACAGGAGTCAGGAT |
| Acat2 | TCCTACTCGACAAGCCAGTG |
| Acat2 | TGCCTTTCACAACTACCACA |
| Aldh7a1 | GGGCTCCGCGAGGATAACGA |
| Aldh7a1 | TGGTGGAGCAGATATCGGGT |
| Aldh7a1 | ACACTAACGAGGGACGTAGT |
| Aldh7a1 | TAGATTCCTGCCCCAAAACG |
| Nr3c2 | GATGATTGGGCTCTTAACGA |
| Nr3c2 | CAAGGAACTTTCAGCCACGG |
| Nr3c2 | TCCACTAATGCATTGACAGG |
| Nr3c2 | GAATGGAAATGGGTTGACTG |
| Pcca | CTGGACAATTATGTGATCCG |
| Pcca | CCTGGATTTCTATATGACGA |
| Pcca | GTAAACCGAAAGACTTGTAG |
| Pcca | TTAGGTTCACGTGAAAATGG |
| Etfb | CATCCACGTGGAGATACCAG |
| Etfb | CCTGAGAGGCGAATGTACCC |
| Etfb | AAAGCCGGACAAGTCTGGAG |
| Etfb | CATGGCCAGAGCAGTTCGGA |
| Etfa | TGGCGTAGCAAAGGTTCTGG |
| Etfa | TTTGTGAGAACTATCTATGC |
| Etfa | GGTAGTGCTAGTTCAGAAAA |
| Etfa | AGTTACACACACATCTGTGC |
| Hlcs | CAGCAATGTGAGAAGACACG |
| Hlcs | ACAGTACCATTGACTCGGTG |
| Hlcs | ATGGCCTATCTTTCTCAGGG |
| Hlcs | CGGCGGCCTTGCAATCACCT |
| Cldn16 | GGCCACGATCAAAAACCCAG |
| Cldn16 | GAACGCTGATGACTCCCTGG |
| Cldn16 | AGTACCTGCAATGAGTAATG |
| Cldn16 | TATATTGTGTAAAACGAGAG |
| Slc13a3 | AGTTTCTTGCCAGTTCGGAA |
| Slc13a3 | TCCAGAGCCAGCCCACCAGT |
| Slc13a3 | GGCTAAAGCGGTGATCCAGG |
| Slc13a3 | CCACCGTACCTTGGGCGGCA |
| Ddhd1 | TCGTTGCGCTACTACAGCGA |
| Ddhd1 | TCAAAGCTTACTCTTGATGG |
| Ddhd1 | GATAAAATACCAGTAATGCG |
| Ddhd1 | ATAAGTGATTACACACCCCA |
| Slc19a2 | GGAGTAGTAGGCGATTTCGG |
| Slc19a2 | GACTCAGCCTGATTGTGACG |
| Slc19a2 | CAAGTGGTGAACTACGCGCA |
| Slc19a2 | GCAGGAACCAGCACTCGCGA |
| Slc2a9 | CGGAGAGGTTGTACCCGTAG |
| Slc2a9 | GCATGCCATCAGCAACGCTG |
| Slc2a9 | CCAACTATGTGGACTCAATG |
| Slc2a9 | TCTCAACGAGATCTCACCCA |
| B3galt6 | GCGCGCCAGGTACAACAGAG |
| B3galt6 | CCTGGTGGACCTACGCGCAC |
| B3galt6 | GTTCTCGTAGGCGTCGCGCA |
| B3galt6 | TGGCGGACCCGAGGACGTGT |
| Mrps2 | CTTGCGGTAGGCCACGTGCG |
| Mrps2 | CCTGTGCCAACTACCTATGG |
| Mrps2 | CACACCCGCTACTTCAAGGG |
| Mrps2 | GCGTATCTAGGATCCTGCTC |
| Pofut1 | TGTTGGTATTCAATCCATGG |
| Pofut1 | GCGCCTCCTACAAAGAACAA |
| Pofut1 | CCGGAGGCCTTACCCATGCA |
| Pofut1 | AGCTCCAGAAGTACATGGTG |
| Slc2a10 | CAGCCCCGTGGCGACCAAGG |
| Slc2a10 | AGAGGCGTAATACAGCACAT |
| Slc2a10 | TGTGCTGGTGTCCCTCTACG |
| Slc2a10 | AGCCAACAGATAAACGGCCC |
| Bbox1 | GTGGTGAGATCATAAAACTT |
| Bbox1 | TACACAGAATGTCAGTACTG |
| Bbox1 | GTTGAAGCTCACCTTACCCC |
| Bbox1 | CAAGATTGATGCCAACAATG |
| Idh3b | TTCACCCTTATACTCCATTG |
| Idh3b | TCGGCACAACAATCTAGACC |
| Idh3b | ACAGGCACAAGATGTGAGGG |
| Idh3b | TAAGGAGCATCATCTGAGCG |
| Rtn4ip1 | ATATTTATCTATCACCCAAG |
| Rtn4ip1 | TGAGGTCATTGATTACACGT |
| Rtn4ip1 | GCAGGTAATGAAAGCATGGG |
| Rtn4ip1 | GAGAGCCACATATGGCAAAG |
| Mfn2 | CCGAGGCAGACGCATCCCAG |
| Mfn2 | GTGGTATGACCAATCCCAGA |
| Mfn2 | ATAAACCTTGAGGACAACTG |
| Mfn2 | AGAACTGGACCCGGTTACCA |
| Sugct | AAGATCTCCTAAATTCATCG |
| Sugct | GGTGATGACACACGATCTTG |
| Sugct | AGGAGTGAGAATCGTCAAAG |
| Sugct | AATCTAAGGATACCTACCTC |
| Mvd | CCCGCCGAGCCACTTCGGAG |
| Mvd | CATCTGGCTGAATGGTCGCG |
| Mvd | CCACGTGCACCTTATAGCTG |
| Mvd | TGTTGTGCCTGAGCGCATGA |
| Sardh | CCTTGACTACGACTACTACG |
| Sardh | CGTCGTCACTGAGCGCATCG |
| Sardh | AACCGAGTAGTTCTTGGCAT |
| Sardh | CGGTGACTGGCATCCGTGTG |
| Klf11 | CAGTGTGATCCGTCACACCG |
| Klf11 | AGCTCACGGAGCCGTCAACA |
| Klf11 | ACCAGCATCACAGTTCCCTG |
| Klf11 | CCTTGCTTGCACAAGTCCGG |
| Tmem199 | TAGCCAATGAGGAGTATAAG |
| Tmem199 | CGGAATGAAACCAGGCGCCG |
| Tmem199 | TTTCTGTTACATACCCGGGG |
| Tmem199 | TGCTCTGGGAAAGAAACACG |
| Alg11 | GGAAACCAGCAAATGCCCTG |
| Alg11 | TTTCCTGGACTGATGATATG |
| Alg11 | AGTGAACATAGCTTCCGACT |
| Alg11 | TAAACCATATCCTCTCACTG |
| Alg1 | GTGAAGTTGACAGATCTGCG |
| Alg1 | TCTGGAAAGGAGAGTACGTG |
| Alg1 | TGTGACCAATGCTATGCGGG |
| Alg1 | TGAGAGGGATTGTCAGAGCG |
| Alg3 | AAGAAGGAATAGCAATCCCG |
| Alg3 | GGAAAAGAAACTGACGACCA |
| Alg3 | GCCACCAGCAATGTGTAGCG |
| Alg3 | ATTGATGAAACCCTCCACCT |
| Slc25a38 | TCCAGGGACACATCTCACAA |
| Slc25a38 | CTGAGAAGGGGGCATCACGG |
| Slc25a38 | AGAGGAAAGCCTTAATCACT |
| Slc25a38 | AGTGATCAAGACACGCTATG |
| Akr1d1 | TTGTCTGGGTGAAATACGGG |
| Akr1d1 | TGAAGACAGCTATTGATGAG |
| Akr1d1 | AACAAATCTGTGTGCCACGT |
| Akr1d1 | GGGCTGGGAGGACCATTGAT |
| Zdhhc9 | TGGAAATTCTTAATACGAGG |
| Zdhhc9 | GTGCTCGAGGAATCACTCCA |
| Zdhhc9 | AGAAGAGTGTACATGTCCCC |
| Zdhhc9 | AAGAGCATAGCGGCAAACAC |
| Pycr1 | CCGGGGTGTTAGTCATACAT |
| Pycr1 | TTGGGGCGAACATTGAGGAC |
| Pycr1 | CACAGGTACTCACGCTCAGG |
| Pycr1 | GGAGGGCCGAGACCGTAGCT |
| Dhtkd1 | CTGATGTTCCGTAAAATGCG |
| Dhtkd1 | GCCACGGTGAAGAGATACGG |
| Dhtkd1 | TCTTCTTTAGGAAAGCTCGT |
| Dhtkd1 | GATGGGGACTACTCCCCGAA |
| Mtrr | GATGAAGCGGTTGTAATCAG |
| Mtrr | GGTGAGCAAAATAGTCCGTG |
| Mtrr | GAACAAAATGAGACAAACAG |
| Mtrr | CTCGACCGTACTCATGTGCA |
| Sepsecs | GAGGACGTCTAAGGACGGCG |
| Sepsecs | TAAGCGTTGTTGACTACATG |
| Sepsecs | CAGGACTTCTGGTCTATCCG |
| Sepsecs | CACCGGATCGTCCAATTCCG |
| Cln5 | GACAAAATTCGAACAGTCGT |
| Cln5 | CCAGCGTTGCCCAGACGTCG |
| Cln5 | GCCCCTTGGTTACACCAGAA |
| Cln5 | ATATTTAAATTCCCAAATCG |
| Suox | AGGGGGACATGCTATCTTCG |
| Suox | TCCTCTTTAGAGTACATCCG |
| Suox | AAACTTATGCAAATCATCCA |
| Suox | CAGTCCTTCAAAACAGACAT |
| Pde12 | AGCTGTCGGAGAACACCGCG |
| Pde12 | AGCTTGGATCGAGACCGGTG |
| Pde12 | TCTTGGTGTACAGATGTCGA |
| Pde12 | AAGAACCGAGCCCACTCGAG |
| Zfyve26 | GAAGTCCTCACCTATCGCCG |
| Zfyve26 | CCTTCCCGAGGACTATGCCG |
| Zfyve26 | CTGAATCCTGATGGTATGCG |
| Zfyve26 | GGCCCTGGACATACTAACTG |
| Aldh4a1 | GTTCAACGCAAAGTTCGCCG |
| Aldh4a1 | CGCTCGGCATTCGAGTACGG |
| Aldh4a1 | CAACTGGTACTGTATATCGG |
| Aldh4a1 | ACCTTTATGACAGGGCAACG |
| Mars2 | ACTCGCAGACCATCTACGTG |
| Mars2 | GAGTGGACATAGATGCGATG |
| Mars2 | TCGATCGCACAGCTCGATCG |
| Mars2 | TGTCCCGTGTCTCTCGAGAG |
| Dse | GTCAAAGTATAAGCATGACC |
| Dse | CAGCACACAGAACATTGCCA |
| Dse | AGTTTCATACATATAGCCCG |
| Dse | TAATGAACGGCACACCATTG |
| Kctd7 | CGGTACTTCATCGATCGAGA |
| Kctd7 | CCGCAGAGTAGACAAGCGCG |
| Kctd7 | GTAATGCCGCCCGCTGAACA |
| Kctd7 | GAGAACATGCAGCCACTGAA |
| Slc39a14 | GAGCGAGCGATCTCAGATCG |
| Slc39a14 | GTAGAGGGTTCCAATCGCCA |
| Slc39a14 | TAAAATGGTTATGCCCGTGA |
| Slc39a14 | GTGACCGAGAAGCTACAGAA |
| Slc18a2 | CGAGCCATACGTACCTACGA |
| Slc18a2 | CCATCTGCTTTGCAAACATG |
| Slc18a2 | GTACATACCTAAGACCCCCA |
| Slc18a2 | ATGCAGAATCCAGCAAACAT |
| Gnptg | GGCCTACCTAACAAAGAGTG |
| Gnptg | CTGTGGAAAGATCAACCGAC |
| Gnptg | TTACCCAAGGATCCCGCTGT |
| Gnptg | ACAATACCTTCAAGGGCATG |
| Aldh5a1 | CCTCTGCCAAAGATAAGCGA |
| Aldh5a1 | ACTGTCAAAGACAATGAATG |
| Aldh5a1 | ACACCAATGGATCGGTACAC |
| Aldh5a1 | AGACACACCCTATTCCGCCC |
| Spg11 | TCCCCGGAAACACAACGCGT |
| Spg11 | ATTCCCGGGGATGTCCCACG |
| Spg11 | TCATTCCGCTCAACGACTGG |
| Spg11 | AGAGCTACAAATCCCGTGCA |
| Pomgnt2 | AAAGCCGTATTGGTACCAGG |
| Pomgnt2 | TCAACCACGGTAGCCCCGCG |
| Pomgnt2 | GTGGCTCTGCTACTCCAATG |
| Pomgnt2 | AGACGTGCATGAGGTTATCG |
| Micu1 | ATGGGATCTCCACTAGACCG |
| Micu1 | TTCTTGCTGTTAACGCATGG |
| Micu1 | TTGAAAGTAATCAACGAACC |
| Micu1 | CCGAACATAAGCCAGACTTG |
| Iba57 | CAGACTCCCAGAGTGTACGG |
| Iba57 | GAAGATCACAGACTCCCTCG |
| Iba57 | ACGGGCTAGGCAGTGACTCG |
| Iba57 | ACCCCCGGACTGCACGTATG |
| Ptrh2 | CTCAAAGAGTGGGAGTACTG |
| Ptrh2 | ACTGACCTTAAAATGGGGAA |
| Ptrh2 | CCCCCAGAACTCAACAAGTG |
| Ptrh2 | GTTGCTTGTGGCATGTGCCT |
| Xylt2 | GTATACAGATGACCCCCTCG |
| Xylt2 | GTGCCACTGACTATCCAACG |
| Xylt2 | TGTAGGCGATGCGTACTGGA |
| Xylt2 | GGACGTCGCTGAGTCCGCCA |
| Nags | GGATGAACTAAGGCACAACG |
| Nags | CAGCTACGGTGGCATCGTCG |
| Nags | GACAGCCAGAAGGTGCCGTG |
| Nags | GCTAGCGGCTGTAATGACTG |
| Mgat2 | GAAGTTCAACTGGTACACCA |
| Mgat2 | TGTAATGGCCGAAGGAGTCA |
| Mgat2 | AAAAGTCTGGGGCTAAGTAG |
| Mgat2 | CTGGACGCGGAGCCCGTACG |
| L2hgdh | CGGACGCCGGTCCACTTGCG |
| L2hgdh | AACAGCGGTGTCATACACAG |
| L2hgdh | GCTATTGATTGTCCATACAC |
| L2hgdh | GACCGTATTTCAGAGTTGAG |
| Coq6 | AGACACCGTGTACGACGTGG |
| Coq6 | GGCCTTGATAATGTTCGACA |
| Coq6 | CTAGGGTAATATGGACCCAA |
| Coq6 | GTGTGATGTGGACCAGACCA |
| Pomt2 | TGGACAGATCCAGTACGGTG |
| Pomt2 | GTAGTCCTTATGCAAATAGG |
| Pomt2 | CCTGCACAGTCACTATCATG |
| Pomt2 | TCTTTATCATCGTGCAAGTG |
| B4galt7 | CACTACAAGACCTATGTGGG |
| B4galt7 | CTACATCGCCATGCACGATG |
| B4galt7 | GGAAGATGGAGCATTTCCGG |
| B4galt7 | TCGGGCAGCACTCATCAATG |
| Alg12 | GTACCAAAGAAGACTGACGG |
| Alg12 | AAGCGAGAGCACATAAACCA |
| Alg12 | TGTAGGGACATATAACCAGG |
| Alg12 | CTGAACAAAAGTTCCAACTG |
| Poglut1 | ATGATATCATGTATCCTGCG |
| Poglut1 | ATTTCTTGACCAAATTAACA |
| Poglut1 | GGACCCCAGAGGATACCTGG |
| Poglut1 | GAGGACCTGACTCCTTTCCG |
| Aars2 | AGGACGCCATAACGACCTGG |
| Aars2 | CTACACTCTTCACCCCAACG |
| Aars2 | GATAGACACAGCGTACCGAG |
| Aars2 | GAGGCTACCTTGTCCGTACA |
| Pex6 | TGTAACTCTCGAGCAAACGG |
| Pex6 | AGGAAATGCCGTCTGCACAT |
| Pex6 | GCACGGGGAGGGTCTCCCCG |
| Pex6 | AGCACACAGCTCATTTACCA |
| Ndufs8 | ATGTGGACAGAACTCATCCG |
| Ndufs8 | TCAGTGGGCCCTTCTCAAAG |
| Ndufs8 | AAGCCTTCATAGCAGCGCAG |
| Ndufs8 | CCGAGCTGCATTGTCAGTTG |
| Trpm6 | AGAGTGCATACAGTTTAGCG |
| Trpm6 | TGTCAACAGTTATGGCCGAG |
| Trpm6 | AGGTCATGATGTAGCGATAG |
| Trpm6 | TATTCTATAGCTCGAGAGAA |
| Cox15 | AGAAAGGGTTGGCTCAACCG |
| Cox15 | AGGCGGTACTGACTGACCCG |
| Cox15 | GGTACATGGAATACTCACAC |
| Cox15 | TAGGGTGGCGTCCAGAACAC |
| Lct | TGAGTTGTTCGATAGCTATG |
| Lct | TCAGCCTATCAAATCGAAGG |
| Lct | CTCCCGTGGATACGCCCCAG |
| Lct | CTTAAACTTGTACATCAACG |
| Dars2 | GTTGCTCGGACCAACACATG |
| Dars2 | TCTTCTAAGAAAATGCCGAC |
| Dars2 | TAGACAGCCAGAGTTCACTC |
| Dars2 | AAAGCAATATGTGTTCATGA |
| Ndufs2 | GCGGCGTATATCCGACCTGG |
| Ndufs2 | TAGCACTCACCATTCCAAGG |
| Ndufs2 | CCTCGGGCACAGTGGATCCG |
| Ndufs2 | CACATCGGGCTCCTGCACCG |
| Slc30a10 | CCATGACAACAGTCAAACTG |
| Slc30a10 | AGCCGTGATGACCACAACCA |
| Slc30a10 | GCACAGCAGTGACTCTCCGG |
| Slc30a10 | TGCTGCAGATGGTCCCCAAG |
| Hhat | CTGAGCCACGCAGAACGCGA |
| Hhat | GCACAGACCCTGGATTGTCA |
| Hhat | GACGGGATAATAGAAGACGT |
| Hhat | AGAGGGCATGCATGTACATG |
| Hibch | AGTCCATGGTCAATTCCGAG |
| Hibch | CAGATGTTGCAGAATAACGC |
| Hibch | CCTGCAACATCCTCAGCTGA |
| Hibch | CTATTGCAGACCTTTAGCTG |
| Ndufs1 | AATGGATCTCTGATAAAACC |
| Ndufs1 | TGTCCTATTTGTGACCAGGG |
| Ndufs1 | TAGAGGTTAGGGCCCCTACA |
| Ndufs1 | TGATGGTCAGTCTGTCATGG |
| Cps1 | TGAGCCTCACAATTTCGTCG |
| Cps1 | ATGCAGACCGAATCATCACA |
| Cps1 | TACAGTATTCCATGGAAGTG |
| Cps1 | GTTGGTGGCATCTCGTGTCG |
| Man1b1 | GAGCACCATTCGCATCTTAG |
| Man1b1 | GGTTACCAAAAGTTTGCTTG |
| Man1b1 | GTGGACTTCAGACAGCACGG |
| Man1b1 | ATGTTCATCAACACGAACAG |
| Coq4 | TCATCGTCCACAAAGCGTGT |
| Coq4 | GCGTACGGGGCCGAAGACTG |
| Coq4 | CCAGGAAGCGAAGATACTCA |
| Coq4 | CTCTACAACCCCTATCGCCA |
| Dolk | GGACCGATACTCCTGGTGCG |
| Dolk | CAGTGTAGGCGAGGTGATCG |
| Dolk | AAAGTATTTCCACTTCATTG |
| Dolk | CCAGCAAAGACCAATAGGCA |
| Agps | GTACCAATGAGTGCAAAGCG |
| Agps | GTAAAACCAAGCCACTAAGT |
| Agps | TCAGAGAAGGGATGTTTGAG |
| Agps | TCCCTGGAATTCAGCACCGT |
| Slc35c1 | AGGGCACCCCTACGTACTTG |
| Slc35c1 | AGGTTAGGCGCCAGATACTG |
| Slc35c1 | CTCACCAATGATGACGCCGC |
| Slc35c1 | GCAGTGAGGTCACCAGGCAT |
| Acad9 | TCTGCCCAAACTGTCGTCTG |
| Acad9 | ATTTCCCGACTGCTCTATCG |
| Acad9 | GGACTCTCGAAAAATTGACC |
| Acad9 | GAGCATACATCGTGTTGGAG |
| Gatb | GTTCATTTGATTATAAGCTG |
| Gatb | GTGACCTCAGGTCGTCATGG |
| Gatb | CTTCCCTAGATAGATACAGT |
| Gatb | AATATATCTGTACATCACCC |
| Msto1 | CTGTTCTGTACAAGACGTCG |
| Msto1 | GGACAGACTACACTTCTATG |
| Msto1 | GGGCGGGGAGTCCTAACCTG |
| Msto1 | AGCCCCCATACCCATCATGG |
| Ampd1 | AATGACAAATACAATCCCGT |
| Ampd1 | CAAAGCACGACTCACCCCCG |
| Ampd1 | CGAGATCTCCCCCTTCGACG |
| Ampd1 | ATGGATGTGAGTATCCACCT |
| Slc25a24 | ATGAGAAAAAATCAGGACAG |
| Slc25a24 | AGAGACTGGACAATTTCAGA |
| Slc25a24 | AAGAAGTTGCTTACCGAGGA |
| Slc25a24 | ATCTTTGTTGACATCGCCAG |
| Slc35a3 | GCTGTAGAAGACAGATAACG |
| Slc35a3 | TGAAGCTCGCTATCCCGTCA |
| Slc35a3 | TCCTGCCATCAGAATTACTA |
| Slc35a3 | CAAACTGTGAGCCTGTTGAA |
| Gba2 | CTCACCTCAAGCCCATACCG |
| Gba2 | CGTCATCCCTCATGACATTG |
| Gba2 | ACCAAGCTACTGCAATTCAG |
| Gba2 | CATCGTATGTGTTGTACATG |
| Aldob | TATCCACAGTTGGACCAAGG |
| Aldob | AATTCCATTAGCCAGAGCAT |
| Aldob | CCGCCTGCAAAGGATAAAGG |
| Aldob | GGTCCCTATTGTTGAGCCAG |
| Pigv | GAGGGCAAAAGGCATCTGCG |
| Pigv | GTAGGCACGATACTGAAAGA |
| Pigv | TTCCTGTTTATCGCTGAGCA |
| Pigv | CAGGAGAAAATGCACCCCCA |
| Slc30a2 | CTGGTCGGGAAGACACCCAG |
| Slc30a2 | TGGAGATTATGAGATCAAAG |
| Slc30a2 | AGGCCACATAGAGTTTGCGT |
| Slc30a2 | GATGGAAAGCACGGACAACA |
| Hadhb | ATATACAAGTCTTACCTCAG |
| Hadhb | CTGACAGCAGAAATGGAATG |
| Hadhb | GCATCGGACCAATATTCCAA |
| Hadhb | TCAAACCAAGCCATGACCAC |
| Slc26a1 | ACAGGCCAACATATGTGATG |
| Slc26a1 | CCAGCCGGAGGATACCCATG |
| Slc26a1 | TACCAGACCCTAAGATAATG |
| Slc26a1 | GCCCACATTAACATGGCGGG |
| Ap5z1 | TGCGTCCAGTACCTTCCATG |
| Ap5z1 | TAGGTCCACACAGACTCCGT |
| Ap5z1 | ATAGCCCGGTCCCATCACTG |
| Ap5z1 | GGCCGCAGACAGCACAGACA |
| Mat2a | CCTGATGCTAAAGTGGCTTG |
| Mat2a | GTGCAATATATGCAAGATCG |
| Mat2a | ACCAAGGCAATGTACCATTG |
| Mat2a | ATTTACCACCTACAGCCAAG |
| Cyp26b1 | ACTGCTGGGTCCCAACACGG |
| Cyp26b1 | GCACCGGTCACACGAATCAG |
| Cyp26b1 | ACTGCTGGTATACCTCAAAG |
| Cyp26b1 | GGCCATCAATGTATATCAGG |
| Slc6a1 | CACCAACATGACCAGCGCCG |
| Slc6a1 | GCAGAAATACACGAGCACCC |
| Slc6a1 | TACCTCTGTGGGAAAAACGG |
| Slc6a1 | TCCATGTGTCCCGGTCAGGG |
| Gys2 | GCATAAGAGTAACGTCACCG |
| Gys2 | GTGGATGCGATGAATAAACA |
| Gys2 | GAGATAACGCCCAAGCAGTG |
| Gys2 | AAAACGACAGCCGATGAGTG |
| Pgap2 | AAACGCCACACGTAGCGTTG |
| Pgap2 | CATTAACTTCAGTCTCAATG |
| Pgap2 | GTAGTGGTTCCAATAGGCGA |
| Pgap2 | GACAACATCTACACACTGTG |
| Xylt1 | CCTGTACGGGAACTATCCTG |
| Xylt1 | CATGCGCTGTAGCTGTCGAG |
| Xylt1 | GTGCCAGTACAAGCATATCG |
| Xylt1 | GATGCCCAGGACAAACGACA |
| Cog7 | GTGCTATCAGACATTACCAG |
| Cog7 | TGAACGATGCTACGATCTGG |
| Cog7 | GTGGCACACACAAACCCAGT |
| Cog7 | GGTGCTGGTAGAGATTGACC |
| Tufm | GAGGAAACTCCAGTCATCGT |
| Tufm | CCGGTAGTTCTAGCCGAGGG |
| Tufm | GTGACAGGTACATTAGAGCG |
| Tufm | TCCACCCAGAATATGATCAC |
| Csgalnact1 | TTATTGTGCCCCTAGCGAGG |
| Csgalnact1 | CAGGGATTTACCGAACCGAA |
| Csgalnact1 | AGGGCTGTTTAGACTCTCCA |
| Csgalnact1 | CTTGACACCGGCATGTACCT |
| Tat | ATGTTCGCGTCAATATTGGC |
| Tat | GGCAAGCTTACCGATAGATG |
| Tat | AACAACCCGTCCAATCCCTG |
| Tat | TGAGCTGTGTCTAGCCGTGT |
| Slc38a8 | TCACGGTGCAATACTACCTG |
| Slc38a8 | ATTTCCCGAAGTGCTGACAG |
| Slc38a8 | GCTGGTCCCCGATCACTCTG |
| Slc38a8 | TCTTTGGTCTTCCTGATCAG |
| Spg7 | TTCCAAATGCAGTCAGACTG |
| Spg7 | GAACTCCCTCAGTACAAGTG |
| Spg7 | ATCAATGTACACTATGCAAG |
| Spg7 | GGATTCCCGTGTCCTACAAG |
| Foxred1 | ACTCCTTCTACGTTTATCCA |
| Foxred1 | CCCAATGAAGACCCCAACTG |
| Foxred1 | TGGAGCAGGACCACACGGTG |
| Foxred1 | TTGAACTGAAGTTCCACAGG |
| Sc5d | CGTATGTATATCCAGCCACG |
| Sc5d | CATGACTTGCAAACGGCGTG |
| Sc5d | ATAGACAAAATAATAGCTGA |
| Sc5d | AAGTAGACCACCTTGTGCAG |
| Dlat | ACTACCGCAACGGACCGCAG |
| Dlat | CAGGCTCTCAAACCCAACAG |
| Dlat | CGACAAGGCCACCATAGGTG |
| Dlat | TTCAGAACCACACCTACCGG |
| Slc17a5 | TCGTCACCCAGATTCCCGGT |
| Slc17a5 | TAGAACGTCTAAGGAGTGTG |
| Slc17a5 | GATCGTTATCTTACCCGCAT |
| Slc17a5 | GCTGGCCGCAGACTTAGGCG |
| Glyctk | GGTGTGATCAGCGTACCCAA |
| Glyctk | GCGGGAAAACCTCTACCTAG |
| Glyctk | CTGTGCTCAACACCATGGCA |
| Glyctk | AGTGATTGCCAGTGGCCCTA |
| Phgdh | CCACACTGAGAGCCTACCTG |
| Phgdh | CTCACTTCTGACCAGACTGT |
| Phgdh | TAGTAGCAGACCGGACAATG |
| Phgdh | CTGCGATTTCCTCCCCACAG |
| Slc13a5 | GATAGCCGCCAATACAGCTG |
| Slc13a5 | GGGTCCCGTCCCGGTCAAGG |
| Slc13a5 | AGTCCAGAACCTTCAAAAGT |
| Slc13a5 | AAGAAGGTGTGTTTACCGTG |
| Apob | GCTGGTAGTGACATCAACAG |
| Apob | AGTCTCACCTGTTAACTGCG |
| Apob | ATGTTGCATGAGTATGCCAA |
| Apob | GAATGGAATCATGCCCATTG |
| Cog5 | GGAGACTGTTACCAGTGTCG |
| Cog5 | GGAGGATTACGTCCTCCCGG |
| Cog5 | CAACTAGCAAAACTTGCCCA |
| Cog5 | TTTCACAGCATATAACTGGA |
| Mtr | TCAGGTGCTCGATATCAACA |
| Mtr | GGGGTCCGAATGAGACACGC |
| Mtr | TGTGGATAGCATCATACACA |
| Mtr | CTTCAAAAACACTAGCAGGG |
| A4galt | TCTTCCTAGAGACATCGGAC |
| A4galt | GCTGATGAAAGGGCTGCCTA |
| A4galt | TCAGGGGCACCCCCAGACAG |
| A4galt | ACTGTTTGAGGACACACCAC |
| Pgap1 | ATAGACAACAATATGCGACA |
| Pgap1 | ATATGTAACTTACCACACGA |
| Pgap1 | AGAATTTGCTCCTACAAGTG |
| Pgap1 | GTCGTCAGACTGTCAGTATG |
| Impad1 | ATTCCTGTGTAGCATCAAGT |
| Impad1 | TCCTTCACGAGAAGTCCAAG |
| Impad1 | CTCCCACTAGATAAATACTG |
| Impad1 | GCTTGGGCAATGGTAGATGG |
| Gabbr2 | CCTGCGACTCTACGACACCG |
| Gabbr2 | CATGGAAGGCTACATCGGAG |
| Gabbr2 | GCTGACAGGATGCTATACAG |
| Gabbr2 | CTTATCCGCAAGAACAGGCG |
| Slc35d1 | GGCGCCGGCCGAAACGCTAA |
| Slc35d1 | TGTGTGTGGAGGATTTCGCG |
| Slc35d1 | ACTGTATTTGCAATGATCAT |
| Slc35d1 | TGGTTCCCAAAATATAGTAG |
| Cldn19 | CCACACGGCTCTTGGCAGTG |
| Cldn19 | CTGGGCACTCACCTGCCAAG |
| Cldn19 | TGCTGACTGGATATGACCTG |
| Cldn19 | ACATCCACAGCCCTTCGTAG |
| Slc45a1 | GAGTACCGGAGTCACGTACG |
| Slc45a1 | AGGAGTACTCACCCGCCATG |
| Slc45a1 | GGAGTGACCGATGTACCTCA |
| Slc45a1 | CCACCCCGCACACTGTCAGG |
| Ppm1k | CAGGTGGGCATAACTCGCGA |
| Ppm1k | GCAGAATTGGCTCATCAATG |
| Ppm1k | CTAGGATCAAGAAATTCGGT |
| Ppm1k | AAAATTAGCCTGGAGAACGT |
| Uroc1 | TGTCACAACCCGTTCAATGG |
| Uroc1 | TCTGCCCCAGCATTGAGATG |
| Uroc1 | GGGAATCGTCCATGGCACAG |
| Uroc1 | CACCCCAATGCATCCAACGA |
| Fkrp | GCTGGACTTGACCTTCGCCG |
| Fkrp | CCAATGGCAGAAGTCCGGCG |
| Fkrp | CCATCCAAAGCGTCGCAGCG |
| Fkrp | CACAAACTCGGTGGCTACGT |
| Nars2 | TCACATCCAACGACTGTGAG |
| Nars2 | GGAACTCTTCAAGGCGACCA |
| Nars2 | CTCAGGGTCCAGGACGCGAG |
| Nars2 | TCAGTCCAAGAGGCAAAATG |
| Cyp2r1 | TAGGCATCAACAAAATGGTG |
| Cyp2r1 | GCAGAGCCGGGTGTATGGCG |
| Cyp2r1 | GATGCTATTGAAACATACAA |
| Cyp2r1 | AAAGGTAAGATGCCAATCCA |
| Snx14 | AGCTGTGCTGCCTAATTACG |
| Snx14 | CTTCAAGAGTTATTTCCGCA |
| Snx14 | GACAGATGCAAAAAACCGTA |
| Snx14 | TCAACATCGATACAAAACAC |
| Slc36a2 | TGACTAGGATGTGCATACAG |
| Slc36a2 | GGTAGTTCTGACGCCCACCA |
| Slc36a2 | AGCAAAGTACCTCCCCCAGT |
| Slc36a2 | AGCCCCGATAGCAATGTACA |
| Fktn | GAGTCTATTCCATCTAGCCG |
| Fktn | CACTCGATAAATCTGGAGTG |
| Fktn | GACTCAAAGGACACATGGAT |
| Fktn | TAATTGATTCCAGAATCCAT |
| Atpaf2 | ACAATGAATTAGCTACAGGG |
| Atpaf2 | AATGGGATCCCGTCATAGAG |
| Atpaf2 | CTATGTCCCGCCAACAGGTG |
| Atpaf2 | TGGCCCCGGCCATACTGAGA |
| Slc5a2 | ATGATTTATACTGTGACAGG |
| Slc5a2 | CTGGCACAAAAAGCCATCCG |
| Slc5a2 | GGTCTCTTCGACAAATACCT |
| Slc5a2 | ATTGGTTCTGAACATAGACT |
| Acsf3 | CACAGCGGCTAGGTTACGGT |
| Acsf3 | CATGAATACGGTAATCTGTG |
| Acsf3 | GGCCCACTGTGCTACAACGT |
| Acsf3 | ATATGGCCATCACACCTACA |
| Gphn | GCTTTGTCCATAGACGTCAG |
| Gphn | TGAAGGAGTAGTGCTAAGGG |
| Gphn | ACCACGAGATGTCACTCCAG |
| Gphn | ATCGGCCATGACATTAAGAG |
| Sptlc1 | AATGTGCCATAGAACCCTCG |
| Sptlc1 | CCCTCCAACCCACAACATCG |
| Sptlc1 | TCCTGCGTACTCTAAGAGAG |
| Sptlc1 | TTTGTCGTAGAATCCTCGCA |
| Rnf31 | GATGGATTGAGTTTCCCCGA |
| Rnf31 | GAACTATGAGTTGTTGGACG |
| Rnf31 | CTACCTCAACACCCTATCCA |
| Rnf31 | GGAGGAACCAAGGTGTTGTG |
| Abat | GAGCAGAGGTAACTACCTAG |
| Abat | GCTCCAGGAGTCCTTGATGT |
| Abat | TTTCGGCAGAGTAAGGAACG |
| Abat | AACGGTGGCTGGAATCATCG |
| Ahcy | GCGCACCTGACAGAAGCTGT |
| Ahcy | TGTCAACGATTCTGTCACCA |
| Ahcy | TGACCCTATCATACCCTCCA |
| Ahcy | TGTGATGATTGCGGGCAAGG |
| Fbxl4 | AGTTGTGCTGCATGGTACGA |
| Fbxl4 | TTGAGGTGGATATATTGCAG |
| Fbxl4 | ACAGTATGTCCTATACCATG |
| Fbxl4 | AGTGGCTGCAGGATAACTCA |
| Vcp | ACTGTCTTCACAGACTCATG |
| Vcp | TATAGGTCGCTTTGACAGAG |
| Vcp | CCAATCGCCTTAAAGAGCGC |
| Vcp | CTTCAGGAGTTGGTTCAGGT |
| Chsy1 | GGTGTAAGGTGATCGCTCGG |
| Chsy1 | GCGTGTGAACCCCATGTACG |
| Chsy1 | AAGAGAAAGTTCCTGTCGCG |
| Chsy1 | CAAGTACGAGTGGTTTATGA |
| Gcdh | GATGATCATACCGGTAGCCG |
| Gcdh | CAGGATCACCAACTCCCCTG |
| Gcdh | GGTCCTTCCAGTCAAACACG |
| Gcdh | CTCCTGGCAGTAGTTACGGA |
| Clpx | AAGAATTCTCGAAAACTACG |
| Clpx | TGGTGATTTGTGTACACACG |
| Clpx | TCTCTTATAATGATTATACA |
| Clpx | TGTTATTTACCTGACCCAGT |
| Pigs | ACCATAATGCCACCCCAGCG |
| Pigs | GGGCGACCTGGACTATGCGA |
| Pigs | GATCTCTCGCTCATGCACAA |
| Pigs | CTACCTAAGCTGGACTTGAG |
| Flad1 | ACAAACTCGCGGAGTCAGGT |
| Flad1 | TCACTCACGTCCTCACCGCG |
| Flad1 | GCTAAGCCTACGCCCAAAGT |
| Flad1 | GAAGCTGATTCTAGACTCCG |
| Pik3r5 | CCACAGGAATCTCCTACCAG |
| Pik3r5 | CGAGAGGCCTGAGACAAAGG |
| Pik3r5 | TGTACTTACACAGAGGACAG |
| Pik3r5 | GCTGAGCGGGACAGTCCAAG |
| Alg6 | TTTGCTCATTTACGTACCCG |
| Alg6 | TCTCATAGCCTCGTGATGTG |
| Alg6 | TAGTCGATAAGAATAAGACC |
| Alg6 | GCTTCATAATCACCAAACAT |
| Ocrl | TTTCGAATCATATTTGTACG |
| Ocrl | CCAATGCAGTAAATATCAGG |
| Ocrl | ATTTACCCCAATATGGAACA |
| Ocrl | ACGTGGAAGAGTTCGAACGA |
| Pgap3 | CATGCAGGTATGGTACATGG |
| Pgap3 | AGTCGCGGTACACGGGCTCG |
| Pgap3 | ACTGCAAGTATGAGTGTATG |
| Pgap3 | GGGACTACAGCACCCGTCTG |
| Trappc11 | GCAGGCCGGGACTTACATCG |
| Trappc11 | AGAGCTTGGAAAACCTAATG |
| Trappc11 | CGAGTGGTACATCCCTAAAG |
| Trappc11 | AGAGGAGTACTACTATGCAA |
| Gfm2 | GGATACACAAGATCATTGGG |
| Gfm2 | ACCAGCAGAGGCGTCAAACA |
| Gfm2 | GGAGCATGGAAAGAAGAGAG |
| Gfm2 | CCAATTACCTGTAAAATCAG |
| Pisd | CAAACTTGCTAGTCACTGGG |
| Pisd | GGTCCCCAGAAAATGGCGTG |
| Pisd | GGGCCCTCGAGCCAATACAG |
| Pisd | CGTATGTGGCTTGCACTGTG |
| Rfx6 | TGAGCATAAAGAATGCACCG |
| Rfx6 | GGACATAGGATGTCTCCACG |
| Rfx6 | GGTAAGATTATGCATCACGC |
| Rfx6 | TCAGAAATGCAGTTAAACAA |
| Serac1 | AGCCATAGTAAAGCATTCGG |
| Serac1 | AACCTGGACCGAGAAACTGT |
| Serac1 | CACCCTGCTATAGTCCACTC |
| Serac1 | GAGCAGCATTTAAAACCTGG |
| Dna2 | ACACGATGCGAAGGATACGG |
| Dna2 | GCCCTTCAGTCCAAACCTAG |
| Dna2 | TGCCTTGCCACAGATAATCG |
| Dna2 | CAGGCGTTAACCAGATACTC |
| Pigl | CTTGTCAATAATCATTACAC |
| Pigl | TTGATTGTAGTAGTTTCCTG |
| Pigl | GTTCCGCTGAGTTCCAAACC |
| Pigl | GCGTGCCAAGCCTAGCATGG |
| Rft1 | AGGACCAAAATATACCCAGT |
| Rft1 | GAATAATCTTGGCTCCCTTG |
| Rft1 | GGTTCTTGCTGAGAGCATGT |
| Rft1 | GGCAAAGCCAAAAACAGTCA |
| Slc5a6 | CTTACTTAATCCCAGAGATG |
| Slc5a6 | CAAACATGACCAGACCAATG |
| Slc5a6 | GCAGTTCACCAACGGTATGG |
| Slc5a6 | ATCCTATAGGTGATATACAT |
| Gmppb | GTGGCTTCTCAACAAAACGG |
| Gmppb | GTGCCGATGAAACTGCACCA |
| Gmppb | TTCCCAGTTATGGCCAAGGA |
| Gmppb | GAGCACTCCGAAGCCATTGG |
| Cndp1 | CACTCACTTATGGAACCCGG |
| Cndp1 | AGTCAGCCATTGGTTCGTTG |
| Cndp1 | CAGCTCAGAAGGACGACGGG |
| Cndp1 | AGCCGGTTCCATCGCCCTGG |
| Fa2h | TTCGTGCGGCATCATCCGGG |
| Fa2h | TCTTGAGCCACAGTTCAAAG |
| Fa2h | CTACTACCGAACCCTCACCC |
| Fa2h | GTGGATGACGTATTCCACAA |
| Gars | AGTCAGCAAATTTGTCTACG |
| Gars | GAGATATTCCAACCTTCGTG |
| Gars | TAAGTTAAAAGGTACCGGAG |
| Gars | CCAACAGTGAGTACCTGACT |
| Lyrm4 | ACAGATCTAAAACTTGTGCG |
| Lyrm4 | CCAGGGCTTGAATTTCTACA |
| Lyrm4 | TCATTAGAATGTATGCTGTC |
| Lyrm4 | CCCGGGTCACCTGTAATTGT |
| Iars2 | CCAAACTAAAACGCTAACAG |
| Iars2 | AAATCGGCGTTTGTCCGCTG |
| Iars2 | AGGCGGCAGATACCGTGACA |
| Iars2 | AATCGATTCCATATGATGAG |
| Pdp1 | GAGGTCAACACCATCTACAT |
| Pdp1 | GTGTACCTCAGACGATTCTG |
| Pdp1 | ACTGGAACACCCAAAAAATG |
| Pdp1 | TGCTTGTGCCACTGAAGGAT |
| B3glct | TGAGGCAGAGTAGTCTACCG |
| B3glct | TGGAACGTGGTGACGCAGCG |
| B3glct | CATACAGCAAGAATTCAGCG |
| B3glct | TGCCAAAAGATTAAAGAGTG |
| Alg8 | TTGTCCGCGTTACTTCCCTG |
| Alg8 | ACAGCCTCCCAATATCTCAG |
| Alg8 | CCGCAGCAGATAGACACCAT |
| Alg8 | AGTTTGGAGCCCAGTAGGCA |
| Rrm2b | GAGAATGTACAAGCAAGCAC |
| Rrm2b | TTGGAAAGATGACGAACCGT |
| Rrm2b | GGAGATAAAATACTTCTCGT |
| Rrm2b | TCTCCTAGGGGAAAGAGTGG |
| Gatc | CAGATGCTCGATTACCGCGG |
| Gatc | GCCCCTAGAGTCGGTACTGG |
| Gatc | CAAAGCGAATCCACAGGGAA |
| Gatc | GGCGTGCAGCTGATCCGCAA |
| Ugt1a1 | AGACAAACTCTTGGGCACGT |
| Ugt1a1 | CAGTGGCAAGTGACCCATAG |
| Ugt1a1 | ACTTTGTGAAAGATTACCCC |
| Ugt1a1 | ACACTAACAGCCTCCCAGCG |
| Opa3 | ATGCGCATAATGGGTTTCCG |
| Opa3 | AGGTCTTGAAGAACTCGCTG |
| Opa3 | TCCTTGATACGGTTGGCCAG |
| Opa3 | TCAGCAGCGCAATAAGGAGG |
| Ndufs6 | GGGGTTCAAGTGTCGCCGAG |
| Ndufs6 | CGAAACCCCGTGCAGCCCTG |
| Ndufs6 | TGGGGAGAGTCAACAGCCGG |
| Ndufs6 | TGATGAAAAAGACTACAGGA |
| Gnptab | TGTTTGCAATGGACGCGTGG |
| Gnptab | TCTGCCCATGCCAATCGACG |
| Gnptab | AGGAGTGAAATATTTACCCG |
| Gnptab | CATGCTGGACCGTTTAAGGG |
| Pnpla1 | TTTGTACGATGTGCTACCGG |
| Pnpla1 | CTGCAGAGATACATCGACGG |
| Pnpla1 | GGTTTCCGAGTACCGATCCA |
| Pnpla1 | AAATGTTCAAACTCCAAACC |
| Pigg | AGGTGGATAAAAATGTCACA |
| Pigg | ACCCCAGCGTCACTCCCATG |
| Pigg | AAAGTACTCAGGCATTACCT |
| Pigg | GATTCGAGGCATAGTAACTG |
| Amt | AAATGGTGGCGTTTGCAGGG |
| Amt | TTGTAAGCAATACTTCTGAG |
| Amt | TGCGGAACTAAGGCCTAACC |
| Amt | TGTAACACCTGTGTTGCCGT |
| Cers3 | TATCATGAATAAACATCACG |
| Cers3 | GTTTAGAAAATGGTTCTGGT |
| Cers3 | ACACCTCTAGCAAATGCACT |
| Cers3 | ATGGGCATATGACCTCTGGG |
| Cyp26c1 | CCCTTGGACGTACCGTTCAG |
| Cyp26c1 | GTCACACACACTACTTGGCG |
| Cyp26c1 | GTGCGCCGCCCAACGACCGG |
| Cyp26c1 | CTGTCCCGTAGCGCTCGCGG |
| Lipt1 | GGTTGATGTTACCCATGTCG |
| Lipt1 | CTGTAAAATGAGCCCATGTG |
| Lipt1 | TGAAGTTCTGATGAGTGCGG |
| Lipt1 | AGTCCGGCCGATCTTTGATG |
| Gpx4 | CGTGTGCATCGTCACCAACG |
| Gpx4 | CATGCCCGATATGCTGAGTG |
| Gpx4 | TGGTCTGGCAGGCACCATGG |
| Gpx4 | TAAGCCAGCACTGCTGTGCG |
| Vps13b | TGTCCATACTACCCAAATCG |
| Vps13b | TAAACACTGCAATACAAGCG |
| Vps13b | GAAACCTCTTCCCGATACAG |
| Vps13b | TGGCAGTAGTCCATGTACTG |
| Pex10 | CAGAAGGACGAGTACTACCT |
| Pex10 | ACCTGGCCAAGAGACTAGCA |
| Pex10 | GTACGTTGGGATCATCCAAG |
| Pex10 | TGGAGGACGCAGCCGATCAG |
| Sco2 | TTCAGCCTACTAGACCACAA |
| Sco2 | CTGTTCCTTCTCAGCCCTCG |
| Sco2 | GCTCATCGGGGCAAATATCA |
| Sco2 | GCATAGAAATTCCCGACACT |
| Epg5 | ACAGCCGACTCGTTGTAACA |
| Epg5 | TCGAGCCAGAAGAACCAATG |
| Epg5 | TGGGTACCATACCCATATTG |
| Epg5 | GAAACGCTGTCTTACACAAG |
| Pet100 | GGAGATCCAGAACATAACCA |
| Pet100 | CTGTGGCCAAGAGAGAAGGA |
| Pet100 | CTTTGATACCCACCTTACGC |
| Pet100 | GGATCTCCAATCAGGCTGAG |
| Nt5c3 | AGAGTAGAAGAAATTATCTG |
| Nt5c3 | CTCACCACTCTACCATGTAA |
| Nt5c3 | CCAGCAGAAAATATGAACAC |
| Nt5c3 | TCTCACTTACTATGACACGT |
| Hprt | CTAGAATGATCAGTCAACGG |
| Hprt | TATACCTAATCATTATGCCG |
| Hprt | AACAAATCTAGGTCATAACC |
| Hprt | AGCCCCCCTTGAGCACACAG |
| Rnaset2a | CTCACCTTGCATACTGTTGG |
| Rnaset2a | AAACATGAGTGGGTTAAACA |
| Rnaset2a | AGAAGACCGGTGAATCACAT |
| Rnaset2a | CAGAAGATTGTAACCAGTCC |
| Oas1a | GGGAGGTACATTCCTCGATG |
| Oas1a | GTTGGTACCAGTGCTTGACC |
| Oas1a | AAAGACAGTGAGCAACTCTA |
| Oas1a | GAGGATCAGTTAAACCGACG |
| Car5a | ATTACAAGAAAGCCTCCGTG |
| Car5a | GACACTGGGCCAGTCCAGAG |
| Car5a | GGAGTTTGACGATTCCTGTG |
| Car5a | AGTGTTTACGAACCTCAGCT |
| Apoa1bp | TGTGGCCCCGGAAATAACGG |
| Apoa1bp | TAACGAGTATCAGTTCAGCG |
| Apoa1bp | CTCTTGGACATAGACGTGGG |
| Apoa1bp | GAGTCACTAGCCCAGTGAAG |
| Carkd | TAGGCCAGGTCCCACGACAA |
| Carkd | CTGACGTCGAAGAAGCACAA |
| Carkd | CATTCTCACCCCCAACCACG |
| Carkd | CCATACAGCACATGCGACAA |
| Ppat | ACCTTGGAATCGGACATACG |
| Ppat | ATAAGACGCCCGATGCAGAG |
| Ppat | TGATCACTCTGGGACTCGTG |
| Ppat | AGGGGTGTATGCGAGTAACT |
| Prosc | TATGCTGGAAACCGTAGACT |
| Prosc | ATGGTCCAGATTAACACCAG |
| Prosc | AGACCCACGAACTCCAGGCT |
| Prosc | TTGCTGACCGCAACGAGCCG |
| Ftl1 | GCTGCTCACCAGAGAGAGGT |
| Ftl1 | TTGGCCGAGGAGAAGCGCGA |
| Ftl1 | GAGTTTCAGAACGATCGCGG |
| Ftl1 | TCGTCAGAATTATTCCACCG |
| Sis | TCAGTCACGAATGACAACAG |
| Sis | AATTTACCGGGAGTAAAAGG |
| Sis | GGGAAGTAAAGTGTAGCGGA |
| Sis | GTAATACGGATGGAATCCAT |
| G6pdx | AGAGGTGGAAACTGACAACG |
| G6pdx | TGCCCGCTCACGACTCACAG |
| G6pdx | ATGACCCCACAGTACCCCAT |
| G6pdx | AGAGATGGTCCAGAATCTCA |
| Ins1 | TAGAGAGCCTCTACCAGGTG |
| Ins1 | CTGGGAGCCCAAACCCACCC |
| Ins1 | ACCCAAGTCCCGCCGTGAAG |
| Ins1 | GTGGAACAACTGGAGCTGGG |
| Gyg | AACAGCACAGGACTACTAGG |
| Gyg | CTTACGCTTTAAATGCCGGG |
| Gyg | CTGGTTATACGTTTCAATGG |
| Gyg | CACTAAAATATGTATTCAGT |
| Pcx | AGGCTGCCATCTCATACACG |
| Pcx | ACGAGCAGAGAGTCATAGTG |
| Pcx | GCGCATGGCAACGTCGAACG |
| Pcx | GACTGGGGCTCACATTGACA |
| Fh1 | AATTGGGCGAACTCACACGC |
| Fh1 | CAGAGCTTCAAACTTATTCG |
| Fh1 | GCGACGTTCGGAGCACACCG |
| Fh1 | CTCGTAGATTCTTGGCATGG |
| Atp5a1 | TGGTCAGAAGCGGTCCACTG |
| Atp5a1 | GCTCCCGCACAGAGATTCGG |
| Atp5a1 | ACTGGGCGTGTGTTAAGCAT |
| Atp5a1 | CCAACAGCTCCTCGCCAACG |
| Atp5d | TCAGGCGCGTACATACGCCG |
| Atp5d | GTGAGTTCGTACCTGCGTCG |
| Atp5d | CCAAAGGCTCCAGTCAGCGT |
| Atp5d | CTCACCAAAGTACTTAGTCG |
| Atp5e | GAGACACCATGGTGGCGTAC |
| Atp5e | GTGGCGTACTGGCGACAGGC |
| Atp5e | AGATCTGTGCAAAAGCAGTG |
| Atp5e | CTGGGAAAACCGGATGTAGC |
| Usmg5 | TTACATTCATTCTACCTGTG |
| Usmg5 | TGTGTCCTGGCCACATATGG |
| Usmg5 | ATGGCTGGTGCAGAAAGTGA |
| Usmg5 | GATGGCCAATTCCAGTTCAC |
| Peo1 | CATGTGCACGTAAATACTGG |
| Peo1 | CTAACCAGAACACAATCCGA |
| Peo1 | ACGTGGCCTGAAGCTACTAG |
| Peo1 | AGGGCGGTACGAAGAATACG |
| 2810006K23Rik | TCCTGATCAGTGGTAGAGGG |
| 2810006K23Rik | GTGCCACCAAACAAGATCTG |
| 2810006K23Rik | CGTGAAAGGACATGGCCCAG |
| 2810006K23Rik | TACCTTGACCACAATGCCGG |
| Timm8a1 | TTCTGAACAGGAGAAGTGCA |
| Timm8a1 | TCAGCCCGACTGTCCAACTT |
| Timm8a1 | GCCCAGAGCCGAGCCCGAAG |
| Timm8a1 | GTTGCAGCATTTCATCGAGG |
| Park2 | CCAAACAGATCACGTGACGG |
| Park2 | TACACATAGTACAGAGACCA |
| Park2 | AAGTGGTTGCTAAGCGACAG |
| Park2 | TTAATTCCAAACCGGATGAG |
| 2410015M20Rik | GGGGCCTAGTGACAAGAGTG |
| 2410015M20Rik | TGGAAATTTAATCTTTGGAG |
| 2410015M20Rik | CTCGAGTGTGGTCGCTAATG |
| 2410015M20Rik | CAATATGTGTGCCAGCAGAC |
| Clk1 | TTCGCAGCACCATTCACACG |
| Clk1 | ATACTTACAAAGTACTATGG |
| Clk1 | ACTACATGGGCTACGAGCCA |
| Clk1 | CTGATATGATCCATTCGAAA |
| Adck3 | AGTTCAGTTCTCAACACCAC |
| Adck3 | CATGCCACTGAAACAGATGA |
| Adck3 | CCAGCGAAGATCCTTCCACC |
| Adck3 | CATTGTGAGTACACTGTGCA |
| Adck4 | GACCTTATGTACAGTCCGGG |
| Adck4 | ACTAAGAAGTCCTTGCCAGG |
| Adck4 | ATGAGTGTGGGCCTGCCAGA |
| Adck4 | GCTAAAGGATGGGACTGAGG |
| Gyk | TCCATCTAGAGTTTAACAGG |
| Gyk | GACCTTGTCCCAGACTACTG |
| Gyk | TACAGCACCAGCTATAGCCA |
| Gyk | CTACCTATAGGTATGGAACA |
| Ept1 | CAATTCCAGCACCCCGTTAG |
| Ept1 | GTGTACTCCATCTTTGGACG |
| Ept1 | TGACTGGGTTTGGATTGTCG |
| Ept1 | GGTCGAAGTATGTCAGGAGT |
| Cyp4f39 | GGGGAAAGAATTATCTGACG |
| Cyp4f39 | CTTCAGGACTTACCTAACCA |
| Cyp4f39 | TCAGCTTGCGACAGGTAATG |
| Cyp4f39 | TGTACTACCTCACAGCCGAT |
| Cyp21a1 | GCTTACCTTGCATCCCCAAG |
| Cyp21a1 | TACCATTTAGCATATGGGGT |
| Cyp21a1 | AGGAGATGATACTACAAGTG |
| Cyp21a1 | ATGATTGACTACATGCTCCA |
| Akr1c21 | AGGACTAACCAAGTCCATTG |
| Akr1c21 | GTATCTCATTCATTACCCAA |
| Akr1c21 | AGATGATTCTGAATAAGCCT |
| Akr1c21 | TACTGAAGATCATGTAGGAG |
| Rab7 | GGAAGTTCTCGGGATCCCGG |
| Rab7 | ACGGTTCCAGTCTCTTGGTG |
| Rab7 | CGACAGACTTGTTACCATGC |
| Rab7 | CACATACTGGTTCATGAGAG |
| Tmem5 | TGCATAAATTAACCACTGGG |
| Tmem5 | GCGCCCGAAGAAGACGTGGT |
| Tmem5 | ATGTCTGCACCATTAATCCA |
| Tmem5 | TCTTTACAGTGACTTACCAA |
| Large | GATGTGTCTGATTTAAAGGT |
| Large | CAGCAATAGAATCAGCAATG |
| Large | GCAGGTAACAGCTCTGAGTG |
| Large | ACAGCAGAGAGGGAACTCAT |
| Chst5 | GTAAGCCTCTGTGCGCAACG |
| Chst5 | GCGAAGGCTTCGCAGACCGG |
| Chst5 | CAGCCATGTGCGCATCGCAG |
| Chst5 | TGAGCGGATCAGGTCACGCA |
| Hykk | CTCCGTATGCCGAACTAAAG |
| Hykk | CCTGAGCCAAAGTCGAAACC |
| Hykk | CCCAGGAAGACCAATAGCTG |
| Hykk | TTTCGTGTTCACATTGCAAG |
| Fdx1l | CGCCCGGGTCCTGTTACGGG |
| Fdx1l | GTTTGTCCCCAGGGTCAACG |
| Fdx1l | CTGGCAAGCGGATCCCGGTG |
| Fdx1l | TCTAGGACGTTCGGGACTAC |
| Zfp143 | GCATTTCGGTGCAAATACGA |
| Zfp143 | CTAATTATAAAAACCATGTG |
| Zfp143 | CCTGACGGAGGCAGTAACCG |
| Zfp143 | AGTGTGAGCATTCAGGCTGT |
| Cyp51 | GCTGTCAAAAGAACCAGCTG |
| Cyp51 | ATGCAGAAGAAGTCTACGGT |
| Cyp51 | TAGAAGTCAACTCAACGAGA |
| Cyp51 | AGGAACATACTGCTTAAAGT |
| Pigyl | GGGATGAGGACGGTCATGGT |
| Pigyl | AGGCCTGCCAGCGAGACCAG |
| Pigyl | TGGGAAGCCTTCCTCCACGG |
| Pigyl | CCACGGAGGCGGAGTAGAGC |
| Fmo3 | GATGCTACAATGATTTGTTC |
| Fmo3 | TTCACAGGACCATATAGAAG |
| Fmo3 | TGTCACAGCCTGAGTTCCCC |
| Fmo3 | CTTGGGTGATGAGTCGAGTC |
| Bckdhb | GCCAAGTATCGCTACCGCTC |
| Bckdhb | CTCGTAAACCAACAGTGCAT |
| Bckdhb | CCACACAACCCCACGGGGCC |
| Bckdhb | CACCGGTGCTACAGCTATTG |
| Agxt2 | CATCACCATGGCCAGGTCGT |
| Agxt2 | TAGCAAAGGCTTCCGGAAAT |
| Agxt2 | AGACAGAACTTGTATGCCAC |
| Agxt2 | AAAAACAGATAGACCGCCTG |
| Nnt | ACGAACGCTTCGTACTCACC |
| Nnt | GTTATACATACCTTCGCAGG |
| Nnt | TTCAATGTTGTCGTGGAATC |
| Nnt | GCTTCGTGAGCGCCTAACGT |
| Slc25a3 | TGCGGCACTTTACTAAGTCC |
| Slc25a3 | TTCAACAGTACGTTCAAAGC |
| Slc25a3 | GCCTTATATAGCAACATACT |
| Slc25a3 | GCCCCGAAGTGAATGTACAA |
| Ndufv1 | GCCGCCTATATCTACATCCG |
| Ndufv1 | TGTACCAGTCACCCCGTCTC |
| Ndufv1 | GCCAATCAGACCTGCTTCAT |
| Ndufv1 | CCCACAGGTAGCTATCCGAG |
| Ndufa12 | CTTAAAGGGCAAATGATATA |
| Ndufa12 | CTGGTGGGAGAAGACAAATA |
| Ndufa12 | TGGTGTAGATGACCCATCGG |
| Ndufa12 | CCTACCTGAAGAAAACCCGT |
| Ndufa12 | TCCACCCTTAGAGATTGTTG |
| Ndufa12 | AGCAATTCCCCGCTGCCTCA |
| Ndufa12 | CATACAGGTGGGATCTCATC |
| Ndufa12 | ATCTGCGGAAATCGCGTGTT |
| Bcs1l | TTGGTTATCCGCGCCGACGT |
| Bcs1l | ACGTCGGCGCGGATAACCAA |
| Bcs1l | TACCTTCTTTATGGGCCCCC |
| Bcs1l | AAGAAGGTAGCCACGTCTGT |
| Tymp | CACCAGGTGCCAATGATCAG |
| Tymp | ATACTTAGGGGCCATGCTAA |
| Tymp | TTCAGGTCACCTGTAATGAG |
| Tymp | TCACATCTCGTGCAGCATAC |
| Sacs | TTTGTAAACCTTCAATCAAA |
| Sacs | TCTCCAATCTTGATCCAATC |
| Sacs | ATATAAAAATTTAACCTCTG |
| Sacs | GTCCGCGTGACCGTGCTCCG |
| Tmlhe | AGTTGTTGCACCAACAAAGC |
| Tmlhe | TATCTAGTCACATGACCATC |
| Tmlhe | ATGCCGGTCTAGAGCTAGCT |
| Tmlhe | CTCCGTAATTTGATGAAGAG |
| Sbf2 | TGTTGGCTGTTTCCGCTCTC |
| Sbf2 | GGATGACACGCCTTTCCCAC |
| Sbf2 | CTTACCAATTCAATTCCCTG |
| Sbf2 | CTAATCTGGATACCAACACC |
| Pikfyve | CTGGCATCCAATATTGCTTC |
| Pikfyve | TGTTGTAATCAAGAAATCCC |
| Pikfyve | GGCTGGTTCTGCTTTCCTTC |
| Pikfyve | CGAAGCTGAACAGCTGTACG |
| Plcg2 | TTGCCAAACTGAGCGTGCTG |
| Plcg2 | CGCTCACTTTGAAGTTGACC |
| Plcg2 | TAGATCAAACCCGAAGAAAC |
| Plcg2 | AGATAAAGGAAATCCGTCCG |
| ApoE | GCCTCTGCGCGATCGCGCCC |
| ApoE | CCGACTCGGAGCCGACATGG |
| ApoE | CTACACAGGATGCCTAGCCG |
| ApoE | GCTTCTGGGATTACCTGCGC |
| Fdft1 | AACCTTGTCCCAGTCCTGTT |
| Fdft1 | CTCTCAGTGAACCGCCACTC |
| Fdft1 | CTGCCATCCCACACCCCATC |
| Fdft1 | CAGTACTGCCACTACGTTGC |
| Abcd1 | TCATCCTGCTTGAGCGCCTA |
| Abcd1 | GTTCCAGCATGACATACCAT |
| Abcd1 | CAGGGTGTACGAGATGTTCC |
| Abcd1 | TCTCTCTACAGGCCAAGTTG |
| Pex26 | GCGCTCTGCGGCCTTCAGCG |
| Pex26 | TCTGGATCCGGCGGAAGAGC |
| Pex26 | CTGGTAGTACCGGAGGACCC |
| Pex26 | GGTCAAACCGTAGAATCAAG |
| Tusc3 | AAAAGCAGATGAATAACGCC |
| Tusc3 | CATTCTGAAGATCGAGCGCC |
| Tusc3 | TAACGAAGAATATCAAATCC |
| Tusc3 | CGCTCGATCTTCAGAATGAA |
| Pigo | CAGGGGACTCATAGAACGCC |
| Pigo | TCCGGGGAAGAGGTCTCTCC |
| Pigo | CCCATCCCGTTTGGGAACAC |
| Pigo | CCACAGTGGACGGCGGTTCG |
| Atp6v0a2 | TGAAGAGCTCGAACGAATAC |
| Atp6v0a2 | CAACAGAGAGTCGTTCTCTA |
| Atp6v0a2 | CTACAGCTGCATGCAGCGGC |
| Atp6v0a2 | TCTTCGTGACCCTCAGCATG |
| Cers1 | GCAGACGACACCGCTTGGCC |
| Cers1 | TGACGTCAGCGATGTGCAGC |
| Cers1 | ATGAACTCACCGGAAGGCGT |
| Cers1 | GTAAGCGCAGTAGCTCCAAC |
| Cers2 | TTGGGAGGGATATCCCATAC |
| Cers2 | GGCGGAACCAGCGTTCTACC |
| Cers2 | CATTGGGAGGTGCCCGCAGT |
| Cers2 | GCCCACTCTGCCGTGACAAA |
| Ears2 | TGCTATCCGCTTCCGTCTAG |
| Ears2 | AGGGGCAGATGAGCAAATCG |
| Ears2 | TTCAGAAGCTCCAGCCGCTG |
| Ears2 | AGTGGCAGGTGATCCGCGTC |
| Hars2 | GGAACCTTGATGACAAAATT |
| Hars2 | GCCTCCAGACCTTTCCAACT |
| Hars2 | GAAGTTGTCTTCATACTTCT |
| Hars2 | AACAGCTGATAATCTTATCG |
| Stat2 | AGTTCTTGGTGAGATCCATC |
| Stat2 | ATAACTTGCGAAAATTCAGC |
| Stat2 | CCTACTTAGACCTTTCCCAA |
| Stat2 | TGAGATTGAAAATCGAATCC |
| BRDN0000737505 | AAAAAGTCCGCGATTACGTC |
| BRDN0000737693 | AAAACGGCTCGATCGGTGAT |
| BRDN0000737637 | AAAACGTAATTATACCGAGC |
| BRDN0000738185 | AAAATTGCACCTTCCCGGCC |
| BRDN0000737801 | AAACCCCCGCGCGGAGCGTC |
| BRDN0000737467 | AAACCTAGCGTAGATTCGGC |
| BRDN0000737848 | AAACGAGGCTGTTCGTACAC |
| BRDN0000737609 | AAACTCATACGTAGCGAATC |
| BRDN0000737434 | AAACTCCCGTGTCAACCGAT |
| BRDN0000738254 | AAAGACGTGCATTCAGCGAG |
| BRDN0000737777 | AACATGTTAAGTCGCGTTAT |
| BRDN0000737611 | AACCAGCATTTGACCGCGCT |
| BRDN0000737528 | AACCCCGGCTGTCATCGCCG |
| BRDN0000738228 | AACCCGCCGGAACAATCAGC |
| BRDN0000737727 | AACCGGCTGCGCGTTTGCAA |
| BRDN0000737483 | AACCGTACTGCGAGGAGCAT |
| BRDN0000737872 | AACCTCGTCTCATGTACGAA |
| BRDN0000737516 | AACGCCCCGGATTTCGTTGA |
| BRDN0000737844 | AACGGCTGCGCCCGCGGCAA |
| BRDN0000737412 | AACGGGCGCAATACCCTTTT |
| BRDN0000737631 | AACGGTAGCGTACCCGTGAA |
| BRDN0000737750 | AACGGTCAAATCCGTGAGGG |
| BRDN0000737875 | AACGTCACCAACCTCGATCC |
| BRDN0000738229 | AACGTTATAGCTTCGTCTCT |
| BRDN0000737806 | AACTAACTCACTACGCACGA |
| BRDN0000738366 | AACTCCTCATCGTACGCTAA |
| BRDN0000737593 | AACTCGCGTGGGAAGTCCGG |
| BRDN0000738128 | AACTTATACGTAATCTGATC |
| BRDN0000738307 | AAGACTCCTACGTATCGAGC |
| BRDN0000737391 | AAGCACAAGAACGGTCCGCC |
| BRDN0000737912 | AAGCAGCGACTACTCGACGC |
| BRDN0000738101 | AAGCCTAACGGAGCTCGCGG |
| BRDN0000738296 | AAGCCTACTTCACCGGTCGG |
| BRDN0000738095 | AAGCGAGCCGCAGACCGTTT |
| BRDN0000737714 | AAGCGTACCCCACTCGTTAA |
| BRDN0000738016 | AAGCGTGAGATTCACCGCCG |
| BRDN0000737416 | AAGGCCTTAACACGTCGACC |
| BRDN0000737993 | AAGGCGTAAACGAGTACACG |
| BRDN0000738351 | AAGGGTAAGTACAGTATCGT |
| BRDN0000738301 | AAGTCCCCTTCGGGAACTCC |
| BRDN0000737395 | AAGTCTATGCGGGGCTCGTA |
| BRDN0000737589 | AATAAGCCTACCCGGCGAGA |

**Supplemental Table 3. Overlap in depleted IEM genes in CD4^+^ T cells expanded in IL-2, restimulated with αCD3/αCD28, or cultured with 2-DG.** List of genes that were depleted when CD4^+^ T cells were expanded in IL-2 alone, after IL-2 mediated expansion and αCD3/αCD28 restimulation, or after IL-2 mediated expansion and treatment with glycolysis inhibitor 2-DG. Mutations in highlighted genes have been shown to cause IEIs.

| IL-2 alone & IL-2 + TCR Restim & IL-2 + 2-DG | IL-2 alone & IL-2 + TCR Restim | IL-2 alone & IL-2 + 2-DG | IL-2 + TCR Restim & IL-2 + 2-DG | IL-2 alone Only | IL-2 + TCR Restim Only | IL-2 + 2-DG Only |
| --- | --- | --- | --- | --- | --- | --- |
| Alpi | Atad3a | Cln3 | Ung | Aicda | Ak2 | Magt1 |
| Atp6ap1 | Slc46a1 | Aars2 | Aldh6a1 | Extl3 | Jagn1 | Abcd4 |
| Mthfd1 | Taz | Ddhd2 | Alg14 | Vps13b | Rbck1 | Acaca |
| Pgm3 | Rnf31 | Lbr | Cog8 | Abcc6 | Slc29a3 | Acad9 |
| Tfrc | Abcg8 | Ndufs3 | Dld | Acbd5 | Trnt1 | Acox1 |
| Rnaseh2a | Acsl4 | Ndufs8 | G6pdx | Afg3l2 | Acadvl | Atp5e |
| 2410015M20Rik | Ahcy | Ppat | Gfer | Aldh18a1 | Atg5 | Bcat2 |
| Adar | Akt2 | Stt3b | Hamp | Apoa1 | Carkd | Cox6b1 |
| Aldoa | Aprt | Tmprss6 | Isca2 | Aqp7 | Cldn10 | Cox7b |
| Alg1 | Asl | Uqcrb | Mrps34 | Asah1 | Cox20 | Eogt |
| Alg2 | Atp6v0a2 | Uqcrq | Ndufa13 | Bbox1 | Cox6a1 | Gatc |
| Atp5a1 | Cad |  | Ndufa6 | C1galt1c1 | Ears2 | Grin2a |
| Alg11 | Clpx |  | Ndufs1 | Cbs | Inpp5k | Mrpl12 |
| Atp6ap2 | Cog1 |  | Ndufs2 | Coa7 | Acy1 | Ndufa11 |
| Atp6v1a | Cog2 |  | Oas1a | Cth | Alg6 | Ndufa4 |
| Bscl2 | Cog7 |  | Slc25a19 | Cyp26c1 | Apoa5 | Ndufa9 |
| Cog4 | Dhfr |  | Sptlc2 | Dgke | Mlycd | Ndufaf4 |
| Cycs | Dpm2 |  | Stt3a | Dpm3 | Apoc2 | Ndufb11 |
| Atp5d | Elovl1 |  | Coa3 | Fut8 | Mpdu1 | Ndufb8 |
| Dhdds | Fdx1l |  |  | Gabra1 | Atp7a | Ndufs6 |
| Atp6v1e1 | Fth1 |  |  | Gad1 | B4galt1 | Ndufs7 |
| Dna2 | Fxn |  |  | Galns | Nars2 | Ndufv1 |
| Ftl1 | Gamt |  |  | Ganab | Cldn16 | Ndufv2 |
| Gars | Gne |  |  | Glul | Cnnm2 | Nfu1 |
| Gmppb | Kars |  |  | Gm2a | Nnt | Pmm2 |
| Cd320 | Ldlr |  |  | Gnptg | Cox15 | Pnpla1 |
| Gpx4 | Ldlrap1 |  |  | Gria3 | Cox4i2 | Rtn4ip1 |
| Isca1 | Lyrm4 |  |  | Gyk | Ogt | Slc25a3 |
| Lias | Msmo1 |  |  | Hadha | Dmgdh | Slc37a4 |
| Lonp1 | Mtr |  |  | Hexb | Elac2 | Timmdc1 |
| Mars2 | Mtrr |  |  | Lipt1 | Ggps1 | Uqcc2 |
| Mat2a | Mvd |  |  | Mboat7 | Gif |  |
| Mmachc | Npc1 |  |  | Mff | Hspd1 |  |
| Cyc1 | Pam16 |  |  | Mgme1 | Lars2 |  |
| Ndufb3 | Pgk1 |  |  | Naglu | Nadk2 |  |
| Ddost | Phgdh |  |  | Neu1 | Nhlrc1 |  |
| Ndufb9 | Pigm |  |  | Ngly1 | Pex11b |  |
| Ogdh | Pisd |  |  | Nr3c2 | Pex14 |  |
| Phykpl | Pmpca |  |  | Opa1 | Pex5 |  |
| Dpagt1 | Rft1 |  |  | Pdss1 | Pgap1 |  |
| Pik3r5 | Sec23b |  |  | Pet100 | Phka1 |  |
| Pnpt1 | Sgpl1 |  |  | Pex12 | Pigl |  |
| Rab7 | Slc17a5 |  |  | Pex6 | Pign |  |
| Rnaseh1 | Slc35a1 |  |  | Plcb3 | Pmpcb |  |
| Rnaset2a | Sptlc1 |  |  | Plcd1 | Scarb1 |  |
| Selenbp1 | Timm22 |  |  | Plin5 | Sdhb |  |
| Slc25a22 | Tmem199 |  |  | Pnpla2 | Serac1 |  |
| Slc33a1 | Ucp2 |  |  | Pomt1 | Suclg1 |  |
| Gfpt1 | Umps |  |  | Psat1 | Taco1 |  |
| Slc7a5 |  |  |  | Ptrh2 | Tars2 |  |
| Surf1 |  |  |  | Pycr1 | Tkt |  |
| Trappc11 |  |  |  | Rmnd1 | Tsfm |  |
| Ugt1a1 |  |  |  | Rrm2b | Tufm |  |
| Hcfc1 |  |  |  | Sbf1 | Urod |  |
| Hspa9 |  |  |  | Sc5d | Usp9x |  |
| Uqcrc2 |  |  |  | Sdha | Wars2 |  |
| Usmg5 |  |  |  | Sdhd | Xylt2 |  |
| Vcp |  |  |  | Sis | Yars2 |  |
| Iscu |  |  |  | Slc10a1 | Abcb7 |  |
| Yme1l1 |  |  |  | Slc25a20 | Aldob |  |
| Ldha |  |  |  | Slc25a26 | Alg13 |  |
| Ndufa1 |  |  |  | Slc29a1 | C1qbp |  |
| Ndufa2 |  |  |  | Slc30a9 | Cpt1a |  |
| Nfs1 |  |  |  | Slc36a2 | Dhodh |  |
| Nus1 |  |  |  | Sms | Ebp |  |
| Pcyt1a |  |  |  | Trip11 | Gck |  |
|  |  |  |  | Vps33a | Hsd11b2 |  |
|  |  |  |  | Zfp143 | Htra2 |  |
|  |  |  |  |  | Acadm |  |
|  |  |  |  |  | Ldhb |  |
|  |  |  |  |  | Ap1s1 |  |
|  |  |  |  |  | Apoa1bp |  |
|  |  |  |  |  | Mecr |  |
|  |  |  |  |  | Mfn2 |  |
|  |  |  |  |  | Mfsd2a |  |
|  |  |  |  |  | Mthfs |  |
|  |  |  |  |  | Mtmr2 |  |
|  |  |  |  |  | Blvra |  |
|  |  |  |  |  | Cant1 |  |
|  |  |  |  |  | Nans |  |
|  |  |  |  |  | Cep89 |  |
|  |  |  |  |  | Dpm1 |  |
|  |  |  |  |  | Eno3 |  |
|  |  |  |  |  | Etfb |  |
|  |  |  |  |  | Pex13 |  |
|  |  |  |  |  | Fbp1 |  |
|  |  |  |  |  | Pex26 |  |
|  |  |  |  |  | Fdps |  |
|  |  |  |  |  | Fdxr |  |
|  |  |  |  |  | Fkrp |  |
|  |  |  |  |  | Pgap2 |  |
|  |  |  |  |  | Gale |  |
|  |  |  |  |  | Ggcx |  |
|  |  |  |  |  | Pigg |  |
|  |  |  |  |  | Glb1 |  |
|  |  |  |  |  | Grin1 |  |
|  |  |  |  |  | Hadhb |  |
|  |  |  |  |  | Iars2 |  |
|  |  |  |  |  | Lfng |  |
|  |  |  |  |  | Mfsd8 |  |
|  |  |  |  |  | Mtpap |  |
|  |  |  |  |  | Neurod1 |  |
|  |  |  |  |  | Pah |  |
|  |  |  |  |  | Pcx |  |
|  |  |  |  |  | Pdhb |  |
|  |  |  |  |  | Pex10 |  |
|  |  |  |  |  | Pex7 |  |
|  |  |  |  |  | Pip5k1c |  |
|  |  |  |  |  | Porcn |  |
|  |  |  |  |  | Psap |  |
|  |  |  |  |  | Pycr2 |  |
|  |  |  |  |  | Rdh12 |  |
|  |  |  |  |  | Rpia |  |
|  |  |  |  |  | Sfxn4 |  |
|  |  |  |  |  | Slc35a2 |  |
|  |  |  |  |  | Smpd1 |  |
|  |  |  |  |  | Sugct |  |
|  |  |  |  |  | Thap11 |  |
|  |  |  |  |  | Timm50 |  |

**Supplemental Table 4. IEM genes and sgRNAs in the Targeted Inborn Errors of Metabolism Genes #1 Library.**

| **Gene Target** | **sgRNA Target Sequence** |
| --- | --- |
| Mat2a | CCTGATGCTAAAGTGGCTTG |
| Mat2a | GTGCAATATATGCAAGATCG |
| Mat2a | ACCAAGGCAATGTACCATTG |
| Mat2a | ATTTACCACCTACAGCCAAG |
| Tfrc | CTACACGCTTACAATAGCCC |
| Tfrc | GAATACATACACTCCTCGTG |
| Tfrc | GGGCTCCTACTACAACATAA |
| Tfrc | AACCCTCGGGAGACTCCACT |
| Atp6ap2 | TTAGCATATTAAGATCGCCA |
| Atp6ap2 | TGAACTTGGGAAGCGTTATG |
| Atp6ap2 | CCGGTGGAATAGGTTACCCA |
| Atp6ap2 | TGGACAGTGCAGCTACGTCT |
| Vcp | ACTGTCTTCACAGACTCATG |
| Vcp | TATAGGTCGCTTTGACAGAG |
| Vcp | CCAATCGCCTTAAAGAGCGC |
| Vcp | CTTCAGGAGTTGGTTCAGGT |
| Atp6v1a | TGACTGCTGATATCCGACAG |
| Atp6v1a | AGTCGGCCATCATACTGACG |
| Atp6v1a | ATGTTGCCCCCACGTAACAG |
| Atp6v1a | CTTACGGGAAAAGGGCATCG |
| Cog4 | GGAAACTCACCAACCCATCG |
| Cog4 | GCTGTGTACGAACGACTCTG |
| Cog4 | GCACAAGTAACGATGAATGT |
| Cog4 | CAACAGAAAAAATTGAACCA |
| Nus1 | GTGGTCGTAGACGCTAATGT |
| Nus1 | TCCAGGTGCCGAAGCGAACG |
| Nus1 | ACAGCACCTTCACTGCCGAA |
| Nus1 | CCAGCGCAGCCGAGGATGGG |
| Gmppb | GTGGCTTCTCAACAAAACGG |
| Gmppb | GTGCCGATGAAACTGCACCA |
| Gmppb | TTCCCAGTTATGGCCAAGGA |
| Gmppb | GAGCACTCCGAAGCCATTGG |
| Ogt | TTTGAGCCCAAATCATGCGG |
| Ogt | AGCATTATCGACATGCCTTG |
| Ogt | ACAGTGCACCACTAGTCCCA |
| Ogt | GAAAGTTTGTACCATCATCC |
| Gfpt1 | TGTGGCACAAGTTACCACGC |
| Gfpt1 | AAGCTGCGGTCTTTCCCGTG |
| Gfpt1 | GGAGAGAGGAGCCTTAACTG |
| Gfpt1 | TCTGTTGTGAACACAATGAG |
| Slc7a5 | GCCCTCCTCGCAGTACATCG |
| Slc7a5 | ACCCCTACTTACGCACGCAG |
| Slc7a5 | AGCGGCCTCTTCGCCTACGG |
| Slc7a5 | GTAGCAGAGTGCGCCCACGA |
| Dhdds | GGATGTCGGGATGAGGAGAG |
| Dhdds | CTATGCCAAGAAGTGTCAGG |
| Dhdds | CTTCAAACGTTCCAAGAGTG |
| Dhdds | GCAGCAGATGCAGATCACCC |
| Alg11 | GGAAACCAGCAAATGCCCTG |
| Alg11 | TTTCCTGGACTGATGATATG |
| Alg11 | AGTGAACATAGCTTCCGACT |
| Alg11 | TAAACCATATCCTCTCACTG |
| Pgm3 | TACGGCCTCACATAACCCTG |
| Pgm3 | AACTTCTTCAAGGTACCGCG |
| Pgm3 | TACACCATGTAGTGCAACTG |
| Pgm3 | CTGCTATGACATACCCTGTG |
| Ddost | GAACTCCCCGTACTTAATGA |
| Ddost | CCCAGATAAACCAATCACCC |
| Ddost | AGGCAACTATGAACTAGCTG |
| Ddost | GGTCAGAAACATCATAGTTG |
| Rnf31 | GATGGATTGAGTTTCCCCGA |
| Rnf31 | GAACTATGAGTTGTTGGACG |
| Rnf31 | CTACCTCAACACCCTATCCA |
| Rnf31 | GGAGGAACCAAGGTGTTGTG |
| Ogdh | TTGGCCCACTCATAGATACG |
| Ogdh | GACTAGTTCGAACTATGTGG |
| Ogdh | GTAAGTGGAAGACCTTGTCA |
| Ogdh | AAAGCTGAACAGTTCTACTG |
| Slc35a2 | TCACCCGCTGTAGTGGACCC |
| Slc35a2 | CTGCTCTTCGCACAAAAGAG |
| Slc35a2 | CTGCAAGGTATAGATGAGAG |
| Slc35a2 | GGCCACTGGATCAGAACCCG |
| Pisd | CAAACTTGCTAGTCACTGGG |
| Pisd | GGTCCCCAGAAAATGGCGTG |
| Pisd | GGGCCCTCGAGCCAATACAG |
| Pisd | CGTATGTGGCTTGCACTGTG |
| Umps | GAGCAGATAACTGTCGCCAG |
| Umps | CCGCAGGTCGATGTAGACTG |
| Umps | AGAGCGTGCACACGGCGTGG |
| Umps | TCTGTCTGCCGATGTGTCGG |
| Gpx4 | CGTGTGCATCGTCACCAACG |
| Gpx4 | CATGCCCGATATGCTGAGTG |
| Gpx4 | TGGTCTGGCAGGCACCATGG |
| Gpx4 | TAAGCCAGCACTGCTGTGCG |
| Gale | ATCTTCACCGATGCGCCCAG |
| Gale | AGAACTTGGACTTGCCGTAG |
| Gale | TTAACTCTATAGTAGTCCAG |
| Gale | CTGGGGGTTCCCGTACACGG |
| Alg13 | GATCTTGTCATTAGCCACGC |
| Alg13 | GTGAATGACTCAGTACGGAA |
| Alg13 | GGTTGTAACCCAGACTCTCG |
| Alg13 | TTTCGACGAGCTCGTCGCAC |
| Hcfc1 | GGTACCCTTCACAACCAATG |
| Hcfc1 | ACAGCAGTATGTGACTCCCG |
| Hcfc1 | TGGAAGGGCTAGTTGCACAA |
| Hcfc1 | CCCAAGGTTAGCCACAACAG |
| Slc25a22 | GTGTCTTAGCCAGGTCGATG |
| Slc25a22 | ATACATGCCGAAGTAGCCCT |
| Slc25a22 | TGATTCAGGTTGGCAAACAG |
| Slc25a22 | GACACCAGCTCTCTAAGGAT |
| Pten | CCTCCAATTCAGGACCCACG |
| Pten | TGTGCATATTTATTGCATCG |
| Pten | ACTATTCCAATGTTCAGTGG |
| Pten | GGTTTGATAAGTTCTAGCTG |
| Asah1 | GCAAGGTGTACGTTACCTAG |
| Asah1 | TAACATTTATAACATACCGC |
| Asah1 | AGTGATAAACCCTACCCACT |
| Asah1 | GAATATAAATAATAACACTT |
| Mmadhc | TCTGTGAGGCAGTCCCATTG |
| Mmadhc | GTGTGGCCTGATGAAACTAT |
| Mmadhc | TCACTAAAGAACAAAATCCT |
| Mmadhc | TATGTAAATGAGTTTCAGGT |
| Agps | GTACCAATGAGTGCAAAGCG |
| Agps | GTAAAACCAAGCCACTAAGT |
| Agps | TCAGAGAAGGGATGTTTGAG |
| Agps | TCCCTGGAATTCAGCACCGT |
| Pex2 | GTATGCTGTGTGCACCATTG |
| Pex2 | TCCCAAAAGACGCTAAATGA |
| Pex2 | GTTCTGGGGCTTGCAAAATA |
| Pex2 | ATGAAAGCACTGAGTAAACT |
| Mtm1 | AGGGAACCACCAAAAGAGCA |
| Mtm1 | TTACCACTCATACCGACAAG |
| Mtm1 | ATGGGAGGCGCGACAAGTAG |
| Mtm1 | TCTGACCGGTGCCATTCAAG |
| Esr2 | TGGCGCTTGGACTAGTAACA |
| Esr2 | TTCGTGACGGCTCTCTACAT |
| Esr2 | GAAGTAGGAATGGTCAAGTG |
| Esr2 | CGGCTCACTAGCACATTGGG |
| Arsb | GGTGGGCAGACTAGGTCTGG |
| Arsb | GAATGTCTGCCGACACGCCG |
| Arsb | AGCACAGACGTATTTATGCA |
| Arsb | GCCAGCACGAAGACCACATG |
| Piga | TCTTCCACGCCAAGACAATG |
| Piga | ATGGGTGCAGGTCCTATCGT |
| Piga | CAGACTGTGAAAGAGAGTCG |
| Piga | CCATGCTTATGGAAATCGAA |
| Coa5 | CCGGTATTATGAGGACAAGC |
| Coa5 | ATCCGCTACCTGGAGCACAC |
| Coa5 | GCGGGCGTGAAGGAGGATCT |
| Coa5 | GCCGGAGGGCGGCGCGTGTG |
| Adar | ACTCCAACAAGCCGCCTACG |
| Adar | AGAGGTAACCCCAGTAACAG |
| Adar | TTCTTGTAGGGTGAACACCG |
| Adar | TGTATCCAGGAATTCCCTAG |
| Pus1 | CCTGCAAAGAGGGTCAAGGG |
| Pus1 | CTTACCCAGAATCCGAATGT |
| Pus1 | TACTCGGGCAAGGGCTACCA |
| Pus1 | TCCAGGCCCTCACGTACGAA |
| Dse | GTCAAAGTATAAGCATGACC |
| Dse | CAGCACACAGAACATTGCCA |
| Dse | AGTTTCATACATATAGCCCG |
| Dse | TAATGAACGGCACACCATTG |
| Nr3c1 | CATTATGGGGTGCTGACGTG |
| Nr3c1 | AGCTTGCCTGGCAATAAACC |
| Nr3c1 | AAAGCCGTTTCACTGTCCAT |
| Nr3c1 | AAAACTGGAATAGGTGCCAA |
| Ext2 | TGTTTCGATGTCTACCGCTG |
| Ext2 | CTGGAGGACTCAATGGAGTG |
| Ext2 | TCCTGCAGAACATCCCACAG |
| Ext2 | AGGGCAGTGTTGTAATCTGG |
| Pigc | TGTACTGACAGACTCCGTCA |
| Pigc | GAATAAAACATACCCAACCA |
| Pigc | TCCGGAAAAACATCTATGCC |
| Pigc | GGCCGACCTGAAGAGTACTC |
| Ehhadh | TTCATATGGATGCTTCACGG |
| Ehhadh | CCACATCATGAGGTTACTAG |
| Ehhadh | GTAAACCCATAGAACCCCGC |
| Ehhadh | TCACTATGGCTCTAACCGTA |
| Mogs | TCTAGGTCATTCTTCCCACG |
| Mogs | TCGGCAGCATATCCACGATG |
| Mogs | TGCCGAAATAGACGTGTGGG |
| Mogs | GAGGTCCTACTACCAGAGAT |
| Slc6a9 | ATACCTCTGCTATCGCAACG |
| Slc6a9 | GTAGTACATGATACCCGTGA |
| Slc6a9 | ATGGTGGTGTCCACATACAT |
| Slc6a9 | TGTGCTACCAGCGTCTACGC |
| Slc39a8 | TCAGCTGCTGTAAGATCGCG |
| Slc39a8 | AGGGGGTTAAAATCAATCCC |
| Slc39a8 | CGTTAGGCTCAGTGACAGCG |
| Slc39a8 | CGGCGCCAACCGGAGCCTGT |
| Mmaa | CGACCTACTCGAAATGTGAG |
| Mmaa | AGGACAGAGAGCTTGCCTAG |
| Mmaa | TCAGGCCCTCTCCTACCAGT |
| Mmaa | ACTTGAGTGAAGGAGCCATG |
| Lipe | GAGTATGTCACGCTACACAA |
| Lipe | CACTTAGAGAGTACGCTCAG |
| Lipe | TGCGGTTAGAAGCCACATAG |
| Lipe | AGAGCGGATATGCCTTGCAG |
| Hprt | CTAGAATGATCAGTCAACGG |
| Hprt | TATACCTAATCATTATGCCG |
| Hprt | AACAAATCTAGGTCATAACC |
| Hprt | AGCCCCCCTTGAGCACACAG |
| BRDN0000737505 | AAAAAGTCCGCGATTACGTC |
| BRDN0000737693 | AAAACGGCTCGATCGGTGAT |
| BRDN0000737637 | AAAACGTAATTATACCGAGC |
| BRDN0000738185 | AAAATTGCACCTTCCCGGCC |
| BRDN0000737801 | AAACCCCCGCGCGGAGCGTC |
| BRDN0000737467 | AAACCTAGCGTAGATTCGGC |
| BRDN0000737848 | AAACGAGGCTGTTCGTACAC |
| BRDN0000737609 | AAACTCATACGTAGCGAATC |
